# Improving patient survival by direct targeting of chimeric protein-protein interaction networks

**DOI:** 10.1101/2022.03.24.485646

**Authors:** Somnath Tagore, Milana Frenkel-Morgenstern

## Abstract

Protein fusions produced by the “slippage” of two genes or by chromosomal translocations are essential diagnostic biomarkers of cancer. Fusions produce novel protein-protein interactions, which eliminate other interactions by changing protein domains in such fusions. The impact of these changes disseminates along protein-protein interaction networks, thereby altering cancer-promoting activity and creating cancer phenotypes. Currently, for most patients, the determination of appropriate drugs is totally empirical. As such, a personalized therapy approach based on unique patient genomic markers is needed. In this study, we considered 672 aliquot IDs containing 3091 fusions from The Cancer Genome Atlas (TCGA), accounting for 25 cancer sub-types assigned as leukemias, lymphomas, sarcomas, melanoma, glioblastoma and carcinomas. Protein-protein interaction maps showed distinct patterns according to cancer sub-type, reflected as different phenotypic traits. We induced site-directed percolations, i.e., critical transitions, by selective knockouts of genes encoding proteins in a given interaction network so as to identify breakdown points. The number of genes that needed to be knocked out in a site-directed manner before inducing a breakdown ranged from 1-7 based on whether the fusion protein was part of a network hub. Quantitatively, in leukemia, lymphoma, melanoma and glioblastoma, breakdown was achieved in 218 fusion networks when only higher degree hubs were percolated, and in 280 networks when a combination of higher and lower degree hubs was targeted, with an FDR < 0.05. These were subsequently addressed in survival studies. We found that patient survival may be improved by considering ‘breakdown points’ as drug targets.

## Introduction

Gene fusions have been recognized as important diagnostic and prognostic biomarkers in malignant hematological disorders and childhood sarcomas^1^. Recently, the biological and clinical impacts of fusions in solid tumors have also been shown^1^. Fusions in cancer are usually produced by chromosomal translocations so as to incorporate parts of two different parental proteins^2^. Arguably, the best-known example is the BCR-ABL fusion, an oncogenic fusion protein considered to be the primary driver of chronic myelogenous leukemia^3^, in addition to BCAS3-BCAS4 in breast cancer^4^ and EWSR1-ETV4 in Ewing sarcoma^5^. To understand the complex activities and dynamics of fusions, knowledge of their protein-protein interactions (PPIs) is essential. Such insight will also aid in the study of different cancer phenotypes. Likewise, these interconnectivities imply that the impact of a specific genetic abnormality is not restricted to a specific protein but rather disseminates along the links of a PPI network, thereby altering the activities of gene products that are otherwise normal^6^.

Using our customized Chimeric Protein-Protein Interaction Server (ChiPPI)^7^, we identified PPIs for fusions, as well as for their parental genes for 25 cancer sub-types. ChiPPI uses a score based on a domain-domain co-occurrence matrix for identifying PPIs. We classified the ∼11,000 known fusion transcripts in cancer^7^ into three groups, namely, leukemia and lymphomas (LL), sarcoma (SC), and solid tumors (ST), together with others. For initial training, we selected 150 fusions from the LL, SC and ST groups (50 each) and their 300 parental proteins (100 each from the LL, SC and ST groups). For the final study, we selected 672 unique aliquot IDs from The Cancer Genome Atlas (TCGA), with 3,091 unique fusions belonging to 6 broad categories, namely, Leukemia (LK), Lymphoma (LY), Sarcoma (SC), Carcinoma (CA), Melanoma (ME) and Glioblastoma (GL), comprising 25 different cancer sub-types (see online Methods).

Identification and validation of PPIs using various experimental and computational approaches have long been possible. In terms of physical interaction predictions, available methods are typified by the approaches of Deng et al.^8^ and Jonsson et al.^9^. In the present work, we used a percolation-based model for understanding the dynamics of fusion PPIs in characterizing the LK, LY, SC, CA, ME and GL cancer phenotypes. Percolation, a popular cooperative phenomenon in physical systems, describes the dynamical properties of various complex networks, by which the system undergoes a critical transition in size or function during its development^*10-12*^. An example of percolation is the epidemic spreading of infectious diseases over a network of towns^13^.

We implemented site-directed and bootstrap percolation by selective knockout of vertices (proteins) or edges (interactions) from the graphed PPI network^14^. A key challenge in such percolations is interpreting the behavior of a graph after performing random or targeted knockouts of edges^15^. Recently, network propagation methods have been used for network mapping of cancer mutations^16-20^. We used power-law modeling to categorize the PPI data into scale-free, hierarchical or random, and identified breakdown points ^15,21,22^, which are instances when knockouts drastically affect the overall functioning of the PPI network^6^. These instances could be specific time points at which knocking down a particular protein may affect its downstream targets. Accordingly, a balance needs to be maintained against perturbations while preserving adaptability in the presence of changes, a property known as robustness^23^. For computational studies of local and global properties, we implemented the local hypothesis and preferential attachment model of Barabási et al.^23^. Recent network-based, pan-cancer analysis studies showed that the topological location of somatic mutations in a PPI network might be closely associated with clinical outcome^24^. To understand the significance of breakdown points, we used gene expression data from TCGA and performed gene set enrichment analysis (GSEA). Overall survival (OS) and event-free survival (EFS) curves were constructed using the Kaplan–Meier method. A log–rank test was used to estimate differences between patients. Lastly, we predicted drug target sites that correspond to breakdown point networks to analyze whether they play a role in gene essentiality. We also addressed a case in which we used a percolation-based model to understand the role of the drug imatinib on a BCR-ABL1 fusion (in both the Philadelphia positive and negative chromosomes), by studying its downstream pathways.

## RESULTS

### Datasets

For the training part of the study, we randomly selected 150 fusions and their parental proteins from the ChiTaRS-3.1 database^7^. We predicted the corresponding interactors for each fusion using ChiPPI^29^ (Supplementary Table S1). For testing, we considered 672 unique aliquot IDs and their corresponding 3,091 unique fusions from TCGA, for 25 cancer sub-types (Supplementary Fig. S1b) (Synapse ID: syn3107127). We classified these cancer sub-types into 6 broad cancer categories, namely, leukemia (LK) consisting of chronic lymphocytic leukemia – ES (CLLE-ES); lymphoma (LY) consisting of lymphoid neoplasm diffuse large B-cell lymphoma – US (DLBC-US), and malignant lymphoma – DE (MALY-DE); sarcoma (SA) consisting of sarcoma – US (SARC-US); carcinoma (CA) consisting of bladder urothelial cancer – US (BLCA-US), breast cancer – US (BRCA-US), cervical squamous cell carcinoma – US (CESC-US), colon adenocarcinoma – US (COAD-US), head and neck squamous cell carcinoma – US (HNSC-US), kidney chromophobe – US (KICH-US), kidney renal clear cell carcinoma – US (KIRC-US), kidney renal papillary cell carcinoma – US (KIRP), liver hepatocellular carcinoma – US (LIHC-US), liver cancer – JP (LIRI-JP), lung squamous cell carcinoma – US (LUSC-US), ovarian cancer – AU (OV-AU), pancreatic cancer endocrine neoplasms – AU (PACA-AU), prostate adenocarcinoma – US (PRAD-US), renal cell cancer - EU/FR (RECA-EU), gastric adenocarcinoma – US (STAD-US), head and neck thyroid carcinoma – US (THCA-US), and uterine corpus endometrial carcinoma – US (UCEC); melanoma (ME), consisting of skin cutaneous melanoma – US (SKCM-US); and glioblastoma (GL) consisting of brain glioblastoma multiforme – US (GBM-US), and brain lower grade glioma – US (LGG-US). We also used the TCGA gene expression files for these aliquot IDs (syn5553991, syn5871617). The gene expression files were used for GSEA. Pathway enrichment and survival analysis were performed for the corresponding breakdown points for each aliquot ID (and cancer sub-type) that we identified using percolation.

### Power-law models of parental and fusion proteins follow Poisson distribution

For training, we generated power-law models for 300 parental and 150 fusion proteins in the LL, SC and ST groups. A higher proportion of PPI networks of parental and fusion proteins belonged to a random category in SC than in ST. This may be due to less connectivity among vertices in SC, as compared to LL, and to the lesser presence of communities and hubs, which are sources of major interactions. Supplementary Fig. S1a depicts an overview of the network categorization for LL, SC and ST fusions. These results indicate that most parental and fusion proteins of LL belong to the scale-free category (118), followed by the hierarchical (24) and random (8) categories (Supplementary Table S2).

In SC, more PPI networks were categorized as random than in LL (30 networks), and almost the same proportion was categorized as hierarchical (31), although a much lower proportion was categorized as scale-free (89) (Supplementary Table S3). Lastly, in ST (Supplementary Table S4), the observations were dramatically different for the scale-free (68), hierarchical (34) and random (48) categories. Moreover, 18 cancer sub-types belonged to CA, two to GL, two to LY, one to ME and LK, and one to SC. The numbers of unique aliquot IDs and fusions, and the average number of fusions per cancer sub-type are represented in Supplementary Fig. S1c. Similarly, Supplementary Fig. S1b illustrates network categorizations for LK, LY, SC, CA, ME and GL fusions, which clearly display Poisson distribution. The results indicate that those fusions are mostly drivers in LK, LY, ME and GL but not in CA. A possible explanation for this observation is that the majority of proteins have fewer interactions in CA, due to the function of fusion PPIs, which depend on ubiquitous rather than central hubs. For example, in prostate cancer, most fusion proteins act as passenger mutations^25^.

### Site-directed percolation and breakdown analysis reveal that higher degree PPI score hubs are soft spots for target-based interaction studies

We next considered higher degree hubs of all PPI networks in LK, LY, SC, CA, ME and GL to determine whether they are drivers of protein functions with fewer interactions. A greater proportion of higher degree hubs was observed in communities in SC, than in LK, LY, ME and GL, where the overall distribution was even. Furthermore, due to the lower average degree of hubs in CA, the proportion of PPI networks in the random category increased. In LK, LY, ME and GL, higher degree hubs ranged from 315 (FUS in a FUS-ERG fusion) to 23 (CBFB in CBFB-MYH11). In *SC*, these ranged from 315 (FUS in FUS-CREB3L1) to 4 (SYT4 in SSX1-SYT4), while and in CA, these ranged from 134 (PRMT1 in BCL2L12-PRMT1) to 1 (EPHA6 in EPHA6-CNTN6). The higher cutoff for higher-degree hubs decreased drastically in CA (Supplementary Table S5), whereas the lower cutoff decreased both in SC (Supplementary Table S6) and CA (Supplementary Table S7). In LK, ME and SC, the overall distributions of higher degree hubs and fusion interactors were considerably greater than in CA (Supplementary Fig. S2b-c). Similarly, the number of ubiquitous hubs increased more in CA than in LK, LY, ME, GL and SA. The lowest number of ubiquitous hubs was observed in CLLE-ES (LK), while the highest number was in RECA-EU (CA). These results indicate that a higher distribution of ubiquitous hubs are found in CA than in other cancers. Supplementary Fig. S2a-d presents the overall distribution of higher and lower degree hubs in LK, LY, ME and GL, SC and CA. For the majority of fusions, breakdown was achieved with a combination of lower and higher degree hubs. For example, in LK, LY, ME and GL, breakdown was achieved in 218 fusion networks when only higher degree hubs were percolated, and in 280 networks when a combination of higher and lower degree hubs was targeted (Supplementary Table S8-S10). In LK and ME, the overall distribution of breakdown points was considerably higher than in CA (Fig. 1).

**Figure 1.**
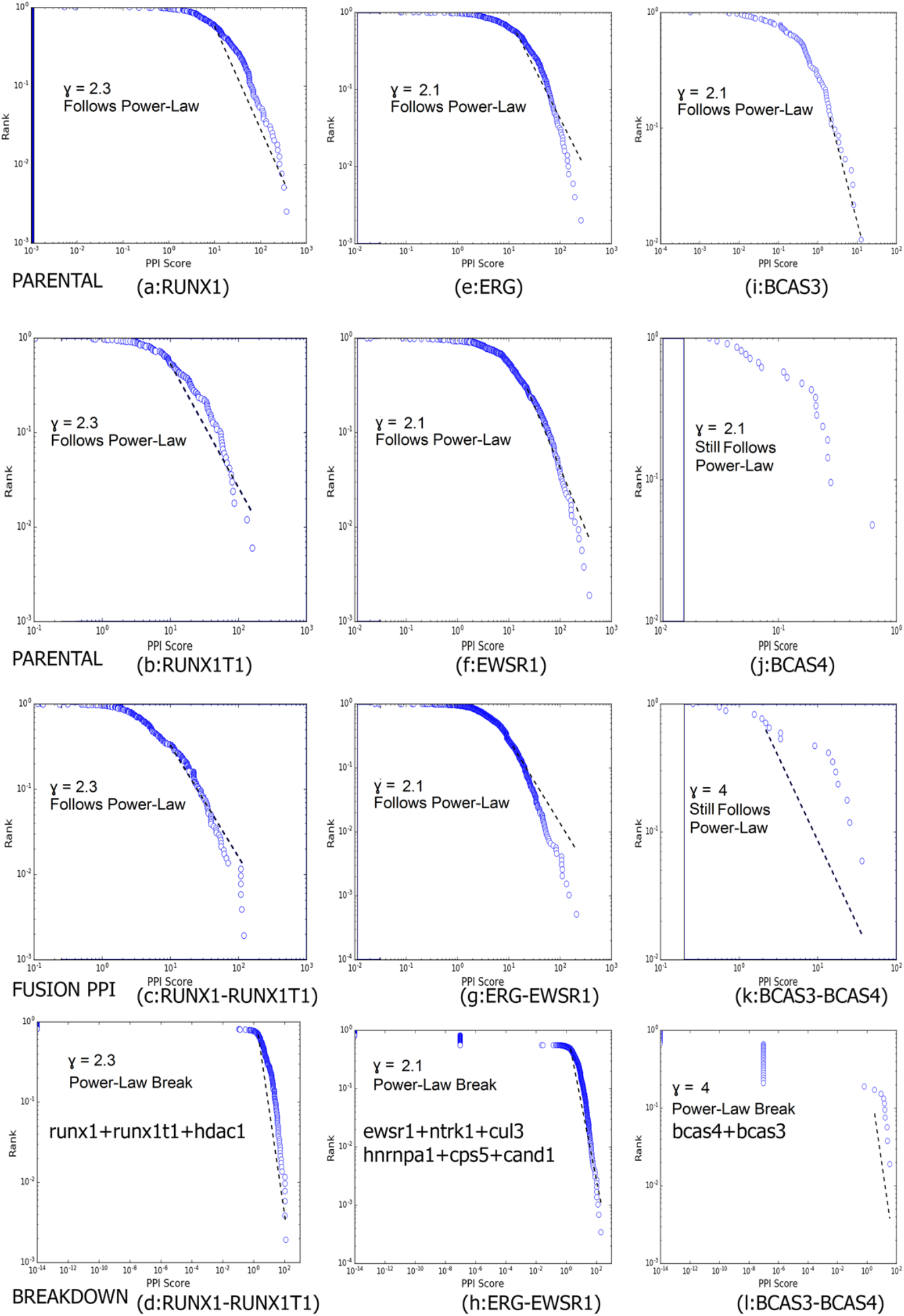
A comparative analysis of power-law breakdown in RUNX1-RUNX1T1 (LK, LY, ME and GL), EWSR1-ERG (SC) and BCAS3-BCAS4 (CA). In LK and ME, the overall distributions of breakdown points/sites were considerably higher than in CA.

We next constructed breakdown point networks for all cancers (Supplementary Fig. S3a). The proportion of fusions under the hierarchical category was greatest in SC, where multiple hubs had to be removed from communities to achieve breakdown. This can be correlated with changes in the functional pathways of fusions when specific hubs are targeted. Likewise, site-directed percolation identified essential and ubiquitous hubs, while targeted removal caused breakdown of PPI networks (Supplementary Fig. S3b).

### Bootstrap percolation detects essential and non-essential communities in fusion PPI networks

The main objective of performing bootstrap percolation was to prune out insignificant branches from fusion PPI networks, for the purpose of identifying communities. For example, in LK, LY, ME and GL, we could not identify non-essential communities for 10 fusion networks (PICALM-MLLT10). However, we identified essential communities for 438 fusion networks and 50 cases (FDR < 0.05) in which neither essential nor non-essential communities could be found (Supplementary Tables S11-13). In LY, the average betweenness centrality for essential community vertices was higher for Malignant Lymphoma - DE (MALY-DE) than for Lymphoid Neoplasm Diffuse Large B-cell Lymphoma - TCGA, US (DLBC-US). This is due to a greater involvement of hubs in fusion PPIs of MALY-DE in the handling of interactors of non-hubs. In GL, GBM-US and LGG-US similar average betweenness centrality for essential community vertices was seen, whereas in LK, CLLE-ES considerably higher average betweenness centrality was noted. Thus, we found that in LK, LY, ME and GL, hub communities have more control over non-hubs in forming novel interactions (Supplementary Fig. S4). Moreover, in SC, we identified essential communities for 426 fusion networks, non-essential communities for 427 fusion networks, as well as 9 cases in which no non-essential community was found and 27 cases in which no essential community was found (Supplementary Table S12). In CA, we identified essential communities for 1,491 fusion networks, non-essential communities for 1,493 fusion networks, and 426 cases in which neither were found (***Supplementary Table S13***). Finally, in CA, we found greater betweenness centrality among fusions in BRCA-US, CESC-US, KIRP-US, OV-AU and RECA-EU. The lowest betweenness centrality among fusions was found in PACA-AU and THCA-US (Supplementary Fig. S4). Overall, fusions in CA had less betweenness centrality than SC or LK, LY, ME and GL (Supplementary Fig. S4). Supplementary Fig. S5 provides a comparative analysis of communities in RUNX1-RUNX1T1 (LK, LY, ME and GL), EWSR1-ERG (SC) and BCAS3-BCAS4 (CA). These results indicate that essential communities have a key role in local hypothesis behavior, although this should be further validated using the preferential attachment model.

### Preferential attachment of proteins follows the Matthew effect

The Mathew effect is reflected as the more the hubs that are clustered, the more interactions they have. We found that essential and non-essential clusters whose average degree PPI score was less than that of the higher degree hubs tended to act as branches of more connected root clusters. We used this connectivity association to further distinguish between interactions in LK, LY, ME and GL, SC and CA PPI networks, based on their preferential attachment scores (PAS) (defined in Methods). For this study, we only considered preferential attachment for essential communities. Particularly, Figure 2 presents the preferential attachment model implemented on the RUNX1-RUNXIT1 fusion PPI (LK, LY, ME and GL) (Supplementary Table S14). Here, the initial size of the cluster was three, forming a clique of size four (SMARCC1, KMT2A, CREBBP, SMARCA4), which plays a significant role in RAS-oncogene-signaling and transcription activation. Similarly, the clique of size six plays a role in apoptosis, followed by successive attachment of new proteins to the original clique, finally forming the size ten community. Supplementary Fig. S6 presents the preferential attachment model implemented on EWSR1-ERGPPI (SC), an important fusion (Supplementary Table S15). In CA, we obtained preferential attachment vertices (PAVs), defined as major points of contact for preferential attachment, specifically, FN1, ATXN10, EGFR (GSTK1); ATXN10, and VCP (ECV) (ATXN10-FBLN1) (Supplementary Table S16). Supplementary Fig. S7 presents the preferential attachment model implemented on the BCAS3-BCAS4 fusion PPI (CA). PAS values for LK, LY, ME and GL, SC and CA are provided in Supplementary Tables S17-19.

**Figure 2.**
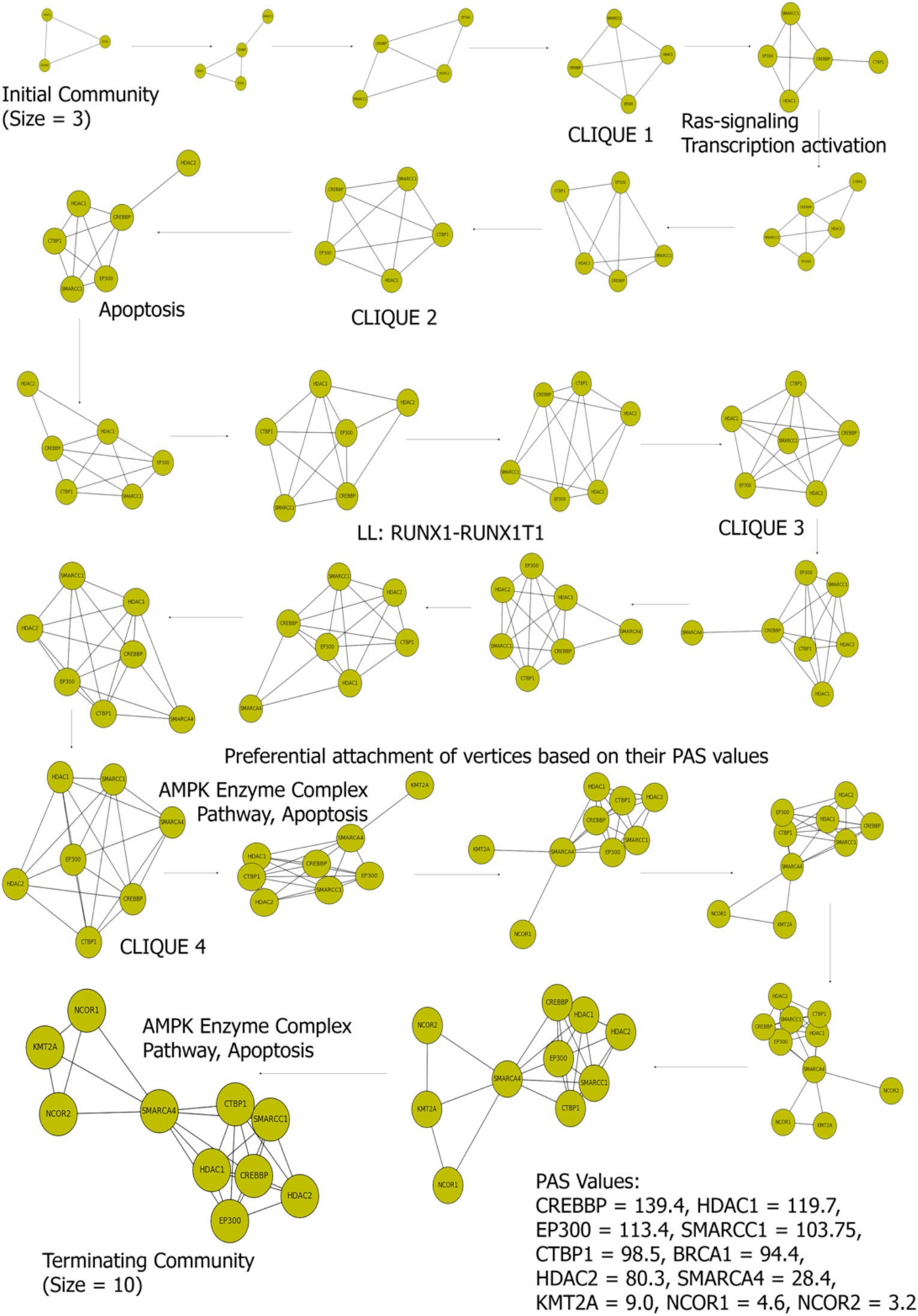
A preferential attachment model of implementation in RUNX1-RUNXIT1 fusion PPI (LK, LY, ME and GL). The initial size of the cluster is three, forming a clique of size four (SMARCC1, KMT2A, CREBBP and SMARCA4), which has a significant role in Ras-signaling and transcription activation. Similarly, the clique of size six plays a role in apoptosis, followed by successive attachment of new proteins to the original clique, eventually forming the size ten community.

To understand correlations with the preferential attachment scores (PAS) of breakdown points, essential communities and essential community vertices, as well as the role of PAS in distinguishing cancer sub-types, we created a cluster of cancer sub-types (Fig. 3). Although the number of aliquot IDs was considerably less in LK, LY, ME and GL, due to the presence of a higher number of communities with preferential attachment vertices (PAVs), network compactness increased. In CA, the highest number of PAVs was found in BRCA-US (5347) and OV-AU (4879), due to the huge size of the network itself. Overall, PAS distribution was lower than in LK, LY, ME and GL. Finally, in SC, SARC-US (5112) had a high number of PAVs due to high network compactness (Supplementary Fig. S8a). In contrast, PAS values were higher in LK, LY, ME and GL, followed by SC and CA (Supplementary Fig. S8b). This suggests a considerably higher compactness ratio among fusion PPIs in LK, LY, ME and GL than in SC and CA, due to larger essential communities. These results indicate that the presence or absence of essential modules, together with attachment patterns, can be distinguished in PPI networks of LK, LY, ME and GL, SC and CA tumor types.

**Figure 3.**
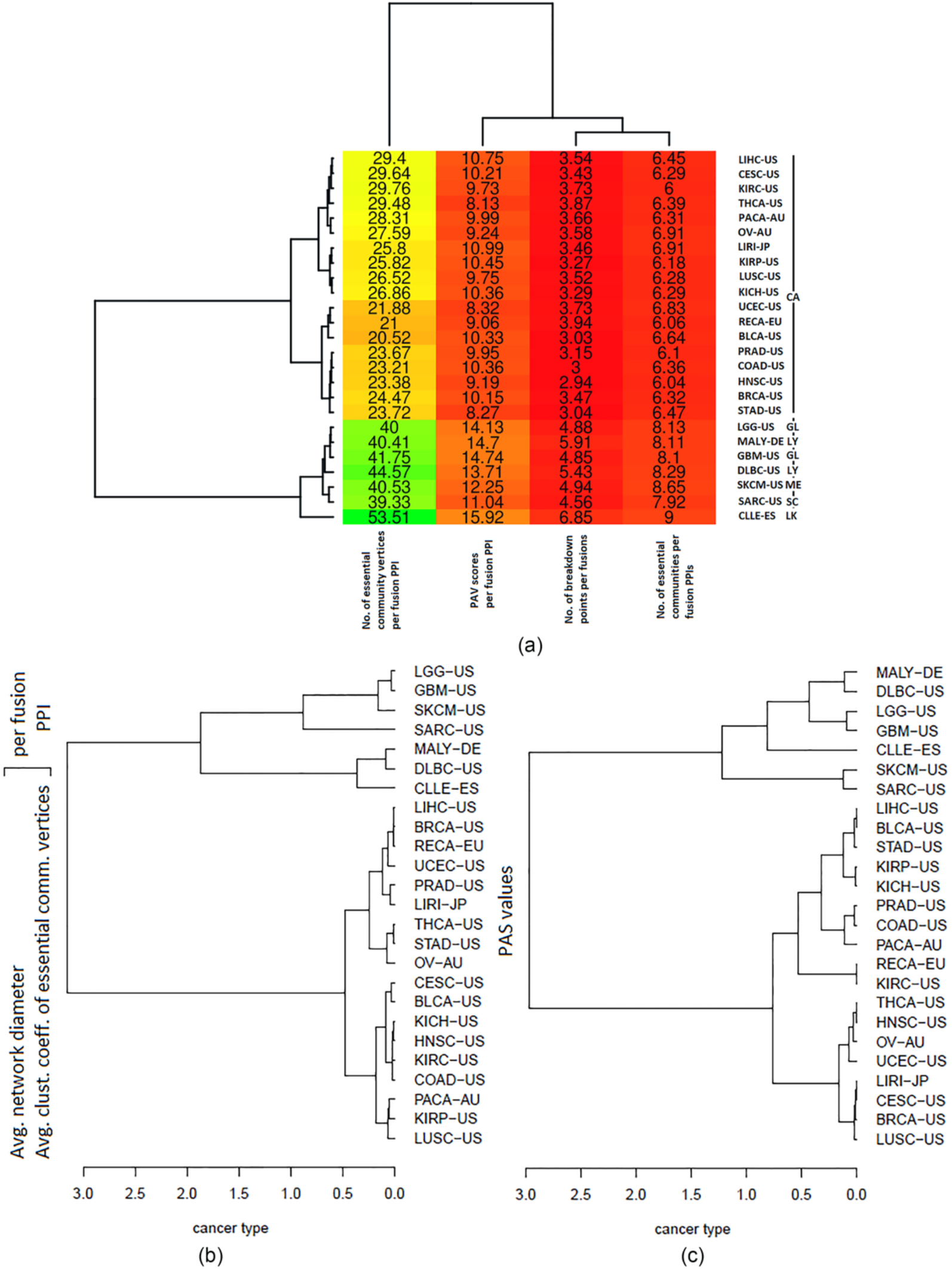
Clustering of cancer-subtypes. (a) We found that PAV presence of breakdown points can indeed be used to distinguish the clusters. (b) To ensure that our observation on preferential attachment is valid, we clustered the cancer sub-types separately using features of essential communities, such as average network diameter and average clustering coefficient. We found that indeed, based on our observations, fusion PPIs in different cancer sub-types can be clustered. (c) We used the PAS values to examine whether fusion PPIs in different cancers can indeed be differentiated; this clearly stratified the cancer sub-types based on their PAS values into six clusters.

### Gene and pathway enrichment analysis highlights that some down-regulated pathways in LK and LY are enriched in CA

We mapped 57,820 genes in the 25 integrated cancer sub-types. Expression profiling data of LK, LY, ME and GL, SC and CA prior to and following normalization were compared between normal and tumor samples. Only limited deviations in the systems were observed (Supplementary Fig. S10). Differentially expressed genes (the top 10) in the six cancer groups are shown in Supplementary Table S20. Genes such as *PRL, C20orf103, RET, GGA2, ICSBP1, ITGA7, DAP, IRAK1* and *PPARG* were over-expressed almost exclusively in fusion protein MYST3-CREBBP, present in CLLE-ES (LK). Of these genes, *PRL* and *RET* were reported to be involved in leukemogenesis^26,27^. Interestingly, the *STAT5A* and *STAT5B* genes were significantly under-expressed in MYST3-CREBBP. This indicates that a negative regulatory effect between prolactin and STAT5 ^26^. Finally, GSEA results showed comparable findings (Supplementary Fig. S10).

In SC, we obtained 36 up-regulated and 30 down-regulated genes (Supplementary Table S18). Likewise, 42 pathways were up-regulated and 10 were down-regulated. In CA, we obtained 67 up-regulated and 72 down-regulated genes (Supplementary Table S18). The cell cycle, DNA replication, spliceosome, proteasomes, mismatch repair, p53 signaling pathway, nucleotide excision repair and another 10 pathways were up-regulated in CA. In a series of 76 aliquot IDs, we observed that TRIM24/TIF-1α mRNA levels were significantly increased, as compared to normal breast tissues. Additionally, the down-regulated pathways in LK and LY were all enriched in CA (olfactory transduction, the renin angiotensin system and neuroactive ligand receptor interaction). However, three pathways (fatty acid metabolism, the adipocytokine signaling pathway and valine, leucine, and isoleucine degradation) were down-regulated in CA but up-regulated in LK and LY. These results indicate that these differentially expressed genes are all components of the breakdown point network, thus highlighting their changes in the expression levels as potential biomarkers in various cancers.

### Survival analysis shows that a family history of cancer and the nature of disease evolution in early prognosis have predictive powers

The study cohort comprised 672 aliquot IDs (Fig. 4, Supplementary Fig. S11). Overall survival (OS) and event-free survival (EFS) curves were constructed using the Kaplan–Meier method. A log–rank test was used to estimate differences between patients. All statistical tests were two-sided and performed at a 5% false discovery rate, p-value < 0.05. For LK (CLLE-ES, 31 aliquot IDs), the median OS was 30.5 months for 28% of patients with unaltered genes (p-value = 0.67, OR = 0.87, 95% CI = 0.3488 – 1.278). The median OS was 40 months for 32% of patients with altered genes. The mean age at diagnosis was lower than that reported in the literature^28^. Our literature search did not reveal a clear cut-off age to define a prognosis of CLLE-ES. As such, this should be considered in further analyses, especially for younger patients.

**Figure 4.**
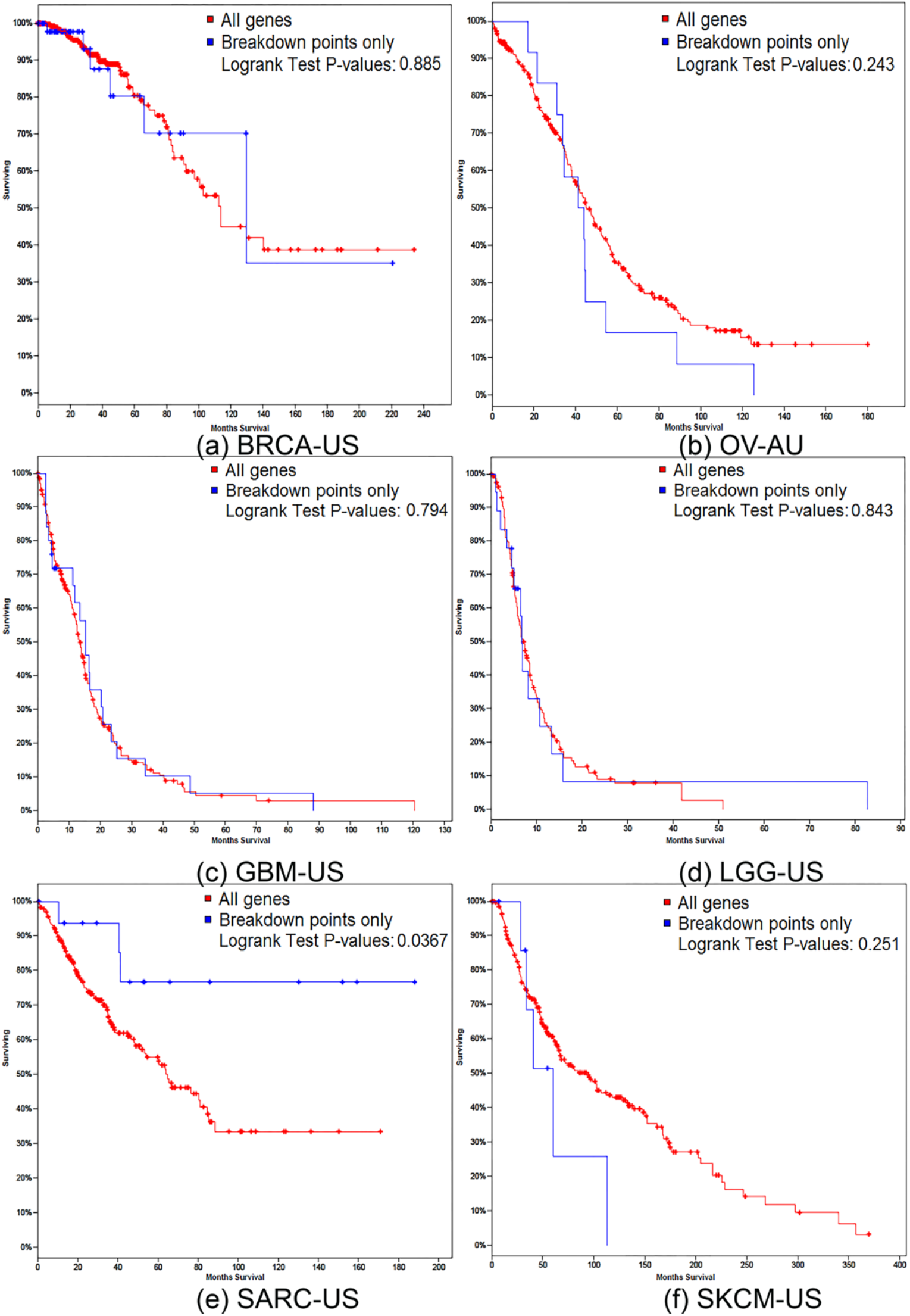
Analysis of OS and EFS. Survival curves of overall survival (OS) and event-free survival (EFS),. constructed by the Kaplan– Meier method, using R. The log–rank test was used to estimate differences between patients. All statistical tests were two-sided and done at the 0.05 significance level. The OS was defined as the time interval between the date of diagnosis and the date of death or the date of the last follow-up visit. The EFS was considered the period between the date of diagnosis and the date of relapse, induction failure or date of death from any cause.

For SC, during a median follow-up period of 20 months among 27 aliquot IDs, 1% had died by the time of analysis. The estimated median OS of all patients was 60 months for altered genes, and 40 months for unaltered genes (t-test, two-sided, p-value = 0.0367; OR = 0.23; 95% CI = 0.0188 – 0.5795), while the median EFS of all patients was 50 months (t-test, two-sided, p-value = 0.014; OR = 0.21; 95% CI = 0.011 – 0.478) (Fig. 4, Supplementary Fig. S11). Some studies have also reported OS in adults as being dramatically worse than in children with sarcoma^28^. Notably, survival studies should be performed post-treatment, as Sultan et al.^28^ reported that adults with sarcoma showed worse survival than children with similar tumors. For CA, the median follow-up for the entire cohort (491) was after 45 months, and the interquartile range was 12-56 months.

Here, we present the results for the two cancer sub-types with the highest number of aliquot IDs, namely, BRCA-US (76) and OV-AU (62). Overall, 2% of patients with breast cancer died, with 90% of the events considered as cancer-related, whereas 3% of patients with ovarian cancer died, with 86% of the events considered as cancer-related. The median OS was 55 months for altered genes and 42 months for unaltered genes in patients with BRCA-US (p-value = 0.885; OR = 0.93; 95% CI = 0.58 – 1.68); and 60 months for unaltered genes and 40 months for altered genes in patients with OV-AU (p-value = 0.243; OR = 0.61; 95% CI = 0.18 – 0.67) (Fig. 4, Supplementary Fig. S11). These results support considering family history of cancer and disease progression at early prognosis. Furthermore, the use of breakdown points as drug-binding targets could enhance patient survival.

### A drug-binding site map identifies breakdown points belonging to communities as potential docking targets

Treatment options for CLLE-US vary greatly, depending on patient age, the disease risk group, and the indication for treatment. Patients who might not be able to tolerate the side effects of strong chemotherapy are often treated with chlorambucil alone or with rituximab or obinutuzumab. Our study identified HDAC2, MDM2 and EGFR as drug-binding sites in the LK (CLLE-US) fusion PPI network (Supplementary Fig. S12); these are all breakdown points for which targeted drugs are available (Supplementary Table S21). DLBC-US tends to grow quickly and is most often treated by chemotherapy, usually with a regimen of four drugs, cyclophosphamide, doxorubicin, vincristine and prednisone, plus the monoclonal antibody rituximab. For LY (DLBC-US, MALY-DE), the predicted binding sites were HDAC1, HDAC5, PARP1 and EGFR (Supplementary Fig. S12). Some melanomas have a genetic mutation that makes them sensitive to a group of medications called BRAF inhibitors. This study identified ESR1, AR and PPARG as drug-binding sites in the ME (SKCM-US) fusion PPI network (Supplementary Fig. S12).

The current standard of care for patients with newly diagnosed glioblastoma includes surgery, radiation therapy and chemotherapy. Approved drug treatments include temozolomide for patients with newly diagnosed GBM. This which provides a median survival advantage of 2.5 months when added to surgery and radiation therapy. Other drugs used are bevacizumab and lomustine, an anti-cancer chemotherapy approved as monotherapy or for use with other drugs to treat recurrent brain cancer. This study identified HDAC2, HDAC1, EGFR, HDAC5, HDAC7, EGFR, NTRK1 and KIT as drug-binding sites for the GL (GBM, LGG) fusion PPI network (Fig. 5).

**Figure 5.**
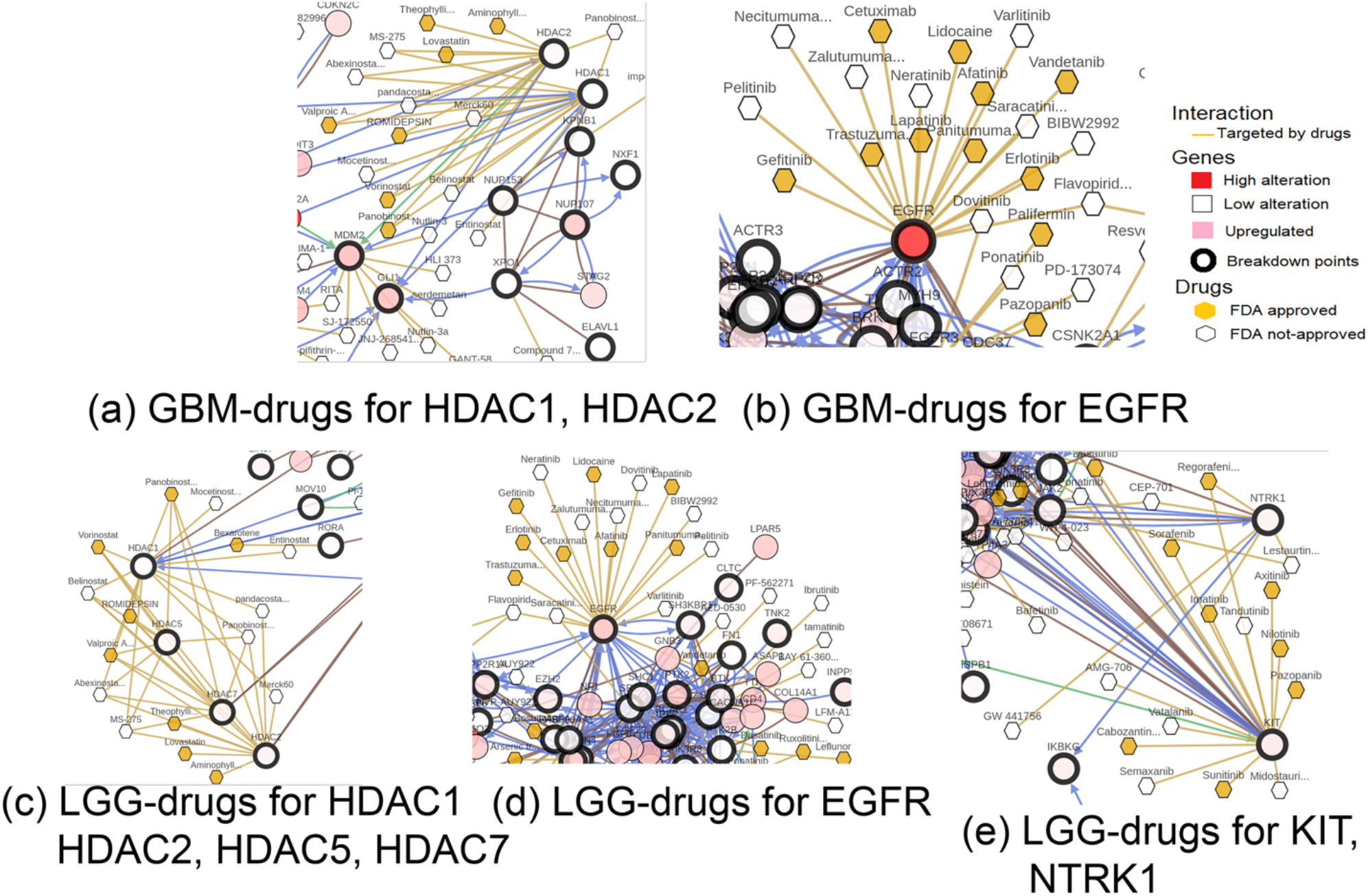
Drug-binding sites for GL. The drug-binding sites for the GL (GBM and LGG) fusion PPI network are HDAC2, HDAC1, EGFR, HDAC5, HDAC7, EGFR, NTRK1 and KIT.

Targeted drugs used to treat one type of sarcoma, namely gastrointestinal stromal tumor, included imatinib, sunitinib and regorafenib. We identified HDAC6, HDAC5 and TOP1 as drug-binding sites for the SC (SARC-US) fusion PPI network (Supplementary Fig. S12). ERBB2, SCN2A, TOP2A, CD74 and EHMT were identified as drug-binding sites for the CA (BRCA-US, OV-AU) fusion PPI network (Supplementary Table S21). Drug-binding sites predicted breakdown points belonging to communities that are potential docking targets.

### Percolation-based analysis of the Philadelphia positive/negative chromosome for chronic myelogenous leukemia (a case study)

In chronic myelogenous leukemia (CML), a constitutively active tyrosine kinase (TK), *Imatinib*, is used to decrease BCR-ABL activity. Imatinib is a 2-phenyl amino pyrimidine derivative that functions as a specific inhibitor of a number of TK enzymes. *Imatinib* occupies the TK active site, which results in decreased activity. TK catalyzes enzymatic activity by means of terminal phosphate transfer from ATP to its tyrosine residue substrates. *Imatinib* attaches tightly to the ATP binding site *of BCR-ABL*, locking it in a closed or self-inhibited conformation, thereby inhibiting the enzyme activity of the protein semi-competitively. Due to this activity, many BCR-ABL mutations can cause resistance to *Imatinib* by shifting the equilibrium toward the open or active conformation^39^. Although imatinib is quite selective for BCR-ABL, it also inhibits c-kit and PDGF-R, as well as ABL2 (ARG), DDR1 TKs and NQO2 (oxidoreductase)^40^. *Imatinib* also inhibits the ABL protein of non-cancer cells, although these cells normally have additional redundant TKs, which enable continued function^41^. In general, most patients under imatinib treatment show major or complete cytogenetic responses, with restoration of a Ph-negative hematopoiesis. However, some studies showed the appearance of clonal chromosomal abnormalities in patients with Ph-negative cells upon imatinib treatment^42-43^.

BCR-ABL is a common point for several downstream pathways that can be targeted when designing novel inhibitors^44^. These include the Ras/MapK pathway, which leads to increased proliferation due to increased growth factor-independent cell growth, the Src/Pax/Fak/Rac pathway, which affects the cytoskeleton and leads to increased cell motility and decreased adhesion, the PI/PI3K/AKT/BCL-2 pathway, wherein bcl-2 is responsible for keeping the mitochondria stable, suppressing cell death by apoptosis and increasing survival, and the JAK/STAT pathway, which is responsible for proliferation. One approach for designing potential inhibitors downstream of the BCR-ABL pathway is to target the PI3K/AKT/mTOR pathway, which is activated by BCR-ABL and additional mechanisms, leading to impaired apoptosis of Ph+ cells. While significantly down-regulating the expression of the main elements in this pathway, such as mTOR, PI3K, PS6K, 4EBP-Thr-37/46′4EBP-Thr-70 and EIF4E, imatinib significantly induces the expression of PP2A. This results in inhibition of the PI3K/AKT/mTOR pathway^45^ (Fig. 7a). Several mouse models have been used to better define the signaling pathways downstream of JAK2 in Ph-negative CMLs and myeloproliferative neoplasms (MPNs). In humans, the fibroblasts are polyclonal, do not carry the JAK2-mutation, and are not part of the malignant clone^46^. However, considerable evidence suggests that one or more cytokines secreted by the clonal megakaryocytes, including PDGF, FGF and TGF-β, are the cause of fibroblast proliferation and fibrosis in CML and MFN ^47^. This has raised great interest as to whether molecularly targeted drugs aimed at the JAK2 signaling pathway can be effective treatments for CMLs.

**Figure 6.**
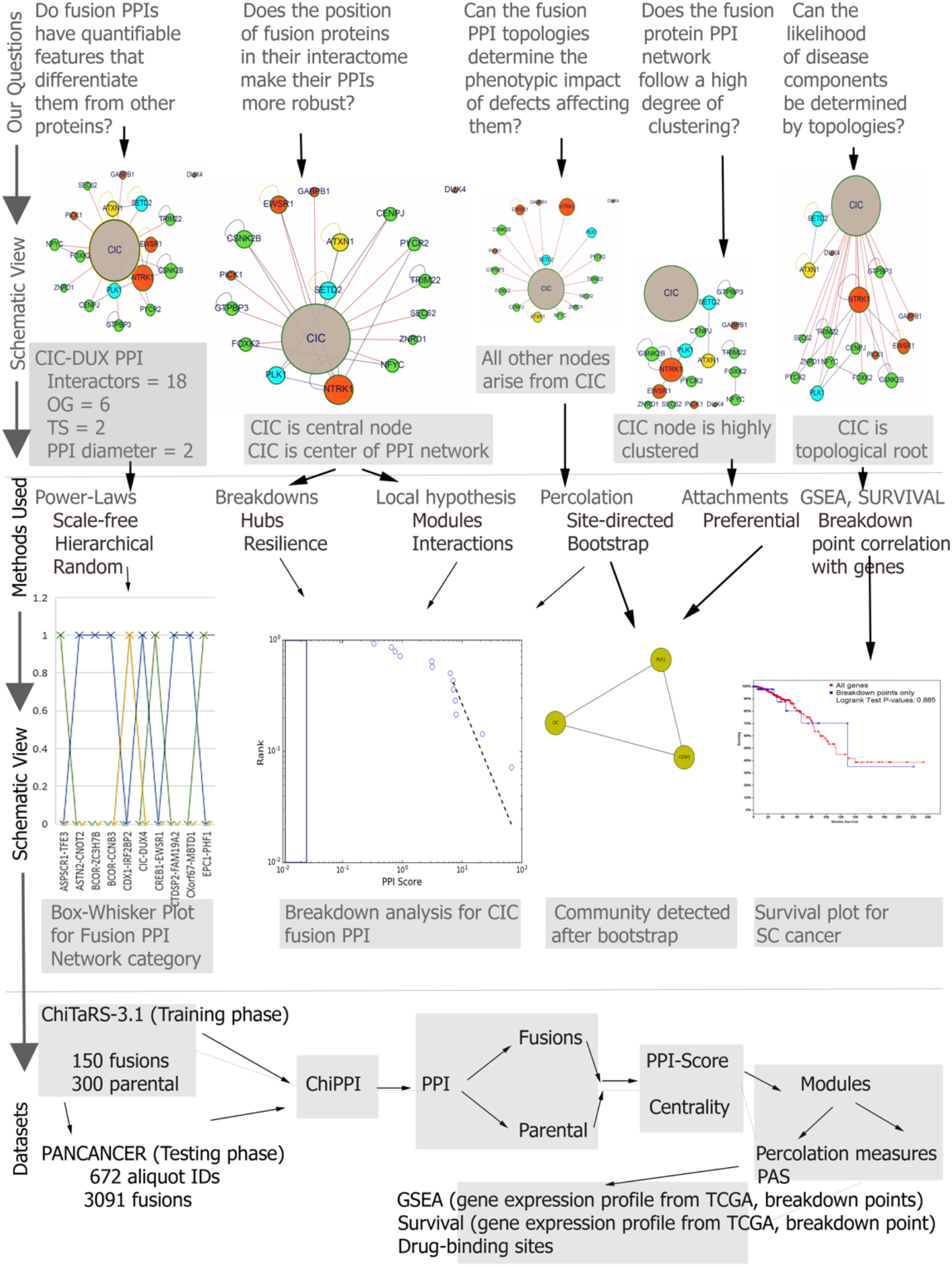
Overall methodology. The overall method initiates with defining the set of questions, focusing on whether the fusion PPIs have features that make them different from normal proteins; followed by implementation of power-laws, breakdowns, percolations, GSEA and survival; ending with the dataset explanation.

**Figure 7.**
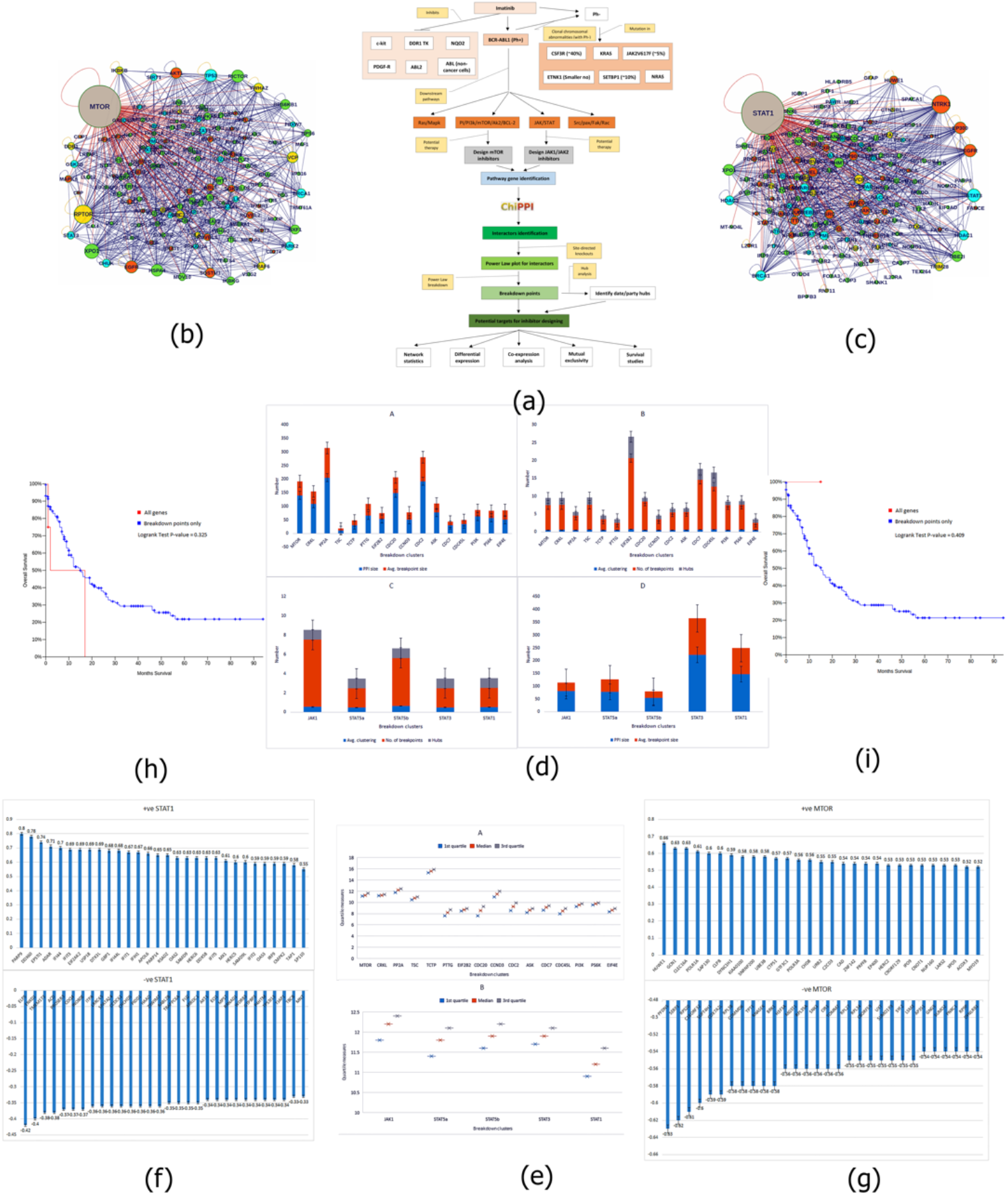
Percolation-based case study. (a) The methodology (b) Breakdown cluster for MTOR; graph size = 140, clustering = 0.524, hubs = MTOR, RPTOR, XPO1 (c) Breakdown cluster for STAT1; graph size = 147, clustering = 0.521, hubs = STAT1 (d) Breakdown point network statistics, ‘A’ MTOR – PPI size = 82.12, avg. breakpoint size = 38.52, ‘B’ MTOR – avg. clustering = 0.55, no. of breakpoints = 7, hubs = 2, ‘C’ JAK – PPI size = 116.6, avg. breakpoint size = 69.97, ‘D’ JAK – avg. clustering = 0.53, no. of breakpoints = 4, hubs = 1 (e) Differential expression plot, ‘A’ MTOR: 1st quartile = 10.43, median = 10.08, 3rd quartile = 9.69 ‘B’ JAK: 1st quartile = 11.48, median = 11.8, 3rd quartile = 12.08 (f) Co-expression in MTOR gene (top 30) ‘A’ +ve, highest in HUWE1 ‘B’ –ve, lowest in PFDN5 (g) Co-expression in STAT1 gene (top 30) ‘A’ +ve, highest in PARP9 ‘B’ –ve, lowest in ELP5 (h-i) Survival plot for MTOR, JAK breakdown points, survival increases when considering breakdown points.

Using our percolation-based algorithm, we identified breakpoints in the PI3K/AKT/mTOR and JAK/STAT pathways, which can act as potential target sites for designing active inhibitors (Fig. 7b-c). Sixteen breakdown clusters were detected for the MTOR pathway, namely, MTOR, CRKL, PP2A, TSC, TCTP, PTTG, EIF2B2, CDC20, CCND3, CDC2, ASK, CDC7, CDC45L, PI3K, PS6K and EIF4E. All these clusters have their own PPIs, with certain proteins acting as hubs. Likewise, five breakdown clusters were identified for the JAK pathway, namely, JAK1, STAT5a, STAT5b, STAT3 and STAT1. Statistical analysis showed that the structure of the breakdown points in both pathways was more compact than in other places, thereby making them more robust against perturbation (Fig. 7d). Thus, targeting the breakdown points to inhibit enzymatic activity has potential, especially with drugs such as imatinib, which have multiple binding sites. Differential expression analysis shows that *MTOR, CRKL, TCTP, TSC, PTTG, EIF2B2, CCND3, CDC2, ASK, CDC7* and *CDC45L* were down-regulated and PP2A, CDC20, PI3K, PS6K and EIF4E were up-regulated in the MTOR pathway. At the same time, *JAK1* was down-regulated and *STAT5a, STAT5b, STAT3* and *STAT1* are up-regulated in the JAK pathway (Fig. 7e). For mutual exclusivity at MTOR breakdown points, we found only two significant cases, namely, PTTG1, MTOR and PTTG1, CDC7. We found no significant cases in JAK breakdown points. Similarly, co-expression with MTOR genes revealed the highest positive correlation for HUWE1 and the lowest negative correlation for PFDN5. Likewise, co-expression with STAT1 genes revealed the highest positive correlation for PARP9 and the lowest negative correlation for ELP5 (Fig. 7f-g). Lastly, we found that survival significantly increased when breakdown point clusters were considered, instead of the entire gene set (Fig. 7h-i). Thus, breakdown points have high potential as targets for future drug development.

## DISCUSSION

PPI networks of fusion proteins have certain quantifiable features, such as high degrees of clustering and robustness, which distinguish them from normal proteins. An essential property of fusions is that they are not randomly placed in the interactome. As a result, their interactions with other proteins tend to be more robust and fragment late. Likewise, in fusion PPIs, lower degree hubs are less resilient against site-directed knockouts than are higher degree hubs. This suggests strong correlation between the local and network topologies of fusion PPIs. We also found that PPI networks of both parental and fusion proteins belonging to the random category were more prevalent in sarcoma (SC) and carcinoma (CA) than in leukemia (LK), lymphoma (LY), melanoma (ME) and glioblastoma (GL). In SC, PPI networks under the hierarchical category were more prevalent, due to the presence of higher degree hubs in communities. Moreover, the effects of ubiquitous hubs was more prominent in SC and CA, due to less clustering. Non-essential communities tended to attach themselves to essential communities in LK, LY, ME and GL. The compactness ratio among fusion PPIs was considerably higher in LK, LY, ME and GL than in SC and CA, due to the presence of larger and higher degrees of connectivity of essential communities. Thus, we can distinguish between the presence and absence of essential modules, and between attachment patterns in PPI networks of LK, LY, ME and GL, SC and CA tumors. GSEA results showed that the down-regulated pathways in LK and LY were all enriched in CA. However, three pathways were down-regulated in CA but up-regulated in LK and LY. Survival analysis demonstrated the contributions of a family history of cancer and disease progression to early prognosis. We presume that the identification of a tumor site as a predictor of survival in the youngest patients is a new finding, especially in CA. Likewise, drug-binding sites predict that breakdown points that belong to communities represent potential docking targets. Thus, our combinatorial approach provides an opportunity to target soft spots in fusion PPIs, due to their significant roles in the overall functionality of the network. This can be applied to target-based drug design, and to identifying cancer phenotype-specific central nodes in fusion networks.

## METHODS

### The power-law model of fusion PPI networks

To study breakdown against targeted knockouts of proteins in PPI networks of all cancer sub-types, we used the power-law model. A power-law is a relation between two entities, e.g., the frequency of proteins vs. their total occurrence in each PPI network. Power law distributions are “long-tailed distributions”, such that if we plot the number of nodes with degree d against d, we find that the area under the tail of the distribution is polynomially small. In contrast, the area under the tail of ‘Gaussian (normal) distribution curve is exponentially small. Essentially, we plotted the power-law for each fusion PPI, with the degree distribution of proteins vs. the ChiPPI PPI score for each protein on a log-log scale. We considered the Barabási–Albert model for studying PPI networks of individual fusion proteins and determined whether they behaved as scale-free networks using a preferential attachment mechanism. We observed that most of the fusion PPI networks could be classified as either scale-free networks, having power-law degree distributions, or as hierarchical. Some PPI networks exhibited random graph models, like the Erdős–Rényi (ER) model and the Watts–Strogatz (WS) model, which do not exhibit power laws^15^. The algorithmic and mathematical descriptions of our power-law model are described in the Mathematical description of methods section, below.

### Network categorization of PPI networks

As discussed in the previous sub-section, we categorized the PPI networks as scale-free, hierarchical and random graphs. The scale-free nature states that the degree sequence in a network satisfies a power law. We observed that LK (CLLE-ES), LY (DLBC-US, MALY-DE), ME (SKCM-US) and GL (GBM-US, LGG-US) belong to the scale-free category. Similarly, a network is hierarchical when two typical neighbors of a vertex/protein are more likely to be connected to each other than to another pair of vertices. We observed that SC (SARC-US) belongs to the hierarchical category. Likewise, random networks follow the classical Erdős-Rényi model. We observed that CA (BLCA-US, BRCA-US, CESC-US, COAD-US, HNSC-US, KICH-US, KIRC-US, KIRP, LIHC-US, LIRI-JP, LUSC-US, OV-AU, PACA-AU, PRAD-US, RECA-EU, STAD-US, THCA-US and UCEC) belong to the random category. We observed that both the parental and fusion proteins PPI follow the classical Poisson distribution; this highlights the fact that the PPI networks in the ChiPPI datasets are noise-free as well as accurate (Supplementary Fig. S1). The algorithmic and mathematical descriptions of our network categorization are described in the Mathematical description of methods section, below.

### The percolation model for studying connectivity in PPI networks

The robustness of a PPI network can be measured through its percolation threshold, which can be interpreted as the critical number of links or node removals that must occur for the network to shift from a regime in which it is only slightly perturbed to a regime in which it is fragmented into small disconnected clusters. The study of robustness encounters two variants. the first is the robustness of the topologies (maintenance of topological connectivity) of networks, called structural robustness, against failures of nodes or links. The second is the robustness of the dynamical processes (maintenance of dynamical processes) running on networks, referred to as dynamical robustness. Albert et al.^30^ observed that scale-free networks display high tolerance to random failures, although such networks are extremely vulnerable to targeted attacks. In the same year, the percolation model of Broadbent et al.^31^ was employed to analyze the structural robustness of networks^14,32^. A series of relevant studies followed^33,34^. In the current study, we considered that each interaction (or protein) in the PPI network as occupied with a probability of *p*. A cluster of interactions was defined as the set of neighboring occupied interactions. Thus, when *p* = 0, all interactions are null. For a small *p*, a sparse population of interactions results in only small clusters. As *p* increases, the mean size of the clusters grows and when *p* = 1, all interactions are occupied. Hence, as *p* increases from 0 to 1, a specific value of *p* appears, at which a large cluster, the incipient percolation cluster, emerges. This enables full connectivity of the network from one side to the other for the first time. If the size of the PPI network approaches infinity, transition from an unconnected to a connected network occurs sharply when *p* crosses a critical threshold called the percolation threshold. Regardless of the property that an interaction represents, this property percolates through the network, with emergence of the percolating cluster representing a phase transition. In this study, we performed both site-directed and bootstrap percolation for targeted knockouts of proteins from PPI networks. Site-directed percolation specifies targeted removal of proteins from a PPI network, whereas bootstrap indicates pruning out less significant links and identifying communities. The algorithmic and mathematical description of our percolation model are described in the Mathematical description of methods section, below.

### Site-directed percolation for systematic knockouts of hubs in PPI networks

For site-directed percolation, we initially selected 1% of the proteins, removed them and calculated the size of the largest connected component. If this size was < 50% of that of the proteins, we stopped here. Selection of proteins was based on their degree of connectivity in the network and was thus targeted. The functioning of complex networks, such as the internet and social networks, has been shown to be crucially dependent upon interconnections between network nodes. These interconnections are such that when some nodes in the network fail, nodes that are connected to the network through them also become disabled and the entire network may collapse. Thus, to understand network robustness of complex PPI networks, we needed to determine whether such networks could continue to function after a fraction of their proteins had been removed by site-directed knockouts. Thus, the robustness of a PPI network under attack is dependent upon the structure of the underlying network and the nature of any attack. Previous research on complex networks focused on two types of initial attack, random attack and hub-targeted attack. In a random attack, each node in the network is attacked with the same probability^14,15,32,34^. In a hub-targeted attack, the probability that high-degree nodes will be attacked is higher than for low-degree nodes^32-34^. However, random and hub-targeted attacks have been shown not to adequately describe many real-world scenarios in which complex networks suffer from localized damage, i.e., a node is affected, then its neighbors, and then their neighbors. In our study on fusion PPI networks, we observed that a hub-targeted attack necessarily destroys the structure of the network, and results in power-law breakdown. We found that the robustness or resilience of PPI networks of LK, LY, ME and GL were high due to strong clusters, compared to SC and CA. Thus, we needed to critically analyze proteins that belong to larger clusters. These clusters may result in high robustness or resilience of PPI networks. The algorithmic and mathematical descriptions of our site-directed percolation model are described in the Mathematical description of methods section, below.

### Bootstrap percolation for identifying communities in PPI networks

Bootstrap percolation initiates with the selecting of random proteins (in our case, proteins with lower degrees of connectivity, ubiquitous) and successively removing them until a compact cluster (community) of proteins is obtained. In our study, we identified essential and non-essential communities. We defined an essential community as one whose size is less than three, and a number of interactions for each participant in the community generates a clique. Similarly, a non-essential community is one whose size is equal to or more than three, but the number of interactions for each participant in the community generates a clique. We used essential communities for further analysis, as they participated in more PPIs for identifying whether proteins tend to interact with them in the PPI network, according to preferential attachment. For growing networks, the precise functional form of preferential attachment can be estimated by the maximum likelihood estimation^35^. In our study, we observed that in networks of LK, LY, ME and GL, compared to networks of SC and CA, proteins with lower degrees of connectivity bind to larger hubs, which have higher degrees of clustering. This preferential attachment nature of lower degree proteins is an important property of fusion PPIs. The algorithmic and mathematical descriptions of our bootstrap percolation model are described in the Mathematical description of methods section, below.

### Using the preferential attachment model to study the “rich get richer” behavior of hubs

Preferential attachment suggests that the more connected is a node, the more likely it is to receive new links. Nodes with higher degrees of connectivity have a greater likelihood of adding links to their network. For example, in a PPI network, an interaction from protein A to B means that protein A tends to interact with protein B. Thus, heavily linked nodes represent proteins with a high degree of connectivity and a relatively high number of interactions. When a novel protein initiates interaction with an already formed community, it is more likely to interact with the hub rather than with proteins with fewer interactions. The Barabási– Albert model assumes that in the World Wide Web, new pages link preferentially to hubs, i.e. to very well-known sites such as Google, rather than to pages that are less known. The model claims that this is consistent with the preferential attachment probability rule. Thus, if a protein has a higher number of interactions, its clustering coefficient increases, resulting in increased compactness. In other words, the more clustered a protein, the more interactions it has. This “rich get richer” phenomena or Matthew effect, describing the preferential attachment of earlier nodes in a network, drives growth of a PPI network^15^. Due to preferential attachment, a node that acquires more connections will increase its connectivity at a higher rate. Thus, an initial difference in the connectivity between two nodes will increase further as the network grows, while the degree of individual nodes will grow proportionally with the square root of time^36^. In our study, we observed some central modules in PPI networks where rigorous attachment of lesser connected vertices occurred. The algorithmic and mathematical descriptions of the application of the preferential attachment model to the current study are described in the Mathematical description of methods section, below.

### Breakdown point network

For each fusion PPI corresponding to each cancer sub-type, we generated the corresponding breakdown point network, which we identified from site-directed and bootstrap percolation. We identified the interactors for each breakdown point using ChiPPI^29^ and used DyNetViewer and the GraphML plugin of Cytoscape to construct the network^36,37^. The algorithmic and mathematical descriptions of constructing the breakdown point network are described in the Mathematical description of methods section, below.

### Gene and pathway enrichment

The GSEA technique, developed by Subramanian and colleagues^38^, is a widely used method for measuring the association between a set of genes and a phenotype in gene expression profiling data sets. GSEA enables detection of gene sets enriched in genes that are significantly associated with a phenotype of interest. Such enrichment is computed using the Kolmogorov-Smirnov (KS) statistic. This statistic compares the anticipated random distribution of a gene set and their actual distribution in a genome-wide list of genes ranked on the basis of their association with the phenotype. The KS statistic is normalized for gene set size and its significance is adjusted to consider multiple hypotheses testing. In our study, we used the gene expression data downloaded from TCGA for 672 aliquot IDs to identify genes and their corresponding pathways that are enriched in all 25 cancer sub-types. We also determined whether our predicted breakdown points were part of these gene sets, which play active roles in this enrichment. The algorithmic and mathematical descriptions of GSEA are described in the Mathematical description of methods section, below.

### Survival analysis

We used Kaplan-Meier analysis to provide an estimate of the overall probability of being recurrence-free following treatment. We surveyed gene expression value distributions in a population totaling 57,820 genes, expressed in 672 aliquot IDs. For each gene, we divided the cancer population into two groups, namely, the gene expression more than normal tissue (log2 lowest normalized value > 0) and the gene expression less than normal tissue (log2 lowest normalized value < 0). Survival rates of two sub-groups of patients for each gene were compared and tested with a log-rank test. Genes with a log-rank test value < 0.005 were filtered out into a gene list for further analysis. The enriched gene clusters that contained not only the identified gene but also its related genes were clustered using gene expression profiles to confirm the pathways or function units that play key roles in tumor evolution and patient survival. The Kaplan Meier plot was generated using R, whereas the log-rank test was conducted based on the survdiff function in the survival package (built-in in R). The algorithmic and mathematical descriptions of survival analysis are described in the Mathematical description of methods section, below.

### Drug-binding site network map

Drug-binding data were taken from the KEGG DRUG, NCI Cancer Drugs and Drugbank. Following breakdown point network generation and obtaining interaction information from ChiPPI, a drug-binding site network map was constructed using the DyNetViewer and the GraphML plugin of Cytoscape^37^.

### Mathematical description of methods

We initiated our methodology by listing the notations used throughout this study (Table 1).

**Table 1.**
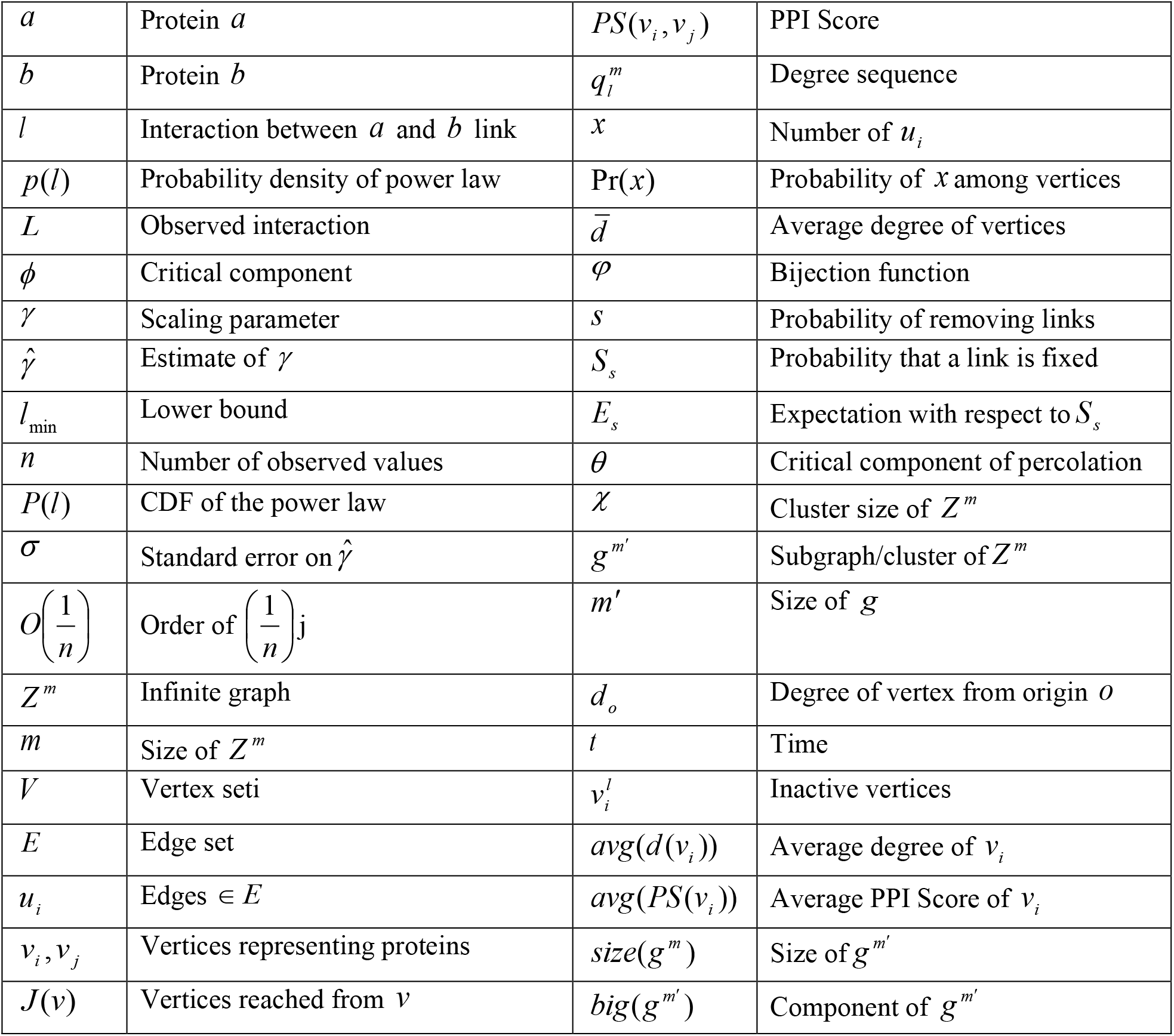
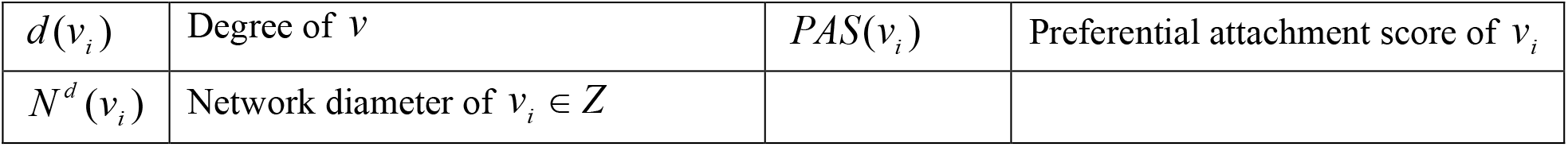
Table of notations.

#### The power-law model

Let *Z* (*V, E*) be a graph, where *V* is the vertex set and *E* is the edge set. More precisely, for *v*_*i*_ ∈*V*, we say that *v*_*j*_ ∈*V* when there exists a path of occupied edges that connects *v*_*i*_ and *v*_*j*_ (***Equation 1***):

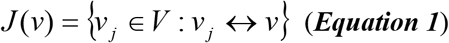

Let *l* represent the link or edge between two proteins *a* and *b*. We define a continuous power-law distribution by a probability density, *p*(*l*), such that (***Equation 2***):

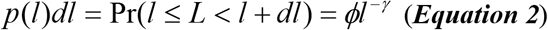

Here, *L* is the observed value, *ϕ* is a normalization constant and *Vγ* is the scaling parameter. This density diverges as *l* → 0 and ***Equation 2*** does not hold ∀*l* ≥ 0. For this purpose, we define a lower bound, *l*_min_ to power-law behavior. Thus, for *γ*>1, we calculate *ϕ* and the probability density of the power-law model by ***Equation 3***.

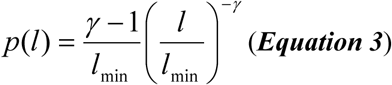

In this work, we considered numerical values of interaction with a probability distribution as follows (***Equation 4***):

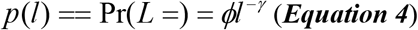

This distribution diverges at zero for the lower bound *l*_min_ > 0 of power-law behavior. We calculated *ϕ* and found the relation between *l*_min_ and *p*(*l*) using ***Equation 5***.

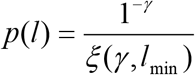

Here, *ξ* (*γ, l*_min_) is the Hurwitz zeta function (***Mező et al***., ***2010***), calculated using ***Equation 6***.

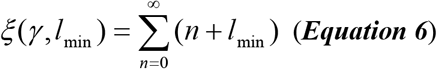

We also considered the complementary cumulative distribution function (CDF), *p*(*l*) = Pr(*L* ≥ *l*), defined by ***Equation 7***:

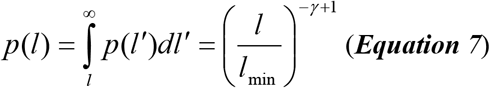

Next, for estimating scaling parameter, *γ*, requires a value for the lower bound, *l*_min_ of power-law behavior in the data. Assuming that *l* ≥ *l*_min_, we derived maximum likelihood estimators (MLEs) of *γ* with ***Equation 8***:

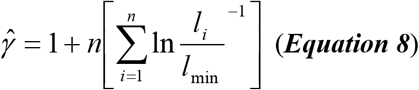

Here, *l*_*i*_, *i* = 1,…, *n* are the observed interaction scores, such that *l*_*i*_ ≥ *l*_min_ and 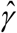 denotes estimates. Likewise, the standard error of 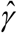, i.e. *σ*, is derived from the width of likelihood maximum (***Equation 9***).

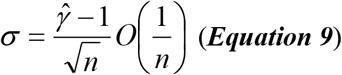

The computational implementation of the power-law model is explained in **Algorithm 1**.

##### Algorithm 1 Power-Law Model

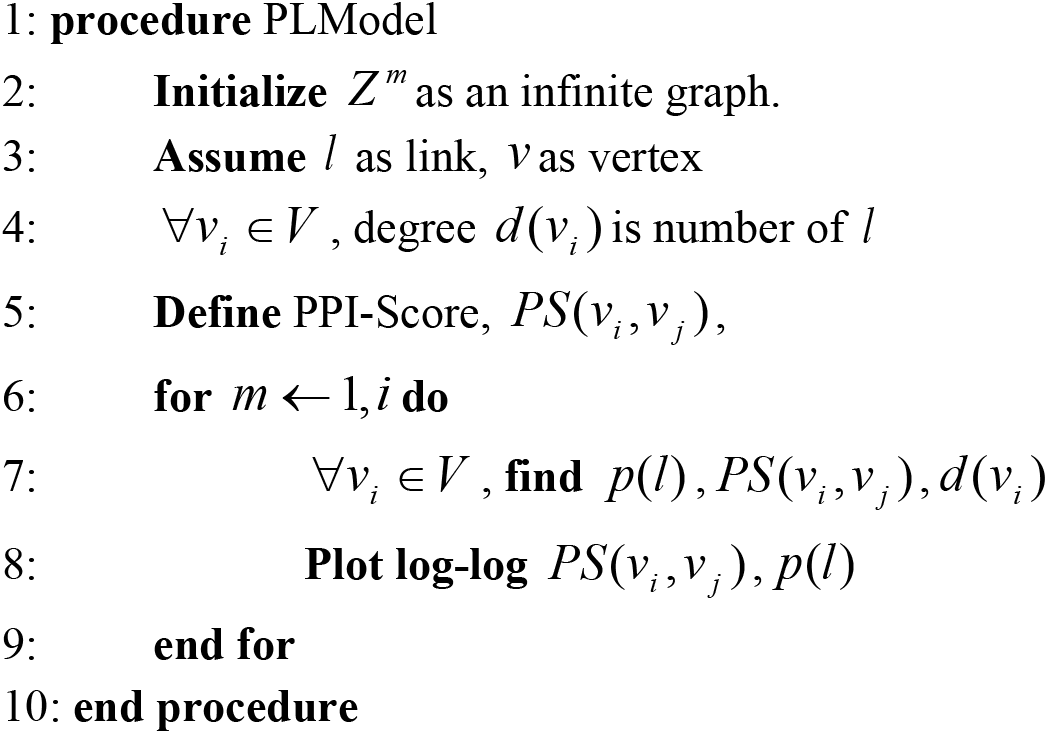

#### Network categorization

In a scale-free network, the degree sequence of a network of size *m*, which we denote by 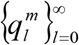, is calculated as follows (***Equation 10***):

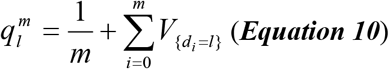

Next, we calculated the relation between *p*(*l*) and 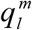 for a scale-free network (***Equation 11***).

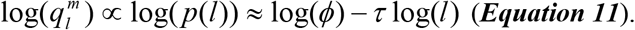

Here, a log-log plot of the degree/link sequence should be close to linear, and the slope is given by − *τ* (scaling parameter). In this case, ***Equation 10*** correlates with ***Equation 4***. Furthermore, ***Equation 4*** is valid for a simple graph of size *m*, where the maximal degree is *m* − 1. Due to complexity in the PPI network graph, there are many self-loops and multiple links.

For a hierarchical network when we draw two vertices uniformly at random from a graph of size *m*, the probability that there is an edge between the drawn vertices is equal to ***Equation 12***:

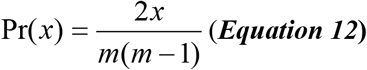

Here, *x* is the number of *l* ∈ *E*. Further, 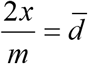, where 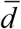 is the average degree of all vertices in the network (***Equation 13***),

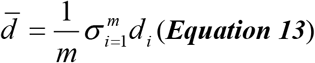

For a random network in the *Z* (*n, r*) model, a graph is constructed by connecting vertices randomly. All graphs with *n* vertices and *x* edges have equal probability of 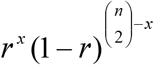 to remain present or absent, independent of other edges.

As *r* (weighting function) increases from 0 to 1, the model becomes more and more likely to include graphs with more edges (***Algorithm 2)***.

##### Algorithm 2 Network Categorization

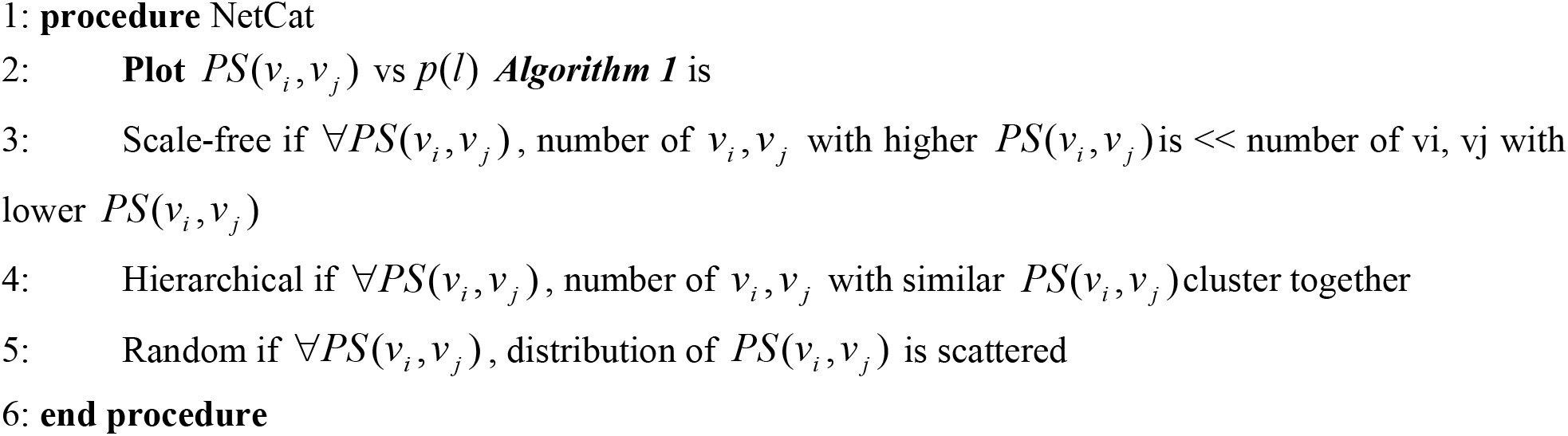

#### The percolation model

In graph *Z* (*V, E*), for every *v*_*i*_, *v* _*j*_ ∈*V*, there exists a bijection *φ* : *V* → *V* for which *φ* (*u*_*i*_ = *u* _*j*_) and {*φ*(*u*_*i*_, *φ* (*u*_*j*_)}∈ *E*. We performed site-directed percolation by independently removing each of the edges and/or vertices with a fixed probability of 1 − *s*. We assumed the probability that an edge/vertex that is occupied is fixed, denoted by *S*_*s*_, while *E*_*s*_ denotes expectation with respect to *S*_*s*_. We defined the percolation function *s* ↦ *θ* (*s*), by ***Equation 14***.

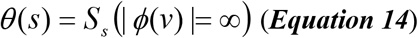

Here, *v* ∈*V* is an arbitrary vertex and the critical value is defined by *s*_*φ*_ (***Equation 15***).

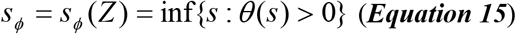

For both site-directed and bootstrap percolation, we identified the critical component with ***Equation 16*** for *s* → *s*_*ϕ*_.

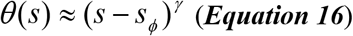

for some *γ* > 0. In our case, *θ* exists as (***Equation 17***).

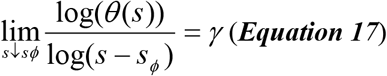

We also have *γ* for the expected cluster size, given by ***Equation 18***,

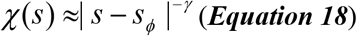

Therefore, *s*_*ϕ*_ defines the threshold after which a PPI network fragments due to systematic knockout of hubs or proteins.

#### Site-directed percolation

Continuing with ***Equation 17***, we can also obtain *γ* for a cluster in graph *Z* by implementing the classical model of site-directed percolation, as defined by ***Broadbent et al. (1957)***. For analyzing the extent of tolerance by PPI networks against site-directed percolation (knockouts), we used resilience, i.e., hoq PPI networks resist or combat perturbation (***Callaway et al***., ***2000; Barabási et al***., ***2011***), ***Algorithm 3***. Next, we calculated the PPI score (*PS*) and degree (*d* (*v*_*i*_)) of the remaining vertices along with their ranks (*p*(*l*)), followed by plotting a power-law on a log-log scale. We repeatedly performed this removal to identify the position at which a sudden breakdown of power-law occurs. The steps for site-directed percolation are given in ***Algorithm 3***.

##### Algorithm 3 Site-directed Percolation

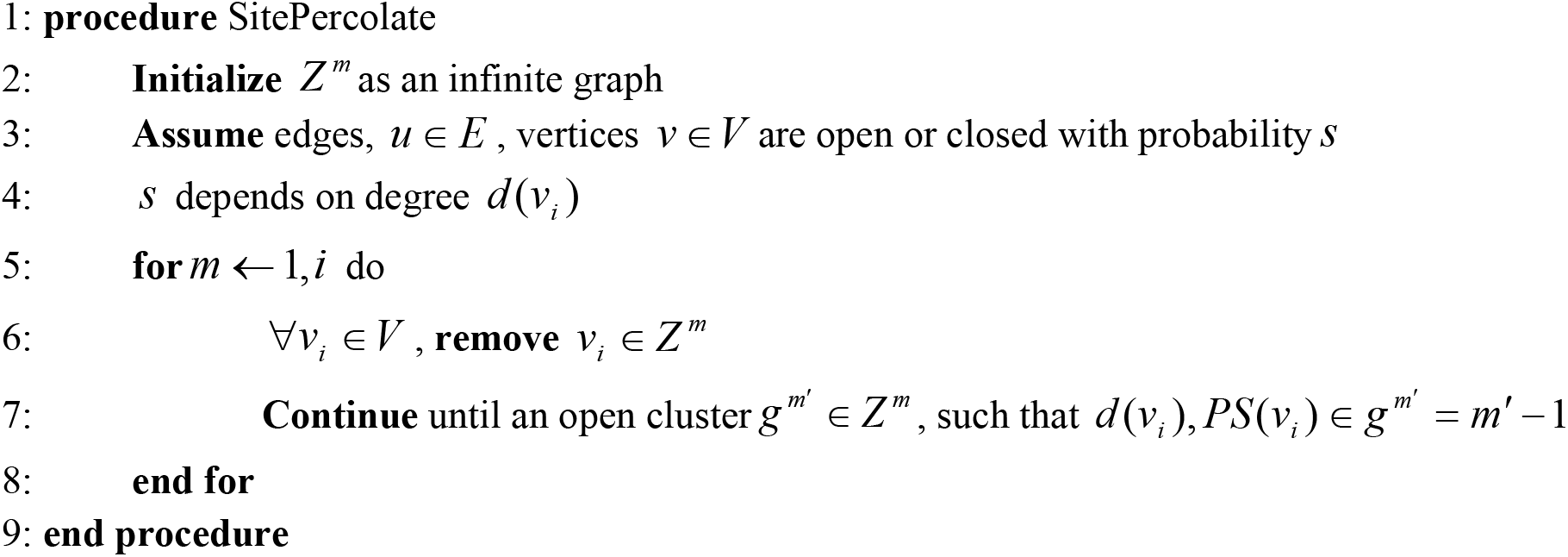

#### Bootstrap percolation

We used the bootstrap percolation model of ***Yukich (2006)*** to identify essential and non-essential communities and create a transitive graph with fixed degree *d* (*v*_*i*_) and percolation with a fixed percolation parameter *s*_*l*_.

Thus, a graph is scale-free for ***Equation 19***.

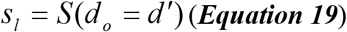

Here, *o* ∈ *Z* ^*d*^ is the origin. Given a graph *H*, we defined H-bootstrap percolation. Given a set *Z* ⊂ *E*(*g* ^*m*^) of initially selected edges on vertex set [*m*], we set *Z*_*o*_ = *Z* and defined

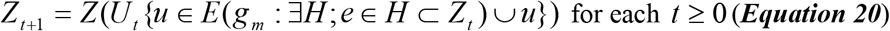

Thus, a new edge *u* is selected at time *t* + 1 if there exists a copy of *H* in *g*_*m*_ for which *u* is the only unselected edge at time *t*. Let ⟨ *Z* ⟩ _*H*_ = ⋃_*t*_ *Z*_*t*_ denote the closure of *Z* under *H* in *g*_*m*_ if ⟨ *Z* ⟩_*H*_ = *E*(*g*_*m*_). Thus, if *V* (*Z*) = [*m*] is chosen independently at random, each with probability *s*, then *s*_*ϕ*_ of *s* = *s*(*m*) at which percolation becomes likely *g*_*m*_, as ***Equation 21***,

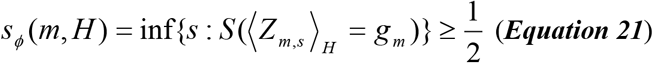

Here, *Z*_*m,s*_ is the PPI network graph, obtained by choosing each edge independently with probability *s*.

##### Algorithm 4 Bootstrap Percolation

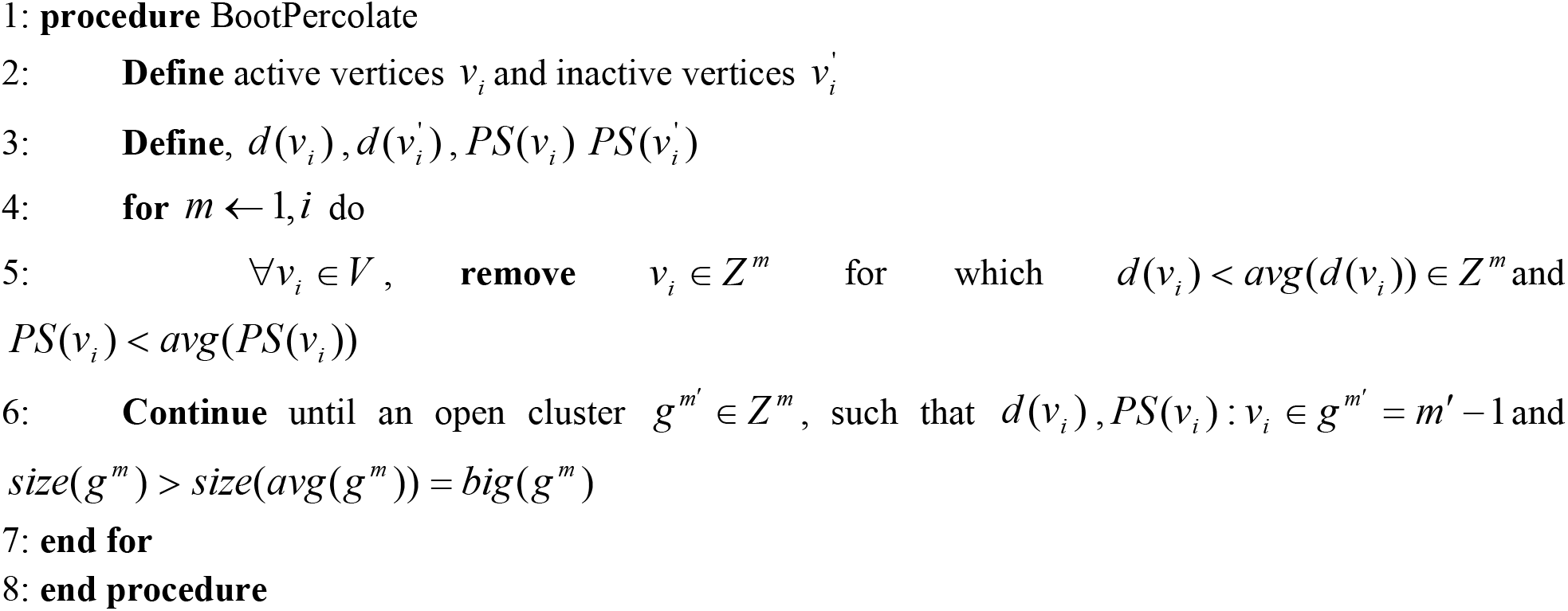

Using bootstrap percolation, we identified essential and non-essential communities after pruning out vertices with lower connectivities. Essential communities have 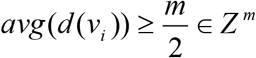, whereas non-essential communities have 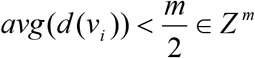 (***Algorithm 4***).

#### Preferential attachment model

Next, we formulated the preferential attachment model, an extension of the model by ***Barabási et al. (2002)*** for the graph process 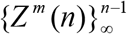. We started by formulating the model for *m* = 1, for which we started with *Z* ^1^ (1), which consisted of a single vertex with a single self-loop. We denoted vertices of the graph *Z*^*m*^(*n*) by 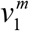, by 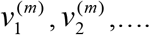 We denoted the degree of vertex 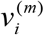 by 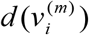, where a self-loop increases the degree by 2. Then, for *m* = 1, and conditionally on *Z* ^1^ (*n*), the growth rule to obtain *Z* ^1^ (*n* + 1) is as follows. We added a single vertex *n* + 1 having a single edge. The other end of this edge is equal to *i* = *n* + 1 with a probability proportional to 1 + *θ*, and to *i* ∈[*n*] with a probability proportional to 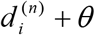, where *θ* ≥− 1is a parameter of the model. Thus, we defined ***Equation 22*** for *S* as the probability that an edge in the PPI network, which is already occupied, is fixed or not.

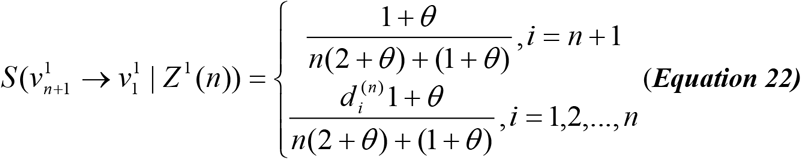

***Algorithm 5*** discusses the steps of the preferential attachment model.

##### Algorithm 5 Preferential Attachment Model

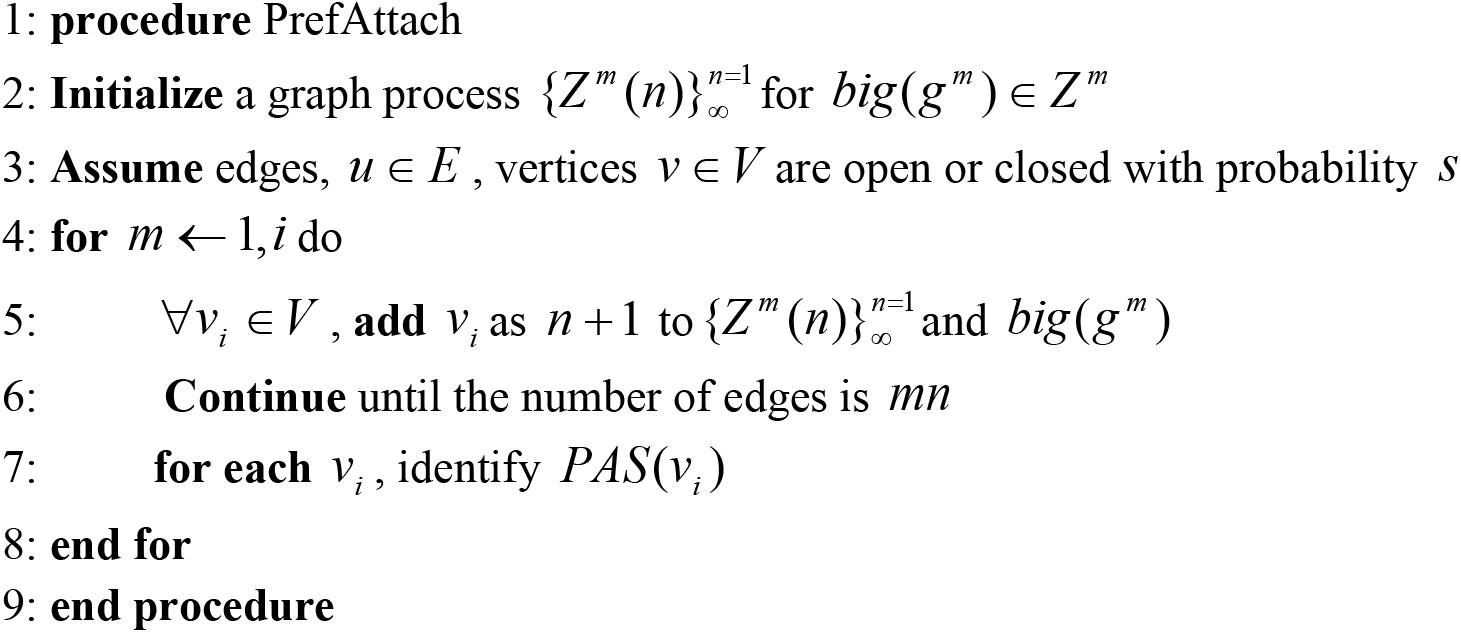

An important parameter that we obtained from this analysis is PAS, defined as the attachment specificity of an incoming vertex to the essential community (see ***Equation 23***).

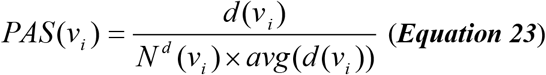

#### Breakdown point network

For each fusion PPI corresponding to a cancer sub-type, we generated the corresponding breakdown point network, identified from site-directed and bootstrap percolation. The steps for site-directed percolation are given in ***Algorithm 6***.

##### Algorithm 6 Breakdown Point Network

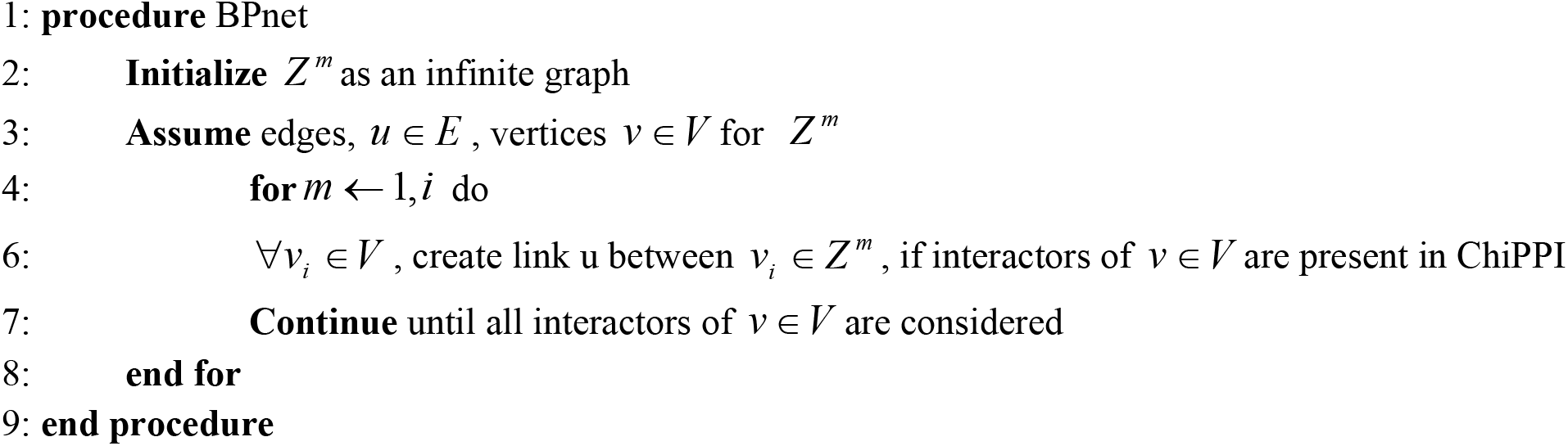

#### Gene and pathway enrichment

GSEA^38^ was originally developed to determine differential expression levels of a pre-defined gene set in two phenotypes. Genome-wide expression profiles from two-class samples were used to rank all genes in the data set. The ranking list was then used to calculate an ES and p-value. The procedure included obtaining the gene-ranking list, calculating an ES, estimating the significance level of the ES, and correcting the significance level for multiple gene sets. Given an *a priori* defined set of genes S (e.g., genes encoding products in a metabolic pathway, located in the same cytogenetic band, or sharing the same GO category), the goal of GSEA is to determine whether the members of S are randomly distributed throughout a list L or primarily located at the top or bottom. ES reflects the degree to which a gene set S is over-represented at the extremes of the entire ranked list L. The score is calculated by moving down the list L, increasing a running sum statistic where we encounter a gene in S, and decreasing it when we encounter genes not in S. The p-value of ES is determined by an empirical phenotype-based permutation test, where the phenotype labels are permuted, and ES is re-computed for the gene set, generating a null distribution for ES. The nominal p-value of the observed ES is then calculated, relative to this null distribution. This is followed by normalizing the ES for each gene set to account for the size of the set, yielding a normalized enrichment score. The false discovery rate was subsequently calculated to control the proportion of false positives.

#### Survival Analysis

The Kaplan–Meier estimator is a statistic, and several estimators are used to approximate its variance. Consider that T = failure time, C = censuring time, distribution function F, for density f, and distribution function G, for density g. We assume that C is independent of T. Thus, we define a function using ***Equations 24 and 25***.

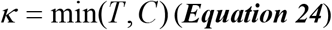

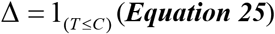

Thus, the density of observing *(k, 1)* is *f* (*k*)(1 − *G*(*k*)), whereas the density of observing *(x, 0)* is *g*(*k*)(1 − *F* (*k*)). Thus, the density of observing *(k, δ)* is

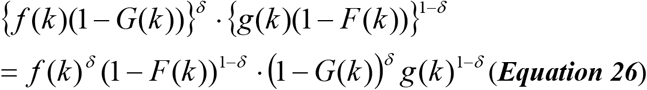

Thus, the likelihood for *F* and *G* of *n* observations (*k*_1_, *δ*_1_),…, (*k*_*n*_, *δ* _*n*_) is

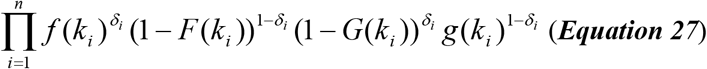

## ACKNOWLEDGMENTS

We acknowledge Dr. Dorith Raviv Shay and Dr. Naamah Bloch for reviewing this manuscript. This work was supported by a PBC (VATAT) Fellowship for outstanding Post-Docs from China and India to S.T. (grants 20027, 22351, 26912, 35174).

## AUTHOR CONTRIBUTIONS

S.T. and M.F.-M. conceived the study, S.T. performed the analysis and both authors approved the manuscript and provided strategic oversight of the work.

