## Supplementary Data for "Improving patient survival by direct targeting of chimeric protein-protein interaction networks"

1. Figure S1a-2b: Network-Category for Training phase (150 fusions), Testing phase (3091 fusions)
2. Figure S2a-S2d: Overall distribution and statistics of higher and lower-degree hubs in LK, LY, ME and GL, SC and CA.
3. Figure S3: an overview of the breakdown points in LK, LY, ME and GL, SC and CA that demonstrate the robust nature of PPIs (training phase).
4. Figure S4a-b: An overview of the community distribution in LK, LY, ME and GL, SC and CA fusions.
5. Figure S5: A comparative analysis of communities in RUNX1-RUNX1T1 (LK, LY, ME and GL), EWSR1-ERG (SC) and BCAS3-BCAS4 (CA).
6. Figure S6: The preferential attachment model implementation on EWSR1-ERG fusion PPI (SC)
7. Figure S7: The preferential attachment model implementation on BCAS3-BCAS4 fusion PPI (CA).
8. Figure S8a-b: Network diameter, number of essential communities, number of essential community vertices, number of preferential attachment vertices; total degree of essential community vertices, total cc of essential community vertices, betweenness centrality of essential community vertices, total PAS
9. Figure S9a-c: GSEA analysis for major cancer sub-types under LK, LY, CA
10. Figure S10a-c: Pathway enrichment maps for LK, LY, ME.
11. Figure S11a-b: Survival plots for LK, LY
12. Figure S12a-d: Drug binding site networks for CLLE-ES (LK), MALY-DE (LY), OV-AU (CA), BRCA-US (CA)
13. Table S1: List of fusion and parental proteins (Training phase)
14. Table S2: K, LY, ME, GL- Network categorization, scalefree_hier_rand (TRAINING)
15. Table S3: SC-Network categorization, scalefree_hier_rand (TRAINING)
16. Table S4: CA- Network categorization, scalefree_hier_rand (TRAINING)
17. Table S5: LK, LY, ME, GL-Hubs (TRAINING)
18. Table S6: SC-Hubs (TRAINING)
19. Table S7: CA-Hubs (TRAINING)
20. Table S8: LK, LY, ME, GL_Breakdown (TRAINING)
21. Table S9: SC_Breakdown (TRAINING)
22. Table S10: CA_Breakdown (TRAINING)
23. Table S11: LK, LY, ME, GL-community (TRAINING)
24. Table S12: SC-community (TRAINING)
25. Table S13: CA-community (TRAINING)
26. Table S14: LK, LY, ME, GL_Preferential_attachment (TRAINING)
27. Table S15: SC_Preferential_attachment (TRAINING)
28. Table S16: CA_Preferential_attachment (TRAINING)
29. Table S17: LK, LY, ME, GL-PAS (TRAINING)
30. Table S18: SC-PAS (TRAINING)
31. Table S19: ST-PAS (TRAINING)

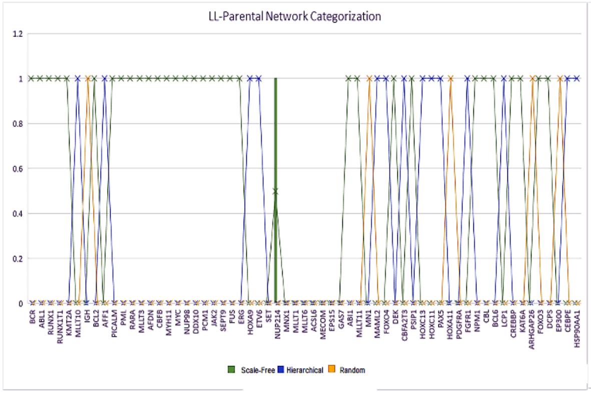

(a)

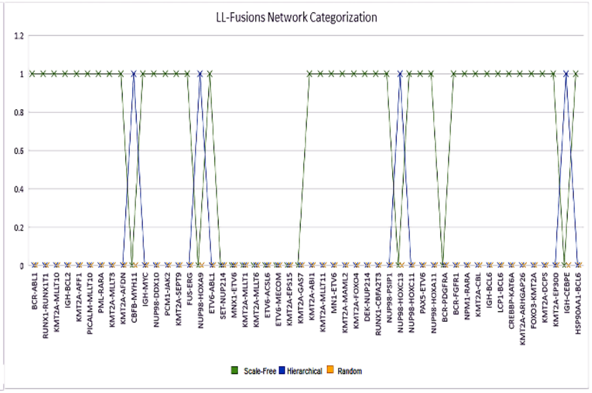

(b)

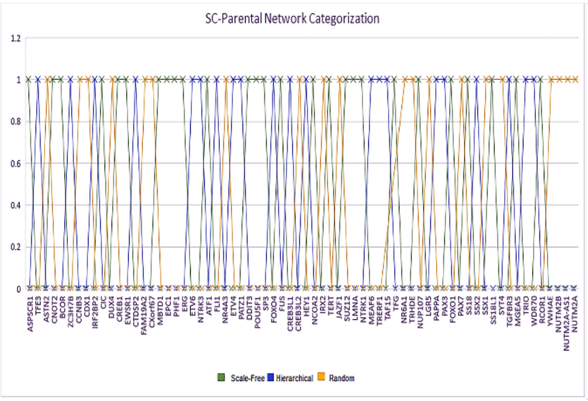

(c)

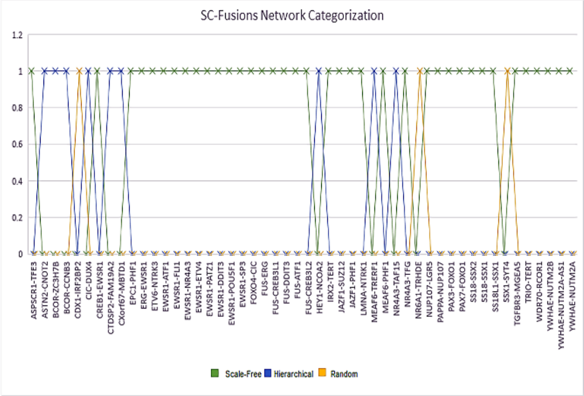

(d)

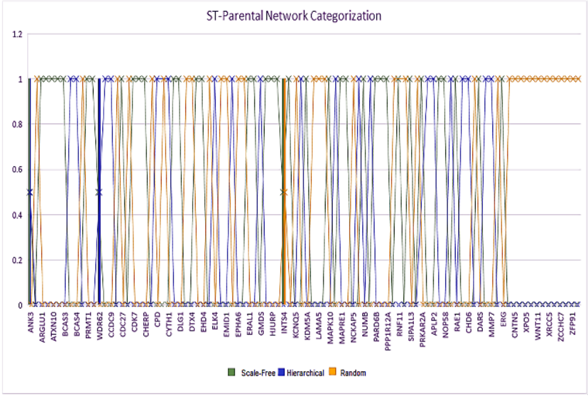

(e)

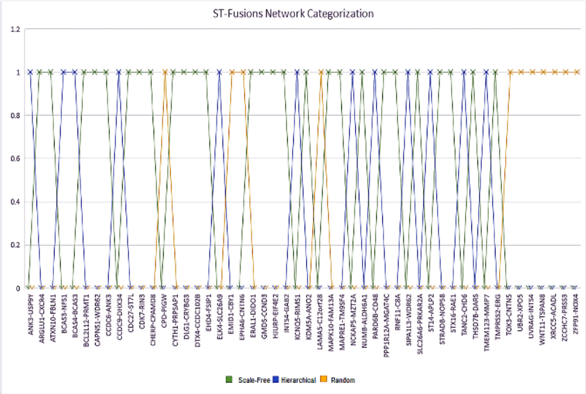

(f)

**Figure S1a: Network-Category for Training phase (150 fusions)**

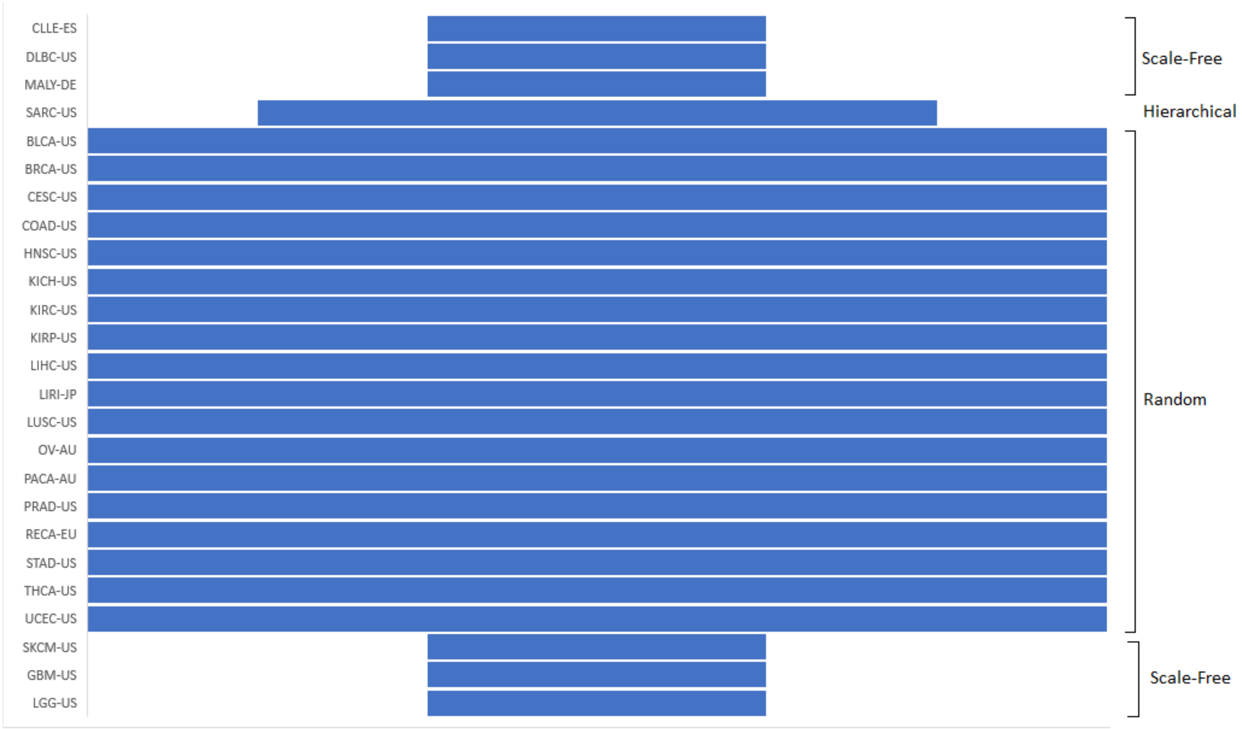

**Figure S1b: Network-Category for Test phase (3091 fusions)**

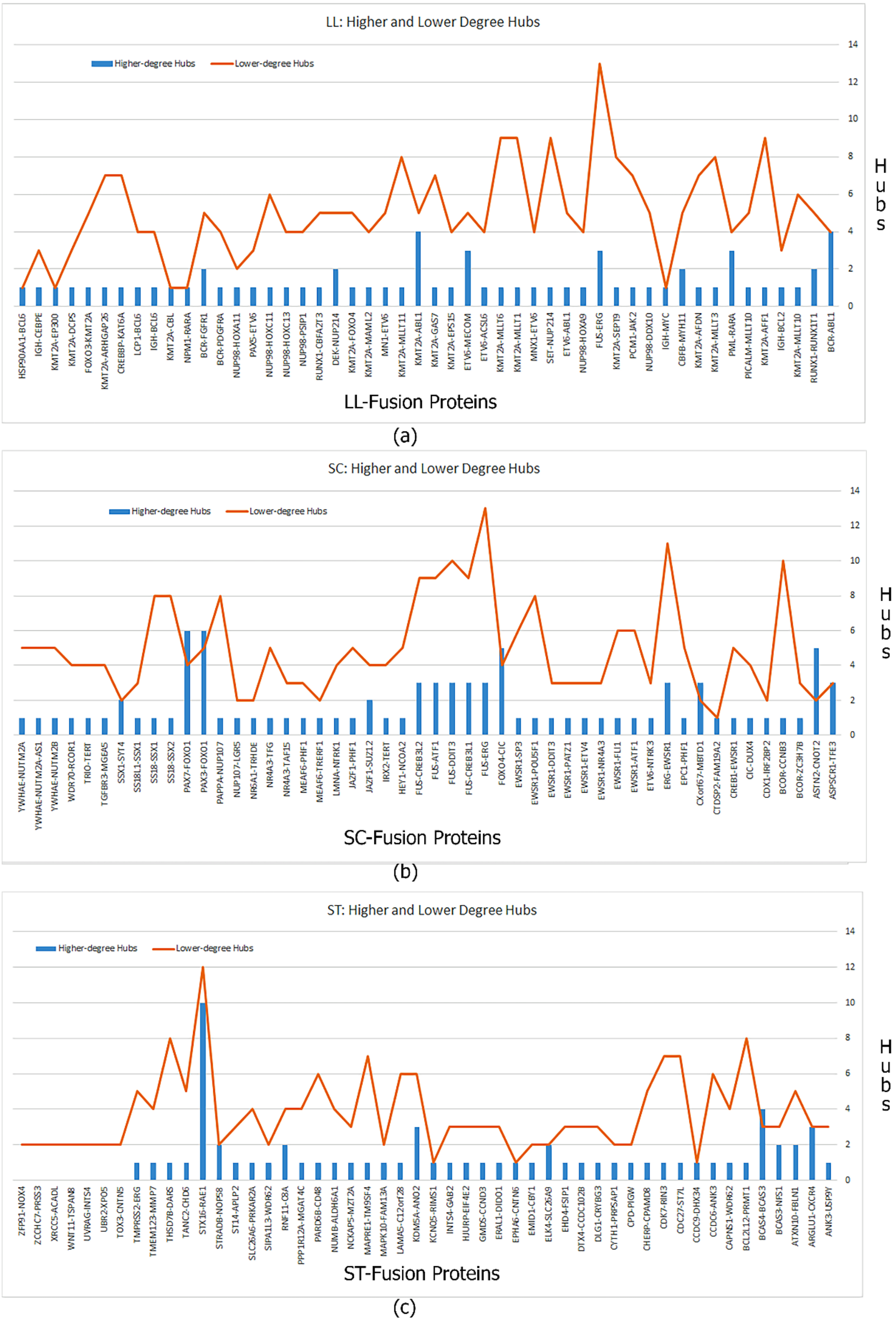

**Figure S2a: Hub distribution (training)**

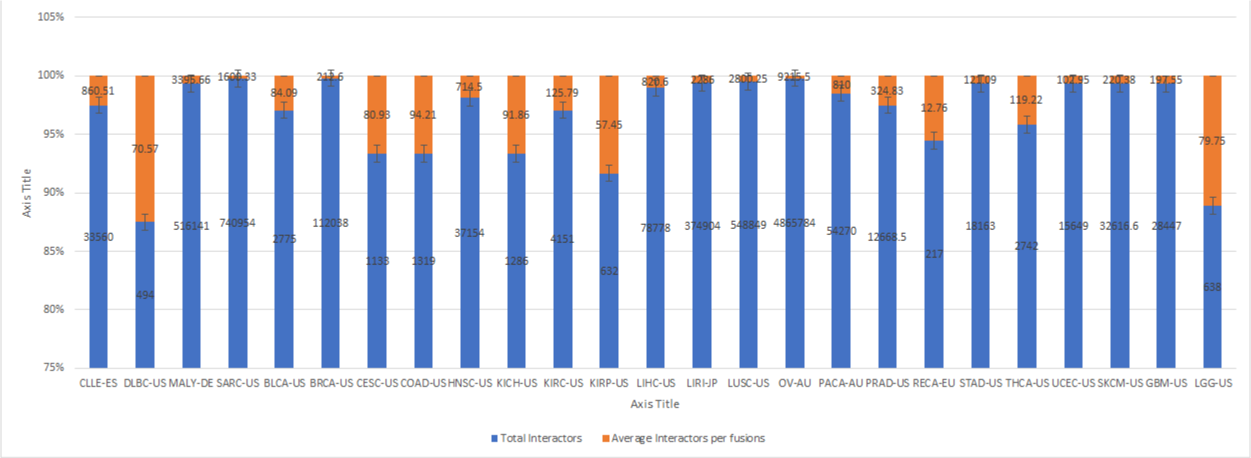

**Figure S2b: Total interactors, avg. interactors per fusion (672 aliquot)**

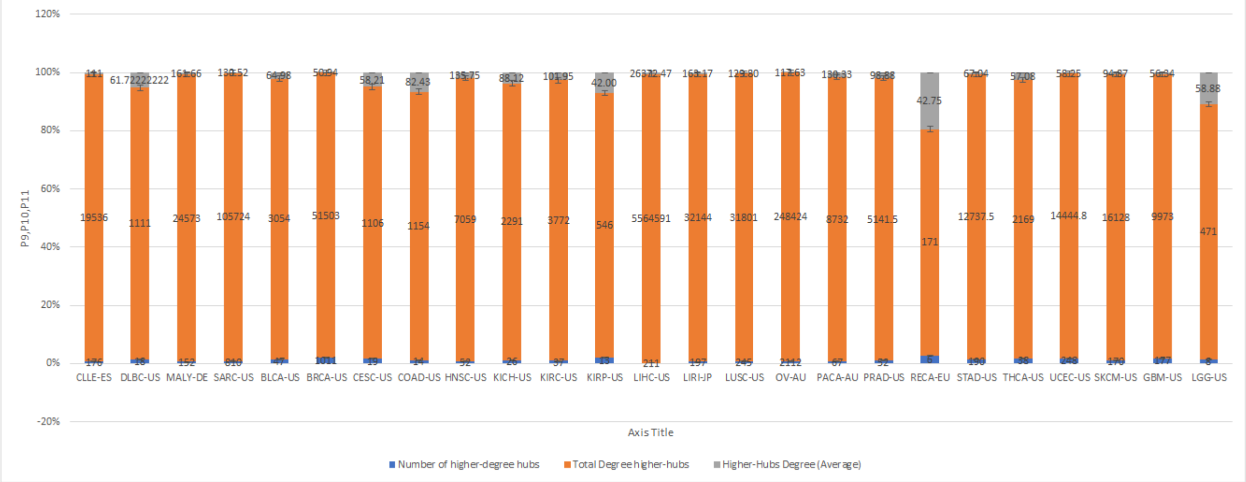

**Figure S2c: No., total degree and avg. degree of higher degree-hubs (672 aliquot)**

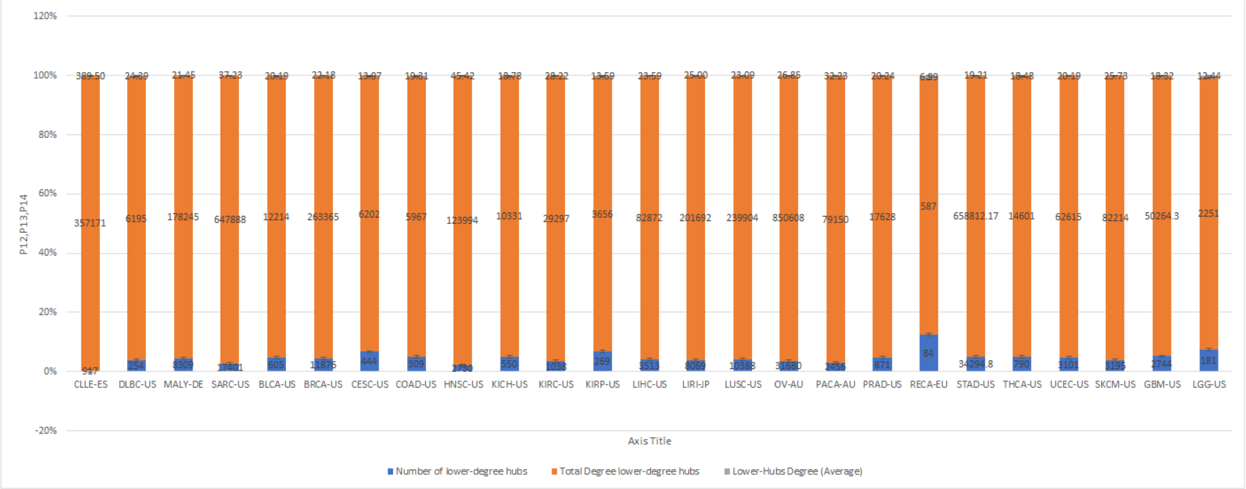

**Figure S2d: No., total degree and avg. degree of lower degree-hubs (672 aliquot)**

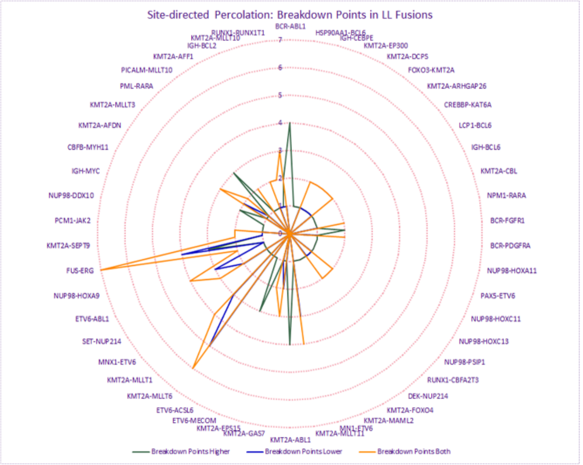

(a)

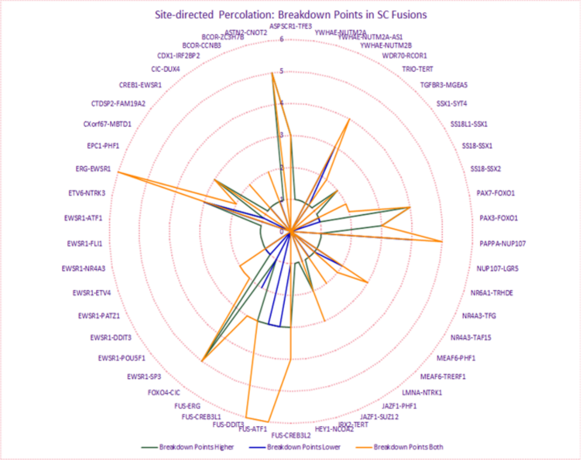

(b)

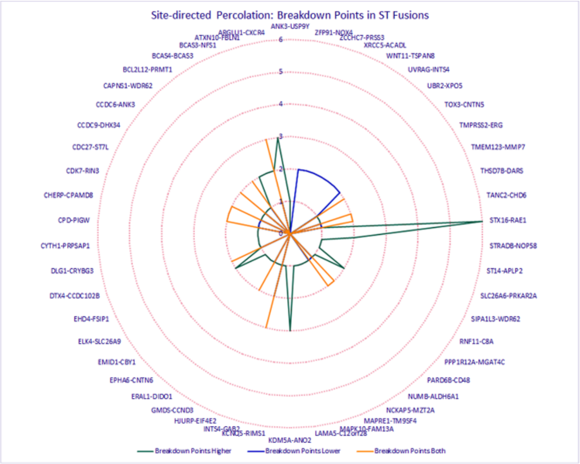

(c)

**Figure S3: Site-directed percolation, breakdown points in the training phase**

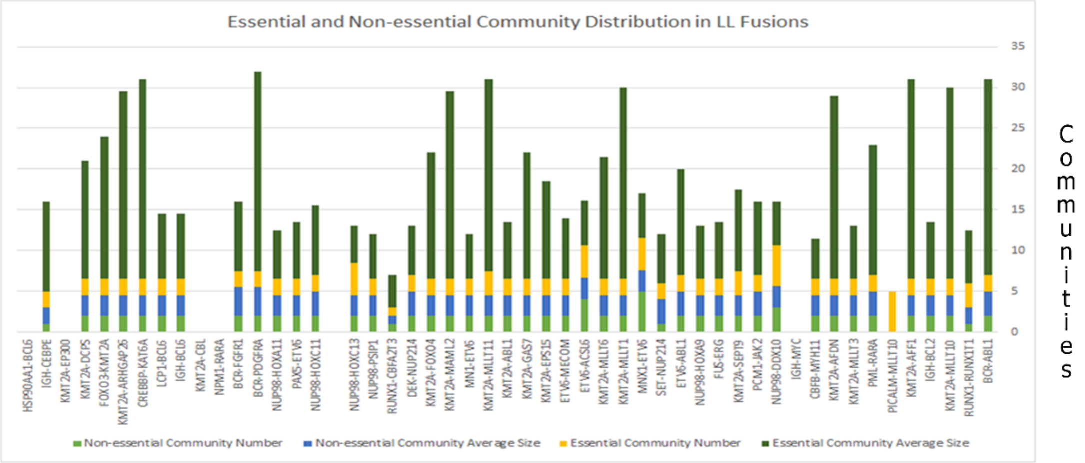

(a)

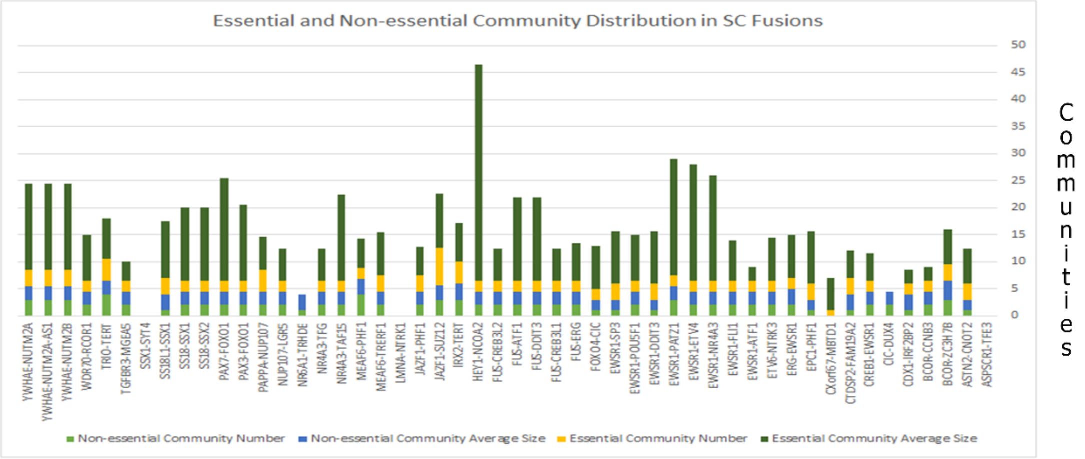

(b)

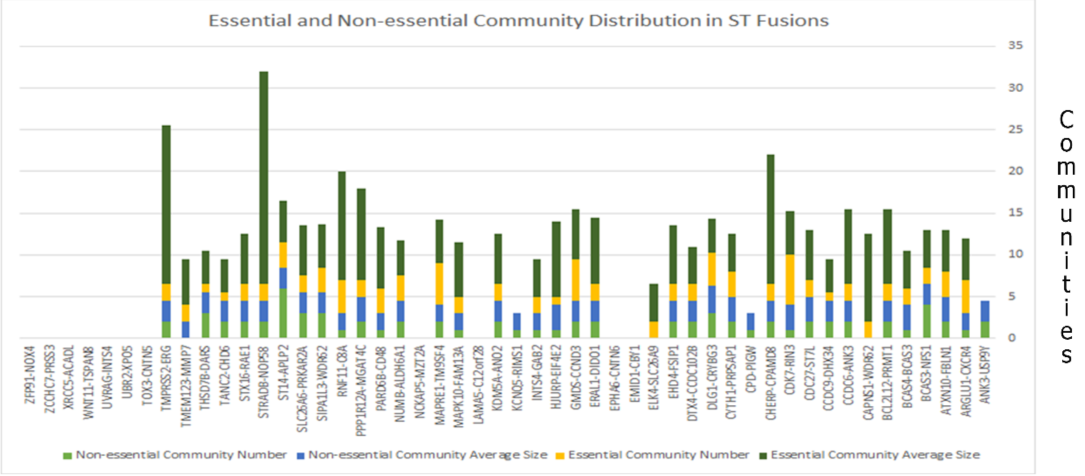

(c)

**Figure S4a: Community distribution (Training)**

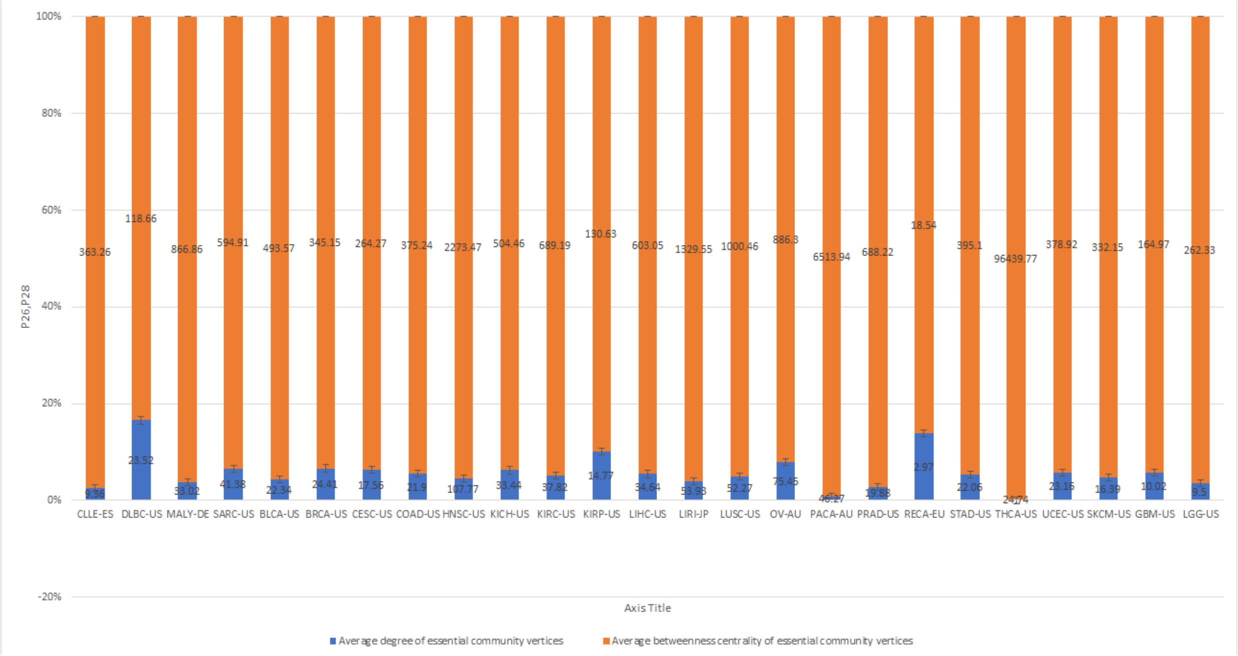

**Figure S4b: Community distribution (672 aliquot)**

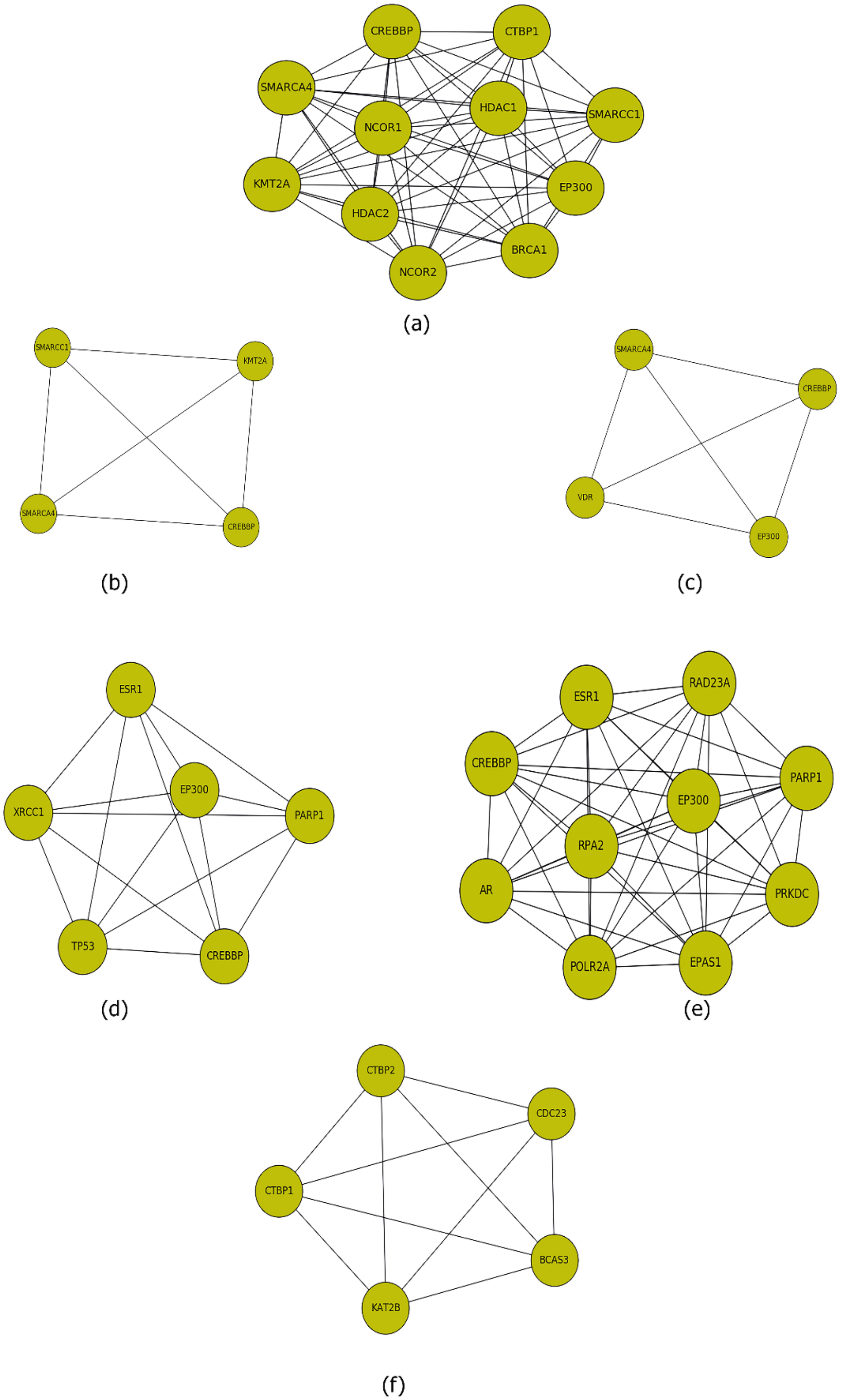

**Figure S5: Essential communities from bootstrap percolation in RUNX1-RUNX1T1 (a-c), ERG-EWSR1 (d-e), BCAS3-BCAS4 (f)**

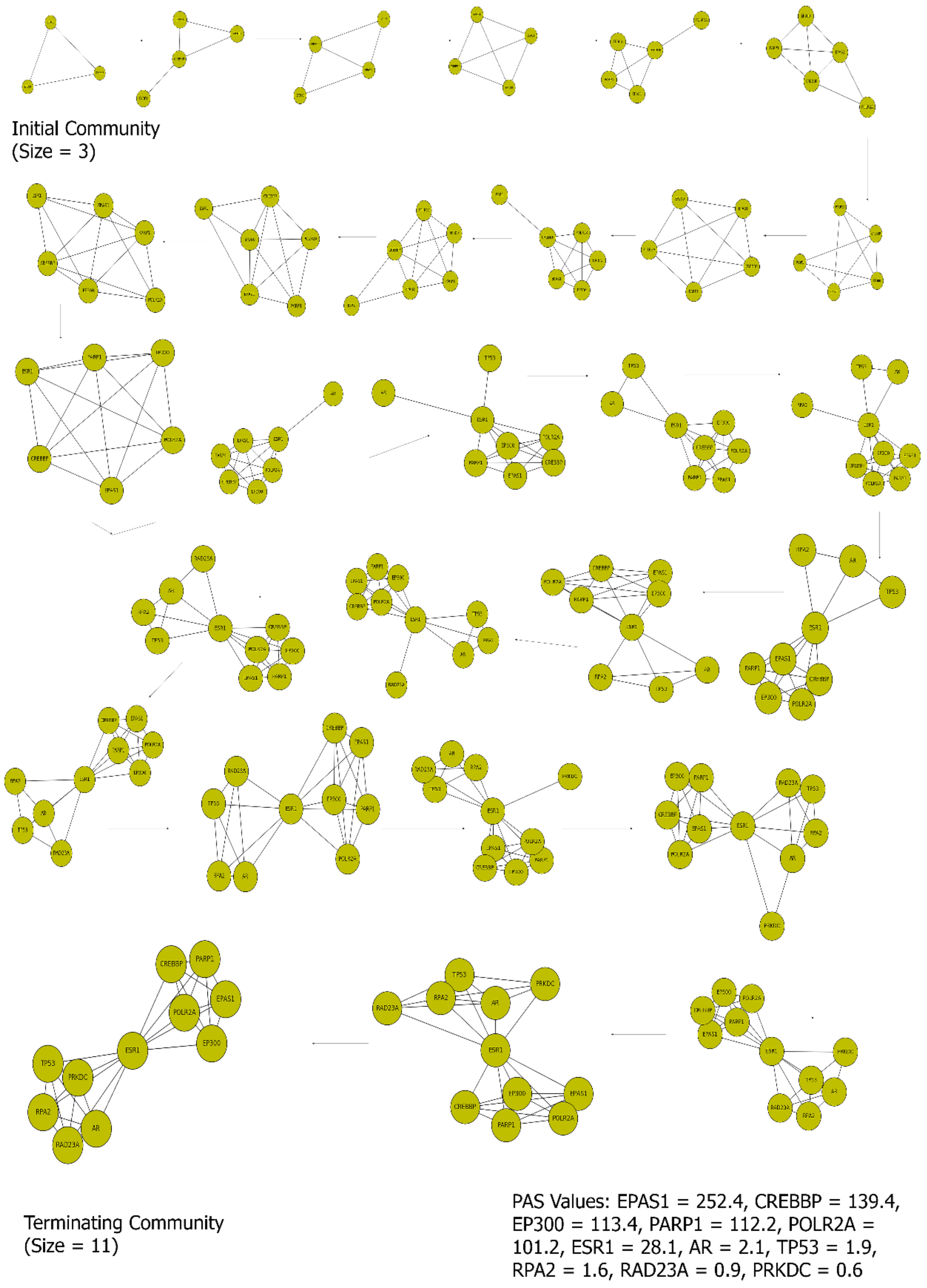

**Figure S6: Preferential attachment ERG-EWSR1 (SC)**

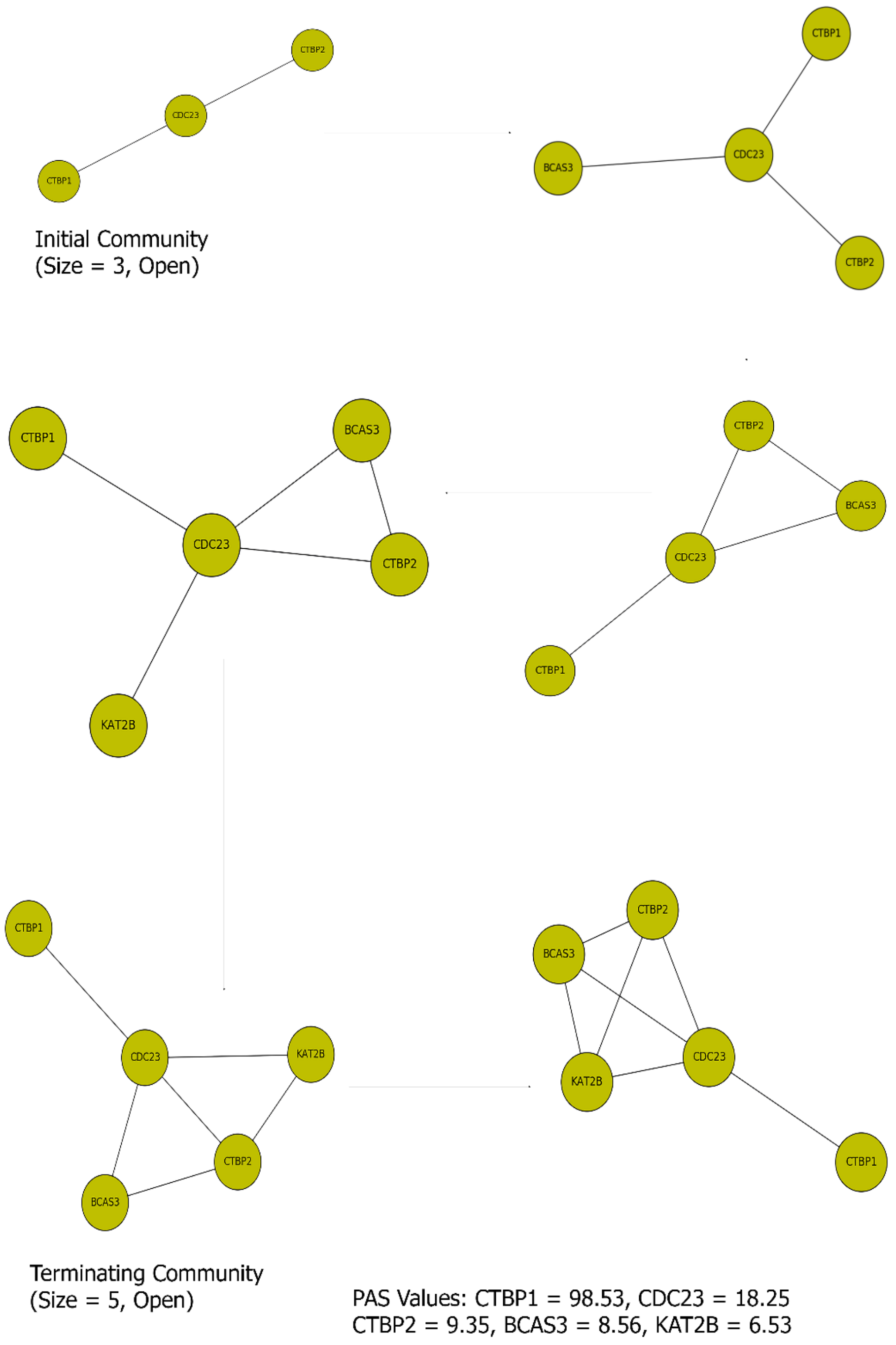

**Figure S7: Preferential attachment BCAS3-BCAS4 (CA)**

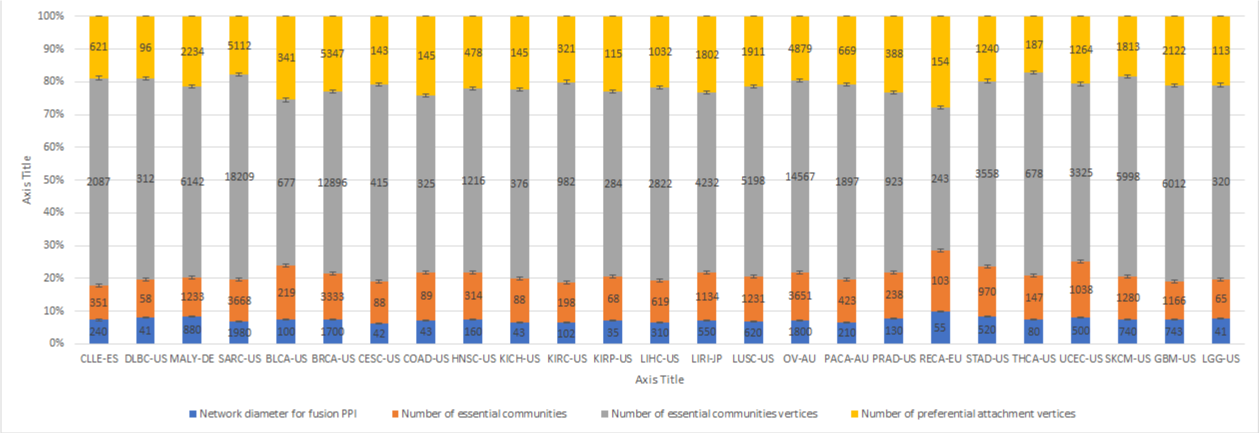

**Figure S8a: Network diameter, number of essential communities, number of essential community vertices, number of preferential attachment vertices**

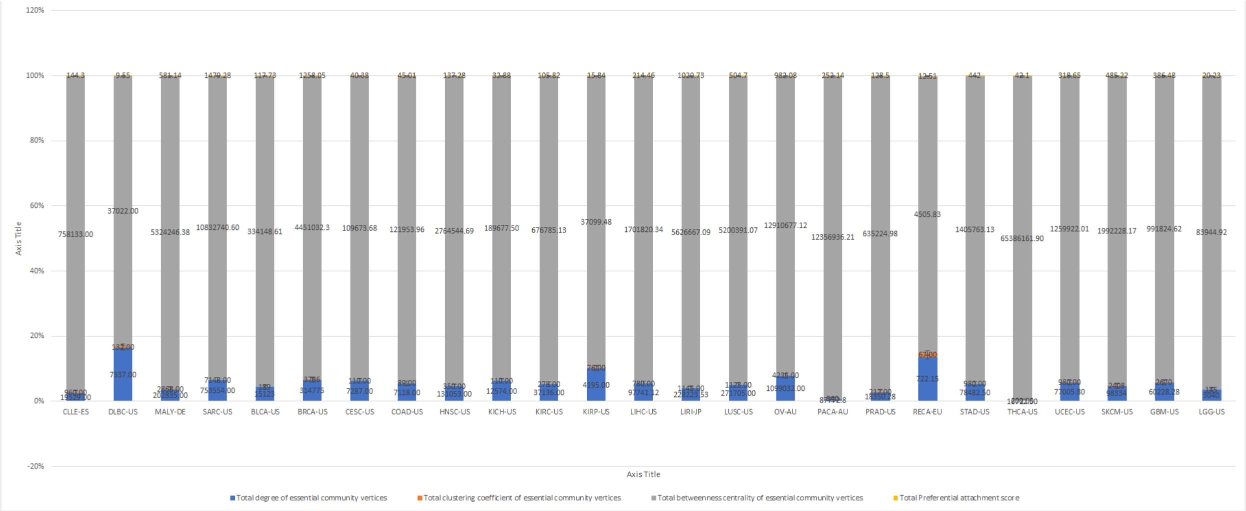

**Figure S8b: Total degree of essential community vertices, total cc of essential community vertices, betweenness centrality of essential community vertices, total PAS**

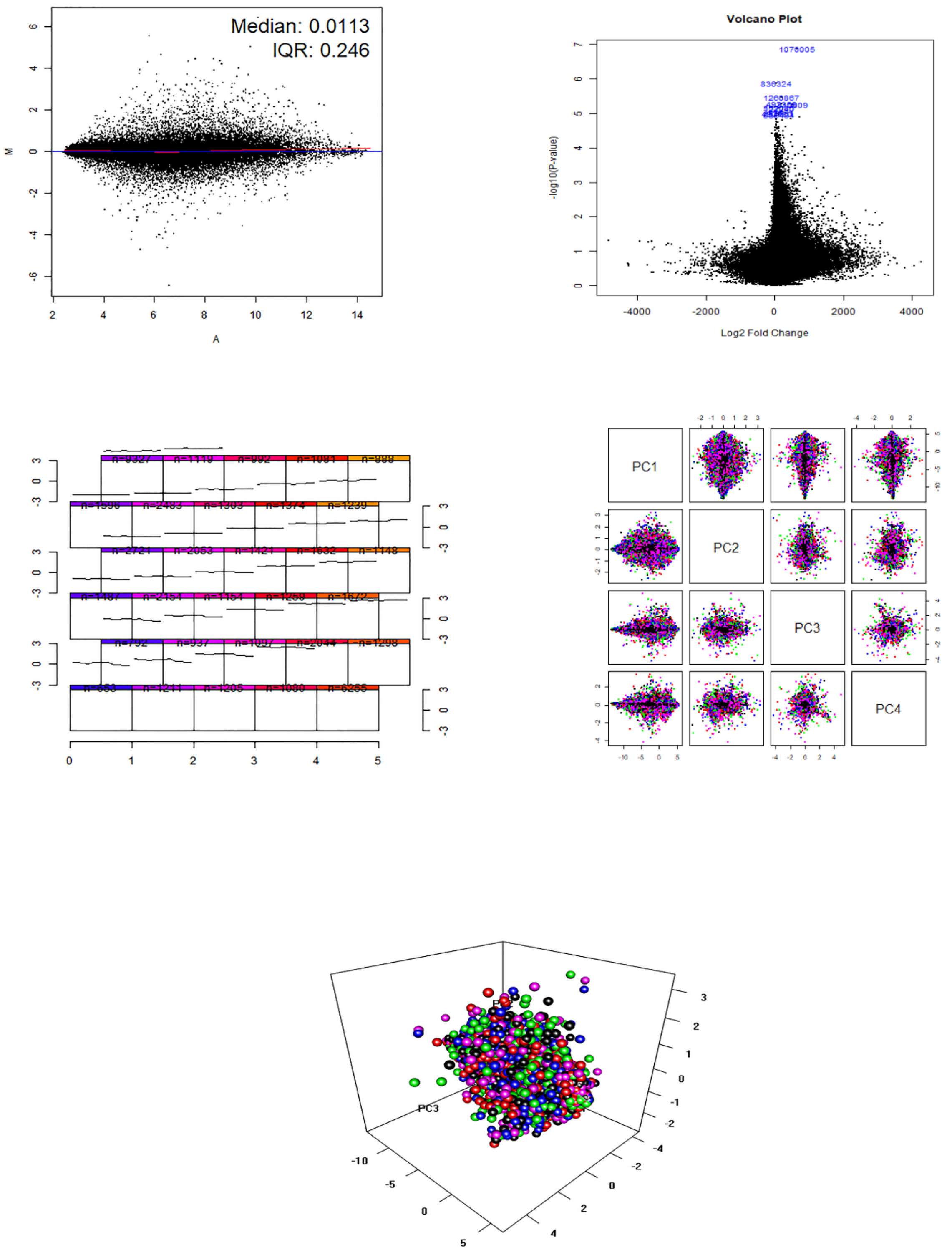

CLLE-ES

**Figure S9a: GSEA, SOM, PCA for CLLE-ES (LK)**

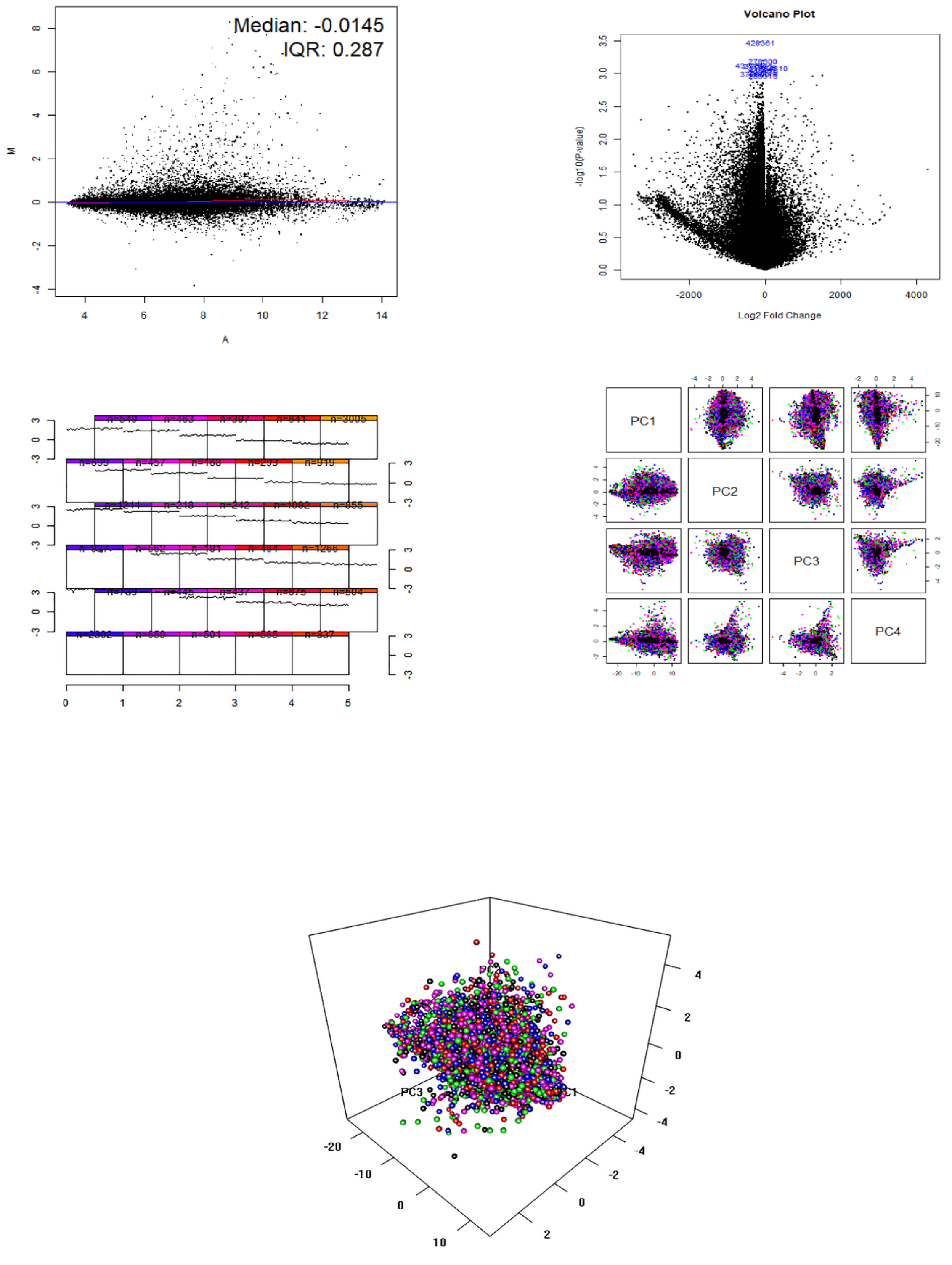

MALY-DE

**Figure S9b: GSEA, SOM, PCA for MALY-DE (LY)**

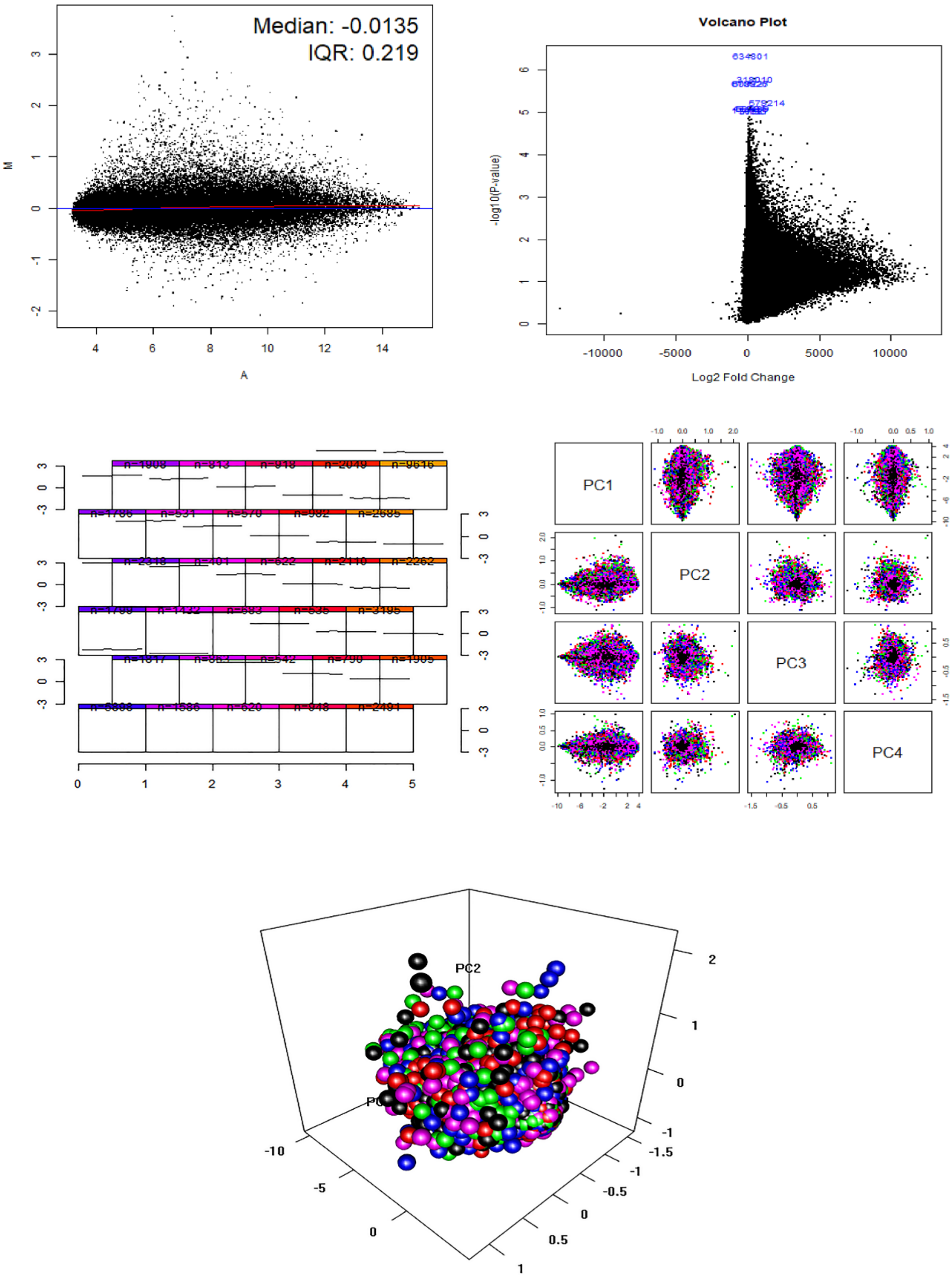

OV-AU

**Figure S9c: GSEA, SOM, PCA for OV-AU (CA)**

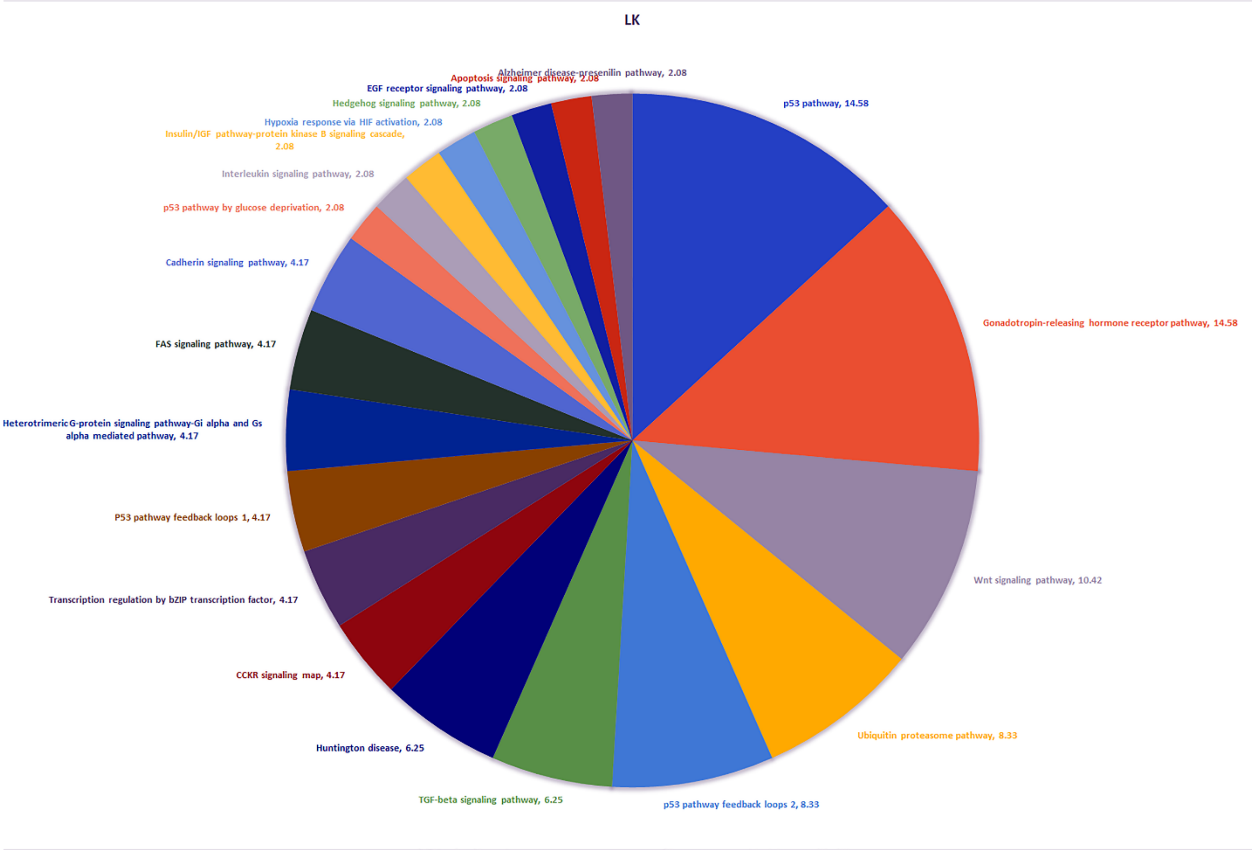

**Figure S10a: Pathway enrichment map for LK**

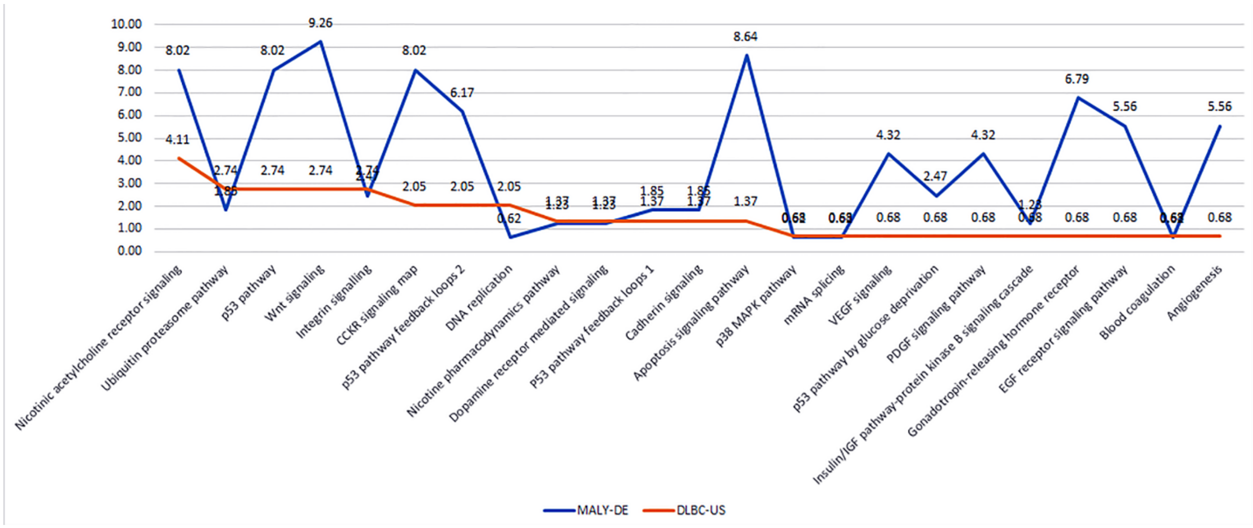

**Figure S10b: Pathway enrichment map for LY**

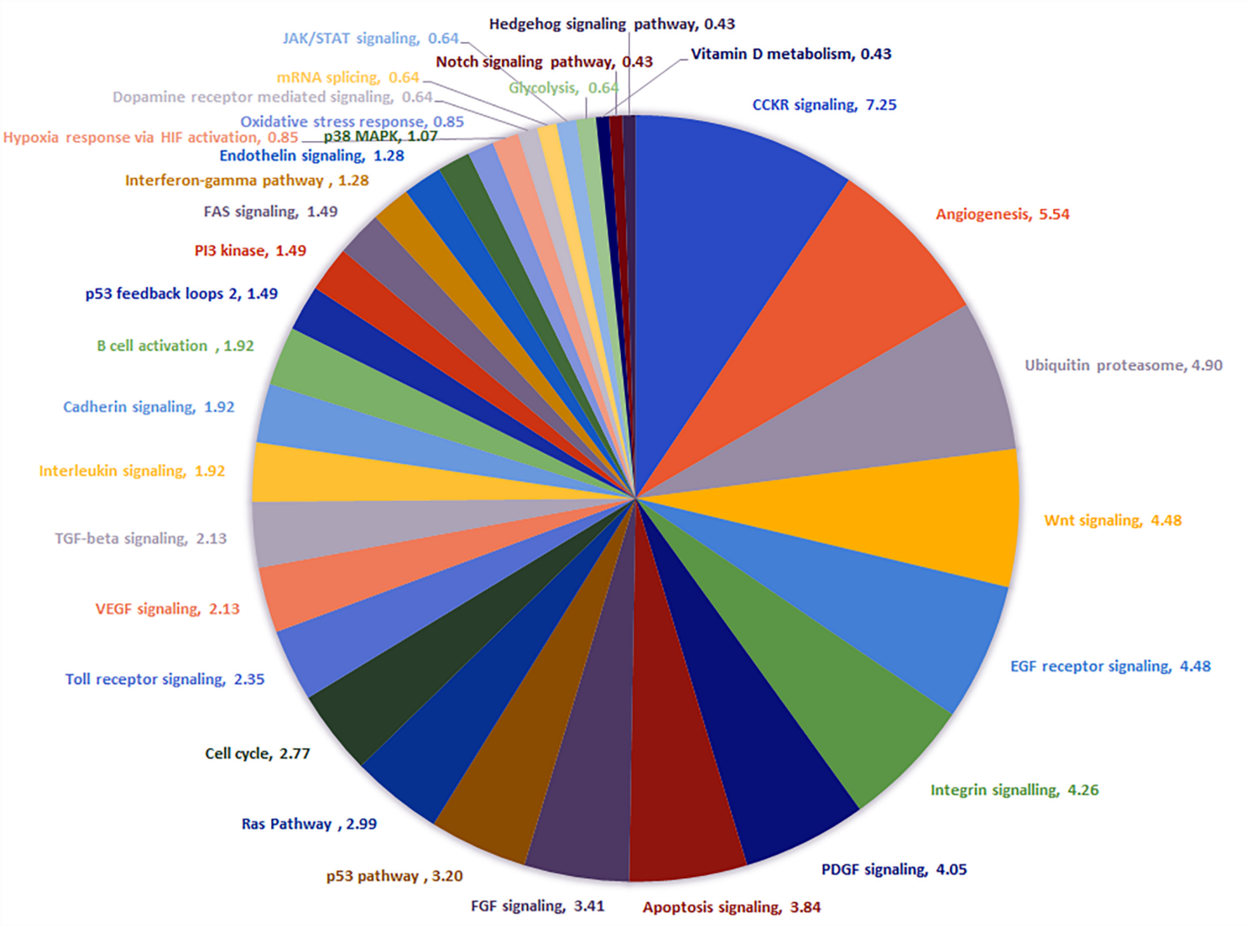

**Figure S10c: Pathway enrichment map for ME**

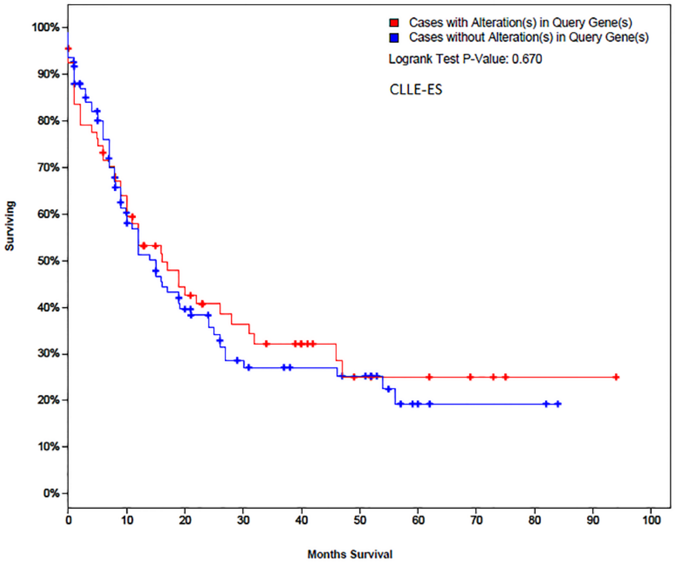

**Figure S11a: Survival Plot for LK**

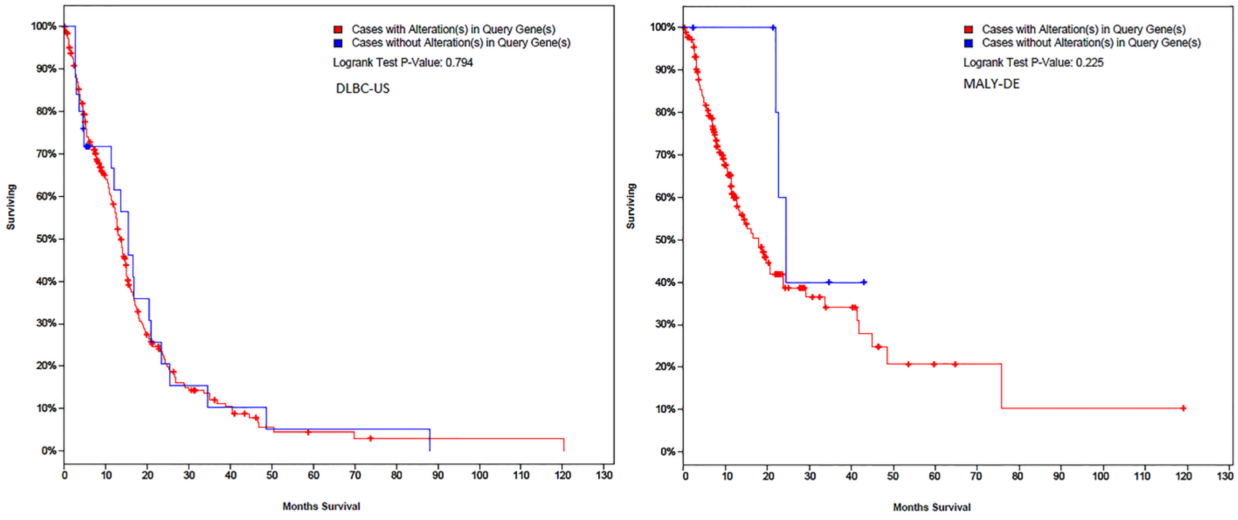

**Figure S11b: Survival Plot for LY**

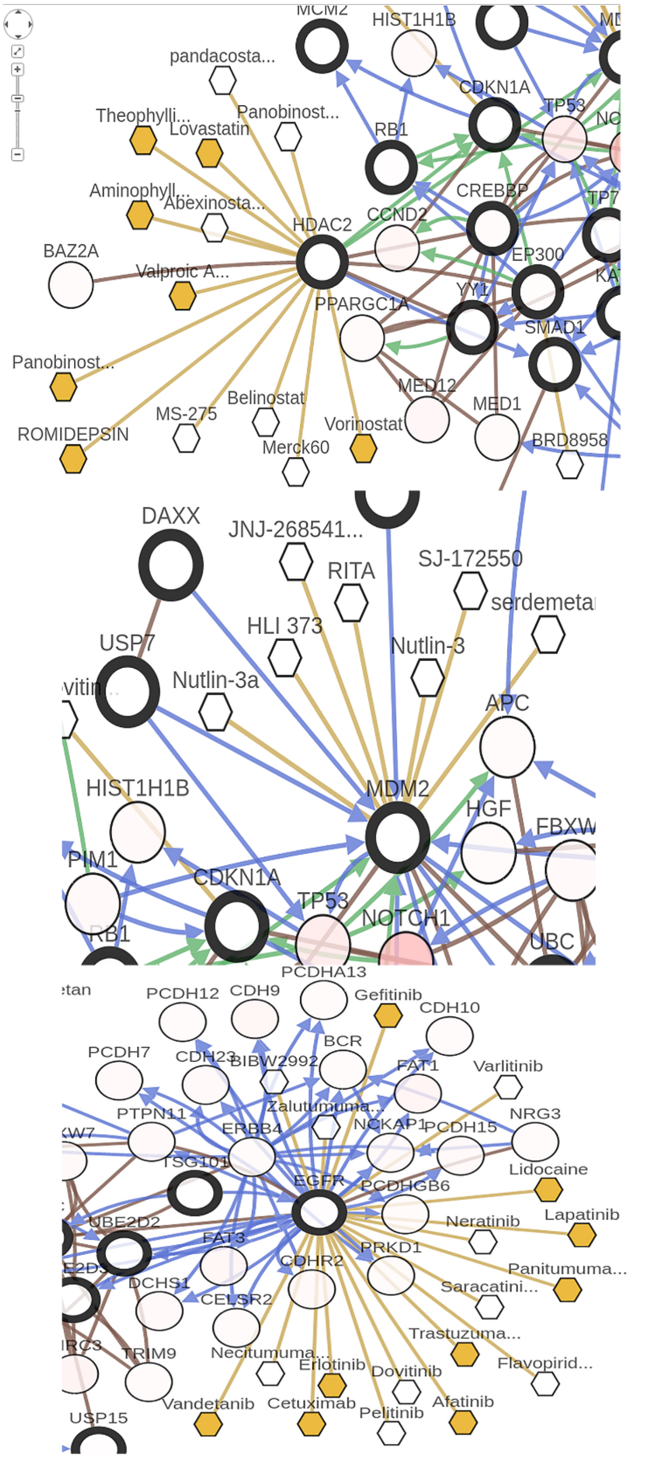

CLLE-ES: Drug binding sites

**Figure S12a: Survival Plot for CLLE-ES (LK)**

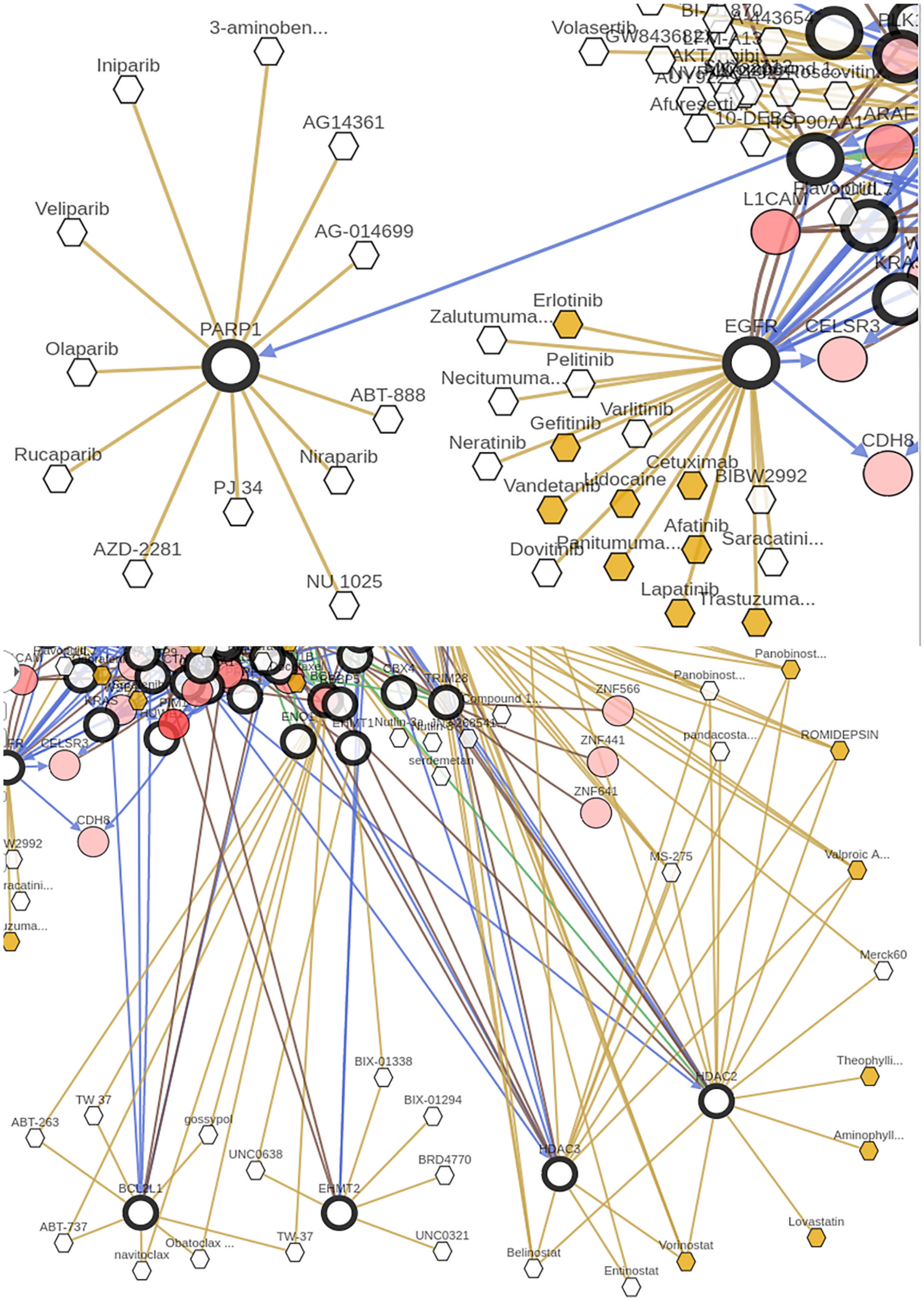

MALY-DE: Drug binding sites

**Figure S12b: Survival Plot for MALY-DE (LY)**

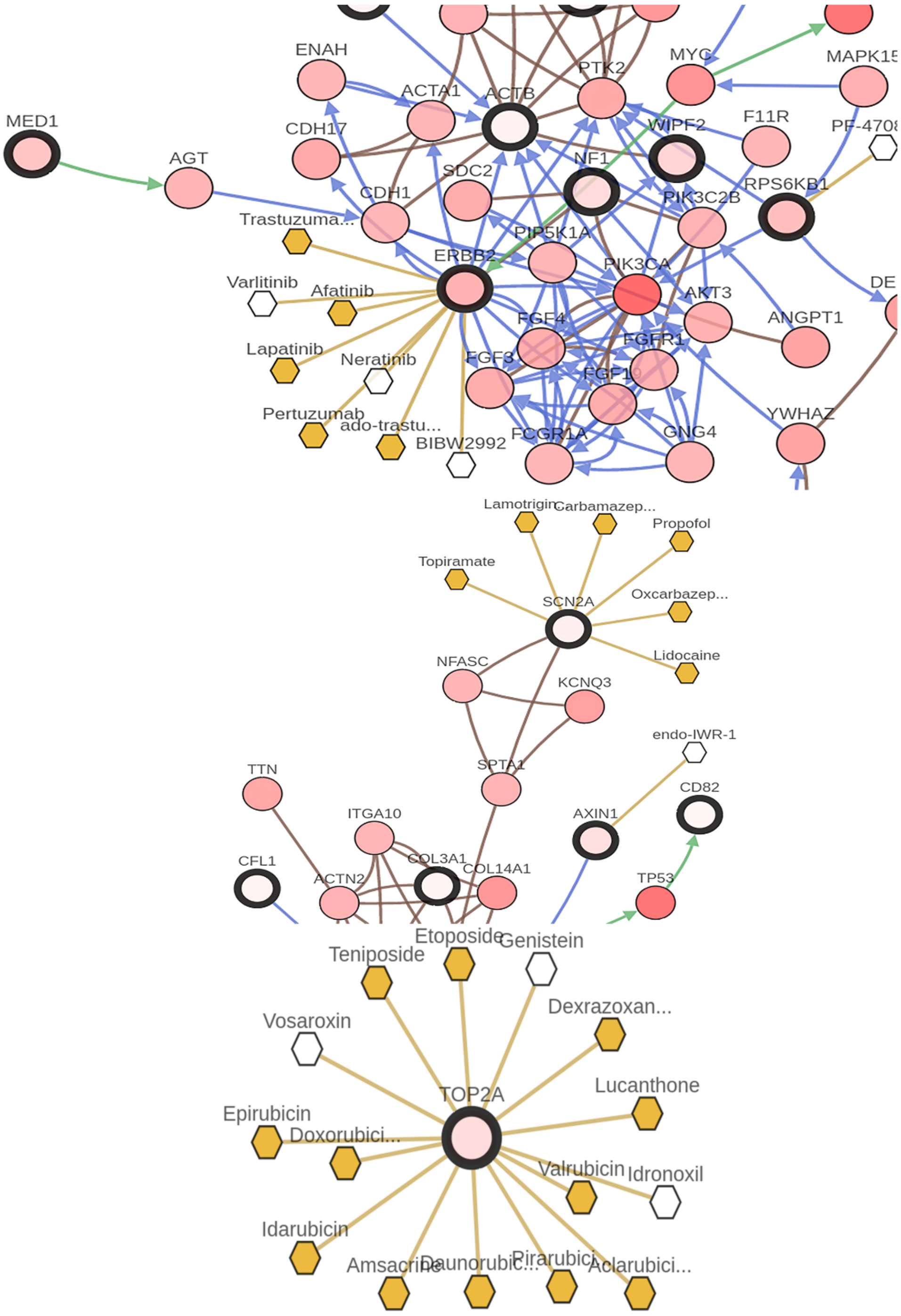

BRCA-US: drug binding sites

**Figure S12c: Survival Plot for OV-AU (CA)**

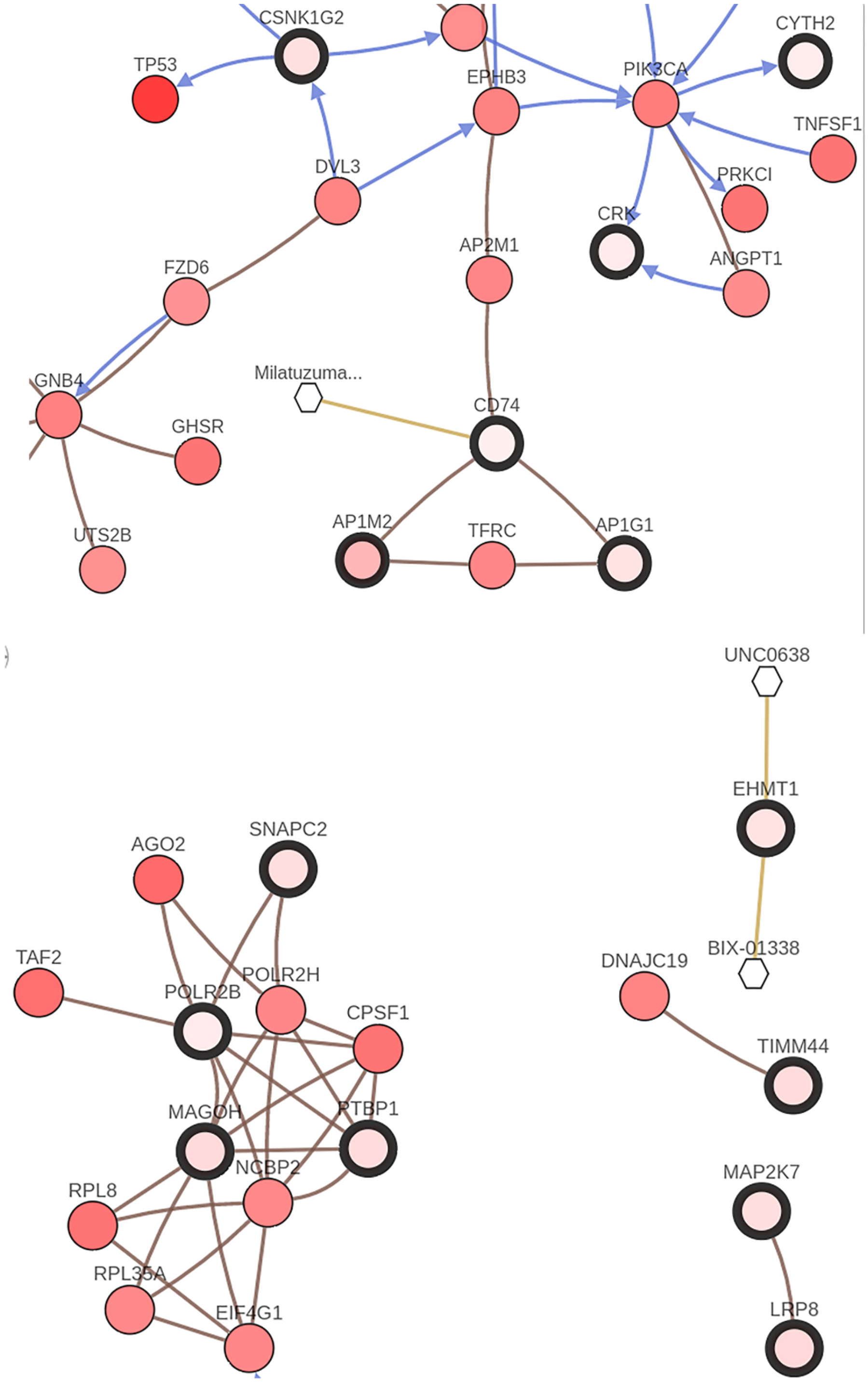

OV-AU: drug binding sites

**Figure S12d: Survival Plot for BRCA-US (CA)**

**Table S1a: List of fusions and parental proteins (Training phase)**

| LK, LY, ME, GL | SC | CA |
|---|---|---|
| BCR | ASPSCR1 | ANK3 |
| ABL1 | TFE3 | USP9Y |
| BCR-ABL1 | ASPSCR1-TFE3 | ANK3-USP9Y |
| RUNX1 | ASTN2 | ARGLU1 |
| RUNX1T1 | CNOT2 | CXCR4 |
| RUNX1-RUNX1T1 | ASTN2-CNOT2 | ARGLU1-CXCR4 |
| KMT2A | BCOR | ATXN10 |
| MLLT10 | ZC3H7B | FBLN1 |
| KMT2A-MLLT10 | BCOR-ZC3H7B | ATXN10-FBLN1 |
| IGH | BCOR | BCAS3 |
| BCL2 | CCNB3 | NFS1 |
| IGH-BCL2 | BCOR-CCNB3 | BCAS3-NFS1 |
| KMT2A | CDX1 | BCAS4 |
| AFF1 | IRF2BP2 | BCAS3 |
| KMT2A-AFF1 | CDX1-IRF2BP2 | BCAS4-BCAS3 |
| PICALM | CIC | BCL2L12 |
| MLLT10 | DUX4 | PRMT1 |
| PICALM-MLLT10 | CIC-DUX4 | BCL2L12-PRMT1 |
| PML | CREB1 | CAPNS1 |
| RARA | EWSR1 | WDR62 |
| PML-RARA | CREB1-EWSR1 | CAPNS1-WDR62 |
| KMT2A | CTDSP2 | CCDC6 |
| MLLT3 | FAM19A2 | ANK3 |
| KMT2A-MLLT3 | CTDSP2-FAM19A2 | CCDC6-ANK3 |
| KMT2A | CXorf67 | CCDC9 |
| AFDN | MBTD1 | DHX34 |
| KMT2A-AFDN | CXorf67-MBTD1 | CCDC9-DHX34 |
| CBFB | EPC1 | CDC27 |
| MYH11 | PHF1 | ST7L |
| CBFB-MYH11 | EPC1-PHF1 | CDC27-ST7L |
| IGH | ERG | CDK7 |
| MYC | EWSR1 | RIN3 |
| IGH-MYC | ERG-EWSR1 | CDK7-RIN3 |
| NUP98 | ETV6 | CHERP |
| DDX10 | NTRK3 | CPAMD8 |
| NUP98-DDX10 | ETV6-NTRK3 | CHERP-CPAMD8 |
| PCM1 | EWSR1 | CPD |
| JAK2 | ATF1 | PIGW |
| PCM1-JAK2 | EWSR1-ATF1 | CPD-PIGW |
| KMT2A | EWSR1 | CYTH1 |

| | | |
|---|---|---|
| SEPT9 | FLI1 | PRPSAP1 |
| KMT2A-SEPT9 | EWSR1-FLI1 | CYTH1-PRPSAP1 |
| FUS | EWSR1 | DLG1 |
| ERG | NR4A3 | CRYBG3 |
| FUS-ERG | EWSR1-NR4A3 | DLG1-CRYBG3 |
| NUP98 | EWSR1 | DTX4 |
| HOXA9 | ETV4 | CCDC102B |
| NUP98-HOXA9 | EWSR1-ETV4 | DTX4-CCDC102B |
| ETV6 | EWSR1 | EHD4 |
| ABL1 | PATZ1 | FSIP1 |
| ETV6-ABL1 | EWSR1-PATZ1 | EHD4-FSIP1 |
| SET | EWSR1 | ELK4 |
| NUP214 | DDIT3 | SLC26A9 |
| SET-NUP214 | EWSR1-DDIT3 | ELK4-SLC26A9 |
| MNX1 | EWSR1 | EMID1 |
| ETV6 | POU5F1 | CBY1 |
| MNX1-ETV6 | EWSR1-POU5F1 | EMID1-CBY1 |
| KMT2A | EWSR1 | EPHA6 |
| MLLT1 | SP3 | CNTN6 |
| KMT2A-MLLT1 | EWSR1-SP3 | EPHA6-CNTN6 |
| KMT2A | FOXO4 | ERAL1 |
| MLLT6 | CIC | DIDO1 |
| KMT2A-MLLT6 | FOXO4-CIC | ERAL1-DIDO1 |
| ETV6 | FUS | GMDS |
| ACSL6 | ERG | CCND3 |
| ETV6-ACSL6 | FUS-ERG | GMDS-CCND3 |
| ETV6 | FUS | HJURP |
| MECOM | CREB3L1 | EIF4E2 |
| ETV6-MECOM | FUS-CREB3L1 | HJURP-EIF4E2 |
| KMT2A | FUS | INTS4 |
| EPS15 | DDIT3 | GAB2 |
| KMT2A-EPS15 | FUS-DDIT3 | INTS4-GAB2 |
| KMT2A | FUS | KCNQ5 |
| GAS7 | ATF1 | RIMS1 |
| KMT2A-GAS7 | FUS-ATF1 | KCNQ5-RIMS1 |
| KMT2A | FUS | KDM5A |
| ABI1 | CREB3L2 | ANO2 |
| KMT2A-ABI1 | FUS-CREB3L2 | KDM5A-ANO2 |
| KMT2A | HEY1 | LAMA5 |
| MLLT11 | NCOA2 | C12orf28 |
| KMT2A-MLLT11 | HEY1-NCOA2 | LAMA5-C12orf28 |
| MN1 | IRX2 | MAPK10 |
| ETV6 | TERT | FAM13A |
| MN1-ETV6 | IRX2-TERT | MAPK10-FAM13A |

| | | |
|---|---|---|
| KMT2A | JAZF1 | MAPRE1 |
| MAML2 | SUZ12 | TM9SF4 |
| KMT2A-MAML2 | JAZF1-SUZ12 | MAPRE1-TM9SF4 |
| KMT2A | JAZF1 | NCKAP5 |
| FOXO4 | PHF1 | MZT2A |
| KMT2A-FOXO4 | JAZF1-PHF1 | NCKAP5-MZT2A |
| DEK | LMNA | NUMB |
| NUP214 | NTRK1 | ALDH6A1 |
| DEK-NUP214 | LMNA-NTRK1 | NUMB-ALDH6A1 |
| RUNX1 | MEAF6 | PARD6B |
| CBFA2T3 | TRERF1 | CD48 |
| RUNX1-CBFA2T3 | MEAF6-TRERF1 | PARD6B-CD48 |
| NUP98 | MEAF6 | PPP1R12A |
| PSIP1 | PHF1 | MGAT4C |
| NUP98-PSIP1 | MEAF6-PHF1 | PPP1R12A-MGAT4C |
| NUP98 | NR4A3 | RNF11 |
| HOXC13 | TAF15 | C8A |
| NUP98-HOXC13 | NR4A3-TAF15 | RNF11-C8A |
| NUP98 | NR4A3 | SIPA1L3 |
| HOXC11 | TFG | WDR62 |
| NUP98-HOXC11 | NR4A3-TFG | SIPA1L3-WDR62 |
| PAX5 | NR6A1 | SLC26A6 |
| ETV6 | TRHDE | PRKAR2A |
| PAX5-ETV6 | NR6A1-TRHDE | SLC26A6-PRKAR2A |
| NUP98 | NUP107 | ST14 |
| HOXA11 | LGR5 | APLP2 |
| NUP98-HOXA11 | NUP107-LGR5 | ST14-APLP2 |
| BCR | PAPPA | STRADB |
| PDGFRA | NUP107 | NOP58 |
| BCR-PDGFRA | PAPPA-NUP107 | STRADB-NOP58 |
| BCR | PAX3 | STX16 |
| FGFR1 | FOXO1 | RAE1 |
| BCR-FGFR1 | PAX3-FOXO1 | STX16-RAE1 |
| NPM1 | PAX7 | TANC2 |
| RARA | FOXO1 | CHD6 |
| NPM1-RARA | PAX7-FOXO1 | TANC2-CHD6 |
| KMT2A | SS18 | THSD7B |
| CBL | SSX2 | DARS |
| KMT2A-CBL | SS18-SSX2 | THSD7B-DARS |
| IGH | SS18 | TMEM123 |
| BCL6 | SSX1 | MMP7 |
| IGH-BCL6 | SS18-SSX1 | TMEM123-MMP7 |
| LCP1 | SS18L1 | TMPRSS2 |
| BCL6 | SSX1 | ERG |

| | | |
|---|---|---|
| LCP1-BCL6 | SS18L1-SSX1 | TMPRSS2-ERG |
| CREBBP | SSX1 | TOX3 |
| KAT6A | SYT4 | CNTN5 |
| CREBBP-KAT6A | SSX1-SYT4 | TOX3-CNTN5 |
| KMT2A | TGFBR3 | UBR2 |
| ARHGAP26 | MGEA5 | XPO5 |
| KMT2A-ARHGAP26 | TGFBR3-MGEA5 | UBR2-XPO5 |
| FOXO3 | TRIO | UVRAG |
| KMT2A | TERT | INTS4 |
| FOXO3-KMT2A | TRIO-TERT | UVRAG-INTS4 |
| KMT2A | WDR70 | WNT11 |
| DCPS | RCOR1 | TSPAN8 |
| KMT2A-DCPS | WDR70-RCOR1 | WNT11-TSPAN8 |
| KMT2A | YWHAE | XRCC5 |
| EP300 | NUTM2B | ACADL |
| KMT2A-EP300 | YWHAE-NUTM2B | XRCC5-ACADL |
| IGH | YWHAE | ZCCHC7 |
| CEBPE | NUTM2A-AS1 | PRSS3 |
| IGH-CEBPE | YWHAE-NUTM2A-AS1 | ZCCHC7-PRSS3 |
| HSP90AA1 | YWHAE | ZFP91 |
| BCL6 | NUTM2A | NOX4 |
| HSP90AA1-BCL6 | YWHAE-NUTM2A | ZFP91-NOX4 |

**Table S2: LK, LY, ME, GL- Network categorization, scalefree_hier_rand (TRAINING)**

| LK, LY, ME, GL | | | |
|---|---|---|---|
| | Scale-Free | Hierarchical | Random |
| BCR | √ | | |
| ABL1 | √ | | |
| BCR-ABL1 | √ | | |
| RUNX1 | √ | | |
| RUNX1T1 | √ | | |
| RUNX1-RUNX1T1 | √ | | |
| KMT2A | √ | | |
| MLLT10 | | √ | |
| KMT2A-MLLT10 | √ | | |
| IGH | | | √ |
| BCL2 | √ | | |
| IGH-BCL2 | √ | | |
| KMT2A | √ | | |
| AFF1 | | √ | |
| KMT2A-AFF1 | √ | | |
| PICALM | √ | | |

| | | | |
|---|---|---|---|
| MLLT10 | | √ | |
| PICALM-MLLT10 | √ | | |
| PML | √ | | |
| RARA | √ | | |
| PML-RARA | √ | | |
| KMT2A | √ | | |
| MLLT3 | √ | | |
| KMT2A-MLLT3 | √ | | |
| KMT2A | √ | | |
| AFDN | √ | | |
| KMT2A-AFDN | √ | | |
| CBFB | √ | | |
| MYH11 | √ | | |
| CBFB-MYH11 | | √ | |
| IGH | | | √ |
| MYC | √ | | |
| IGH-MYC | √ | | |
| NUP98 | √ | | |
| DDX10 | √ | | |
| NUP98-DDX10 | √ | | |
| PCM1 | √ | | |
| JAK2 | √ | | |
| PCM1-JAK2 | √ | | |
| KMT2A | √ | | |
| SEPT9 | √ | | |
| KMT2A-SEPT9 | √ | | |
| FUS | √ | | |
| ERG | √ | | |
| FUS-ERG | √ | | |
| NUP98 | √ | | |
| HOXA9 | | √ | |
| NUP98-HOXA9 | | √ | |
| ETV6 | | √ | |
| ABL1 | √ | | |
| ETV6-ABL1 | √ | | |
| SET | | | |
| NUP214 | | | |
| SET-NUP214 | | | |
| MNX1 | | | |
| ETV6 | | √ | |
| MNX1-ETV6 | | | |
| KMT2A | √ | | |
| MLLT1 | | | |
| KMT2A-MLLT1 | | | |

| | | | |
|---|---|---|---|
| KMT2A | √ | | |
| MLLT6 | | | |
| KMT2A-MLLT6 | | | |
| ETV6 | | √ | |
| ACSL6 | | | |
| ETV6-ACSL6 | | | |
| ETV6 | | √ | |
| MECOM | | | |
| ETV6-MECOM | | | |
| KMT2A | √ | | |
| EPS15 | | | |
| KMT2A-EPS15 | | | |
| KMT2A | √ | | |
| GAS7 | | | |
| KMT2A-GAS7 | | | |
| KMT2A | √ | | |
| ABI1 | √ | | |
| KMT2A-ABI1 | √ | | |
| KMT2A | √ | | |
| MLLT11 | √ | | |
| KMT2A-MLLT11 | √ | | |
| MN1 | | | √ |
| ETV6 | | √ | |
| MN1-ETV6 | √ | | |
| KMT2A | √ | | |
| MAML2 | | √ | |
| KMT2A-MAML2 | √ | | |
| KMT2A | √ | | |
| FOXO4 | | √ | |
| KMT2A-FOXO4 | √ | | |
| DEK | √ | | |
| NUP214 | √ | | |
| DEK-NUP214 | √ | | |
| RUNX1 | √ | | |
| CBFA2T3 | | √ | |
| RUNX1-CBFA2T3 | √ | | |
| NUP98 | √ | | |
| PSIP1 | √ | | |
| NUP98-PSIP1 | √ | | |
| NUP98 | √ | | |
| HOXC13 | | √ | |
| NUP98-HOXC13 | | √ | |
| NUP98 | √ | | |
| HOXC11 | | √ | |

| | | | |
|---|---|---|---|
| NUP98-HOXC11 | √ | | |
| PAX5 | | √ | |
| ETV6 | | √ | |
| PAX5-ETV6 | √ | | |
| NUP98 | √ | | |
| HOXA11 | | | √ |
| NUP98-HOXA11 | √ | | |
| BCR | √ | | |
| PDGFRA | | | |
| BCR-PDGFRA | | | |
| BCR | √ | | |
| FGFR1 | | √ | |
| BCR-FGFR1 | √ | | |
| NPM1 | √ | | |
| RARA | √ | | |
| NPM1-RARA | √ | | |
| KMT2A | √ | | |
| CBL | √ | | |
| KMT2A-CBL | √ | | |
| IGH | | | √ |
| BCL6 | √ | | |
| IGH-BCL6 | √ | | |
| LCP1 | | √ | |
| BCL6 | √ | | |
| LCP1-BCL6 | √ | | |
| CREBBP | √ | | |
| KAT6A | √ | | |
| CREBBP-KAT6A | √ | | |
| KMT2A | √ | | |
| ARHGAP26 | | | √ |
| KMT2A-ARHGAP26 | √ | | |
| FOXO3 | √ | | |
| KMT2A | √ | | |
| FOXO3-KMT2A | √ | | |
| KMT2A | √ | | |
| DCPS | √ | | |
| KMT2A-DCPS | √ | | |
| KMT2A | √ | | |
| EP300 | | | √ |
| KMT2A-EP300 | √ | | |
| IGH | | | √ |
| CEBPE | | √ | |
| IGH-CEBPE | | √ | |
| HSP90AA1 | | √ | |

| | |
|---|---|
| BCL6 | √ |
| HSP90AA1-BCL6 | √ |

**Table S3: SC-Network categorization, scalefree_hier_rand (TRAINING)**

| SC | | | |
|---|---|---|---|
| | Scale-Free | Hierarchical | Random |
| ASPSCR1 | √ | | |
| TFE3 | | √ | |
| ASPSCR1-TFE3 | √ | | |
| ASTN2 | | | √ |
| CNOT2 | √ | | |
| ASTN2-CNOT2 | | √ | |
| BCOR | √ | | |
| ZC3H7B | | √ | |
| BCOR-ZC3H7B | | √ | |
| BCOR | √ | | |
| CCNB3 | | | √ |
| BCOR-CCNB3 | | √ | |
| CDX1 | | | √ |
| IRF2BP2 | | √ | |
| CDX1-IRF2BP2 | | | √ |
| CIC | √ | | |
| DUX4 | | | √ |
| CIC-DUX4 | | √ | |
| CREB1 | √ | | |
| EWSR1 | √ | | |
| CREB1-EWSR1 | √ | | |
| CTDSP2 | | √ | |
| FAM19A2 | | | √ |
| CTDSP2-FAM19A2 | | √ | |
| CXorf67 | | | √ |
| MBTD1 | √ | | |
| CXorf67-MBTD1 | | √ | |
| EPC1 | √ | | |
| PHF1 | √ | | |
| EPC1-PHF1 | √ | | |
| ERG | √ | | |
| EWSR1 | √ | | |
| ERG-EWSR1 | √ | | |
| ETV6 | | √ | |
| NTRK3 | | √ | |
| ETV6-NTRK3 | √ | | |
| EWSR1 | √ | | |

| | | | |
|---|---|---|---|
| ATF1 | √ | | |
| EWSR1-ATF1 | √ | | |
| EWSR1 | √ | | |
| FLI1 | | √ | |
| EWSR1-FLI1 | √ | | |
| EWSR1 | √ | | |
| NR4A3 | | | √ |
| EWSR1-NR4A3 | √ | | |
| EWSR1 | √ | | |
| ETV4 | | √ | |
| EWSR1-ETV4 | √ | | |
| EWSR1 | √ | | |
| PATZ1 | | √ | |
| EWSR1-PATZ1 | √ | | |
| EWSR1 | √ | | |
| DDIT3 | √ | | |
| EWSR1-DDIT3 | √ | | |
| EWSR1 | √ | | |
| POU5F1 | √ | | |
| EWSR1-POU5F1 | √ | | |
| EWSR1 | √ | | |
| SP3 | √ | | |
| EWSR1-SP3 | √ | | |
| FOXO4 | | √ | |
| CIC | √ | | |
| FOXO4-CIC | √ | | |
| FUS | √ | | |
| ERG | √ | | |
| FUS-ERG | √ | | |
| FUS | √ | | |
| CREB3L1 | | √ | |
| FUS-CREB3L1 | √ | | |
| FUS | √ | | |
| DDIT3 | √ | | |
| FUS-DDIT3 | √ | | |
| FUS | √ | | |
| ATF1 | √ | | |
| FUS-ATF1 | √ | | |
| FUS | √ | | |
| CREB3L2 | | | √ |
| FUS-CREB3L2 | √ | | |
| HEY1 | | √ | |
| NCOA2 | √ | | |
| HEY1-NCOA2 | | √ | |

| | | | |
|---|---|---|---|
| IRX2 | | | √ |
| TERT | √ | | |
| IRX2-TERT | √ | | |
| JAZF1 | | | √ |
| SUZ12 | √ | | |
| JAZF1-SUZ12 | √ | | |
| JAZF1 | | | √ |
| PHF1 | √ | | |
| JAZF1-PHF1 | √ | | |
| LMNA | √ | | |
| NTRK1 | √ | | |
| LMNA-NTRK1 | √ | | |
| MEAF6 | | √ | |
| TRERF1 | | √ | |
| MEAF6-TRERF1 | | √ | |
| MEAF6 | | √ | |
| PHF1 | √ | | |
| MEAF6-PHF1 | √ | | |
| NR4A3 | | | √ |
| TAF15 | | √ | |
| NR4A3-TAF15 | | √ | |
| NR4A3 | | | √ |
| TFG | √ | | |
| NR4A3-TFG | √ | | |
| NR6A1 | | | √ |
| TRHDE | | | √ |
| NR6A1-TRHDE | | | √ |
| NUP107 | √ | | |
| LGR5 | | | √ |
| NUP107-LGR5 | √ | | |
| PAPPA | | √ | |
| NUP107 | √ | | |
| PAPPA-NUP107 | √ | | |
| PAX3 | | √ | |
| FOXO1 | √ | | |
| PAX3-FOXO1 | √ | | |
| PAX7 | | | √ |
| FOXO1 | √ | | |
| PAX7-FOXO1 | √ | | |
| SS18 | √ | | |
| SSX2 | | √ | |
| SS18-SSX2 | √ | | |
| SS18 | √ | | |
| SSX1 | | | √ |

| | | | |
|---|---|---|---|
| SS18-SSX1 | √ | | |
| SS18L1 | √ | | |
| SSX1 | | | √ |
| SS18L1-SSX1 | √ | | |
| SSX1 | | | √ |
| SYT4 | | | √ |
| SSX1-SYT4 | | | √ |
| TGFBR3 | | √ | |
| MGEA5 | √ | | |
| TGFBR3-MGEA5 | √ | | |
| TRIO | | √ | |
| TERT | √ | | |
| TRIO-TERT | √ | | |
| WDR70 | | √ | |
| RCOR1 | √ | | |
| WDR70-RCOR1 | √ | | |
| YWHAE | | | √ |
| NUTM2B | | | √ |
| YWHAE-NUTM2B | √ | | |
| YWHAE | | | √ |
| NUTM2A-AS1 | | | √ |
| YWHAE-NUTM2A-AS1 | √ | | |
| YWHAE | | | √ |
| NUTM2A | | | √ |
| YWHAE-NUTM2A | √ | | |

**Table S4: CA- Network categorization, scalefree_hier_rand (TRAINING)**

| CA | | | |
|---|---|---|---|
| | Scale-Free | Hierarchical | Random |
| ANK3 | | √ | |
| USP9Y | | | √ |
| ANK3-USP9Y | | √ | |
| ARGLU1 | √ | | |
| CXCR4 | √ | | |
| ARGLU1-CXCR4 | √ | | |
| ATXN10 | √ | | |
| FBLN1 | √ | | |
| ATXN10-FBLN1 | √ | | |
| BCAS3 | √ | | |
| NFS1 | | √ | |
| BCAS3-NFS1 | | √ | |
| BCAS4 | | √ | |
| BCAS3 | √ | | |
| BCAS4-BCAS3 | | √ | |

| | | | |
|---|---|---|---|
| BCL2L12 | | | √ |
| PRMT1 | √ | | |
| BCL2L12-PRMT1 | √ | | |
| CAPNS1 | √ | | |
| WDR62 | √ | | |
| CAPNS1-WDR62 | √ | | |
| CCDC6 | | √ | |
| ANK3 | √ | | |
| CCDC6-ANK3 | √ | | |
| CCDC9 | | √ | |
| DHX34 | | | √ |
| CCDC9-DHX34 | | √ | |
| CDC27 | √ | | |
| ST7L | | | √ |
| CDC27-ST7L | √ | | |
| CDK7 | √ | | |
| RIN3 | √ | | |
| CDK7-RIN3 | √ | | |
| CHERP | √ | | |
| CPAMD8 | | | √ |
| CHERP-CPAMD8 | √ | | |
| CPD | | √ | |
| PIGW | | | √ |
| CPD-PIGW | | | √ |
| CYTH1 | | √ | |
| PRPSAP1 | √ | | |
| CYTH1-PRPSAP1 | √ | | |
| DLG1 | √ | | |
| CRYBG3 | | | √ |
| DLG1-CRYBG3 | √ | | |
| DTX4 | | | √ |
| CCDC102B | √ | | |
| DTX4-CCDC102B | √ | | |
| EHD4 | √ | | |
| FSIP1 | | | √ |
| EHD4-FSIP1 | √ | | |
| ELK4 | | √ | |
| SLC26A9 | | | √ |
| ELK4-SLC26A9 | | √ | |
| EMID1 | | | √ |
| CBY1 | | √ | |
| EMID1-CBY1 | | | √ |
| EPHA6 | | | √ |
| CNTN6 | | | √ |
| EPHA6-CNTN6 | | | √ |
| ERAL1 | √ | | |
| DIDO1 | √ | | |
| ERAL1-DIDO1 | √ | | |

| | | | |
|---|---|---|---|
| GMDS | | √ | |
| CCND3 | √ | | |
| GMDS-CCND3 | √ | | |
| HJURP | √ | | |
| EIF4E2 | √ | | |
| HJURP-EIF4E2 | √ | | |
| INTS4 | √ | | |
| GAB2 | √ | | |
| INTS4-GAB2 | √ | | |
| KCNQ5 | | | √ |
| RIMS1 | | √ | |
| KCNQ5-RIMS1 | | √ | |
| KDM5A | √ | | |
| ANO2 | | | √ |
| KDM5A-ANO2 | √ | | |
| LAMA5 | | | √ |
| C12orf28 | | | √ |
| LAMA5-C12orf28 | | | √ |
| MAPK10 | √ | | |
| FAM13A | | √ | |
| MAPK10-FAM13A | √ | | |
| MAPRE1 | √ | | |
| TM9SF4 | √ | | |
| MAPRE1-TM9SF4 | √ | | |
| NCKAP5 | | | √ |
| MZT2A | | √ | |
| NCKAP5-MZT2A | | √ | |
| NUMB | √ | | |
| ALDH6A1 | | √ | |
| NUMB-ALDH6A1 | √ | | |
| PARD6B | √ | | |
| CD48 | √ | | |
| PARD6B-CD48 | | √ | |
| PPP1R12A | √ | | |
| MGAT4C | | | √ |
| PPP1R12A-MGAT4C | √ | | |
| RNF11 | √ | | |
| C8A | | | √ |
| RNF11-C8A | √ | | |
| SIPA1L3 | √ | | |
| WDR62 | | √ | |
| SIPA1L3-WDR62 | | √ | |
| SLC26A6 | | | √ |
| PRKAR2A | √ | | |
| SLC26A6-PRKAR2A | √ | | |
| ST14 | | √ | |
| APLP2 | | √ | |

| | | | |
|---|---|---|---|
| ST14-APLP2 | | √ | |
| STRADB | √ | | |
| NOP58 | √ | | |
| STRADB-NOP58 | √ | | |
| STX16 | | √ | |
| RAE1 | √ | | |
| STX16-RAE1 | √ | | |
| TANC2 | | √ | |
| CHD6 | | √ | |
| TANC2-CHD6 | | √ | |
| THSD7B | | | √ |
| DARS | √ | | |
| THSD7B-DARS | √ | | |
| TMEM123 | | √ | |
| MMP7 | | √ | |
| TMEM123-MMP7 | | √ | |
| TMPRSS2 | | | √ |
| ERG | √ | | |
| TMPRSS2-ERG | √ | | |
| TOX3 | | | √ |
| CNTN5 | | | √ |
| TOX3-CNTN5 | | | √ |
| UBR2 | | | √ |
| XPO5 | | | √ |
| UBR2-XPO5 | | | √ |
| UVRAG | | | √ |
| INTS4 | | | √ |
| UVRAG-INTS4 | | | √ |
| WNT11 | | | √ |
| TSPAN8 | | | √ |
| WNT11-TSPAN8 | | | √ |
| XRCC5 | | | √ |
| ACADL | | | √ |
| XRCC5-ACADL | | | √ |
| ZCCHC7 | | | √ |
| PRSS3 | | | √ |
| ZCCHC7-PRSS3 | | | √ |
| ZFP91 | | | √ |
| NOX4 | | | √ |
| ZFP91-NOX4 | | | √ |

**Table S5: LK, LY, ME, GL-Hubs (TRAINING)**

| LK, LY, ME, GL | Higher-degree Hubs | | Lower-degree Hubs | |
|---|---|---|---|---|
| | Hub | Degree | Hub | Degree |
| BCR-ABL1 | abl1 | 172 | bcr | 53 |
| | ntrk1 | 86 | mdm2 | 53 |
| | egfr | 71 | tp53 | 53 |

| | | | | |
|---|---|---|---|---|
| | grb2 | 67 | cbl | 50 |
| RUNX1-RUNX1T1 | runx1 | 68 | hdac1 | 40 |
| | runx1t1 | 58 | ep300 | 39 |
| | | | crebbp | 35 |
| | | | hdac2 | 33 |
| | | | smad3 | 33 |
| KMT2A-MLLT10 | kmt2a | 90 | tp53 | 37 |
| | | | elavl1 | 33 |
| | | | hdac1 | 30 |
| | | | hdac2 | 28 |
| | | | rnf2 | 28 |
| | | | crebbp | 27 |
| IGH-BCL2 | bcl2 | 92 | bcl2l1 | 41 |
| | | | tp53 | 29 |
| | | | hsp90aa1 | 23 |
| | | | myc | 21 |
| KMT2A-AFF1 | kmt2a | 90 | tp53 | 41 |
| | | | hdac1 | 33 |
| | | | rnf2 | 30 |
| | | | aff1 | 29 |
| | | | hist3h3 | 29 |
| | | | crebbp | 28 |
| | | | hdac2 | 28 |
| | | | hist1h3a | 27 |
| | | | sin3a | 27 |
| PICALM-MLLT10 | picalm | 56 | ntrk1 | 31 |
| | | | fn1 | 24 |
| | | | eef1a1 | 20 |
| | | | itga4 | 20 |
| | | | hnrnpa1 | 19 |
| PML-RARA | pml | 182 | ep300 | 77 |
| | rara | 113 | hdac1 | 75 |
| | tp53 | 106 | ube2i | 75 |
| | | | crebbp | 66 |
| KMT2A-MLLT3 | kmt2a | 90 | tp53 | 40 |
| | | | mllt3 | 39 |
| | | | rnf2 | 33 |
| | | | hdac1 | 31 |
| | | | hist3h3 | 29 |
| | | | hdac2 | 28 |
| | | | sin3a | 28 |
| | | | crebbp | 26 |
| KMT2A-AFDN | kmt2a | 90 | mllt4 | 55 |
| | | | tp53 | 44 |
| | | | ntrk1 | 42 |
| | | | hdac1 | 33 |
| | | | hdac2 | 32 |
| | | | hist1h3a | 29 |

| | | | | |
|---|---|---|---|---|
| | | | hist3h3 | 29 |
| CBFB-MYH11 | cbfb | 23 | actb | 16 |
| | myh11 | 23 | elavl1 | 13 |
| | | | rpa1 | 10 |
| | | | rpa2 | 10 |
| | | | rpa3 | 10 |
| IGH-MYC | igh | 50 | tp53 | 29 |
| NUP98-DDX10 | nup98 | 51 | ntrk1 | 28 |
| | | | sirt7 | 25 |
| | | | nxf1 | 24 |
| | | | obsl1 | 24 |
| | | | npm1 | 22 |
| PCM1-JAK2 | jak2 | 93 | ntrk1 | 46 |
| | | | egfr | 45 |
| | | | grb2 | 44 |
| | | | pcm1 | 40 |
| | | | hsp90aa1 | 35 |
| | | | src | 32 |
| | | | shc1 | 30 |
| KMT2A-SEPT9 | kmt2a | 90 | sept9 | 46 |
| | | | tp53 | 43 |
| | | | ntrk1 | 42 |
| | | | hdac1 | 34 |
| | | | rnf2 | 33 |
| | | | hdac2 | 31 |
| | | | sin3a | 31 |
| | | | crebbp | 29 |
| FUS-ERG | fus | 315 | cul7 | 169 |
| | ntrk1 | 227 | fn1 | 154 |
| | cul3 | 202 | cops5 | 151 |
| | | | obsl1 | 145 |
| | | | cdk2 | 138 |
| | | | cand1 | 135 |
| | | | mdm2 | 133 |
| | | | cul1 | 130 |
| | | | itga4 | 127 |
| | | | hnrnpa1 | 124 |
| | | | vcam1 | 120 |
| | | | esr1 | 107 |
| | | | cul2 | 105 |
| NUP98-HOXA9 | nup98 | 51 | nxf1 | 22 |
| | | | tp53 | 21 |
| | | | obsl1 | 20 |
| | | | sirt7 | 19 |
| ETV6-ABL1 | abl1 | 172 | ntrk1 | 86 |
| | | | egfr | 72 |
| | | | grb2 | 63 |
| | | | tp53 | 56 |

| | | | | |
|---|---|---|---|---|
| | | | mdm2 | 53 |
| SET-NUP214 | set | 97 | nup214 | 54 |
| | | | ntrk1 | 45 |
| | | | cul3 | 39 |
| | | | tp53 | 38 |
| | | | cdk2 | 34 |
| | | | xrcc6 | 34 |
| | | | elavl1 | 33 |
| | | | nxf1 | 33 |
| | | | app | 30 |
| MNX1-ETV6 | etv6 | 32 | elavl1 | 9 |
| | | | grb2 | 8 |
| | | | hdac3 | 8 |
| | | | ube2i | 8 |
| KMT2A-MLLT1 | kmt2a | 90 | tp53 | 39 |
| | | | hdac1 | 32 |
| | | | rnf2 | 30 |
| | | | hdac2 | 29 |
| | | | hist1h3a | 28 |
| | | | hist3h3 | 28 |
| | | | mllt1 | 28 |
| | | | crebbp | 27 |
| | | | sin3a | 27 |
| KMT2A-MLLT6 | kmt2a | 90 | tp53 | 38 |
| | | | elavl1 | 37 |
| | | | crebbp | 29 |
| | | | hdac1 | 29 |
| | | | rnf2 | 29 |
| | | | hdac2 | 28 |
| | | | hist3h3 | 28 |
| | | | wdr5 | 28 |
| | | | sin3a | 27 |
| ETV6-ACSL6 | etv6 | 32 | elavl1 | 8 |
| | | | grb2 | 8 |
| | | | hdac3 | 8 |
| | | | ube2i | 8 |
| ETV6-MECOM | etv6 | 32 | elavl1 | 16 |
| | hdac1 | 20 | ube2i | 16 |
| | mecom | 20 | smad3 | 15 |
| | | | crebbp | 14 |
| | | | hdac4 | 14 |
| KMT2A-EPS15 | eps15 | 112 | kmt2a | 90 |
| | | | ntrk1 | 62 |
| | | | tp53 | 59 |
| | | | grb2 | 57 |
| KMT2A-GAS7 | kmt2a | 90 | tp53 | 39 |
| | | | hdac1 | 31 |
| | | | rnf2 | 30 |

| | | | | |
|---|---|---|---|---|
| | | | hist3h3 | 28 |
| | | | sin3a | 28 |
| | | | crebbp | 27 |
| | | | hdac2 | 27 |
| KMT2A-ABL1 | abl1 | 172 | egfr | 71 |
| | ntrk1 | 105 | mdm2 | 67 |
| | kmt2a | 90 | grb2 | 63 |
| | tp53 | 80 | brca1 | 51 |
| | | | hsp90aa1 | 50 |
| KMT2A-MLLT11 | kmt2a | 90 | tp53 | 37 |
| | | | hdac1 | 29 |
| | | | hist3h3 | 27 |
| | | | rnf2 | 27 |
| | | | crebbp | 26 |
| | | | hdac2 | 26 |
| | | | sin3a | 26 |
| | | | wdr5 | 26 |
| MN1-ETV6 | etv6 | 32 | ep300 | 12 |
| | | | elavl1 | 9 |
| | | | grb2 | 9 |
| | | | hdac3 | 9 |
| | | | ube2i | 9 |
| KMT2A-MAML2 | kmt2a | 90 | tp53 | 37 |
| | | | hdac1 | 31 |
| | | | ep300 | 30 |
| | | | crebbp | 29 |
| KMT2A-FOXO4 | kmt2a | 90 | tp53 | 45 |
| | | | esr1 | 37 |
| | | | crebbp | 35 |
| | | | hdac1 | 35 |
| | | | sin3a | 31 |
| DEK-NUP214 | nup214 | 54 | ntrk1 | 40 |
| | dek | 46 | nxf1 | 35 |
| | | | cul3 | 33 |
| | | | cops5 | 32 |
| | | | sirt7 | 32 |
| RUNX1-CBFA2T3 | runx1 | 68 | ep300 | 36 |
| | | | hdac1 | 35 |
| | | | crebbp | 31 |
| | | | hdac2 | 27 |
| | | | smad3 | 27 |
| NUP98-PSIP1 | nup98 | 51 | psip1 | 39 |
| | | | obsl1 | 33 |
| | | | nxf1 | 32 |
| | | | cul7 | 28 |
| NUP98-HOXC13 | nup98 | 51 | nxf1 | 24 |
| | | | obsl1 | 20 |
| | | | cul7 | 17 |

| | | | | |
|---|---|---|---|---|
| | | | sirt7 | 17 |
| NUP98-HOXC11 | nup98 | 51 | nxf1 | 24 |
| | | | obsl1 | 20 |
| | | | kpnb1 | 18 |
| | | | sirt7 | 18 |
| | | | cul7 | 17 |
| | | | crebbp | 16 |
| | | | elavl1 | 16 |
| PAX5-ETV6 | etv6 | 32 | ep300 | 16 |
| | | | ube2i | 13 |
| | | | hdac3 | 12 |
| NUP98-HOXA11 | nup98 | 51 | nxf1 | 23 |
| | | | obsl1 | 20 |
| BCR-PDGFRA | bcr | 53 | egfr | 35 |
| | | | grb2 | 33 |
| | | | ntrk1 | 31 |
| | | | abl1 | 29 |
| BCR-FGFR1 | bcr | 53 | grb2 | 37 |
| | fgfr1 | 44 | abl1 | 33 |
| | | | ntrk1 | 32 |
| | | | cbl | 31 |
| | | | hsp90aa1 | 31 |
| NPM1-RARA | rara | 113 | hdac1 | 75 |
| KMT2A-CBL | kmt2a | 90 | tp53 | 37 |
| IGH-BCL6 | bcl6 | 111 | hdac1 | 31 |
| | | | tp53 | 28 |
| | | | crebbp | 23 |
| | | | hdac2 | 22 |
| LCP1-BCL6 | bcl6 | 111 | hdac1 | 32 |
| | | | tp53 | 29 |
| | | | crebbp | 23 |
| | | | hdac2 | 23 |
| CREBBP-KAT6A | kmt2a | 90 | tp53 | 41 |
| | | | hdac1 | 33 |
| | | | rnf2 | 30 |
| | | | aff1 | 29 |
| | | | hist3h3 | 29 |
| | | | crebbp | 28 |
| | | | hdac2 | 28 |
| KMT2A-ARHGAP26 | kmt2a | 90 | tp53 | 40 |
| | | | hdac1 | 29 |
| | | | hist3h3 | 27 |
| | | | rnf2 | 27 |
| | | | crebbp | 26 |
| | | | hdac2 | 26 |
| | | | sin3a | 26 |
| FOXO3-KMT2A | kmt2a | 90 | tp53 | 56 |
| | | | elavl1 | 45 |

| | | | | |
|---|---|---|---|---|
| | | | ep300 | 42 |
| | | | foxo3 | 42 |
| | | | crebbp | 41 |
| KMT2A-DCPS | kmt2a | 90 | ntrk1 | 49 |
| | | | tp53 | 39 |
| | | | dcps | 37 |
| KMT2A-EP300 | kmt2a | 90 | tp53 | 56 |
| IGH-CEBPE | cebpe | 45 | jun | 20 |
| | | | atf4 | 15 |
| | | | cebpa | 15 |
| HSP90AA1-BCL6 | bcl6 | 111 | hsp90aa1 | 52 |
| | | 86.0152 | | 37.3307 |

**Table S6: SC-Hubs (TRAINING)**

| SC | Higher-degree Hubs | | Lower-degree Hubs | |
|---|---|---|---|---|
| | **Hub** | **Degree** | **Hub** | **Degree** |
| ASPSCR1-TFE3 | aspscr1 | 31 | ubc | 16 |
| | ntrk1 | 24 | vcp | 15 |
| | tfe3 | 21 | ago2 | 10 |
| ASTN2-CNOT2 | cnot2 | 23 | cnot6 | 10 |
| | cnot1 | 13 | cnot6l | 10 |
| | cnot3 | 12 | | |
| | cnot7 | 12 | | |
| | cnot8 | 11 | | |
| BCOR-ZC3H7B | bcor | 34 | rnf2 | 18 |
| | | | pcgf1 | 14 |
| | | | ring1 | 13 |
| BCOR-CCNB3 | bcor | 34 | rnf2 | 17 |
| | | | pcgf1 | 13 |
| | | | ring1 | 13 |
| | | | hdac3 | 12 |
| | | | usp7 | 12 |
| | | | eed | 11 |
| | | | hdac1 | 11 |
| | | | hdac4 | 10 |
| | | | rybp | 10 |
| | | | sp1 | 10 |
| CDX1-IRF2BP2 | irf2bp2 | 16 | elavl1 | 9 |
| | | | ntrk1 | 8 |
| CIC-DUX4 | cic | 14 | ntrk1 | 6 |
| | | | atxn1 | 4 |
| | | | csnk2b | 4 |
| | | | setd2 | 4 |
| CREB1-EWSR1 | ewsr1 | 234 | creb1 | 96 |
| | | | ntrk1 | 76 |
| | | | elavl1 | 75 |
| | | | cul3 | 72 |

| | | | | |
|---|---|---|---|---|
| | | | tp53 | 69 |
| CTDSP2-FAM19A2 | ctdsp2 | 22 | polr2a | 6 |
| CXorf67-MBTD1 | kat5 | 11 | mrgbp | 9 |
| | mbtd1 | 11 | yeats4 | 9 |
| | morf4l1 | 11 | | |
| EPC1-PHF1 | phf1 | 39 | kat5 | 23 |
| | | | epc1 | 21 |
| | | | morf4l1 | 21 |
| | | | eed | 20 |
| | | | hdac1 | 19 |
| ERG-EWSR1 | ewsr1 | 234 | hnrnpa1 | 87 |
| | ntrk1 | 123 | cops5 | 83 |
| | cul3 | 112 | cand1 | 80 |
| | | | cul7 | 79 |
| | | | erg | 77 |
| | | | elavl1 | 76 |
| | | | obsl1 | 75 |
| | | | tp53 | 74 |
| | | | cul1 | 71 |
| | | | rpa1 | 71 |
| | | | fn1 | 70 |
| ETV6-NTRK3 | etv6 | 32 | ntrk1 | 20 |
| | | | grb2 | 15 |
| | | | hsp90aa1 | 14 |
| EWSR1-ATF1 | ewsr1 | 234 | ntrk1 | 69 |
| | | | cul3 | 63 |
| | | | elavl1 | 59 |
| | | | hnrnpa1 | 52 |
| | | | fus | 48 |
| | | | tp53 | 46 |
| EWSR1-FLI1 | ewsr1 | 234 | ntrk1 | 67 |
| | | | cul3 | 62 |
| | | | elavl1 | 59 |
| | | | hnrnpa1 | 53 |
| | | | tp53 | 50 |
| | | | fus | 48 |
| EWSR1-NR4A3 | ewsr1 | 234 | ntrk1 | 68 |
| | | | cul3 | 63 |
| | | | elavl1 | 58 |
| EWSR1-ETV4 | ewsr1 | 234 | ntrk1 | 69 |
| | | | cul3 | 64 |
| | | | elavl1 | 59 |
| EWSR1-PATZ1 | ewsr1 | 234 | ntrk1 | 67 |
| | | | cul3 | 64 |
| | | | elavl1 | 59 |
| EWSR1-DDIT3 | ewsr1 | 234 | ntrk1 | 76 |
| | | | cul3 | 72 |
| | | | ddit3 | 70 |

| | | | | |
|---|---|---|---|---|
| EWSR1-POU5F1 | ewsr1 | 234 | cul3 | 87 |
| | | | ntrk1 | 86 |
| | | | elavl1 | 68 |
| | | | hnrnpa1 | 65 |
| | | | cops5 | 62 |
| | | | tp53 | 62 |
| | | | cand1 | 61 |
| | | | pou5f1 | 61 |
| EWSR1-SP3 | ewsr1 | 234 | ntrk1 | 71 |
| | | | elavl1 | 64 |
| | | | cul3 | 62 |
| | | | tp53 | 53 |
| | | | hnrnpa1 | 52 |
| | | | fus | 50 |
| FOXO4-CIC | foxo4 | 17 | ctnnb1 | 12 |
| | cic | 14 | akt1 | 11 |
| | crebbp | 13 | esr1 | 11 |
| | mdm2 | 13 | ntrk1 | 11 |
| | smad3 | 13 | | |
| FUS-ERG | fus | 315 | cul7 | 169 |
| | ntrk1 | 227 | fn1 | 154 |
| | cul3 | 202 | cops5 | 151 |
| | | | obsl1 | 145 |
| | | | cdk2 | 138 |
| | | | cand1 | 135 |
| | | | mdm2 | 133 |
| | | | cul1 | 130 |
| | | | itga4 | 127 |
| | | | hnrnpa1 | 124 |
| | | | vcam1 | 120 |
| | | | esr1 | 107 |
| | | | cul2 | 105 |
| FUS-CREB3L1 | fus | 315 | cul7 | 130 |
| | ntrk1 | 181 | fn1 | 127 |
| | cul3 | 166 | cops5 | 119 |
| | | | cdk2 | 114 |
| | | | mdm2 | 109 |
| | | | cul1 | 108 |
| | | | obsl1 | 106 |
| | | | cand1 | 104 |
| | | | itga4 | 102 |
| FUS-DDIT3 | fus | 315 | cul7 | 134 |
| | ntrk1 | 189 | fn1 | 132 |
| | cul3 | 173 | cops5 | 126 |
| | | | cdk2 | 121 |
| | | | mdm2 | 116 |
| | | | cul1 | 113 |
| | | | obsl1 | 111 |

| | | | | |
|---|---|---|---|---|
| | | | cand1 | 106 |
| | | | itga4 | 103 |
| | | | hnrnpa1 | 100 |
| FUS-ATF1 | fus | 315 | cul7 | 131 |
| | ntrk1 | 183 | fn1 | 129 |
| | cul3 | 165 | cops5 | 118 |
| | | | cdk2 | 116 |
| | | | mdm2 | 109 |
| | | | obsl1 | 108 |
| | | | cul1 | 107 |
| | | | cand1 | 104 |
| | | | itga4 | 103 |
| FUS-CREB3L2 | fus | 315 | cul7 | 130 |
| | ntrk1 | 182 | fn1 | 129 |
| | cul3 | 165 | cops5 | 119 |
| | | | cdk2 | 114 |
| | | | cul1 | 108 |
| | | | mdm2 | 108 |
| | | | obsl1 | 107 |
| | | | cand1 | 105 |
| | | | itga4 | 103 |
| HEY1-NCOA2 | ncoa2 | 60 | ep300 | 39 |
| | | | ncoa1 | 39 |
| | | | ncoa3 | 36 |
| | | | crebbp | 30 |
| | | | tp53 | 30 |
| IRX2-TERT | tert | 64 | mdm2 | 28 |
| | | | tp53 | 26 |
| | | | hsp90aa1 | 16 |
| | | | stub1 | 16 |
| JAZF1-SUZ12 | SUZ12 | 331 | EED | 140 |
| | RNF2 | 202 | EZH2 | 131 |
| | | | ELAVL1 | 76 |
| | | | NXF1 | 71 |
| JAZF1-PHF1 | phf1 | 39 | eed | 18 |
| | | | ezh2 | 15 |
| | | | rbbp4 | 15 |
| | | | hdac1 | 14 |
| | | | suz12 | 14 |
| LMNA-NTRK1 | lmna | 222 | ntrk1 | 99 |
| | | | obsl1 | 79 |
| | | | cul7 | 73 |
| | | | tp53 | 54 |
| MEAF6-TRERF1 | meaf6 | 25 | kat5 | 16 |
| | | | morf4l1 | 16 |
| MEAF6-PHF1 | phf1 | 39 | meaf6 | 25 |
| | | | morf4l1 | 20 |
| | | | kat5 | 19 |

| | | | | |
|---|---|---|---|---|
| NR4A3-TAF15 | taf15 | 52 | cul3 | 26 |
| | | | ddb1 | 26 |
| | | | fus | 25 |
| NR4A3-TFG | tfg | 72 | cul3 | 25 |
| | | | ntrk1 | 25 |
| | | | ewsr1 | 24 |
| | | | cops5 | 23 |
| | | | cul1 | 20 |
| NR6A1-TRHDE | nr6a1 | 5 | akt1 | 3 |
| | | | ncoa1 | 3 |
| NUP107-LGR5 | nup107 | 31 | ntrk1 | 17 |
| | | | obsl1 | 16 |
| PAPPA-NUP107 | nup107 | 31 | ntrk1 | 18 |
| | | | obsl1 | 17 |
| | | | cul7 | 16 |
| | | | elavl1 | 16 |
| | | | sirt7 | 16 |
| | | | kpnb1 | 15 |
| | | | cul3 | 14 |
| | | | nup160 | 14 |
| PAX3-FOXO1 | foxo1 | 33 | akt1 | 18 |
| | crebbp | 23 | hdac1 | 18 |
| | ep300 | 23 | smad3 | 18 |
| | mdm2 | 22 | pax3 | 17 |
| | ar | 21 | smad4 | 16 |
| | esr1 | 19 | | |
| PAX7-FOXO1 | foxo1 | 33 | mdm2 | 17 |
| | ep300 | 23 | smad3 | 16 |
| | crebbp | 20 | sirt1 | 15 |
| | esr1 | 20 | smad4 | 15 |
| | akt1 | 18 | | |
| | ar | 17 | | |
| SS18-SSX2 | ss18 | 42 | smarcc1 | 27 |
| | | | smarca4 | 26 |
| | | | smarcb1 | 26 |
| | | | smarcc2 | 26 |
| | | | smarce1 | 26 |
| | | | smarca2 | 25 |
| | | | smarcd1 | 25 |
| | | | ssx2 | 22 |
| SS18-SSX1 | ss18 | 42 | smarcc1 | 27 |
| | | | smarcb1 | 26 |
| | | | smarcc2 | 25 |
| | | | smarcd1 | 25 |
| | | | smarca2 | 24 |
| | | | smarca4 | 24 |
| | | | smarce1 | 22 |
| | | | arid1a | 21 |

| | | | | |
|---|---|---|---|---|
| SS18L1-SSX1 | ss18l1 | 42 | smarca4 | 14 |
| | | | smarcc1 | 14 |
| | | | crebbp | 12 |
| SSX1-SYT4 | ntrk1 | 5 | nbr1 | 3 |
| | syt4 | 4 | ssx1 | 3 |
| TGFBR3-MGEA5 | mgea5 | 28 | ntrk1 | 16 |
| | | | hsp90aa1 | 14 |
| | | | tgfbr3 | 13 |
| | | | tpm3 | 13 |
| TRIO-TERT | tert | 64 | ntrk1 | 35 |
| | | | mdm2 | 30 |
| | | | tp53 | 27 |
| | | | trio | 21 |
| WDR70-RCOR1 | rcor1 | 46 | hdac2 | 33 |
| | | | hdac1 | 31 |
| | | | kdm1a | 31 |
| | | | ntrk1 | 19 |
| YWHAE-NUTM2B | ywhae | 337 | ywhaz | 138 |
| | | | ywhab | 128 |
| | | | ywhag | 126 |
| | | | ywhaq | 120 |
| | | | ntrk1 | 118 |
| YWHAE-NUTM2A-AS1 | YWHAE | 337 | YWHAZ | 138 |
| | | | YWHAB | 128 |
| | | | YWHAG | 126 |
| | | | YWHAQ | 120 |
| | | | NTRK1 | 118 |
| YWHAE-NUTM2A | YWHAE | 337 | YWHAZ | 138 |
| | | | YWHAB | 128 |
| | | | YWHAG | 126 |
| | | | YWHAQ | 120 |
| | | | NTRK1 | 118 |
| | | 106.43 | | 37.057 |

**Table S7: CA-Hubs (TRAINING)**

| **CA** | **Higher-degree Hubs** | | **Lower-degree Hubs** | |
|---|---|---|---|---|
| | **Hub** | **Degree** | **Hub** | **Degree** |
| ANK3-USP9Y | ank3 | 13 | usp9y | 7 |
| | | | ubc | 6 |
| | | | mapk6 | 5 |
| ARGLU1-CXCR4 | arglu1 | 23 | elavl1 | 15 |
| | cxcr4 | 22 | cul7 | 13 |
| | ntrk1 | 22 | jak2 | 12 |
| ATXN10-FBLN1 | fbln1 | 42 | fn1 | 14 |
| | atxn10 | 23 | app | 13 |

| | | | | |
|---|---|---|---|---|
| | | | cul3 | 13 |
| | | | egfr | 13 |
| | | | vcp | 12 |
| BCAS3-NFS1 | nfs1 | 15 | ctbp1 | 7 |
| | bcas3 | 10 | hdac5 | 7 |
| | | | esr1 | 6 |
| BCAS4-BCAS3 | bcas4 | 11 | bloc1s2 | 7 |
| | bcas3 | 10 | bloc1s3 | 7 |
| | bloc1s6 | 9 | dtnbp1 | 7 |
| | bloc1s1 | 8 | | |
| BCL2L12-PRMT1 | prmt1 | 134 | ntrk1 | 37 |
| | | | ep300 | 32 |
| | | | cand1 | 29 |
| | | | hnrnpk | 29 |
| | | | ncl | 28 |
| | | | cul3 | 27 |
| | | | ilf3 | 26 |
| | | | fus | 23 |
| | | | brca1 | 22 |
| CAPNS1-WDR62 | capns1 | 58 | ywhae | 25 |
| | | | ywhah | 22 |
| | | | ywhaz | 20 |
| | | | asns | 18 |
| CCDC6-ANK3 | ccdc6 | 33 | ntrk1 | 19 |
| | | | elavl1 | 17 |
| | | | hdac1 | 16 |
| | | | nr3c1 | 14 |
| | | | ank3 | 13 |
| | | | trim28 | 12 |
| CCDC9-DHX34 | ccdc9 | 11 | eif4a3 | 7 |
| CDC27-ST7L | cdc27 | 86 | cdc20 | 46 |
| | | | anapc4 | 39 |
| | | | fzr1 | 38 |
| | | | anapc7 | 37 |
| | | | cdc16 | 37 |
| | | | anapc1 | 33 |
| | | | cdc23 | 33 |
| CDK7-RIN3 | cdk7 | 78 | tp53 | 29 |
| | | | ccnh | 28 |
| | | | cdk9 | 28 |
| | | | brca1 | 27 |
| | | | cdk2 | 27 |
| | | | myc | 27 |
| | | | polr2a | 27 |
| CHERP-CPAMD8 | cherp | 46 | ntrk1 | 25 |
| | | | prpf40a | 22 |
| | | | obsl1 | 19 |
| | | | rbm39 | 19 |

| | | | | |
|---|---|---|---|---|
| | | | srpk2 | 19 |
| CPD-PIGW | cpd | 13 | elavl1 | 6 |
| | | | ntrk1 | 5 |
| CYTH1-PRPSAP1 | prpsap1 | 26 | cops5 | 10 |
| | | | cyth1 | 10 |
| DLG1-CRYBG3 | dlg1 | 45 | ntrk1 | 12 |
| | | | cask | 9 |
| | | | lin7a | 9 |
| DTX4-CCDC102B | ccdc102b | 55 | trim27 | 23 |
| | | | trim54 | 22 |
| | | | kifc3 | 17 |
| EHD4-FSIP1 | ehd4 | 32 | ntrk1 | 16 |
| | | | ehd1 | 13 |
| | | | egfr | 11 |
| ELK4-SLC26A9 | elk4 | 8 | mapk1 | 5 |
| | brca1 | 6 | mapk3 | 5 |
| EMID1-CBY1 | cby1 | 8 | emid1 | 6 |
| | | | kras | 4 |
| EPHA6-CNTN6 | epha6 | 1 | cntn6 | 1 |
| ERAL1-DIDO1 | dido1 | 30 | eral1 | 21 |
| | | | cul3 | 18 |
| | | | cand1 | 17 |
| GMDS-CCND3 | ccnd3 | 48 | cdk2 | 19 |
| | | | gmds | 19 |
| | | | cdkn1a | 15 |
| HJURP-EIF4E2 | eif4e2 | 58 | hjurp | 19 |
| | | | aes | 17 |
| | | | huwe1 | 17 |
| INTS4-GAB2 | gab2 | 33 | ntrk1 | 26 |
| | | | grb2 | 25 |
| | | | ints4 | 25 |
| KCNQ5-RIMS1 | rims1 | 9 | kcnq5 | 5 |
| KDM5A-ANO2 | KDM5A | 22 | RBBP7 | 13 |
| | HDAC1 | 20 | MORF4L1 | 11 |
| | HDAC2 | 20 | EZH2 | 11 |
| | | | HIST3H3 | 11 |
| | | | RB1 | 10 |
| | | | SUZ12 | 10 |
| LAMA5-C12orf28 | LAMA5 | 16 | USP4 | 3 |
| | | | SMAD2 | 3 |
| | | | PLAT | 3 |
| | | | MYOC | 3 |
| | | | FBLN2 | 3 |
| | | | MEP1A | 3 |
| MAPK10-FAM13A | mapk10 | 44 | mapk9 | 19 |
| | | | tp53 | 14 |
| MAPRE1-TM9SF4 | mapre1 | 73 | elavl1 | 26 |
| | | | ntrk1 | 26 |

| | | | | |
|---|---|---|---|---|
| | | | cops5 | 22 |
| | | | tubb | 17 |
| | | | app | 16 |
| | | | tm9sf4 | 16 |
| | | | ywhaz | 16 |
| NCKAP5-MZT2A | mzt2a | 6 | apc | 4 |
| | | | tubgcp2 | 4 |
| | | | tubgcp3 | 4 |
| NUMB-ALDH6A1 | numb | 20 | egfr | 15 |
| | | | aldh6a1 | 13 |
| | | | app | 12 |
| | | | mdm2 | 10 |
| PARD6B-CD48 | pard6b | 30 | pard6a | 12 |
| | | | prkci | 12 |
| | | | prkcz | 12 |
| | | | npm1 | 11 |
| | | | pard6g | 10 |
| | | | ywhah | 10 |
| PPP1R12A-MGAT4C | ppp1r12a | 47 | rpa1 | 21 |
| | | | ntrk1 | 17 |
| | | | elavl1 | 15 |
| | | | nsun2 | 14 |
| RNF11-C8A | rnf11 | 94 | app | 22 |
| | ubc | 60 | psmd4 | 20 |
| | | | eps15 | 19 |
| | | | nedd4 | 19 |
| SIPA1L3-WDR62 | wdr62 | 16 | elavl1 | 8 |
| | | | sipa1l3 | 8 |
| SLC26A6-PRKAR2A | prkar2a | 37 | prkaca | 16 |
| | | | prkar2b | 13 |
| | | | elavl1 | 12 |
| | | | prkacb | 10 |
| ST14-APLP2 | aplp2 | 23 | brca1 | 7 |
| | | | elavl1 | 6 |
| | | | jun | 6 |
| STRADB-NOP58 | nop58 | 94 | rps4x | 46 |
| | fbl | 64 | snu13 | 46 |
| STX16-RAE1 | nop56 | 63 | cul7 | 45 |
| | rnf2 | 56 | eed | 45 |
| | nop2 | 55 | rpl7 | 45 |
| | cand1 | 52 | rps7 | 43 |
| | sirt7 | 52 | rps8 | 43 |
| | rpl5 | 50 | gnl3 | 41 |
| | ntrk1 | 49 | rpl11 | 41 |
| | obsl1 | 49 | rpl4 | 41 |
| | ddx18 | 47 | hnrnpu | 40 |
| | rpl6 | 47 | nifk | 40 |

| | | | | |
|---|---|---|---|---|
| | | | rpl18a | 40 |
| | | | rps6 | 40 |
| TANC2-CHD6 | rae1 | 120 | obsl1 | 40 |
| | | | cul3 | 39 |
| | | | nxf1 | 39 |
| | | | cul7 | 37 |
| | | | tp53 | 25 |
| THSD7B-DARS | dars | 68 | ntrk1 | 40 |
| | | | iars | 38 |
| | | | eprs | 34 |
| | | | mars | 32 |
| | | | qars | 32 |
| | | | rplp0 | 32 |
| | | | cul3 | 31 |
| | | | fn1 | 30 |
| TMEM123-MMP7 | mmp7 | 23 | elavl1 | 12 |
| | | | cnot7 | 11 |
| | | | cnot6 | 9 |
| | | | gid8 | 9 |
| TMPRSS2-ERG | erg | 77 | hnrnpa1 | 42 |
| | | | elavl1 | 26 |
| | | | npm1 | 23 |
| | | | hnrnpm | 21 |
| | | | top1 | 21 |
| TOX3-CNTN5 | | | toxc3 | 1 |
| | | | cntn5 | 1 |
| UBR2-XPO5 | | | ubr2 | 1 |
| | | | xpo5 | 1 |
| UVRAG-INTS4 | | | uvrag | 1 |
| | | | ints4 | 1 |
| WNT11-TSPAN8 | | | wnt11 | 1 |
| | | | tspan8 | 1 |
| XRCC5-ACADL | | | xrcc5 | 1 |
| | | | acadl | 1 |
| ZCCHC7-PRSS3 | | | zcchc7 | 1 |
| | | | prss3 | 1 |
| ZFP91-NOX4 | | | zfp91 | 1 |
| | | | nox4 | 1 |
| | | 39.375 | | 13.404 |

**Table S8: LK, LY, ME, GL_Breakdown (TRAINING)**

| **LK, LY, ME, GL** | **Site-Directed Percolation Breakdown Points** |
|---|---|
| BCR-ABL1 | abl1+ntrk1+egfr+grb2 |
| RUNX1-RUNX1T1 | runx1+runx1t1+hdac1 |
| KMT2A-MLLT10 | kmt2a+tp53 |
| IGH-BCL2 | bcl2+tp53 |

| | |
|---|---|
| KMT2A-AFF1 | kmt2a |
| PICALM-MLLT10 | picalm+ntrk1 |
| PML-RARA | pml+rara+tp53 |
| KMT2A-MLLT3 | kmt2a+tp53 |
| KMT2A-AFDN | kmt2a+mllt4+tp53 |
| CBFB-MYH11 | cbfb+myh11 |
| IGH-MYC | igh |
| NUP98-DDX10 | nup98 |
| PCM1-JAK2 | jak2+ntrk1 |
| KMT2A-SEPT9 | kmt2a+sept9 |
| FUS-ERG | fus+ntrk1+cul3+cul7+fn1+cops5+prpf8 |
| NUP98-HOXA9 | nup98+nxf1 |
| ETV6-ABL1 | abl1+ntrk1+egfr+grb2 |
| SET-NUP214 | set+nup214+ntrk1 |
| MNX1-ETV6 | etv6 |
| KMT2A-MLLT1 | kmt2a+tp53+hdac1+rnf2 |
| KMT2A-MLLT6 | kmt2a+tp53+elavl1+crebbp+hdac1+rnf2 |
| ETV6-ACSL6 | etv6 |
| ETV6-MECOM | etv6+hdac1+mecom |
| KMT2A-EPS15 | eps15+kmt2a |
| KMT2A-GAS7 | kmt2a+tp53+crebbp |
| KMT2A-ABL1 | abl1+ntrk1+kmt2a+tp53 |
| KMT2A-MLLT11 | kmt2a+tp53+hdac1+hist3h3 |
| MN1-ETV6 | etv6 |
| KMT2A-MAML2 | kmt2a |
| KMT2A-FOXO4 | kmt2a |
| DEK-NUP214 | nup214+dek |
| RUNX1-CBFA2T3 | runx1+ep300 |
| NUP98-PSIP1 | nup98+psip1 |
| NUP98-HOXC13 | nup98 |
| NUP98-HOXC11 | nup98 |
| PAX5-ETV6 | etv6 |
| NUP98-HOXA11 | nup98 |
| BCR-PDGFRA | bcr+egfr |
| BCR-FGFR1 | bcr+fgfr1 |
| NPM1-RARA | rara+hdac1 |
| KMT2A-CBL | kmt2a |
| IGH-BCL6 | bcl6 |
| LCP1-BCL6 | bcl6 |
| CREBBP-KAT6A | kmt2a+tp53 |
| KMT2A-ARHGAP26 | kmt2a+tp53 |
| FOXO3-KMT2A | kmt2a+tp53 |
| KMT2A-DCPS | kmt2a+ntrk1 |
| KMT2A-EP300 | kmt2a+tp53 |
| IGH-CEBPE | cebpe |
| HSP90AA1-BCL6 | bcl6 |

**Table S9: SC_Breakdown (TRAINING)**

| SC | Site-Directed Percolation |
|---|---|
| | Breakdown Points |
| ASPSCR1-TFE3 | aspscr1+ntrk1+tfe3 |
| ASTN2-CNOT2 | cnot2+cnot1+cnot3+cnot7+cnot8 |
| BCOR-ZC3H7B | bcor |
| BCOR-CCNB3 | bcor+rnf2 |
| CDX1-IRF2BP2 | irf2bp2 |
| CIC-DUX4 | cic |
| CREB1-EWSR1 | ewsr1+creb1 |
| CTDSP2-FAM19A2 | ctdsp2 |
| CXorf67-MBTD1 | kat5+mbtd1+morf4l1 |
| EPC1-PHF1 | phf1+kat5 |
| ERG-EWSR1 | ewsr1+ntrk1+cul3+hnrnpa1+cps5+cand1 |
| ETV6-NTRK3 | etv6 |
| EWSR1-ATF1 | ewsr1 |
| EWSR1-FLI1 | ewsr1 |
| EWSR1-NR4A3 | ewsr1 |
| EWSR1-ETV4 | ewsr1 |
| EWSR1-PATZ1 | ewsr1 |
| EWSR1-DDIT3 | ewsr1+ntrk1 |
| EWSR1-POU5F1 | ewsr1+cul3 |
| EWSR1-SP3 | ewsr1+ntrk1 |
| FOXO4-CIC | foxo4+cic+crebbp+mdm2+smad3 |
| FUS-ERG | fus+ntrk1+cul3 |
| FUS-CREB3L1 | fus+ntrk1+cul3 |
| FUS-DDIT3 | fus+ntrk1+cul3+cul7+fn1+cops5 |
| FUS-ATF1 | fus+ntrk1+cul3+cul7+fn1+cops5 |
| FUS-CREB3L2 | fus+ntrk1+cul3+cul7 |
| HEY1-NCOA2 | ncoa2 |
| IRX2-TERT | tert |
| JAZF1-SUZ12 | SUZ12+rnf2+eed |
| JAZF1-PHF1 | phf1 |
| LMNA-NTRK1 | lmna+ntrk1 |
| MEAF6-TRERF1 | meaf6 |
| MEAF6-PHF1 | phf1+meaf6 |
| NR4A3-TAF15 | taf15+cul3+ddb1 |
| NR4A3-TFG | tfg |
| NR6A1-TRHDE | nr6a1 |
| NUP107-LGR5 | nup107 |
| PAPPA-NUP107 | nup107+ntrk1+obsl1+cul7+elavl1 |
| PAX3-FOXO1 | foxo1+crebbp+ep300 |
| PAX7-FOXO1 | foxo1+crebbp+ep300+esr1 |
| SS18-SSX2 | ss18+smarcc1 |
| SS18-SSX1 | ss18+smarcc1 |
| SS18L1-SSX1 | ss18l1 |
| SSX1-SYT4 | ntrk1+syt4 |
| TGFBR3-MGEA5 | mgea5 |
| TRIO-TERT | tert+ntrk1 |

| | |
|---|---|
| WDR70-RCOR1 | rcor1+hdac1+hdac2+kdm1a |
| YWHAE-NUTM2B | ywhae |
| YWHAE-NUTM2A-AS1 | YWHAE |
| YWHAE-NUTM2A | YWHAE |

**Table S10: CA_Breakdown (TRAINING)**

| CA | Site-Directed Percolation |
|---|---|
| | **Breakdown Points** |
| ANK3-USP9Y | ank3 |
| ARGLU1-CXCR4 | arglu1+cxcr4+ntrk1 |
| ATXN10-FBLN1 | fbln1+atxn10+egfr |
| BCAS3-NFS1 | nfs1+bcas3 |
| BCAS4-BCAS3 | bcas4+bcas3 |
| BCL2L12-PRMT1 | prmt1+ntrk1 |
| CAPNS1-WDR62 | capns1 |
| CCDC6-ANK3 | ccdc6+ntrk1 |
| CCDC9-DHX34 | ccdc9 |
| CDC27-ST7L | cdc27+cdc20 |
| CDK7-RIN3 | cdk7+tp53 |
| CHERP-CPAMD8 | cherp+ntrk1 |
| CPD-PIGW | cpd |
| CYTH1-PRPSAP1 | prpsap1 |
| DLG1-CRYBG3 | dlg1 |
| DTX4-CCDC102B | ccdc102b |
| EHD4-FSIP1 | ehd4+ntrk1 |
| ELK4-SLC26A9 | elk4+brca1 |
| EMID1-CBY1 | cby1 |
| EPHA6-CNTN6 | epha6 |
| ERAL1-DIDO1 | dido1 |
| GMDS-CCND3 | ccnd3+cdk2 |
| HJURP-EIF4E2 | eif4e2 |
| INTS4-GAB2 | gab2+ntrk1+grb2 |
| KCNQ5-RIMS1 | rims1 |
| KDM5A-ANO2 | KDM5A+hdac1+hdac2 |
| LAMA5-C12orf28 | LAMA5 |
| MAPK10-FAM13A | mapk10 |
| MAPRE1-TM9SF4 | mapre1 |
| NCKAP5-MZT2A | mzt2a |
| NUMB-ALDH6A1 | numb+egfr |
| PARD6B-CD48 | pard6b+pard6a |
| PPP1R12A-MGAT4C | ppp1r12a |
| RNF11-C8A | rnf11+ubc |
| SIPA1L3-WDR62 | wdr62 |
| SLC26A6-PRKAR2A | prkar2a |
| ST14-APLP2 | aplp2 |

| | |
|---|---|
| STRADB-NOP58 | nop58+fbl |
| STX16-RAE1 | nop56+rnf2+nop2+cand1+ntrk1+ddx18 |
| TANC2-CHD6 | rae1+tp53 |
| THSD7B-DARS | dars+ntrk1 |
| TMEM123-MMP7 | mmp7 |
| TMPRSS2-ERG | erg+hnrnpa1 |
| TOX3-CNTN5 | toxc3+cntn5 |
| UBR2-XPO5 | ubr2+xpo5 |
| UVRAG-INTS4 | uvrag+ints4 |
| WNT11-TSPAN8 | wnt11+tspan8 |
| XRCC5-ACADL | xrcc5+acadl |
| ZCCHC7-PRSS3 | zcchc7+prss3 |
| ZFP91-NOX4 | zfp91+nox4 |

**Table S11: LK, LY, ME, GL-community (TRAINING)**

| | Essential Community |
|---|---|
| BCR-ABL1 | |
| RUNX1-RUNX1T1 | HDAC1 BRCA1 KMT2A SMARCC1 HDAC2 CREBBP CTBP1 |
| | EP300 SMARCA4 NCOR2 NCOR1 |
| | VDR EP300 SMARCA4 CREBBP |
| KMT2A-MLLT10 | SMARCC2 KMT2A HDAC2 CHD3 SMARCC1 POLR2A |
| | SMARCA2 CREBBP SIN3A |
| | CTBP1 KMT2A CREBBP |
| IGH-BCL2 | TP53 CASP3 PARP1 CASP8 HIF1A BCL2 |
| | BAG3 PARP1 BCL2 |
| KMT2A-AFF1 | SMARCC2 KMT2A HDAC2 CHD3 SMARCC1 POLR2A |
| | SMARCA2 CREBBP SIN3A |
| | CTBP1 KMT2A CREBBP |
| PICALM-MLLT10 | FN1 EEF1A1 EGFR NTRK1 PLCG1 DNM2 ILVBL PICALM |
| | HNRNPD FUS FN1 DDX1 |
| | SEC24D PICALM NTRK1 SEC24C |
| PML-RARA | NCOA2 NR3C1 NR4A1 KAT2B RXRA PPARG TP53 MDM2 EP300 |
| | SMARCA4 RELA RARA NCOA3 NCOA1 STAT3 ARNT NPAS2 |
| | PARP1 NFKB1 TRIP4 CREBBP NFKB1 EP300 PARP1 |
| KMT2A-MLLT3 | SMARCC2 KMT2A HDAC2 CHD3 SMARCC1 POLR2A SMARCA2 |
| | CREBBP SIN3A CTBP1 KMT2A CREBBP |
| KMT2A-AFDN | SMARCC2 KMT2A HDAC2 CHD3 SMARCC1 POLR2A SMARCA2 |
| | CREBBP SIN3A |
| | CTBP1 KMT2A CREBBP |
| CBFB-MYH11 | ACTA2 MYH11 MYO1E RPA1 RPA2 ACTB |
| | RPA1 ELAVL1 RPA2 |
| IGH-MYC | |

| | |
|---|---|
| NUP98-DDX10 | SIRT7 DDX10 APP DDX56 NTRK1 DDX54 PUM3 PWP1 CSNK2A1 |
| | HDAC1 CTNNB1 MAPK8 EP300 CREBBP |
| | PUM3 NXF1 EED KPNB1 HNRNPUL1 |
| | HDAC1 CREBBP APC CSNK2A1 |
| | CTNNB1 APC CREBBP |
| | USP7 NTRK1 CDC37 |
| PCM1-JAK2 | ERBB2 ERBB3 VAV1 TEC EGFR INSR PLCG1 JAK2 |
| | STAT5A JAK2 INSR |
| KMT2A-SEPT9 | SMARCC2 KMT2A HDAC2 CHD3 SMARCC1 POLR2A SMARCA2 |
| | CREBBP SIN3A |
| | CTBP1 KMT2A CREBBP |
| FUS-ERG | RPA1 SF3B2 PRKDC PRPF8 SF3A2 DHX15 RPA2 CUL3 |
| | ABL1 PARP1 PRKDC |
| NUP98-HOXA9 | HDAC1 TP53 SMAD4 CTNNB1 CSNK2A1 MAPK8 EP300 CREBBP |
| | CTNNB1 APC CREBBP |
| ETV6-ABL1 | ERBB2 CBLB UBASH3B SOS1 ERBB4 SRC VAV1 EGFR CBL |
| | PLCG1 PIK3R2 PIK3R1 ABL1 |
| | ABL1 ABL2 JAK1 |
| SET-NUP214 | NXF1 FAF1 SUPT5H GART CUL2 CUL3 |
| | NXF1 CUL2 RANBP2 |
| MNX1-ETV6 | HDAC3 ETV6 HDAC9 PIN1 NCOR1 SIN3A |
| | L3MBTL1 ETV7 ETV6 |
| KMT2A-MLLT1 | SMARCC2 KMT2A HDAC2 CHD3 SMARCC1 POLR2A |
| | SMARCA2 CREBBP SIN3A |
| KMT2A-MLLT6 | SMARCC2 KMT2A HDAC2 CHD3 SMARCC1 POLR2A |
| | SMARCA2 CREBBP SIN3A |
| | CTBP1 KMT2A CREBBP |
| ETV6-ACSL6 | HDAC3 ETV6 HDAC9 PIN1 NCOR1 SIN3A |
| | L3MBTL1 ETV7 ETV6 |
| ETV6-MECOM | HDAC1 UBE2I HDAC3 EHMT2 ELAVL1 |
| | SMAD3 KAT2B SUV39H1 NCOR1 CTBP1 |
| | MECOM SMAD1 SMAD2 |
| | CREBBP MECOM SUV39H1 HDAC4 |
| KMT2A-EPS15 | SMARCC2 KMT2A HDAC2 CHD3 SMARCC1 |
| | POLR2A SMARCA2 CREBBP SIN3A |
| | CTBP1 KMT2A CREBBP |
| KMT2A-GAS7 | SMARCC2 KMT2A HDAC2 CHD3 SMARCC1 |
| | POLR2A SMARCA2 CREBBP SIN3A |
| | CTBP1 KMT2A CREBBP |
| KMT2A-ABL1 | SMARCC2 KMT2A SMARCC1 CHD3 POLR2A |
| | SMARCA2 CREBBP ABL1 CBLB CBL |
| KMT2A-MLLT11 | SMARCC2 KMT2A HDAC2 CHD3 SMARCC1 |
| | POLR2A SMARCA2 CREBBP SIN3A |

| | |
|---|---|
| | CTBP1 KMT2A CREBBP |
| MN1-ETV6 | HDAC3 ETV6 HDAC9 PIN1 EP300 NCOR1 SIN3A |
| | HDAC3 EP300 KAT5 |
| KMT2A-MAML2 | SMARCC2 KMT2A SMARCC1 CHD3 SMARCA2 CREBBP |
| | SMARCA2 EP300 CREBBP |
| KMT2A-FOXO4 | SMARCC2 KMT2A HDAC2 CHD3 SMARCC1 |
| | POLR2A SMARCA2 CREBBP SIN3A |
| | CTBP1 KMT2A CREBBP |
| DEK-NUP214 | ESR1 KAT2B SMAD2 SMAD3 CDK2 DEK EP300 CREBBP |
| | NXF1 DHX15 CUL2 CUL3 |
| RUNX1-CBFA2T3 | KMT2A SMARCC1 SMARCA4 CREBBP |
| | EP300 CREBBP |
| NUP98-PSIP1 | HDAC1 KMT2A ESR1 CTNNB1 EP300 CREBBP |
| | NXF1 EIF4A3 SON |
| NUP98-HOXC13 | HDAC1 CTNNB1 MAPK8 EP300 CREBBP |
| | NXF1 HNRNPAB EED KPNB1 HNRNPUL1 |
| | HDAC1 CREBBP APC CSNK2A1 |
| | CTNNB1 APC CREBBP |
| NUP98-HOXC11 | HDAC1 STAT3 SP1 SMAD3 CTNNB1 MAPK8 EP300 CREBBP |
| | CTNNB1 APC CREBBP |
| PAX5-ETV6 | UBE2I HDAC3 TBP KAT5 RB1 PAX5 RUNX1 PIN1 EP300 NCOR1 |
| | MAPK1 EP300 HDAC6 |
| NUP98-HOXA11 | HDAC1 HDAC2 YY1 CTNNB1 CSNK2A1 MAPK8 EP300 CREBBP |
| | CTNNB1 APC CREBBP |
| BCR-PDGFRA | TGFBR2 PDGFRA SHC1 CRKL EGFR PLCG1 FES CRK ABL1 |
| | HCK PTPN6 BCR UBASH3B INPP5D SOS1 GRB2 NTRK1 |
| | CBL KIT DOK1 PIK3R2 PIK3R1 |
| | ABL1 TP53 RB1 BCR |
| BCR-FGFR1 | SRC ITK SOS1 ERBB3 VAV1 CBL PLCG1 ABL1 |
| | ABL1 HCK BCR CBL |
| NPM1-RARA | |
| KMT2A-CBL | |
| IGH-BCL6 | HDAC1 TP53 CTBP1 EP300 NCOR2 CREBBP |
| | HDAC2 SMARCA4 CREBBP |
| LCP1-BCL6 | HDAC1 TP53 CTBP1 EP300 NCOR2 CREBBP |
| | HDAC2 SMARCA4 CREBBP |
| CREBBP-KAT6A | SMARCC2 KMT2A HDAC2 CHD3 SMARCC1 POLR2A SMARCA2 |
| | CREBBP SIN3A |
| | CTBP1 KMT2A CREBBP |
| KMT2A-ARHGAP26 | SMARCC2 KMT2A HDAC2 CHD3 SMARCC1 POLR2A SMARCA2 |

| | |
|---|---|
| | CREBBP SIN3A |
| | CTBP1 KMT2A CREBBP |
| FOXO3-KMT2A | SMARCC2 KMT2A SMARCC1 CHD3 SMARCA2 CREBBP |
| | SMARCA2 EP300 CREBBP |
| KMT2A-DCPS | SMARCC2 KMT2A HDAC2 CHD3 SMARCC1 POLR2A SMARCA2 |
| | CREBBP SIN3A |
| | CTBP1 KMT2A CREBBP |
| KMT2A-EP300 | |
| IGH-CEBPE | UBE2I BATF DDIT3 CEBPG CEBPE FOS JUN STAT6 |
| | RB1 PIAS1 FOSL1 BATF3 BATF2 ATF4 MYB ATF3 |
| HSP90AA1-BCL6 | |

**Table S12: SC-community (TRAINING)**

| | Essential Community |
|---|---|
| ASPSCR1 -TFE3 | |
| | xxxxxxxxxxxxxxxxxx |
| ASTN2-CNOT2 | CNOT6L AURKA CNOT8 TNRC6C TNRC6B CNOT3 CNOT2 CNOT1 CNOT7 AGO2 |
| | xxxxx |
| | HDAC3 GPS2 CNOT2 NCOR2 NCOR1 |
| | xxxxxxxxxxxxxxxxxx |
| BCOR-ZC3H7B | HDAC3 HDAC4 SP1 CTBP1 NACC1 NCOR2 |
| | xxxxx |
| | HDAC1 CTBP1 NCOR2 |
| | xxxxxxxxxxxxxxxxxx |
| BCOR-CCNB3 | HDAC3 HDAC4 SP1 CTBP1 NACC1 NCOR2 |
| | HDAC1 CTBP1 NCOR2 |
| | xxxxxxxxxxxxxxxxxx |
| CDX1-IRF2BP2 | ELAVL1 NTRK1 IRF2BPL IRF2BP2 RBM39 |
| | xxxxxxxxxxxxxxxxxx |
| CIC-DUX4 | |
| | xxxxxxxxxxxxxxxxxx |
| CREB1-EWSR1 | BRCA1 EPAS1 MYOD1 ESR1 JUN POLR2A EP300 SMARCA4 EWSR1 CREBBP |
| | xxxxx |
| | NR3C1 EP300 SMARCA4 CREBBP |
| | xxxxx |
| | NONO SMARCA4 CUL3 |
| | xxxxxxxxxxxxxxxxxx |
| CTDSP2-FAM19A2 | SETD1A POLR2A INTS6 CTDSP1 CTDSP2 |

| | |
|---|---|
| | xxxxxxxxxxxxxxxxxx |
| CXorf67-MBTD1 | |
| | xxxxxxxxxxxxxxxxxx |
| EPC1-PHF1 | HDAC1 DHX9 TP53 ELAVL1 E2F6 RBBP7 RBBP4 EZH1 EZH2 EED XRCC6 XRCC5 PHF1 |
| | xxxxx |
| | HDAC1 TP53 YEATS4 TRIM27 KAT5 TRIM23 HIST1H2BA DMAP1 MYC XRCC6 MORF4L1 ING3 |
| | xxxxxxxxxxxxxxxxxx |
| ERG-EWSR1 | TP53 ESR1 PARP1 EP300 CREBBP XRCC6 |
| | xxxxx |
| | EPAS1 EP300 CREBBP |
| | xxxxxxxxxxxxxxxxxx |
| ETV6-NTRK3 | PDGFRB SHC1 CRKL GAB2 NTRK1 PLCG1 GRB2 |
| | xxxxx |
| | HDAC3 PIN1 ETV6 |
| | xxxxxxxxxxxxxxxxxx |
| EWSR1-ATF1 | PDGFRB SHC1 CRKL GAB2 NTRK1 PLCG1 GRB2 |
| | xxxxx |
| | HDAC3 PIN1 ETV6 |
| | xxxxxxxxxxxxxxxxxx |
| EWSR1-FLI1 | BRCA1 POLR2A ESR1 EP300 KAT2B CREBBP |
| | xxxxx |
| | EPAS1 EP300 CREBBP |
| | xxxxxxxxxxxxxxxxxx |
| EWSR1-NR4A3 | TSG101 DHX9 HDAC3 TRIM28 FUS RAD23A JUN PRMT1 ILK BMI1 ELK1 POLR2A CHERP ATXN3 EP300 IRF3 EPAS1 TP53 ESR1 NONO RPA1 NTRK1 RPA2 CUL4A CUL4B FASN CREBBP HBP1 CUL5 HLTF EWSR1 YBX1 CUL1 CUL2 CUL3 |
| | xxxxx |
| | HDAC2 CREBBP ESR1 |
| | xxxxx |
| | NONO FXR2 CUL3 |
| | xxxxxxxxxxxxxxxxxx |
| EWSR1-ETV4 | TSG101 DHX9 HDAC3 RFWD2 FUS RAD23A JUN PRMT1 ILK SMAD2 BMI1 ELK1 POLR2A CHERP ATXN3 EP300 IRF3 EPAS1 TP53 ESR1 NONO RPA1 NTRK1 RPA2 CUL4A CUL4B FASN CREBBP HBP1 CUL5 HLTF EWSR1 YBX1 CUL1 CUL2 CUL3 |
| | xxxxx |
| | HDAC2 CREBBP ESR1 |
| | xxxxx |
| | NONO FXR2 CUL3 |
| | xxxxxxxxxxxxxxxxxx |

| | |
|---|---|
| EWSR1-PATZ1 | TSG101 DHX9 HDAC3 RFWD2 FUS RAD23A JUN PRMT1 ILK SMAD2 BMI1 ELK1 POLR2A CHERP ATXN3 EP300 IRF3 EPAS1 TP53 ESR1 NONO RPA1 NTRK1 RPA2 CUL4A CUL4B FASN CREBBP HBP1 CUL5 HLTF EWSR1 YBX1 CUL1 CUL2 CUL3 |
| | xxxxx |
| | HDAC2 CREBBP ESR1 |
| | xxxxx |
| | NONO FXR2 CUL3 |
| | xxxxxxxxxxxxxxxxxx |
| EWSR1-DDIT3 | HDAC1 EPAS1 HDAC3 DDIT3 CEBPB ESR1 FOS JUN HBP1 POLR2A IRF3 TP53 EP300 EWSR1 CREBBP |
| | xxxxx |
| | DHX9 CUL4A CUL4B CUL5 CUL1 CUL2 CUL3 |
| | xxxxxxxxxxxxxxxxxx |
| EWSR1-POU5F1 | IRF3 EPAS1 HDAC3 TP53 ESR1 JUN HBP1 ETS2 CTNNB1 POLR2A EP300 EWSR1 CREBBP |
| | xxxxx |
| | NONO CUL2 CUL3 |
| | xxxxxxxxxxxxxxxxxx |
| EWSR1-SP3 | HDAC1 EPAS1 HDAC3 CEBPB ESR1 JUN HBP1 POLR2A IRF3 TP53 RELA EP300 EWSR1 CREBBP |
| | xxxxx |
| | DHX9 CUL4A CUL4B CUL5 CUL1 CUL2 CUL3 |
| | xxxxxxxxxxxxxxxxxx |
| FOXO4-CIC | XPO1 CTNNB1 VDR ESR1 SMAD4 SMAD3 SFN AKT1 FOXO4 MDM2 NLK CREBBP |
| | xxxxxxxxxxxxxxxxxx |
| FUS-ERG | RPA1 SF3B2 PRKDC PRPF8 SF3A2 DHX15 RPA2 CUL3 |
| | ABL1 PARP1 PRKDC |
| | xxxxxxxxxxxxxxxxxx |
| FUS-CREB3L1 | NONO CUL4A CUL4B DHX15 CUL5 CUL1 CUL2 CUL3 |
| | VCP FBXW11 CUL1 |
| | xxxxxxxxxxxxxxxxxx |
| FUS-DDIT3 | HDAC1 DDX17 EPAS1 RELA ESR1 JUN TP73 EWSR1 CTNNB1 DDX5 CDK2 DDIT3 MDM2 EP300 TRIP4 CREBBP |
| | VCP FBXW11 CUL1 |
| | xxxxxxxxxxxxxxxxxx |
| FUS-ATF1 | HDAC1 DDX17 EPAS1 RELA ESR1 JUN TP73 EWSR1 CTNNB1 DDX5 CDK2 DDIT3 MDM2 EP300 TRIP4 CREBBP |
| | VCP FBXW11 CUL1 |
| | xxxxxxxxxxxxxxxxxx |
| FUS-CREB3L2 | NONO CUL4A CUL4B DHX15 CUL5 CUL1 CUL2 CUL3 |
| | VCP FBXW11 CUL1 |
| | xxxxxxxxxxxxxxxxxx |

| | |
|---|---|
| HEY1-NCOA2 | BRCA1 RARA NR3C1 VDR STAT6 HNF4A PRMT1 CARM1 RXRA PPARG PPARD AR PPARA ESR2 EP300 NCOA2 NCOA3 NCOA1 TP53 ESR1 AHR ARNT THRB THRA NR1I3 NR1I2 PGR CREBBP PIAS3 |
| | NCOA2 NCOA1 UBR5 |
| | xxxxxxxxxxxxxxxxxx |
| IRX2-TERT | YWHAZ AKT1 RPS6KB1 ENO1 MTOR MDM2 TERT XRCC6 |
| | TERF1 STUB1 TERT POT1 |
| | TERT MTOR YWHAQ RUVBL2 |
| | TPP1 TERT POT1 |
| | xxxxxxxxxxxxxxxxxx |
| JAZF1-SUZ12 | DHX9 RBM5 FBXW11 DDX3X NXF1 SF3B4 PRMT1 SF3B1 SF3B2 PRPF8 EED CRNKL1 RNPS1 UBE2I SNRNP200 SRSF7 SNRPD3 RALY DDX5 PRPF19 RNF2 SNRPA1 SON EFTUD2 U2AF1 SF3A1 EPRS ILF2 EIF4A3 CDC40 ILF3 RANBP2 |
| | DNMT3B HDAC1 HDAC2 TRIM28 CHD4 UHRF1 DNMT1 MTA1 NR2C2 EZH2 GATAD2B EED CBX5 RBBP4 CBX3 SETDB1 |
| | VCP BRCA1 MTOR NXF1 RUVBL2 |
| | VCP FBXW11 SKP1 BTRC EZH2 |
| | DHX9 ADAR EZH2 FBXW11 |
| | HDAC2 NXF1 JARID2 SETDB1 |
| | RELA FBXW11 BTRC |
| | BTRC CSNK2B NXF1 |
| | xxxxxxxxxxxxxxxxxx |
| JAZF1-PHF1 | DHX9 EZH1 PPARG EZH2 EED XRCC6 XRCC5 PHF1 |
| | HDAC1 PPARG EZH2 PHF1 |
| | PHF1 TP53 XRCC6 |
| | xxxxxxxxxxxxxxxxxx |
| LMNA-NTRK1 | |
| | xxxxxxxxxxxxxxxxxx |
| MEAF6-TRERF1 | HDAC1 TRERF1 KAT5 ING3 CREBBP YEATS4 EP300 MORF4L1 HIST1H2BA |
| | ELAVL1 TRERF1 KAT6A CREBBP NR5A1 |
| | SOX2 HDAC1 TRERF1 |
| | xxxxxxxxxxxxxxxxxx |
| MEAF6-PHF1 | DHX9 EZH1 EZH2 EED XRCC6 XRCC5 PHF1 |
| | PHF1 TP53 XRCC6 |
| | KAT6A TP53 ELAVL1 |
| | HDAC1 EZH2 PHF1 |
| | xxxxxxxxxxxxxxxxxx |
| NR4A3-TAF15 | TRIM28 FUS PRMT1 COPS6 COPS5 POLR2C POLR2A TAF15 SF1 NEDD8 POLR2E RPA1 RPA2 CUL4A CUL4B CUL5 CUL1 CUL2 CUL3 |
| | EZH2 TRIM28 CUL1 |
| | xxxxxxxxxxxxxxxxxx |
| NR4A3-TFG | CUL4A CUL4B CUL5 CUL1 CUL2 CUL3 |

| | |
|---|---|
| | TRIM28 CUL1 CUL3 |
| | xxxxxxxxxxxxxxxxxx |
| NR6A1-TRHDE | |
| | xxxxxxxxxxxxxxxxxx |
| NUP107-LGR5 | NUP153 KPNB1 NTRK1 CUL3 |
| | NUP153 EIF4B CUL3 |
| | EED KPNB1 TP53BP1 |
| | xxxxxxxxxxxxxxxxxx |
| PAPPA-NUP107 | NUP153 ELAVL1 KPNB1 NTRK1 SMAD3 VCP CUL3 EIF4B |
| | SMAD9 SKIL PAPPA SMAD2 SMAD3 |
| | TP53BP1 ELAVL1 EED KPNB1 |
| | NUP214 SMAD2 SMAD3 |
| | xxxxxxxxxxxxxxxxxx |
| PAX3-FOXO1 | NCOA1 ESR1 PARP1 AR EP300 CREBBP |
| | TRIM28 PARP1 CREBBP |
| | xxxxxxxxxxxxxxxxxx |
| PAX7-FOXO1 | RARA NCOA1 MYOD1 ESR1 HNF4A PARP1 SMAD3 FOXO1 AR MDM2 EP300 CREBBP |
| | AKT1 EP300 CREBBP |
| | xxxxxxxxxxxxxxxxxx |
| SS18-SSX2 | DPF2 SMARCC2 SMARCC1 PHF10 ELAVL1 DPF3 ARID2 DPF1 SMARCD3 SMARCE1 SMARCD1 EED SMARCA2 EP300 SMARCA4 HDAC1 ARID1B ARID1A ACTL6A HDAC2 CUL3 RNF2 SMARCD2 |
| | GRB2 YWHAG CUL3 |
| | xxxxxxxxxxxxxxxxxx |
| SS18-SSX1 | SMARCC2 DPF2 SMARCC1 PHF10 ELAVL1 DPF3 DPF1 ARID2 HDAC2 SMARCD3 SMARCE1 SMARCD1 EED SMARCA2 EP300 SMARCA4 HDAC1 ARID1B ARID1A ACTL6A CUL3 SMARCD2 |
| | GRB2 YWHAG CUL3 |
| | xxxxxxxxxxxxxxxxxx |
| SS18L1-SSX1 | SMARCC1 STAT3 BMI1 WHSC1L1 SMAD3 HDAC2 SMAD1 EP300 SMARCA4 CREBBP |
| | DPF2 SMARCC1 SMARCE1 SMARCA4 CUL3 |
| | xxxxxxxxxxxxxxxxxx |
| SSX1-SYT4 | |
| | xxxxxxxxxxxxxxxxxx |
| TGFBR3-MGEA5 | MAST1 RNF32 PAXIP1 |
| | CBX8 CSNK2B PAXIP1 |
| | xxxxxxxxxxxxxxxxxx |
| TRIO-TERT | YWHAZ AKT1 RPS6KB1 ENO1 MTOR MDM2 TERT XRCC6 |
| | TERF1 STUB1 TERT POT1 |

| | |
|---|---|
| | TERT MTOR YWHAQ RUVBL2 |
| | xxxxxxxxxxxxxxxxxx |
| WDR70-RCOR1 | SMARCC2 HDAC1 HDAC3 HDAC2 KDM1A RCOR1 NR2C1 SMARCE1 CTBP1 NR2E1 SMARCA4 CTBP2 KDM5B MTA3 |
| | KDM1A HDAC3 CTBP1 |
| | xxxxxxxxxxxxxxxxxx |
| YWHAE-NUTM2B | LARP1 NOS2 YWHAQ YWHAG HUWE1 MAST2 NTRK1 VCP AKT1 MAP2K1 FBXW11 ARAF CUL3 PARK2 RAF1 YWHAZ UBXN1 TP53 RUVBL2 BTRC TUBB CDC37 YWHAH MAST3 BRAF YWHAB KSR1 YWHAE CUL1 MAPK7 |
| | YWHAZ IGF1R IRS1 YWHAQ MST1R NTRK1 CBL TUBB YWHAB YWHAH GRB2 ABL1 SORBS2 BCAR1 YWHAE |
| | VCP CDK2 CUL1 |
| | CDC37 LRRK2 YWHAE |
| | xxxxxxxxxxxxxxxxxx |
| YWHAE-NUTM2A-AS1 | LARP1 NOS2 YWHAQ YWHAG HUWE1 MAST2 NTRK1 VCP AKT1 MAP2K1 FBXW11 ARAF CUL3 PARK2 RAF1 YWHAZ UBXN1 TP53 RUVBL2 BTRC TUBB CDC37 YWHAH MAST3 BRAF YWHAB KSR1 YWHAE CUL1 MAPK7 |
| | YWHAZ IGF1R IRS1 YWHAQ MST1R NTRK1 CBL TUBB YWHAB YWHAH GRB2 ABL1 SORBS2 BCAR1 YWHAE |
| | VCP CDK2 CUL1 |
| | CDC37 LRRK2 YWHAE |
| | xxxxxxxxxxxxxxxxxx |
| YWHAE-NUTM2A | LARP1 NOS2 YWHAQ YWHAG HUWE1 MAST2 NTRK1 VCP AKT1 MAP2K1 FBXW11 ARAF CUL3 PARK2 RAF1 YWHAZ UBXN1 TP53 RUVBL2 BTRC TUBB CDC37 YWHAH MAST3 BRAF YWHAB KSR1 YWHAE CUL1 MAPK7 |
| | YWHAZ IGF1R IRS1 YWHAQ MST1R NTRK1 CBL TUBB YWHAB YWHAH GRB2 ABL1 SORBS2 BCAR1 YWHAE |
| | VCP CDK2 CUL1 |
| | CDC37 LRRK2 YWHAE |
| | xxxxxxxxxxxxxxxxx |

**Table S13: CA-community (TRAINING)**

| | Essential Community |
|---|---|
| ANK3-USP9Y | |
| ARGLU1-CXCR4 | APP CHERP SNRNP70 SRPK1 SRPK2 |
| | PTK2 JAK2 SOCS3 PTPN11 NTRK1 |
| | PTK2 ELAVL1 SRPK1 NTRK1 |
| | PTK2 JAK2 JAK3 SOCS3 PTPN11 STAM |
| ATXN10-FBLN1 | FN1 ATXN10 EGFR GSTK1 |
| | ATXN10 VCP BSG CUL3 |
| | VCP ABCE1 ATXN10 APP YWHAQ CUL3 |

| | |
|---|---|
| BCAS3-NFS1 | CTBP1 CTBP2 BCAS3 KAT2B |
| | CTBP1 CTBP2 CDC23 KAT2B BCAS3 |
| BCAS4-BCAS3 | CTBP1 CTBP2 BCAS3 KAT2B |
| | CTBP1 CTBP2 CDC23 KAT2B BCAS3 |
| BCL2L12-PRMT1 | NCOA2 NCOA3 NCOA1 TP53 ESR1 BRCA1 PRMT1 THRB CARM1 NR1I2 AR PPARA EP300 |
| | NCOA2 NCOA3 NCOA1 EP300 PARP1 |
| CAPNS1-WDR62 | YWHAZ FBXW11 FN1 HUWE1 YWHAQ GAPDH FERMT2 YWHAH VCAM1 YWHAB PAFAH1B1 YWHAG PAK2 YWHAE |
| | OGFOD1 MYO1E ASNS TBCB CAPN2 PROSC YWHAE |
| CCDC6-ANK3 | HDAC1 NR3C1 TRIM28 SF3A1 HNRNPR BRCC3 |
| | HDAC1 NR3C1 TRIM28 ELAVL1 SKP1 HNRNPR PPP1CA BRCC3 NTRK1 SF3A1 CUL1 FBXW7 |
| CCDC9-DHX34 | EIF4A3 SNIP1 CCDC9 PRPF40A |
| CDC27-ST7L | CDC16 CDC27 MDC1 CDC20 CREBBP ANAPC2 ANAPC7 |
| | CREBBP E2F1 RB1 TFDP1 |
| | CDC16 CDC27 MDC1 SMAD2 TP53BP1 CREBBP UBE2S ANAPC2 ANAPC7 |
| CDK7-RIN3 | BRCA1 TP53 RUVBL2 SUPT5H ESR1 RPA1 HNRNPU RPA2 CDK2 POLR2A |
| | GTF2H1 RPA1 RPA2 POLR2A CCNH |
| | HDAC2 TP53 ESR1 MTA1 |
| | GTF2H1 ERCC3 POLR2A ERCC5 |
| | BRCA1 POLR2A RUVBL2 GTF2H1 RPA1 RPA2 |
| | PRKCI APP CDK7 CDC37 |
| CHERP-CPAMD8 | DHX8 U2AF1 RPA1 PRPF40A RPA2 CHERP SF3A2 RBM39 EWSR1 |
| | DHX8 AGGF1 RNPS1 SNIP1 SF3B4 NTRK1 CHERP SRPK1 SRPK2 U2AF1 U2AF2 RPA1 APBB1 PRPF40A RPA2 TTC14 RBM23 SF3A2 SNRNP70 RBM39 EWSR1 WBP4 |
| CPD-PIGW | |
| CYTH1-PRPSAP1 | DDX17 DDX5 ILK COPS5 |
| | CYTH1 ARRB2 ARF6 ARRB1 |
| | DDX5 FBXW11 DDX17 ILK ITGB2 COPS5 |
| DLG1-CRYBG3 | DLG1 LIN7A LIN7C APBA1 CASK |
| | DLG1 NTRK1 CASK EPB41 |
| | DLG1 KHDRBS1 LCK NTRK1 |
| | MAPK1 ARRB2 ARRB1 |
| DTX4-CCDC102B | MCM7 CDK18 LENG1 TRIM54 TRIM27 KIFC3 |
| | SFN MARK1 CCDC102B |

| | |
|---|---|
| EHD4-FSIP1 | EHD4 CTPS2 EHD1 EGFR NTRK1 WARS PLCG1 UBA2 ADSL UQCRC2 |
| | PLCG1 EHD4 EGFR NTRK1 |
| ELK4-SLC26A9 | BRCA1 MAPK3 MAPK1 ELK4 |
| | BRCA1 MAPK3 MAPK1 ELK4 BLM |
| EMID1-CBY1 | xx |
| EPHA6-CNTN6 | xx |
| ERAL1-DIDO1 | HNRNPDL RPA1 RPA2 HNRNPK RBM15 CUL3 DIDO1 FUS |
| | FUS APP RPA1 RPA2 RBM15 CUL3 DIDO1 SRPK2 |
| GMDS-CCND3 | PCNA PPP1CC RBL2 PPP1CA CCND3 RB1 POLD1 CDK2 CDK4 CDK6 CREBBP |
| | GMDS NSFL1C CTH CAPN2 ATIC |
| | RARA NCOA2 VDR CCND3 CREBBP |
| | MCM10 RBX1 CCND3 APP |
| | NCOA2 RARA VDR CCND3 CREBBP |
| HJURP-EIF4E2 | FBXW11 TP53 GIGYF2 APP HUWE1 EIF4E2 YWHAB SHMT2 YWHAE |
| INTS4-GAB2 | SRC PLCG1 GRB2 ZAP70 NTRK1 SHC1 |
| | PIK3CB PIK3R2 PIK3R1 |
| KCNQ5-RIMS1 | |
| KDM5A-ANO2 | HDAC1 HDAC2 RBL1 TBP RB1 VDR KDM5A MORF4L1 |
| | HDAC2 EZH2 KDM5A ESR1 |
| LAMA5-C12orf28 | |
| MAPK10-FAM13A | HDAC1 TP53 HDAC9 JUN DDX5 ELK1 MAPK10 RELA CREBBP ATF2 |
| | APP MAPK10 MAP2K4 |
| MAPRE1-TM9SF4 | YWHAZ FN1 APP TUBB NTRK1 VCAM1 UNK COPS5 |
| | CDK5RAP2 PRKACA AKAP9 PRKACB |
| | CLIP1 TUBB TUBA1A HDAC6 |
| | PDE4DIP PRKACA PRKACB CDK5RAP2 AKAP9 |
| | TERF1 SPTAN1 DST MAPRE1 |
| NCKAP5-MZT2A | |
| NUMB-ALDH6A1 | TP53 NUMB ITCH MDM2 EGFR |
| | EPS15 EGFR AP2A1 NUMB |
| | PRKCZ NUMB APP EGFR |
| PARD6B-CD48 | PRKCI RASSF8 PARD3 PARD6G APP PARD6B PARD6A YWHAH PRKCZ WWC1 |
| | PRKCI PARD3 PARD6G APP PARD6B PARD6A YWHAH PRKCZ |
| | RAC1 PARD6G PARD6B PARD6A |

| | |
|---|---|
| PPP1R12A-MGAT4C | KDM1A ELAVL1 RPA1 RPA2 PPP1R12A CUL1 |
| | KDM1A NUDT5 TP53 ELAVL1 PUS1 NUAK1 AARSD1 RPA1 NTRK1 RPA2 RPRD1B PPP1R12A TRIM47 PAXIP1 ACTR3 CUL1 |
| RNF11-C8A | CBLB RNF11 ITCH SMAD4 EPN1 RABGEF1 UBE2E1 UBE2D3 UBE2E3 HGS GGA1 AKT1 GGA3 GGA2 AP2A1 EPN3 UBE2D1 AP2B1 CSNK2A1 SMURF1 EPS15 SMURF2 UBQLN2 STAM2 NEDD4 UBQLN4 NEDD4L APP |
| | RNF11 PSMD4 PSMD7 PSMD6 PSMD11 PSMD10 PSMD3 PSMD12 PSMD13 USP14 PSMD14 PSMD1 PSMD2 APP |
| | GGA1 GGA3 GGA2 APP |
| | AKT1 RNF11 TBK1 APP NEDD4 |
| SIPA1L3-WDR62 | MAPK10 WDR62 MAPK8 MAPK9 |
| | SFN YWHAB SIPA1L3 YWHAQ |
| | YWHAB FBXW11 MAPK10 ELAVL1 WDR62 TBP MAPK8 MAPK9 |
| SLC26A6-PRKAR2A | AKAP7 AKAP9 PRKAR2A PRKAR2B PRKACA PRKACB |
| | PRKAR2A AKAP7 PRKACA PRKACB PRKAR2B GCH1 |
| ST14-APLP2 | BRCA1 APLP2 ETS1 JUN |
| | SFN HDAC5 APLP2 RPL26 |
| | BRCA1 JUNB JUN APBB1 APBB2 KAT5 ETS1 APLP2 MAPK8 |
| STRADB-NOP58 | NIFK NOP56 HNRNPU RUVBL2 NOLC1 SNU13 NTRK1 PUM3 RPS15A RSL1D1 KRR1 DDX18 EIF6 DHX15 RPS4X EED PRPF3 TARDBP RPL11 EIF2S2 DDX27 NOP58 FN1 DDX24 U2AF1 RPL30 ESR1 DDX56 FTSJ3 DDX47 KPNA6 GTPBP4 WDR36 KPNA1 DKC1 RRP12 BOP1 FBL TBL3 |
| | WDR36 NOP58 DHX15 NOP56 |
| STX16-RAE1 | FBXW11 NXF1 FAF1 RAE1 ILF3 CUL1 HNRNPUL1 CUL3 |
| | NXF1 CUL1 ILF3 CUL3 |
| TANC2-CHD6 | ZFYVE9 PPP1CC PPP1CA TANC2 |
| THSD7B-DARS | ZFYVE9 PPP1CC PPP1CA TANC2 |
| TMEM123-MMP7 | MAEA RANBP9 MKLN1 MMP7 RMND5A |
| | MAEA RANBP9 MKLN1 RANBP10 MMP7 RMND5A |
| TMPRSS2-ERG | CDC5L DDX3X ELAVL1 CAD NEDD4 SF3B1 PARP1 PRKDC PRPF8 SFPQ ERG POLR2A TOP1 CLTC SF3B2 XRCC5 XRCC6 DDX23 SNRNP200 DDX21 TUBB NONO JUN HNRNPU PRPF40A AR NCL HNRNPM HNRNPC TOP2B ILF3 ILF2 |
| | PRPF8 ERG SF3B2 SF3B1 PARP1 PRKDC |
| TOX3-CNTN5 | |

| |
|---|
| UBR2-XPO5 |
| UVRAG-INTS4 |
| WNT11-TSPAN8 |
| XRCC5-ACADL |
| ZCCHC7-PRSS3 |
| ZFP91-NOX4 |

**Table S14: LK, LY, ME, GL_Preferential_attachment (TRAINING)**

| | Essential Community Vertices | Preferential attachment vertices |
|---|---|---|
| BCR-ABL1 | | |
| RUNX1-RUNX1T1 | HDAC1 BRCA1 | KMT2A SMARCC1 SMARCA4 CREBBP |
| KMT2A-MLLT10 | KMT2A HDAC2 SMARCC1 POLR2A SMARCA2 CREBBP SIN3A | HDAC1 SMARCC2 KMT2A SMARCC1 WDR5 ELAVL1 CBX4 DOT1L SENP3 BMI1 CTNNB1 HECW2 HDAC2 POLR2A CCNT1 SMARCA2 MAP3K5 CSNK2A2 KAT8 TAF6 CHD3 E2F4 TP53 TAF1 RAN TBP RBBP7 TOP1 RBBP4 CXXC1 RUNX1 MBD3 CTBP1 KAT6A MYB RNF2 CREBBP SIN3A |
| IGH-BCL2 | TP53 CASP3 BCL2 | BCL2L1 BAG3 TP53 CASP3 HIF1A PARP1 CASP8 BCL2 |
| KMT2A-AFF1 | SMARCC2 KMT2A HDAC2 CREBBP SIN3A | HDAC1 SMARCC2 KMT2A SMARCC1 WDR5 HECW2 CBX4 DOT1L SENP3 BMI1 CARM1 CTNNB1 HDAC2 POLR2A CCNT1 SMARCA2 MAP3K5 CSNK2A2 KAT8 TAF6 CHD3 E2F4 TP53 TAF1 RAN TBP RBBP7 TOP1 RBBP4 KAT6A CXXC1 RUNX1 MBD3 NSD1 CTBP1 RELA MYB RNF2 CREBBP SIN3A |
| PICALM-MLLT10 | FN1 EGFR NTRK1 PICALM | FN1 ITSN1 EGFR NTRK1 PLCG1 CLTC DNM2 ILVBL PICALM |
| PML-RARA | NR4A1 KAT2B PPARG TP53 RARA CREBBP | NCOA2 NCOA3 NCOA1 STAT3 ARNT WRN PARP1 NPAS2 EP300 SMARCA4 CREBBP |
| KMT2A-MLLT3 | KMT2A HDAC2 CREBBP SIN3A | KMT2A SMARCA2 CHD3 CREBBP |
| KMT2A-AFDN | KMT2A HDAC2 CREBBP SIN3A | SMARCC2 HDAC1 KMT2A SMARCC1 WDR5 HECW2 DOT1L SENP3 BMI1 SMAD2 CTNNB1 HDAC2 POLR2A CCNT1 SMARCA2 MAP3K5 CSNK2A2 TAF6 CHD3 CBX4 TP53 TAF1 RAN TBP RBBP7 RBBP4 CXXC1 RUNX1 E2F4 MBD3 CTBP1 KAT6A MYB RNF2 CREBBP SIN3A |
| CBFB-MYH11 | MYH11 RPA1 RPA2 ACTB | MYH11 MYO1E RPA1 RPA2 |
| IGH-MYC | | |
| NUP98-DDX10 | SIRT7 APP NTRK1 CSNK2A1 | CTNNB1 HDAC1 EP300 CREBBP |

| | | |
|---|---|---|
| PCM1-JAK2 | EGFR JAK2 | ERBB2 ERBB3 PTK2 VAV1 TEC EGFR INSR PLCG1 JAK2 STAT5A |
| KMT2A-SEPT9 | KMT2A HDAC2 SMARCA2 CREBBP | SMARCC2 KMT2A SMARCC1 TAF1 CHD3 HIF1A POLR2A CTBP1 SMARCA2 KAT6A CREBBP |
| FUS-ERG | RPA1 SF3B2 RPA2 CUL3 | PRPF8 DHX15 CUL3 RPA2 RPA1 SF3B2 |
| NUP98-HOXA9 | TP53 CREBBP | CTNNB1 HDAC1 TP53 EP300 CREBBP |
| ETV6-ABL1 | ERBB2 SRC EGFR CBL ABL1 | ERBB2 ERBB3 SHC1 ERBB4 CBLB SPTAN1 EGFR SRC SPTA1 SORBS1 PLCG1 CRK ABL1 HCK ABL2 BCAR1 RASA1 BCR NCK1 UBASH3B INPPL1 SOS1 VAV1 GRB10 EPHB2 NTRK1 CBL SORBS2 GRB2 ZAP70 PIK3R1 JAK1 CRKL PIK3R2 |
| SET-NUP214 | NXF1CUL2 CUL3 | FARSB NXF1 FAF1 SUPT5H GART CUL3 |
| MNX1-ETV6 | ETV6 NCOR1 SIN3A | SOX2 HDAC3 SOCS3 ETV6 HDAC9 PIN1 NCOR1 SIN3A |
| KMT2A-MLLT1 | KMT2A SMARCA2 CREBBP SIN3A | HDAC1 SMARCC2 KMT2A SMARCC1 WDR5 HECW2 CBX4 DOT1L SENP3 BMI1 CTNNB1 HDAC2 POLR2A CCNT1 SMARCA2 MAP3K5 CSNK2A2 CSNK2A1 KAT8 TAF6 CHD3 E2F4 TP53 TAF1 RAN TBP RBBP7 TOP1 RBBP4 CXXC1 RUNX1 MBD3 CTBP1 KAT6A MYB RNF2 CREBBP SIN3A |
| KMT2A-MLLT6 | KMT2A HDAC2 SMARCA2 CREBBP SIN3A | HDAC1 SMARCC2 KMT2A SMARCC1 HDAC2 POLR2A SMARCA2 TAF6 CHD3 TP53 TAF1 RAN TBP WHSC1L1 RUNX1 MBD3 CTBP1 KAT6A MYB CREBBP SIN3A |
| ETV6-ACSL6 | HDAC3 ETV6 HDAC9 | SOX2 HDAC3 SOCS3 ETV6 HDAC9 PIN1 NCOR1 SIN3A |
| ETV6-MECOM | HDAC1 UBE2I HDAC3 ELAVL1 MECOM CREBBP | HDAC1 HDAC3 CTBP1 KAT2B CREBBP |
| KMT2A-EPS15 | KMT2A HDAC2 CREBBP SIN3A | SMARCC2 HDAC1 KMT2A SMARCC1 TP53 TAF1 TBP CHD3 HDAC2 POLR2A CTBP1 SMARCA2 KAT6A CREBBP SIN3A |
| KMT2A-GAS7 | KMT2A HDAC2 SMARCA2 CREBBP | HDAC1 SMARCC2 KMT2A SMARCC1 CTNNB1 HDAC2 POLR2A SMARCA2 TAF6 CHD3 TP53 TAF1 RAN TBP MAP3K5 RUNX1 MBD3 CTBP1 KAT6A MYB CREBBP SIN3A |
| KMT2A-ABL1 | KMT2A SMARCC1 SMARCA2 CREBBP | SMARCC2 NCOA3 KMT2A SMARCA2 CHD3 CREBBP PARP1 |
| KMT2A-MLLT11 | KMT2A HDAC2 SMARCA2 CREBBP | HDAC1 SMARCC2 KMT2A SMARCC1 WDR5 HECW2 CBX4 CUL3 SENP3 BMI1 CTNNB1 HDAC2 POLR2A DOT1L CCNT1 SMARCA2 MAP3K5 CSNK2A2 KAT8 TAF6 CHD3 E2F4 TP53 TAF1 RAN TBP RBBP7 TOP1 RBBP4 CXXC1 RUNX1 MBD3 CTBP1 KAT6A MYB RNF2 CREBBP SIN3A |
| MN1-ETV6 | HDAC3 ETV6 EP300 | NCOA3 PIN1 EP300 NCOR1 |
| KMT2A-MAML2 | KMT2A SMARCA2 CREBBP | HDAC1 SMARCC2 HDAC2 WDR5 CTNNB1 SMARCC1 CCNT1 SMARCA2 CBX4 MYB CHD3 MAML2 MAML1 RAN MAP3K5 DOT1L SIN3A KMT2A MBD3 SENP3 BMI1 POLR2A EP300 CSNK2A2 KAT8 TAF6 E2F4 TP53 TAF1 TBP RBBP7 TOP1 RBBP4 CXXC1 RUNX1 HECW2 CTBP1 KAT6A RNF2 CREBBP |
| KMT2A-FOXO4 | KMT2A HDAC2 SMARCC1 CREBBP SIN3A | HDAC1 SMARCC2 KMT2A SMARCC1 HDAC2 POLR2A SMARCA2 MDM2 TAF6 CHD3 TP53 TAF1 |

| | | |
|---|---|---|
| | | ESR1 RAN TBP RUNX1 MBD3 CTBP1 KAT6A MYB CREBBP SIN3A |
| DEK-NUP214 | ESR1 DEK CREBBP | DEK EP300 KAT2B CREBBP |
| RUNX1-CBFA2T3 | KMT2A CREBBP | KMT2A SMARCC1 SMARCA4 CREBBP |
| NUP98-PSIP1 | HDAC1 KMT2A CREBBP | HDAC1 KMT2A ESR1 EP300 CREBBP |
| NUP98-HOXC13 | HDAC1 EP300 CREBBP | HDAC1 CTNNB1 EP300 CREBBP |
| NUP98-HOXC11 | HDAC1 EP300 CREBBP | HDAC1 STAT3 TBX21 SP1 SMAD3 CTNNB1 MAPK8 EP300 CREBBP |
| PAX5-ETV6 | HDAC3 RUNX1 EP300 | HDAC3 TBP EP300 RB1 |
| NUP98-HOXA11 | HDAC1 MAPK8 EP300 CREBBP | HDAC1 CTNNB1 EP300 CREBBP |
| BCR-PDGFRA | PDGFRA EGFR ABL1 BCR | PDGFRB PDGFRA SHC1 CRKL TGFBR2 EGFR PLCG1 RB1 CRK ABL1 HCK PTPN6 BCR INPP5D UBASH3B TP53 SOS1 STUB1 GRB2 NTRK1 CBL KIT DOK1 PIK3R2 PIK3R1 FES CDC42 |
| BCR-FGFR1 | SRC ABL1 | SRC ITK SOS1 ERBB3 VAV1 CBL PLCG1 ABL1 BCR |
| NPM1-RARA | | |
| KMT2A-CBL | | |
| IGH-BCL6 | HDAC1 TP53 CREBBP | HDAC1 CHD3 HDAC2 TP53 CREBBP CTBP1 EP300 SMARCA4 NCOR2 NCOR1 |
| LCP1-BCL6 | HDAC1 TP53 CREBBP | HDAC1 CHD3 HDAC2 TP53 CREBBP CTBP1 EP300 SMARCA4 NCOR2 NCOR1 |
| CREBBP-KAT6A | KMT2A HDAC2 CREBBP | HDAC1 SMARCC2 KMT2A HDAC2 WDR5 HECW2 CBX4 DOT1L SENP3 BMI1 CARM1 CTNNB1 SMARCC1 POLR2A CCNT1 SMARCA2 MAP3K5 CSNK2A2 KAT8 TAF6 CHD3 E2F4 TP53 TAF1 RAN TBP RBBP7 TOP1 RBBP4 KAT6A CXXC1 RUNX1 MBD3 NSD1 CTBP1 RELA MYB RNF2 CREBBP SIN3A |
| KMT2A-ARHGAP26 | KMT2A HDAC2 CREBBP | HDAC1 SMARCC2 KMT2A SMARCC1 WDR5 HECW2 CBX4 DOT1L SENP3 BMI1 CTNNB1 HDAC2 POLR2A CCNT1 SMARCA2 MAP3K5 CSNK2A2 KAT8 TAF6 CHD3 E2F4 TP53 TAF1 RAN TBP RBBP7 TOP1 RBBP4 CXXC1 RUNX1 MBD3 CTBP1 KAT6A MYB RNF2 CREBBP SIN3A |
| FOXO3-KMT2A | KMT2A CREBBP | HDAC1 SMARCC2 KMT2A HDAC2 SMAD1 BRCA1 SMAD3 AKT1 SMARCC1 POLR2A SMARCA2 EP300 MDM2 TAF6 CHD3 TP53 TAF1 RAN TBP MAP3K5 RUNX1 PCNA MBD3 VDR CTBP1 KAT6A MYB CREBBP SIN3A |
| KMT2A-DCPS | KMT2A HDAC2 CREBBP | HDAC1 SMARCC2 KMT2A HDAC2 SMARCC1 POLR2A SMARCA2 TAF6 CHD3 TP53 TAF1 RAN TBP RUNX1 MBD3 CTBP1 KAT6A MYB CREBBP SIN3A |

| | | |
|---|---|---|
| KMT2A-EP300 | | |
| IGH-CEBPE | DDIT3 ATF4 MYB | STAT6 E2F1 CEBPE FOS JUN RB1 |
| HSP90AA1-BCL6 | | |

**Table S15: SC_Preferential_attachment (TRAINING)**

| | | |
|---|---|---|
| | Essential Community Vertices | Preferential attachment vertices |
| ASPSCR1-TFE3 | | |
| ASTN2-CNOT2 | CNOT2 CNOT1 CNOT7 | HDAC3 GPS2 NCOR1 NCOR2 |
| BCOR-ZC3H7B | HDAC3 SP1 NCOR2 | HDAC1 HDAC3 HDAC4 CTBP1 NCOR2 |
| BCOR-CCNB3 | HDAC3 SP1 NCOR2 | HDAC1 HDAC3 HDAC4 CTBP1 NCOR2 |
| CDX1-IRF2BP2 | NTRK1 IRF2BP2 | IRF2 ELAVL1 NTRK1 IRF2BPL IRF2BP2 DEK EIF4H FOSL2 RBM39 SH3KBP1 VGLL4 |
| CIC-DUX4 | | |
| CREB1-EWSR1 | BRCA1 EWSR1 CREBBP | BRCA1 EPAS1 NR3C1 HDAC2 TP53 MYOD1 ESR1 HBP1 AKT1 POLR2A NCOR1 EP300 SMARCA4 CREBBP RPS6KA5 |
| CTDSP2-FAM19A2 | INTS6 CTDSP1 CTDSP2 | BRCA1 EPAS1 MYOD1 ESR1 JUN POLR2A EP300 SMARCA4 EWSR1 CREBBP |
| CXorf67-MBTD1 | | |
| EPC1-PHF1 | HDAC1 TP53 | NEK6 TRIM27 KAT5 TRIM23 |
| ERG-EWSR1 | TP53 ESR1 CREBBP | EPAS1 RAD23A ESR1 PARP1 PRKDC AR POLR2A RPA2 EP300 CREBBP |
| ETV6-NTRK3 | NTRK1 GRB2 | SQSTM1 TNK2 NTRK1 NTRK3 |
| EWSR1-ATF1 | GAB2 NTRK1 GRB2 | SQSTM1 TNK2 NTRK1 NTRK3 |
| EWSR1-FLI1 | BRCA1 ESR1 EP300 CREBBP | BRCA1 EPAS1 TP53 ESR1 KAT2B HBP1 POLR2A EP300 CREBBP |
| EWSR1-NR4A3 | FUS ELK1 EP300 TP53 CREBBP | EPAS1 POLR2A EP300 CREBBP |
| EWSR1-ETV4 | FUS ELK1 EP300 | EPAS1 TP53 ESR1 POLR2A EP300 CREBBP HBP1 |
| EWSR1-PATZ1 | FUS ELK1 EP300 CREBBP | EPAS1 TP53 ESR1 POLR2A EP300 CREBBP HBP1 |
| EWSR1-DDIT3 | HDAC1 HDAC3 DDIT3 TP53 EWSR1 | EPAS1 TP53 ESR1 POLR2A EP300 CREBBP HBP1 |
| EWSR1-POU5F1 | HDAC3 TP53 ESR1 EWSR1 | EPAS1 POLR2A EP300 CREBBP |
| EWSR1-SP3 | HDAC1 HDAC3 CEBPB ESR1 TP53 EWSR1 | EPAS1 TP53 ESR1 POLR2A RELA EP300 CREBBP HBP1 |
| FOXO4-CIC | ESR1 CREBBP | AKT1 ESR1 MDM2 CREBBP |
| FUS-ERG | DHX15 CUL3 | PRPF8 DHX15 CUL3 RPA2 RPA1 SF3B2 |
| FUS-CREB3L1 | CUL1 CUL2 CUL3 | NONO FASN CUL2 CUL3 |
| FUS-DDIT3 | ESR1 EWSR1 CREBBP | HDAC1 EPAS1 DDX17 PCNA ESR1 TP73 EWSR1 DDX5 CDK2 MDM2 RELA EP300 TRIP4 CREBBP ATF2 |

| | | |
|---|---|---|
| FUS-ATF1 | ESR1 EWSR1 EP300 CREBBP | HDAC1 EPAS1 DDX17 PCNA ESR1 TP73 EWSR1 DDX5 CDK2 MDM2 RELA EP300 TRIP4 CREBBP ATF2 |
| FUS-CREB3L2 | NONO CUL1 CUL2 CUL3 | NONO FASN CUL2 CUL3 |
| HEY1-NCOA2 | BRCA1 RARA TP53 ESR1 CREBBP | VDR ELAVL1 NR2F1 CARM1 SMAD3 ATR CCNT1 NKX2-1 ANKRD11 RARA STAT6 HIST4H4 NCOA6 MYOD1 NR1I2 FOS THRB THRA NR1I3 PGR PML CAD MEF2C BRCA1 NCOA2 NR3C1 ESRRB NCOA3 WDHD1 PRMT1 RXRA PPARG PPARD NCOA1 PPARA ESR2 EP300 UBR5 ETV1 TP53 ARNT CCND3 ESR1 HNF4A AHR PIAS3 AR KANK2 CREBBP |
| IRX2-TERT | TERT | YWHAZ TP53 RUVBL2 STUB1 YWHAQ RPS6KB1 POT1 AKT1 TERF1 MTOR UBR5 TERT ENO1 |
| JAZF1-SUZ12 | PRMT1 EED DDX5 | DHX9 RBM5 NXF1 PRPF8 |
| JAZF1-PHF1 | PPARG PHF1 | DHX9 PPARG PHF1 XRCC6 |
| LMNA-NTRK1 | | |
| MEAF6-TRERF1 | HDAC1 TRERF1 CREBBP | HDAC1 ELAVL1 TRERF1 KAT5 MORF4L1 NR5A1 ING5 KAT6A EP300 CREBBP |
| MEAF6-PHF1 | DHX9 PHF1 | DHX9 EZH2 XRCC6 PHF1 |
| NR4A3-TAF15 | TRIM28 FUS PRMT1 | TRIM28 FUS SAFB RPA1 RPA2 CUL4A CUL4B POLR2A TAF15 CUL5 CUL1 CUL2 CUL3 |
| NR4A3-TFG | CUL1 CUL2 CUL3 | TRIM28 SEC24A FBXO11 CUL1 CUL2 CUL3 |
| NR6A1-TRHDE | | NSD1 NCOA1 NR6A1 |
| NUP107-LGR5 | NUP153 NTRK1 | NUP153 VCP KPNB1 CUL3 NTRK1 C1QBP EIF4B |
| PAPPA-NUP107 | NTRK1 SMAD3 | NUP153 VCP KPNB1 ELAVL1 CUL3 NTRK1 C1QBP EIF4B |
| PAX3-FOXO1 | NCOA1 ESR1 CREBBP | HDAC1 IRF3 TRIM28 POU3F2 SMAD4 PARP1 SMAD3 AKT1 FOXO1 AR MDM2 EP300 RARA CEBPB NCOA1 ESR1 TBP YWHAQ HNF4A FHL2 PML CREBBP |
| PAX7-FOXO1 | RARA NCOA1 CREBBP | IRF3 WDR5 ASH2L SMAD4 TRIM27 PARP1 SMAD3 AKT1 FOXO1 AR MDM2 EP300 RARA CEBPB NCOA1 SKP2 MYOD1 ESR1 YWHAQ HNF4A FHL2 PML NLK YWHAZ YWHAG CREBBP |
| SS18-SSX2 | DPF2 SMARCC2 SMARCC1 PHF10 ELAVL1 DPF3 ARID2 DPF1 SMARCD3 SMARCE1 SMARCD1 EED SMARCA2 EP300 SMARCA4 HDAC1 ARID1B ARID1A ACTL6A HDAC2 CUL3 RNF2 SMARCD2 | SMARCC2 SMARCC1 SMARCA2 SMARCA4 |
| SS18-SSX1 | SMARCC2 HDAC2 EP300 | SMARCC2 SMARCC1 SMARCA2 SMARCA4 |
| SS18L1-SSX1 | SMARCC1 STAT3 SMAD3 SMAD1 CREBBP | DPF2 DPF3 HDAC2 STAT3 CUL3 BMI1 WHSC1L1 SMAD1 SMAD3 SMARCC1 ATF3 SMARCD1 EP300 SMARCA4 CREBBP SMARCE1 |
| SSX1-SYT4 | | |
| TGFBR3-MGEA5 | MAST1 | CBX8 CSNK2B NTRK1 PAXIP1 |
| TRIO-TERT | YWHAZ AKT1 TERT | YWHAZ TP53 RUVBL2 STUB1 YWHAQ RPS6KB1 POT1 AKT1 TERF1 MTOR UBR5 TERT ENO1 |
| WDR70-RCOR1 | RCOR1 | HDAC1 SMARCC2 HDAC3 NR2C1 SMARCE1 HDAC2 SMARCA4 RCOR1 |

| | | |
|---|---|---|
| YWHAE-NUTM2B | YWHAQ YWHAG HUWE1 NTRK1 YWHAE | ARAF BRAF RAF1 CDC37 |
| YWHAE-NUTM2A-AS1 | YWHAQ YWHAG HUWE1 NTRK1 YWHAE | ARAF BRAF RAF1 CDC37 |
| YWHAE-NUTM2A | YWHAQ YWHAG HUWE1 NTRK1 YWHAE | ARAF BRAF RAF1 CDC37 |

**Table S16: CA_Preferential_attachment (TRAINING)**

| | Essential Community Vertices | Preferential Attachment Vertices |
|---|---|---|
| ANK3-USP9Y | ANK3 SMAD2 SMAD3 | SMAD2 SMAD3 |
| ARGLU1-CXCR4 | APP CHERP SNRNP70 SRPK1 SRPK2 | PTK2 JAK2 JAK3 SOCS3 PTPN11 STAM |
| | PTK2 JAK2 SOCS3 PTPN11 NTRK1 | STAM ITCH |
| | PTK2 ELAVL1 SRPK1 NTRK1 | |
| ATXN10-FBLN1 | FN1 ATXN10 EGFR GSTK1 | VCP ABCE1 ATXN10 APP YWHAQ CUL3 |
| | ATXN10 VCP BSG CUL3 | FBLN1 NID1 FGB |
| | SMAD4 SKIL SMAD3 | |
| BCAS3-NFS1 | CTBP1 CTBP2 BCAS3 KAT2B | CTBP1 CTBP2 CDC23 KAT2B BCAS3 |
| | | CTBP1 HDAC5 ESR1 |
| | | PELP1 BCAS3 ESR1 |
| | | NFS1 SUCLA2 ACADM |
| | | PSMC2 SUCLA2 |
| BCAS4-BCAS3 | CTBP1 CTBP2 BCAS3 KAT2B | CTBP1 CTBP2 CDC23 KAT2B BCAS3 |
| | | PELP1 BCAS3 ESR1 |
| BCL2L12-PRMT1 | NCOA2 NCOA3 NCOA1 TP53 ESR1 BRCA1 PRMT1 THRB CARM1 NR1I2 AR PPARA EP300 | NCOA2 NCOA3 NCOA1 EP300 PARP1 |
| | EP300 STAT5A NCOA1 | NXF1 PARP1 |
| CAPNS1-WDR62 | YWHAZ FBXW11 FN1 HUWE1 YWHAQ GAPDH FERMT2 YWHAH VCAM1 YWHAB PAFAH1B1 YWHAG PAK2 YWHAE | YWHAZ FBXW11 FN1 HUWE1 YWHAQ GAPDH FERMT2 YWHAH VCAM1 YWHAB PAFAH1B1 YWHAG PAK2 YWHAE |
| | OGFOD1 MYO1E ASNS TBCB CAPN2 PROSC YWHAE | OGFOD1 MYO1E ASNS TBCB CAPN2 PROSC YWHAE |
| CCDC6-ANK3 | HDAC1 NR3C1 TRIM28 SF3A1 HNRNPR BRCC3 | HDAC1 NR3C1 TRIM28 ELAVL1 SKP1 HNRNPR PPP1CA BRCC3 NTRK1 SF3A1 CUL1 FBXW7 |
| | ANK3 SMAD2 SMAD3 | CUL1 SMAD3 |
| CCDC9-DHX34 | | EIF4A3 SNIP1 CCDC9 PRPF40A |
| | | TERF1 CCDC9 POT1 |
| | | DHX34 GSK3B |
| CDC27-ST7L | CDC16 CDC27 MDC1 CDC20 CREBBP ANAPC2 ANAPC7 | CDC16 CDC27 MDC1 SMAD2 TP53BP1 CREBBP UBE2S ANAPC2 ANAPC7 |

| | | |
|---|---|---|
| | ANAPC2 SMAD2 CREBBP | CREBBP E2F1 RB1 TFDP1 |
| | | TRIM33 ANAPC2 BUB1B |
| CDK7-RIN3 | BRCA1 TP53 RUVBL2 SUPT5H ESR1 RPA1 HNRNPU RPA2 CDK2 POLR2A | BRCA1 POLR2A RUVBL2 GTF2H1 RPA1 RPA2 |
| | GTF2H1 RPA1 RPA2 POLR2A CCNH | PRKCI APP CDC37 |
| | HDAC2 TP53 ESR1 MTA1 | |
| | GTF2H1 ERCC3 POLR2A ERCC5 | |
| | PRKCI APP CDK7 CDC37 | |
| CHERP-CPAMD8 | DHX8 U2AF1 RPA1 PRPF40A RPA2 CHERP SF3A2 RBM39 EWSR1 | DHX8 AGGF1 RNPS1 SNIP1 SF3B4 NTRK1 CHERP SRPK1 SRPK2 U2AF1 U2AF2 RPA1 APBB1 PRPF40A RPA2 TTC14 RBM23 SF3A2 SNRNP70 RBM39 EWSR1 WBP4 |
| | AGGF1 CHERP SF3A2 | CHERP HNRNPH3 |
| CPD-PIGW | | ELAVL1 NTRK1 |
| CYTH1-PRPSAP1 | DDX17 DDX5 ILK COPS5 | DDX5 FBXW11 DDX17 ILK ITGB2 COPS5 |
| | CYTH1 ARRB2 ARF6 ARRB1 | ARRB1 ARRB2 YWHAE |
| | MAPK1 ARRB2 ARRB1 | |
| DLG1-CRYBG3 | DLG1 LIN7A LIN7C APBA1 CASK | DDX17 DDX5 ILK COPS5 |
| | DLG1 NTRK1 CASK EPB41 | CYTH1 ARRB2 ARF6 ARRB1 |
| | DLG1 KHDRBS1 LCK NTRK1 | MAPK1 ARRB2 ARRB1 |
| | DLG1 UBE3A LCK | |
| DTX4-CCDC102B | | MCM7 CDK18 LENG1 TRIM54 TRIM27 KIFC3 |
| | | SFN MARK1 CCDC102B |
| | | NXT2 TRIM54 LNX1 |
| | | TRIM54 EHHADH |
| EHD4-FSIP1 | EHD4 CTPS2 EHD1 EGFR NTRK1 WARS PLCG1 UBA2 ADSL UQCRC2 | PLCG1 EHD4 EGFR NTRK1 |
| | EHD4 MTMR2 CTPS2 | TGFBR2 APP |
| ELK4-SLC26A9 | BRCA1 MAPK3 MAPK1 ELK4 | BRCA1 MAPK3 MAPK1 ELK4 BLM |
| EMID1-CBY1 | xx | xx |
| EPHA6-CNTN6 | xx | xx |
| ERAL1-DIDO1 | HNRNPDL RPA1 RPA2 HNRNPK RBM15 CUL3 DIDO1 FUS | FUS APP RPA1 RPA2 RBM15 CUL3 DIDO1 SRPK2 |
| | APP SRPK2 CUL3 | DIDO1 WWP1 |
| GMDS-CCND3 | PCNA PPP1CC RBL2 PPP1CA CCND3 RB1 POLD1 CDK2 CDK4 CDK6 CREBBP | NCOA2 RARA VDR CCND3 CREBBP |
| | GMDS NSFL1C CTH CAPN2 ATIC | CDK2 CREBBP |
| | RARA NCOA2 VDR CCND3 CREBBP | |
| | MCM10 RBX1 CCND3 APP | |

| | | |
|---|---|---|
| | BAG3 GMDS MAT2B | |
| HJURP-EIF4E2 | FBXW11 TP53 GIGYF2 APP HUWE1 EIF4E2 YWHAB SHMT2 YWHAE | FBXW11 TP53 YWHAB APP HUWE1 SHMT2 YWHAE |
| | RUVBL1 NXF1 RUVBL2 | RUVBL1 NXF1 RUVBL2 |
| INTS4-GAB2 | SRC PLCG1 GRB2 ZAP70 NTRK1 SHC1 | SRC PLCG1 GRB2 ZAP70 NTRK1 SHC1 |
| | PIK3CB PIK3R2 PIK3R1 | SRC PIK3R1 |
| KCNQ5-RIMS1 | | YWHAH RIMS1 |
| KDM5A-ANO2 | HDAC1 HDAC2 RBL1 TBP RB1 VDR KDM5A MORF4L1 | HDAC1 VDR RBL1 HDAC2 TBP KDM5A RB1 |
| | HDAC2 EZH2 KDM5A ESR1 | KDM5A ESR1 |
| | | NR3C1 KDM5A |
| LAMA5-C12orf28 | | |
| MAPK10-FAM13A | HDAC1 TP53 HDAC9 JUN DDX5 ELK1 MAPK10 RELA CREBBP ATF2 | HDAC1 TP53 APP JUN DDX5 MAPK10 RELA CREBBP |
| | APP MAPK10 MAP2K4 | CREBBP ATF2 |
| MAPRE1-TM9SF4 | YWHAZ FN1 APP TUBB NTRK1 VCAM1 UNK COPS5 | YWHAZ FN1 APP TUBB NTRK1 CDK2 OLA1 VCAM1 UNK COPS5 |
| | CDK5RAP2 PRKACA AKAP9 PRKACB | PDE4DIP PRKACA PRKACB CDK5RAP2 AKAP9 |
| | CLIP1 TUBB TUBA1A HDAC6 | TERF1 SPTAN1 DST MAPRE1 |
| | | CLIP1 HDAC6 TUBA1A TUBB |
| | | ABCE1 MACF1 |
| | | BAG3 TERF1 |
| NCKAP5-MZT2A | | |
| NUMB-ALDH6A1 | TP53 NUMB ITCH MDM2 EGFR | PRKCZ NUMB APP EGFR |
| | EPS15 EGFR AP2A1 NUMB | EGFR ITCH |
| | TPI1 EGFR APP | |
| PARD6B-CD48 | PRKCI RASSF8 PARD3 PARD6G APP PARD6B PARD6A YWHAH PRKCZ WWC1 | PRKCI PARD3 PARD6G APP PARD6B PARD6A YWHAH PRKCZ |
| | RAC1 PARD6G PARD6B PARD6A | EEF1D APP |
| PPP1R12A-MGAT4C | KDM1A ELAVL1 RPA1 RPA2 PPP1R12A CUL1 | KDM1A NUDT5 TP53 ELAVL1 PUS1 NUAK1 AARSD1 RPA1 NTRK1 RPA2 RPRD1B PPP1R12A TRIM47 PAXIP1 ACTR3 CUL1 |
| | KDM1A TP53 RPA1 | PPP1CB PPP1R12A HDAC7 |
| RNF11-C8A | CBLB RNF11 ITCH SMAD4 EPN1 RABGEF1 UBE2E1 UBE2D3 UBE2E3 HGS GGA1 AKT1 GGA3 GGA2 AP2A1 EPN3 UBE2D1 AP2B1 CSNK2A1 SMURF1 EPS15 SMURF2 UBQLN2 STAM2 NEDD4 UBQLN4 NEDD4L APP | GGA1 GGA3 GGA2 APP |

| | | |
|---|---|---|
| | RNF11 PSMD4 PSMD7 PSMD6 PSMD11 PSMD10 PSMD3 PSMD12 PSMD13 USP14 PSMD14 PSMD1 PSMD2 APP | UBQLN2 UBQLN4 |
| | AKT1 RNF11 TBK1 APP NEDD4 | |
| SIPA1L3-WDR62 | MAPK10 WDR62 MAPK8 MAPK9 | YWHAB FBXW11 MAPK10 ELAVL1 WDR62 TBP MAPK8 MAPK9 |
| | YWHAB FBXW11 ELAVL1 | SFN YWHAB SIPA1L3 YWHAQ |
| | WDR62 YWHAB FBXW11 | LATS2 AURKA |
| SLC26A6-PRKAR2A | AKAP7 AKAP9 PRKAR2A PRKAR2B PRKACA PRKACB | PRKAR2A AKAP7 PRKACA PRKACB PRKAR2B GCH1 |
| | PRKAR2A PRKAR2B GCH1 | PRKAR2A RUNX1T1 CBFA2T3 |
| | | VCP FAF2 |
| ST14-APLP2 | BRCA1 APLP2 ETS1 JUN | BRCA1 JUNB JUN APBB1 APBB2 KAT5 ETS1 APLP2 MAPK8 |
| | APLP2 APBB1 APBB2 | SFN HDAC5 APLP2 RPL26 |
| | SFN APLP2 HDAC5 | APLP2 DEDD |
| | APLP2 ETS1 ELAVL1 | APLP2 VKORC1 |
| | BRCA1 APLP2 KAT5 | |
| STRADB-NOP58 | NIFK NOP56 HNRNPU RUVBL2 NOLC1 SNU13 NTRK1 PUM3 RPS15A RSL1D1 KRR1 DDX18 EIF6 DHX15 RPS4X EED PRPF3 TARDBP RPL11 EIF2S2 DDX27 NOP58 FN1 DDX24 U2AF1 RPL30 ESR1 DDX56 FTSJ3 DDX47 KPNA6 GTPBP4 WDR36 KPNA1 DKC1 RRP12 BOP1 FBL TBL3 | WDR36 NOP58 DHX15 NOP56 |
| | XIAP MAP3K7 TRAF6 | PRPF3 NOP58 |
| STX16-RAE1 | FBXW11 NXF1 FAF1 RAE1 ILF3 CUL1 HNRNPUL1 CUL3 | NXF1 CUL1 ILF3 CUL3 |
| | USP11 CUL1 NXF1 | BUB1 NXF1 |
| TANC2-CHD6 | SOX2 CHD3 CHD6 | ZFYVE9 PPP1CC PPP1CA TANC2 |
| | | SOX2 CHD3 CHD6 |
| | | AIRE CHD6 |
| THSD7B-DARS | SOX2 CHD3 CHD6 | ZFYVE9 PPP1CC PPP1CA TANC2 |
| | | SOX2 CHD3 CHD6 |
| | | AIRE CHD6 |
| TMEM123-MMP7 | MAEA RANBP9 MKLN1 MMP7 RMND5A | MAEA RANBP9 MKLN1 RANBP10 MMP7 RMND5A |
| TMPRSS2-ERG | CDC5L DDX3X ELAVL1 CAD NEDD4 SF3B1 PARP1 PRKDC PRPF8 SFPQ ERG POLR2A TOP1 CLTC SF3B2 XRCC5 XRCC6 DDX23 SNRNP200 DDX21 TUBB NONO JUN HNRNPU PRPF40A AR NCL HNRNPM HNRNPC TOP2B ILF3 ILF2 | PRPF8 ERG SF3B2 SF3B1 PARP1 PRKDC |
| | ACTB ERG CAD | PRPF8 NEDD4 |
| TOX3-CNTN5 | | |

| |
|---|
| UBR2-XPO5 |
| UVRAG-INTS4 |
| WNT11-TSPAN8 |
| XRCC5-ACADL |
| ZCCHC7-PRSS3 |
| ZFP91-NOX4 |

**Table S17: LK, LY, ME, GL-PAS (TRAINING)**

| | Essential Community Vertices | D | CC | BC | Avg.D (avgD) | Net.Diam (ND) | PAS | PAS/D | D/PAS |
|---|---|---|---|---|---|---|---|---|---|
| RUNX1-RUNX1T1 | HDAC1 | 478 | 0 | 212982 | 1.996 | 2 | 119.739 | 0.2505 | 3.992 |
| | BRCA1 | 17 | 0.191 | 187.233 | 0.09 | 2 | 94.4444 | 5.55556 | 0.18 |
| | KMT2A | 394 | 0.109 | 5267.838 | 21.87 | 2 | 9.00695 | 0.02286 | 43.744 |
| | SMARCC1 | 116 | 0.148 | 4333.831 | 0.559 | 2 | 103.757 | 0.89445 | 1.118 |
| | HDAC2 | 301 | 0.067 | 40883.166 | 1.874 | 2 | 80.3095 | 0.26681 | 3.748 |
| | CREBBP | 296 | 0.058 | 48179.347 | 1.061 | 2 | 139.491 | 0.47125 | 2.122 |
| | CTBP1 | 148 | 0.058 | 14901.592 | 0.751 | 2 | 98.5353 | 0.66578 | 1.502 |
| | EP300 | 453 | 0 | 203852 | 1.996 | 2 | 113.477 | 0.2505 | 3.992 |
| | SMARCA4 | 200 | 0.088 | 20379.264 | 3.512 | 2 | 28.4738 | 0.14237 | 7.024 |
| | NCOR2 | 104 | 0.136 | 4824.426 | 15.81 | 2 | 3.28906 | 0.03163 | 31.62 |
| | NCOR1 | 151 | 0.095 | 11956.904 | 16.08 | 2 | 4.69557 | 0.0311 | 32.158 |
| | VDR | 89 | 0.109 | 4933.604 | 0.371 | 2 | 119.946 | 1.34771 | 0.742 |
| | SMARCA4 | 200 | 0.088 | 20379.264 | 3.512 | 2 | 28.4738 | 0.14237 | 7.024 |
| KMT2A-MLLT10 | SMARCC2 | 129 | 0.129 | 6769.42 | 0.623 | 2 | 103.531 | 0.80257 | 1.246 |
| | KMT2A | 394 | 0.109 | 5267.838 | 21.87 | 2 | 9.00695 | 0.02286 | 43.744 |
| | HDAC2 | 301 | 0.067 | 40883.166 | 1.874 | 2 | 80.3095 | 0.26681 | 3.748 |
| | CHD3 | 146 | 0.075 | 13797.187 | 12.66 | 2 | 5.7671 | 0.0395 | 25.316 |
| | SMARCC1 | 116 | 0.148 | 4333.831 | 0.559 | 2 | 103.757 | 0.89445 | 1.118 |
| | POLR2A | 251 | 0.074 | 34918.732 | 1.239 | 2 | 101.291 | 0.40355 | 2.478 |
| | SMARCA2 | 103 | 0.141 | 4685.809 | 16.25 | 2 | 3.16923 | 0.03077 | 32.5 |
| | CREBBP | 296 | 0.058 | 48179.347 | 1.061 | 2 | 139.491 | 0.47125 | 2.122 |
| | SIN3A | 192 | 0.072 | 20856.124 | 15.68 | 2 | 6.12284 | 0.03189 | 31.358 |
| | CTBP1 | 148 | 0.058 | 14901.592 | 0.751 | 2 | 98.5353 | 0.66578 | 1.502 |
| | KMT2A | 394 | 0.109 | 5267.838 | 21.87 | 2 | 9.00695 | 0.02286 | 43.744 |
| | CREBBP | 296 | 0.058 | 48179.347 | 1.061 | 2 | 139.491 | 0.47125 | 2.122 |
| IGH-BCL2 | TP53 | 961 | 0 | 920640 | 1.998 | 2 | 240.49 | 0.25025 | 3.996 |
| | CASP3 | 93 | 0.077 | 5925.317 | 0.503 | 2 | 92.4453 | 0.99404 | 1.006 |
| | PARP1 | 219 | 0.08 | 25769.813 | 1.233 | 2 | 88.8078 | 0.40552 | 2.466 |
| | CASP8 | 134 | 0.057 | 13584.185 | 0.388 | 2 | 172.68 | 1.28866 | 0.776 |

| | | | | | | | | | |
|---|---|---|---|---|---|---|---|---|---|
| | HIF1A | 145 | 0.089 | 12651.423 | 0.74 | 2 | 97.973 | 0.67568 | 1.48 |
| | BCL2 | 94 | 0.069 | 6004.406 | 8.234 | 2 | 5.70804 | 0.06072 | 16.468 |
| | BAG3 | 448 | 0 | 199362 | 30 | 2 | 7.46766 | 0.01667 | 59.992 |
| | PARP1 | 219 | 0.08 | 25769.813 | 1.233 | 2 | 88.8078 | 0.40552 | 2.466 |
| | BCL2 | 94 | 0.069 | 6004.406 | 8.234 | 2 | 5.70804 | 0.06072 | 16.468 |
| KMT2A-AFF1 | SMARCC2 | 129 | 0.129 | 6769.42 | 0.623 | 2 | 103.531 | 0.80257 | 1.246 |
| | KMT2A | 394 | 0.109 | 5267.838 | 21.87 | 2 | 9.00695 | 0.02286 | 43.744 |
| | HDAC2 | 301 | 0.067 | 40883.166 | 1.874 | 2 | 80.3095 | 0.26681 | 3.748 |
| | CHD3 | 146 | 0.075 | 13797.187 | 12.66 | 2 | 5.7671 | 0.0395 | 25.316 |
| | SMARCC1 | 116 | 0.148 | 4333.831 | 0.559 | 2 | 103.757 | 0.89445 | 1.118 |
| | POLR2A | 251 | 0.074 | 34918.732 | 1.239 | 2 | 101.291 | 0.40355 | 2.478 |
| | SMARCA2 | 103 | 0.141 | 4685.809 | 16.25 | 2 | 3.16923 | 0.03077 | 32.5 |
| | CREBBP | 296 | 0.058 | 48179.347 | 1.061 | 2 | 139.491 | 0.47125 | 2.122 |
| | SIN3A | 192 | 0.072 | 20856.124 | 15.68 | 2 | 6.12284 | 0.03189 | 31.358 |
| | CTBP1 | 148 | 0.058 | 14901.592 | 0.751 | 2 | 98.5353 | 0.66578 | 1.502 |
| | KMT2A | 394 | 0.109 | 5267.838 | 21.87 | 2 | 9.00695 | 0.02286 | 43.744 |
| | CREBBP | 296 | 0.058 | 48179.347 | 1.061 | 2 | 139.491 | 0.47125 | 2.122 |
| PICALM-MLLT10 | FN1 | 23 | 0.134 | 343.967 | 0.109 | 2 | 105.505 | 4.58716 | 0.218 |
| | EEF1A1 | 370 | 0.082 | 71411.66 | 1.167 | 2 | 158.526 | 0.42845 | 2.334 |
| | EGFR | 833 | 0 | 691392 | 52 | 2 | 8.00992 | 0.00962 | 103.996 |
| | NTRK1 | 166 | 0.045 | 22876.817 | 9.512 | 2 | 8.72582 | 0.05257 | 19.024 |
| | PLCG1 | 112 | 0.071 | 8175.982 | 9.805 | 2 | 5.71137 | 0.05099 | 19.61 |
| | DNM2 | 88 | 0.062 | 6030.786 | 7.348 | 2 | 5.98802 | 0.06805 | 14.696 |
| | ILVBL | 41 | 0.133 | 1028.658 | 7.143 | 2 | 2.86994 | 0.07 | 14.286 |
| | PICALM | 89 | 0.147 | 4243.12 | 14.76 | 2 | 3.01572 | 0.03388 | 29.512 |
| | HNRNPD | 269 | 0.076 | 42696.344 | 22.19 | 2 | 6.06047 | 0.02253 | 44.386 |
| | FUS | 319 | 0.082 | 50309.802 | 8 | 2 | 19.9375 | 0.0625 | 16 |
| | FN1 | 23 | 0.134 | 343.967 | 0.109 | 2 | 105.505 | 4.58716 | 0.218 |
| | DDX1 | 127 | 0.168 | 5869.092 | 0.706 | 2 | 89.9433 | 0.70822 | 1.412 |
| | SEC24D | 26 | 0.142 | 450.267 | 5.333 | 2 | 2.43765 | 0.09376 | 10.666 |
| | PICALM | 89 | 0.147 | 4243.12 | 14.76 | 2 | 3.01572 | 0.03388 | 29.512 |
| | NTRK1 | 166 | 0.045 | 22876.817 | 9.512 | 2 | 8.72582 | 0.05257 | 19.024 |
| | SEC24C | 64 | 0.125 | 2418.599 | 9.754 | 2 | 3.28071 | 0.05126 | 19.508 |
| PML-RARA | NCOA2 | 64 | 0.121 | 2191.888 | 9.446 | 2 | 3.38768 | 0.05293 | 18.892 |
| | NR3C1 | 157 | 0.069 | 15984.767 | 12.61 | 2 | 6.22472 | 0.03965 | 25.222 |
| | NR4A1 | 94 | 0.051 | 7257.643 | 0.253 | 2 | 185.771 | 1.97628 | 0.506 |
| | KAT2B | 175 | 0.066 | 18317.722 | 13.38 | 2 | 6.53815 | 0.03736 | 26.766 |
| | RXRA | 109 | 0.094 | 6829.11 | 11.91 | 2 | 4.57676 | 0.04199 | 23.816 |
| | PPARG | 130 | 0.086 | 10094.308 | 12.88 | 2 | 5.04776 | 0.03883 | 25.754 |
| | TP53 | 961 | 0 | 920640 | 1.998 | 2 | 240.49 | 0.25025 | 3.996 |
| | MDM2 | 189 | 0.06 | 22876.817 | 9.512 | 2 | 9.93482 | 0.05257 | 19.024 |
| | EP300 | 453 | 0 | 203852 | 1.996 | 2 | 113.477 | 0.2505 | 3.992 |
| | SMARCA4 | 200 | 0.088 | 20379.264 | 3.512 | 2 | 28.4738 | 0.14237 | 7.024 |
| | RELA | 178 | 0.06 | 22556.817 | 9.512 | 2 | 9.3566 | 0.05257 | 19.024 |
| | RARA | 108 | 0.084 | 7253.63 | 10.8 | 2 | 5.00185 | 0.04631 | 21.592 |
| | NCOA3 | 106 | 0.109 | 5991.77 | 13.36 | 2 | 3.96588 | 0.03741 | 26.728 |
| | NCOA1 | 101 | 0.12 | 5004.528 | 13.74 | 2 | 3.6746 | 0.03638 | 27.486 |

| | | | | | | | | | |
|---|---|---|---|---|---|---|---|---|---|
| | STAT3 | 223 | 0.044 | 37462.83 | 5.641 | 2 | 19.766 | 0.08864 | 11.282 |
| | ARNT | 42 | 0.07 | 1457.586 | 4.744 | 2 | 4.42664 | 0.1054 | 9.488 |
| | NPAS2 | 26 | 0.151 | 470.9 | 0.156 | 2 | 83.3333 | 3.20513 | 0.312 |
| | PARP1 | 219 | 0.08 | 25769.813 | 1.233 | 2 | 88.8078 | 0.40552 | 2.466 |
| | NFKB1 | 141 | 0.08 | 11734.095 | 3.021 | 2 | 23.3366 | 0.16551 | 6.042 |
| | TRIP4 | 41 | 0.248 | 760.708 | 11.62 | 2 | 1.76435 | 0.04303 | 23.238 |
| | CREBBP | 296 | 0.058 | 48179.347 | 1.061 | 2 | 139.491 | 0.47125 | 2.122 |
| | NFKB1 | 141 | 0.08 | 11734.095 | 3.021 | 2 | 23.3366 | 0.16551 | 6.042 |
| | EP300 | 453 | 0 | 203852 | 1.996 | 2 | 113.477 | 0.2505 | 3.992 |
| | PARP1 | 219 | 0.08 | 25769.813 | 1.233 | 2 | 88.8078 | 0.40552 | 2.466 |
| KMT2A-MLLT3 | SMARCC2 | 129 | 0.129 | 6769.42 | 0.623 | 2 | 103.531 | 0.80257 | 1.246 |
| | KMT2A | 394 | 0.109 | 5267.838 | 21.87 | 2 | 9.00695 | 0.02286 | 43.744 |
| | HDAC2 | 301 | 0.067 | 40883.166 | 1.874 | 2 | 80.3095 | 0.26681 | 3.748 |
| | CHD3 | 146 | 0.075 | 13797.187 | 12.66 | 2 | 5.7671 | 0.0395 | 25.316 |
| | SMARCC1 | 116 | 0.148 | 4333.831 | 0.559 | 2 | 103.757 | 0.89445 | 1.118 |
| | POLR2A | 251 | 0.074 | 34918.732 | 1.239 | 2 | 101.291 | 0.40355 | 2.478 |
| | SMARCA2 | 103 | 0.141 | 4685.809 | 16.25 | 2 | 3.16923 | 0.03077 | 32.5 |
| | CREBBP | 296 | 0.058 | 48179.347 | 1.061 | 2 | 139.491 | 0.47125 | 2.122 |
| | SIN3A | 192 | 0.072 | 20856.124 | 15.68 | 2 | 6.12284 | 0.03189 | 31.358 |
| | CTBP1 | 148 | 0.058 | 14901.592 | 0.751 | 2 | 98.5353 | 0.66578 | 1.502 |
| | KMT2A | 394 | 0.109 | 5267.838 | 21.87 | 2 | 9.00695 | 0.02286 | 43.744 |
| | CREBBP | 296 | 0.058 | 48179.347 | 1.061 | 2 | 139.491 | 0.47125 | 2.122 |
| KMT2A-AFDN | SMARCC2 | 129 | 0.129 | 6769.42 | 0.623 | 2 | 103.531 | 0.80257 | 1.246 |
| | KMT2A | 394 | 0.109 | 5267.838 | 21.87 | 2 | 9.00695 | 0.02286 | 43.744 |
| | HDAC2 | 301 | 0.067 | 40883.166 | 1.874 | 2 | 80.3095 | 0.26681 | 3.748 |
| | CHD3 | 146 | 0.075 | 13797.187 | 12.66 | 2 | 5.7671 | 0.0395 | 25.316 |
| | SMARCC1 | 116 | 0.148 | 4333.831 | 0.559 | 2 | 103.757 | 0.89445 | 1.118 |
| | POLR2A | 251 | 0.074 | 34918.732 | 1.239 | 2 | 101.291 | 0.40355 | 2.478 |
| | SMARCA2 | 103 | 0.141 | 4685.809 | 16.25 | 2 | 3.16923 | 0.03077 | 32.5 |
| | CREBBP | 296 | 0.058 | 48179.347 | 1.061 | 2 | 139.491 | 0.47125 | 2.122 |
| | SIN3A | 192 | 0.072 | 20856.124 | 15.68 | 2 | 6.12284 | 0.03189 | 31.358 |
| | CTBP1 | 148 | 0.058 | 14901.592 | 0.751 | 2 | 98.5353 | 0.66578 | 1.502 |
| | KMT2A | 394 | 0.109 | 5267.838 | 21.87 | 2 | 9.00695 | 0.02286 | 43.744 |
| | CREBBP | 296 | 0.058 | 48179.347 | 1.061 | 2 | 139.491 | 0.47125 | 2.122 |
| CBFB-MYH11 | ACTA2 | 82 | 0.092 | 4895.233 | 9.325 | 2 | 4.39678 | 0.05362 | 18.65 |
| | MYH11 | 44 | 0.071 | 1556.867 | 4.933 | 2 | 4.45976 | 0.10136 | 9.866 |
| | MYO1E | 55 | 0.174 | 1893.75 | 11.18 | 2 | 2.45997 | 0.04473 | 22.358 |
| | RPA1 | 463 | 0 | 212982 | 1.996 | 2 | 115.982 | 0.2505 | 3.992 |
| | RPA2 | 226 | 0.06 | 22556.817 | 9.512 | 2 | 11.8797 | 0.05257 | 19.024 |
| | ACTB | 223 | 0.076 | 28937.125 | 18.66 | 2 | 5.97407 | 0.02679 | 37.328 |
| | RPA1 | 463 | 0 | 212982 | 1.996 | 2 | 115.982 | 0.2505 | 3.992 |
| | ELAVL1 | 213 | 0.076 | 28937.125 | 18.66 | 2 | 5.70617 | 0.02679 | 37.328 |
| | RPA2 | 226 | 0.06 | 22556.817 | 9.512 | 2 | 11.8797 | 0.05257 | 19.024 |
| NUP98-DDX10 | SIRT7 | 172 | 0.045 | 22876.817 | 9.512 | 2 | 9.04121 | 0.05257 | 19.024 |
| | DDX10 | 127 | 0.168 | 5869.092 | 23.02 | 2 | 2.75895 | 0.02172 | 46.032 |

| | | | | | | | | | |
|---|---|---|---|---|---|---|---|---|---|
| | APP | 169 | 0.045 | 22876.817 | 13.51 | 2 | 6.2537 | 0.037 | 27.024 |
| | DDX56 | 82 | 0.238 | 2197.27 | 21.06 | 2 | 1.94682 | 0.02374 | 42.12 |
| | NTRK1 | 166 | 0.045 | 22876.817 | 9.512 | 2 | 8.72582 | 0.05257 | 19.024 |
| | DDX54 | 43 | 0.126 | 1231.913 | 7.136 | 2 | 3.01289 | 0.07007 | 14.272 |
| | PUM3 | 65 | 0.358 | 586.736 | 24.52 | 2 | 1.32572 | 0.0204 | 49.03 |
| | PWP1 | 36 | 0.298 | 471.093 | 12.11 | 2 | 1.48662 | 0.0413 | 24.216 |
| | CSNK2A1 | 382 | 0 | 144780 | 22 | 2 | 8.68379 | 0.02273 | 43.99 |
| | HDAC1 | 478 | 0 | 212982 | 1.996 | 2 | 119.739 | 0.2505 | 3.992 |
| | CTNNB1 | 286 | 0.057 | 51076.364 | 18.22 | 2 | 7.85024 | 0.02745 | 36.432 |
| | MAPK8 | 172 | 0.045 | 22876.817 | 9.512 | 2 | 9.04121 | 0.05257 | 19.024 |
| | EP300 | 453 | 0 | 203852 | 1.996 | 2 | 113.477 | 0.2505 | 3.992 |
| | CREBBP | 296 | 0.058 | 48179.347 | 1.061 | 2 | 139.491 | 0.47125 | 2.122 |
| | PUM3 | 65 | 0.358 | 586.736 | 24.52 | 2 | 1.32572 | 0.0204 | 49.03 |
| | NXF1 | 61 | 0.312 | 489.736 | 24.52 | 2 | 1.24414 | 0.0204 | 49.03 |
| | EED | 66 | 0.322 | 479.736 | 0.167 | 2 | 197.605 | 2.99401 | 0.334 |
| | KPNB1 | 211 | 0.072 | 26556.162 | 16.9 | 2 | 6.2426 | 0.02959 | 33.8 |
| | HNRNPUL1 | 112 | 0.14 | 6163.826 | 17.23 | 2 | 3.24977 | 0.02902 | 34.464 |
| | HDAC1 | 478 | 0 | 212982 | 1.996 | 2 | 119.739 | 0.2505 | 3.992 |
| | CREBBP | 296 | 0.058 | 48179.347 | 1.061 | 2 | 139.491 | 0.47125 | 2.122 |
| | APC | 168 | 0.03 | 24110.735 | 6.869 | 2 | 12.2289 | 0.07279 | 13.738 |
| | CSNK2A1 | 382 | 0 | 144780 | 22 | 2 | 8.68379 | 0.02273 | 43.99 |
| | CTNNB1 | 286 | 0.057 | 51076.364 | 18.22 | 2 | 7.85024 | 0.02745 | 36.432 |
| | APC | 168 | 0.03 | 24110.735 | 6.869 | 2 | 12.2289 | 0.07279 | 13.738 |
| | CREBBP | 296 | 0.058 | 48179.347 | 1.061 | 2 | 139.491 | 0.47125 | 2.122 |
| | USP7 | 41 | 0.248 | 760.708 | 11.62 | 2 | 1.76435 | 0.04303 | 23.238 |
| | NTRK1 | 166 | 0.045 | 22876.817 | 9.512 | 2 | 8.72582 | 0.05257 | 19.024 |
| | CDC37 | 44 | 0.083 | 1464.862 | 5.364 | 2 | 4.10142 | 0.09321 | 10.728 |
| PCM1-JAK2 | ERBB2 | 23 | 0.134 | 343.967 | 4.609 | 2 | 2.49512 | 0.10848 | 9.218 |
| | ERBB3 | 23 | 0.134 | 343.967 | 8.609 | 2 | 1.33581 | 0.05808 | 17.218 |
| | VAV1 | 41 | 0.248 | 760.708 | 9.619 | 2 | 2.1312 | 0.05198 | 19.238 |
| | TEC | 156 | 0.071 | 15458.811 | 11.86 | 2 | 6.57728 | 0.04216 | 23.718 |
| | EGFR | 833 | 0 | 691392 | 52 | 2 | 8.00992 | 0.00962 | 103.996 |
| | INSR | 41 | 0.133 | 1028.658 | 9.143 | 2 | 2.24215 | 0.05469 | 18.286 |
| | PLCG1 | 112 | 0.071 | 8175.982 | 9.805 | 2 | 5.71137 | 0.05099 | 19.61 |
| | JAK2 | 41 | 0.133 | 1028.658 | 14.14 | 2 | 1.44948 | 0.03535 | 28.286 |
| | STAT5A | 223 | 0.044 | 37462.83 | 6.641 | 2 | 16.7896 | 0.07529 | 13.282 |
| | JAK2 | 41 | 0.133 | 1028.658 | 14.14 | 2 | 1.44948 | 0.03535 | 28.286 |
| | INSR | 41 | 0.133 | 1028.658 | 9.143 | 2 | 2.24215 | 0.05469 | 18.286 |
| KMT2A-SEPT9 | SMARCC2 | 129 | 0.129 | 6769.42 | 0.623 | 2 | 103.531 | 0.80257 | 1.246 |
| | KMT2A | 394 | 0.109 | 5267.838 | 21.87 | 2 | 9.00695 | 0.02286 | 43.744 |
| | HDAC2 | 301 | 0.067 | 40883.166 | 1.874 | 2 | 80.3095 | 0.26681 | 3.748 |
| | CHD3 | 146 | 0.075 | 13797.187 | 12.66 | 2 | 5.7671 | 0.0395 | 25.316 |
| | SMARCC1 | 116 | 0.148 | 4333.831 | 0.559 | 2 | 103.757 | 0.89445 | 1.118 |
| | POLR2A | 251 | 0.074 | 34918.732 | 1.239 | 2 | 101.291 | 0.40355 | 2.478 |
| | SMARCA2 | 103 | 0.141 | 4685.809 | 16.25 | 2 | 3.16923 | 0.03077 | 32.5 |
| | CREBBP | 296 | 0.058 | 48179.347 | 1.061 | 2 | 139.491 | 0.47125 | 2.122 |
| | SIN3A | 192 | 0.072 | 20856.124 | 15.68 | 2 | 6.12284 | 0.03189 | 31.358 |
| | CTBP1 | 148 | 0.058 | 14901.592 | 0.751 | 2 | 98.5353 | 0.66578 | 1.502 |

| | | | | | | | | | |
|---|---|---|---|---|---|---|---|---|---|
| | KMT2A | 394 | 0.109 | 5267.838 | 21.87 | 2 | 9.00695 | 0.02286 | 43.744 |
| | CREBBP | 296 | 0.058 | 48179.347 | 1.061 | 2 | 139.491 | 0.47125 | 2.122 |
| FUS-ERG | RPA1 | 463 | 0 | 212982 | 1.996 | 2 | 115.982 | 0.2505 | 3.992 |
| | SF3B2 | 26 | 0.142 | 450.267 | 7.333 | 2 | 1.77281 | 0.06818 | 14.666 |
| | PRKDC | 50 | 0.358 | 586.736 | 24.52 | 2 | 1.01978 | 0.0204 | 49.03 |
| | PRPF8 | 90 | 0.071 | 8175.982 | 9.805 | 2 | 4.5895 | 0.05099 | 19.61 |
| | SF3A2 | 26 | 0.142 | 450.267 | 6.333 | 2 | 2.05274 | 0.07895 | 12.666 |
| | DHX15 | 82 | 0.238 | 2197.27 | 21.06 | 2 | 1.94682 | 0.02374 | 42.12 |
| | RPA2 | 226 | 0.06 | 22556.817 | 9.512 | 2 | 11.8797 | 0.05257 | 19.024 |
| | CUL3 | 70 | 0.081 | 3676.928 | 10.37 | 2 | 3.3748 | 0.04821 | 20.742 |
| | ABL1 | 173 | 0.076 | 17246.646 | 14.9 | 2 | 5.80459 | 0.03355 | 29.804 |
| | PARP1 | 219 | 0.08 | 25769.813 | 1.233 | 2 | 88.8078 | 0.40552 | 2.466 |
| | PRKDC | 50 | 0.358 | 586.736 | 24.52 | 2 | 1.01978 | 0.0204 | 49.03 |
| NUP98-HOXA9 | HDAC1 | 478 | 0 | 212982 | 1.996 | 2 | 119.739 | 0.2505 | 3.992 |
| | TP53 | 961 | 0 | 920640 | 1.998 | 2 | 240.49 | 0.25025 | 3.996 |
| | SMAD4 | 103 | 0.141 | 4685.809 | 2.01 | 2 | 25.6219 | 0.24876 | 4.02 |
| | CTNNB1 | 286 | 0.057 | 51076.364 | 18.22 | 2 | 7.85024 | 0.02745 | 36.432 |
| | CSNK2A1 | 382 | 0 | 144780 | 22 | 2 | 8.68379 | 0.02273 | 43.99 |
| | MAPK8 | 172 | 0.045 | 22876.817 | 9.512 | 2 | 9.04121 | 0.05257 | 19.024 |
| | EP300 | 453 | 0 | 203852 | 1.996 | 2 | 113.477 | 0.2505 | 3.992 |
| | CREBBP | 296 | 0.058 | 48179.347 | 1.061 | 2 | 139.491 | 0.47125 | 2.122 |
| | CTNNB1 | 286 | 0.057 | 51076.364 | 18.22 | 2 | 7.85024 | 0.02745 | 36.432 |
| | APC | 168 | 0.03 | 24110.735 | 6.869 | 2 | 12.2289 | 0.07279 | 13.738 |
| | CREBBP | 296 | 0.058 | 48179.347 | 1.061 | 2 | 139.491 | 0.47125 | 2.122 |
| ETV6-ABL1 | ERBB2 | 23 | 0.134 | 343.967 | 4.609 | 2 | 2.49512 | 0.10848 | 9.218 |
| | CBLB | 44 | 0.083 | 1464.862 | 6.364 | 2 | 3.45695 | 0.07857 | 12.728 |
| | UBASH3B | 109 | 0.097 | 6392.818 | 12.26 | 2 | 4.44644 | 0.04079 | 24.514 |
| | SOS1 | 44 | 0.174 | 955.71 | 9.289 | 2 | 2.36839 | 0.05383 | 18.578 |
| | ERBB4 | 23 | 0.134 | 343.967 | 0.609 | 2 | 18.8834 | 0.82102 | 1.218 |
| | SRC | 44 | 0.174 | 955.71 | 5.289 | 2 | 4.15958 | 0.09454 | 10.578 |
| | VAV1 | 41 | 0.248 | 760.708 | 9.619 | 2 | 2.1312 | 0.05198 | 19.238 |
| | EGFR | 833 | 0 | 691392 | 52 | 2 | 8.00992 | 0.00962 | 103.996 |
| | CBL | 233 | 0.061 | 32363.102 | 6.043 | 2 | 19.2785 | 0.08274 | 12.086 |
| | PLCG1 | 112 | 0.071 | 8175.982 | 9.805 | 2 | 5.71137 | 0.05099 | 19.61 |
| | PIK3R2 | 109 | 0.111 | 6174.794 | 13.89 | 2 | 3.9234 | 0.03599 | 27.782 |
| | PIK3R1 | 145 | 0.079 | 12813.777 | 13.15 | 2 | 5.51541 | 0.03804 | 26.29 |
| | ABL1 | 173 | 0.076 | 17246.646 | 14.9 | 2 | 5.80459 | 0.03355 | 29.804 |
| | ABL1 | 173 | 0.076 | 17246.646 | 14.9 | 2 | 5.80459 | 0.03355 | 29.804 |
| | ABL2 | 153 | 0.076 | 17246.646 | 14.9 | 2 | 5.13354 | 0.03355 | 29.804 |
| | JAK1 | 41 | 0.133 | 1028.658 | 14.14 | 2 | 1.44948 | 0.03535 | 28.286 |
| SET-NUP214 | NXF1 | 61 | 0.312 | 489.736 | 24.52 | 2 | 1.24414 | 0.0204 | 49.03 |
| | FAF1 | 23 | 0.134 | 343.967 | 2.609 | 2 | 4.40782 | 0.19164 | 5.218 |
| | SUPT5H | 61 | 0.05 | 3258.819 | 5.935 | 2 | 5.13901 | 0.08425 | 11.87 |
| | GART | 28 | 0.204 | 437.762 | 7.241 | 2 | 1.93343 | 0.06905 | 14.482 |
| | CUL2 | 833 | 0 | 691392 | 2.998 | 2 | 138.926 | 0.16678 | 5.996 |
| | CUL3 | 70 | 0.081 | 3676.928 | 10.37 | 2 | 3.3748 | 0.04821 | 20.742 |
| | NXF1 | 61 | 0.312 | 489.736 | 24.52 | 2 | 1.24414 | 0.0204 | 49.03 |
| | CUL2 | 833 | 0 | 691392 | 2.998 | 2 | 138.926 | 0.16678 | 5.996 |

| | | | | | | | | | |
|---|---|---|---|---|---|---|---|---|---|
| | RANBP2 | 80 | 0.071 | 8175.982 | 9.805 | 2 | 4.07955 | 0.05099 | 19.61 |
| MNX1-ETV6 | HDAC3 | 23 | 0.134 | 343.967 | 0.509 | 2 | 22.5933 | 0.98232 | 1.018 |
| | ETV6 | 23 | 0.134 | 343.967 | 0.309 | 2 | 37.2168 | 1.61812 | 0.618 |
| | HDAC9 | 145 | 0.089 | 12651.423 | 28.74 | 2 | 2.52262 | 0.0174 | 57.48 |
| | PIN1 | 223 | 0.037 | 40989.149 | 1.089 | 2 | 102.388 | 0.45914 | 2.178 |
| | NCOR1 | 151 | 0.095 | 11956.904 | 16.08 | 2 | 4.69557 | 0.0311 | 32.158 |
| | SIN3A | 192 | 0.072 | 20856.124 | 15.68 | 2 | 6.12284 | 0.03189 | 31.358 |
| | L3MBTL1 | 211 | 0.072 | 26556.162 | 26.9 | 2 | 3.92193 | 0.01859 | 53.8 |
| | ETV7 | 23 | 0.134 | 343.967 | 3.609 | 2 | 3.18648 | 0.13854 | 7.218 |
| | ETV6 | 23 | 0.134 | 343.967 | 0.309 | 2 | 37.2168 | 1.61812 | 0.618 |
| KMT2A-MLLT1 | SMARCC2 | 129 | 0.129 | 6769.42 | 0.623 | 2 | 103.531 | 0.80257 | 1.246 |
| | KMT2A | 394 | 0.109 | 5267.838 | 21.87 | 2 | 9.00695 | 0.02286 | 43.744 |
| | HDAC2 | 301 | 0.067 | 40883.166 | 1.874 | 2 | 80.3095 | 0.26681 | 3.748 |
| | CHD3 | 146 | 0.075 | 13797.187 | 12.66 | 2 | 5.7671 | 0.0395 | 25.316 |
| | SMARCC1 | 116 | 0.148 | 4333.831 | 0.559 | 2 | 103.757 | 0.89445 | 1.118 |
| | POLR2A | 251 | 0.074 | 34918.732 | 1.239 | 2 | 101.291 | 0.40355 | 2.478 |
| | SMARCA2 | 103 | 0.141 | 4685.809 | 16.25 | 2 | 3.16923 | 0.03077 | 32.5 |
| | CREBBP | 296 | 0.058 | 48179.347 | 1.061 | 2 | 139.491 | 0.47125 | 2.122 |
| | SIN3A | 192 | 0.072 | 20856.124 | 15.68 | 2 | 6.12284 | 0.03189 | 31.358 |
| | CTBP1 | 148 | 0.058 | 14901.592 | 0.751 | 2 | 98.5353 | 0.66578 | 1.502 |
| | KMT2A | 394 | 0.109 | 5267.838 | 21.87 | 2 | 9.00695 | 0.02286 | 43.744 |
| | CREBBP | 296 | 0.058 | 48179.347 | 1.061 | 2 | 139.491 | 0.47125 | 2.122 |
| KMT2A-MLLT6 | SMARCC2 | 129 | 0.129 | 6769.42 | 0.623 | 2 | 103.531 | 0.80257 | 1.246 |
| | KMT2A | 394 | 0.109 | 5267.838 | 21.87 | 2 | 9.00695 | 0.02286 | 43.744 |
| | HDAC2 | 301 | 0.067 | 40883.166 | 1.874 | 2 | 80.3095 | 0.26681 | 3.748 |
| | CHD3 | 146 | 0.075 | 13797.187 | 12.66 | 2 | 5.7671 | 0.0395 | 25.316 |
| | SMARCC1 | 116 | 0.148 | 4333.831 | 0.559 | 2 | 103.757 | 0.89445 | 1.118 |
| | POLR2A | 251 | 0.074 | 34918.732 | 1.239 | 2 | 101.291 | 0.40355 | 2.478 |
| | SMARCA2 | 103 | 0.141 | 4685.809 | 16.25 | 2 | 3.16923 | 0.03077 | 32.5 |
| | CREBBP | 296 | 0.058 | 48179.347 | 1.061 | 2 | 139.491 | 0.47125 | 2.122 |
| | SIN3A | 192 | 0.072 | 20856.124 | 15.68 | 2 | 6.12284 | 0.03189 | 31.358 |
| | CTBP1 | 148 | 0.058 | 14901.592 | 0.751 | 2 | 98.5353 | 0.66578 | 1.502 |
| | KMT2A | 394 | 0.109 | 5267.838 | 21.87 | 2 | 9.00695 | 0.02286 | 43.744 |
| | CREBBP | 296 | 0.058 | 48179.347 | 1.061 | 2 | 139.491 | 0.47125 | 2.122 |
| ETV6-ACSL6 | HDAC3 | 23 | 0.134 | 343.967 | 0.509 | 2 | 22.5933 | 0.98232 | 1.018 |
| | ETV6 | 23 | 0.134 | 343.967 | 0.309 | 2 | 37.2168 | 1.61812 | 0.618 |
| | HDAC9 | 145 | 0.089 | 12651.423 | 28.74 | 2 | 2.52262 | 0.0174 | 57.48 |
| | PIN1 | 223 | 0.037 | 40989.149 | 1.089 | 2 | 102.388 | 0.45914 | 2.178 |
| | NCOR1 | 151 | 0.095 | 11956.904 | 16.08 | 2 | 4.69557 | 0.0311 | 32.158 |
| | SIN3A | 192 | 0.072 | 20856.124 | 15.68 | 2 | 6.12284 | 0.03189 | 31.358 |
| | L3MBTL1 | 211 | 0.072 | 26556.162 | 26.9 | 2 | 3.92193 | 0.01859 | 53.8 |
| | ETV7 | 23 | 0.134 | 343.967 | 3.609 | 2 | 3.18648 | 0.13854 | 7.218 |
| | ETV6 | 23 | 0.134 | 343.967 | 0.309 | 2 | 37.2168 | 1.61812 | 0.618 |
| ETV6-MECOM | HDAC1 | 478 | 0 | 212982 | 1.996 | 2 | 119.739 | 0.2505 | 3.992 |
| | UBE2I | 429 | 0 | 182756 | 1.995 | 2 | 107.519 | 0.25063 | 3.99 |

| | | | | | | | | | |
|---|---|---|---|---|---|---|---|---|---|
| | HDAC3 | 23 | 0.134 | 343.967 | 0.509 | 2 | 22.5933 | 0.98232 | 1.018 |
| | EHMT2 | 833 | 0 | 691392 | 42 | 2 | 9.91714 | 0.01191 | 83.996 |
| | ELAVL1 | 213 | 0.076 | 28937.125 | 18.66 | 2 | 5.70617 | 0.02679 | 37.328 |
| | MECOM | 189 | 0.06 | 22876.817 | 12.51 | 2 | 7.55275 | 0.03996 | 25.024 |
| | SMAD1 | 192 | 0.072 | 20856.124 | 11.68 | 2 | 8.21988 | 0.04281 | 23.358 |
| | SMAD2 | 103 | 0.141 | 4685.809 | 2.25 | 2 | 22.8889 | 0.22222 | 4.5 |
| | SMAD3 | 103 | 0.141 | 4685.809 | 26.25 | 2 | 1.9619 | 0.01905 | 52.5 |
| | KAT2B | 175 | 0.066 | 18317.722 | 13.38 | 2 | 6.53815 | 0.03736 | 26.766 |
| | SUV39H1 | 61 | 0.05 | 3258.819 | 3.935 | 2 | 7.75095 | 0.12706 | 7.87 |
| | NCOR1 | 151 | 0.095 | 11956.904 | 16.08 | 2 | 4.69557 | 0.0311 | 32.158 |
| | CTBP1 | 148 | 0.058 | 14901.592 | 0.751 | 2 | 98.5353 | 0.66578 | 1.502 |
| | CREBBP | 296 | 0.058 | 48179.347 | 1.061 | 2 | 139.491 | 0.47125 | 2.122 |
| | MECOM | 189 | 0.06 | 22876.817 | 12.51 | 2 | 7.55275 | 0.03996 | 25.024 |
| | SUV39H1 | 61 | 0.05 | 3258.819 | 3.935 | 2 | 7.75095 | 0.12706 | 7.87 |
| | HDAC4 | 145 | 0.089 | 12651.423 | 16.74 | 2 | 4.33094 | 0.02987 | 33.48 |
| KMT2A-EPS15 | SMARCC2 | 129 | 0.129 | 6769.42 | 0.623 | 2 | 103.531 | 0.80257 | 1.246 |
| | KMT2A | 394 | 0.109 | 5267.838 | 21.87 | 2 | 9.00695 | 0.02286 | 43.744 |
| | HDAC2 | 301 | 0.067 | 40883.166 | 1.874 | 2 | 80.3095 | 0.26681 | 3.748 |
| | CHD3 | 146 | 0.075 | 13797.187 | 12.66 | 2 | 5.7671 | 0.0395 | 25.316 |
| | SMARCC1 | 116 | 0.148 | 4333.831 | 0.559 | 2 | 103.757 | 0.89445 | 1.118 |
| | POLR2A | 251 | 0.074 | 34918.732 | 1.239 | 2 | 101.291 | 0.40355 | 2.478 |
| | SMARCA2 | 103 | 0.141 | 4685.809 | 16.25 | 2 | 3.16923 | 0.03077 | 32.5 |
| | CREBBP | 296 | 0.058 | 48179.347 | 1.061 | 2 | 139.491 | 0.47125 | 2.122 |
| | SIN3A | 192 | 0.072 | 20856.124 | 15.68 | 2 | 6.12284 | 0.03189 | 31.358 |
| | CTBP1 | 148 | 0.058 | 14901.592 | 0.751 | 2 | 98.5353 | 0.66578 | 1.502 |
| | KMT2A | 394 | 0.109 | 5267.838 | 21.87 | 2 | 9.00695 | 0.02286 | 43.744 |
| | CREBBP | 296 | 0.058 | 48179.347 | 1.061 | 2 | 139.491 | 0.47125 | 2.122 |
| KMT2A-GAS7 | SMARCC2 | 129 | 0.129 | 6769.42 | 0.623 | 2 | 103.531 | 0.80257 | 1.246 |
| | KMT2A | 394 | 0.109 | 5267.838 | 21.87 | 2 | 9.00695 | 0.02286 | 43.744 |
| | HDAC2 | 301 | 0.067 | 40883.166 | 1.874 | 2 | 80.3095 | 0.26681 | 3.748 |
| | CHD3 | 146 | 0.075 | 13797.187 | 12.66 | 2 | 5.7671 | 0.0395 | 25.316 |
| | SMARCC1 | 116 | 0.148 | 4333.831 | 0.559 | 2 | 103.757 | 0.89445 | 1.118 |
| | POLR2A | 251 | 0.074 | 34918.732 | 1.239 | 2 | 101.291 | 0.40355 | 2.478 |
| | SMARCA2 | 103 | 0.141 | 4685.809 | 16.25 | 2 | 3.16923 | 0.03077 | 32.5 |
| | CREBBP | 296 | 0.058 | 48179.347 | 1.061 | 2 | 139.491 | 0.47125 | 2.122 |
| | SIN3A | 192 | 0.072 | 20856.124 | 15.68 | 2 | 6.12284 | 0.03189 | 31.358 |
| | CTBP1 | 148 | 0.058 | 14901.592 | 0.751 | 2 | 98.5353 | 0.66578 | 1.502 |
| | KMT2A | 394 | 0.109 | 5267.838 | 21.87 | 2 | 9.00695 | 0.02286 | 43.744 |
| | CREBBP | 296 | 0.058 | 48179.347 | 1.061 | 2 | 139.491 | 0.47125 | 2.122 |
| KMT2A-ABL1 | SMARCC2 | 129 | 0.129 | 6769.42 | 0.623 | 2 | 103.531 | 0.80257 | 1.246 |
| | KMT2A | 394 | 0.109 | 5267.838 | 21.87 | 2 | 9.00695 | 0.02286 | 43.744 |
| | SMARCC1 | 116 | 0.148 | 4333.831 | 0.559 | 2 | 103.757 | 0.89445 | 1.118 |
| | CHD3 | 146 | 0.075 | 13797.187 | 12.66 | 2 | 5.7671 | 0.0395 | 25.316 |
| | POLR2A | 251 | 0.074 | 34918.732 | 1.239 | 2 | 101.291 | 0.40355 | 2.478 |
| | SMARCA2 | 103 | 0.141 | 4685.809 | 16.25 | 2 | 3.16923 | 0.03077 | 32.5 |
| | CREBBP | 296 | 0.058 | 48179.347 | 1.061 | 2 | 139.491 | 0.47125 | 2.122 |

| | | | | | | | | | |
|---|---|---|---|---|---|---|---|---|---|
| | ABL1 | 173 | 0.076 | 17246.646 | 14.9 | 2 | 5.80459 | 0.03355 | 29.804 |
| | CBLB | 44 | 0.083 | 1464.862 | 6.364 | 2 | 3.45695 | 0.07857 | 12.728 |
| | CBL | 233 | 0.061 | 32363.102 | 6.043 | 2 | 19.2785 | 0.08274 | 12.086 |
| KMT2A-MLLT11 | SMARCC2 | 129 | 0.129 | 6769.42 | 0.623 | 2 | 103.531 | 0.80257 | 1.246 |
| | KMT2A | 394 | 0.109 | 5267.838 | 21.87 | 2 | 9.00695 | 0.02286 | 43.744 |
| | HDAC2 | 301 | 0.067 | 40883.166 | 1.874 | 2 | 80.3095 | 0.26681 | 3.748 |
| | CHD3 | 146 | 0.075 | 13797.187 | 12.66 | 2 | 5.7671 | 0.0395 | 25.316 |
| | SMARCC1 | 116 | 0.148 | 4333.831 | 0.559 | 2 | 103.757 | 0.89445 | 1.118 |
| | POLR2A | 251 | 0.074 | 34918.732 | 1.239 | 2 | 101.291 | 0.40355 | 2.478 |
| | SMARCA2 | 103 | 0.141 | 4685.809 | 16.25 | 2 | 3.16923 | 0.03077 | 32.5 |
| | CREBBP | 296 | 0.058 | 48179.347 | 1.061 | 2 | 139.491 | 0.47125 | 2.122 |
| | SIN3A | 192 | 0.072 | 20856.124 | 15.68 | 2 | 6.12284 | 0.03189 | 31.358 |
| | CTBP1 | 148 | 0.058 | 14901.592 | 0.751 | 2 | 98.5353 | 0.66578 | 1.502 |
| | KMT2A | 394 | 0.109 | 5267.838 | 21.87 | 2 | 9.00695 | 0.02286 | 43.744 |
| | CREBBP | 296 | 0.058 | 48179.347 | 1.061 | 2 | 139.491 | 0.47125 | 2.122 |
| MN1-ETV6 | HDAC3 | 23 | 0.134 | 343.967 | 0.509 | 2 | 22.5933 | 0.98232 | 1.018 |
| | ETV6 | 23 | 0.134 | 343.967 | 0.309 | 2 | 37.2168 | 1.61812 | 0.618 |
| | HDAC9 | 145 | 0.089 | 12651.423 | 28.74 | 2 | 2.52262 | 0.0174 | 57.48 |
| | PIN1 | 223 | 0.037 | 40989.149 | 1.089 | 2 | 102.388 | 0.45914 | 2.178 |
| | EP300 | 453 | 0 | 203852 | 1.996 | 2 | 113.477 | 0.2505 | 3.992 |
| | NCOR1 | 151 | 0.095 | 11956.904 | 16.08 | 2 | 4.69557 | 0.0311 | 32.158 |
| | SIN3A | 192 | 0.072 | 20856.124 | 15.68 | 2 | 6.12284 | 0.03189 | 31.358 |
| | HDAC3 | 23 | 0.134 | 343.967 | 0.509 | 2 | 22.5933 | 0.98232 | 1.018 |
| | EP300 | 453 | 0 | 203852 | 1.996 | 2 | 113.477 | 0.2505 | 3.992 |
| | KAT5 | 210 | 0.054 | 31401.393 | 3.238 | 2 | 32.4274 | 0.15442 | 6.476 |
| KMT2A-MAML2 | SMARCC2 | 129 | 0.129 | 6769.42 | 0.623 | 2 | 103.531 | 0.80257 | 1.246 |
| | KMT2A | 394 | 0.109 | 5267.838 | 21.87 | 2 | 9.00695 | 0.02286 | 43.744 |
| | SMARCC1 | 116 | 0.148 | 4333.831 | 0.559 | 2 | 103.757 | 0.89445 | 1.118 |
| | CHD3 | 146 | 0.075 | 13797.187 | 12.66 | 2 | 5.7671 | 0.0395 | 25.316 |
| | SMARCA2 | 103 | 0.141 | 4685.809 | 16.25 | 2 | 3.16923 | 0.03077 | 32.5 |
| | CREBBP | 296 | 0.058 | 48179.347 | 1.061 | 2 | 139.491 | 0.47125 | 2.122 |
| | SMARCA2 | 103 | 0.141 | 4685.809 | 16.25 | 2 | 3.16923 | 0.03077 | 32.5 |
| | EP300 | 453 | 0 | 203852 | 1.996 | 2 | 113.477 | 0.2505 | 3.992 |
| | CREBBP | 296 | 0.058 | 48179.347 | 1.061 | 2 | 139.491 | 0.47125 | 2.122 |
| KMT2A-FOXO4 | SMARCC2 | 129 | 0.129 | 6769.42 | 0.623 | 2 | 103.531 | 0.80257 | 1.246 |
| | KMT2A | 394 | 0.109 | 5267.838 | 21.87 | 2 | 9.00695 | 0.02286 | 43.744 |
| | HDAC2 | 301 | 0.067 | 40883.166 | 1.874 | 2 | 80.3095 | 0.26681 | 3.748 |
| | CHD3 | 146 | 0.075 | 13797.187 | 12.66 | 2 | 5.7671 | 0.0395 | 25.316 |
| | SMARCC1 | 116 | 0.148 | 4333.831 | 0.559 | 2 | 103.757 | 0.89445 | 1.118 |
| | POLR2A | 251 | 0.074 | 34918.732 | 1.239 | 2 | 101.291 | 0.40355 | 2.478 |
| | SMARCA2 | 103 | 0.141 | 4685.809 | 16.25 | 2 | 3.16923 | 0.03077 | 32.5 |
| | CREBBP | 296 | 0.058 | 48179.347 | 1.061 | 2 | 139.491 | 0.47125 | 2.122 |
| | SIN3A | 192 | 0.072 | 20856.124 | 15.68 | 2 | 6.12284 | 0.03189 | 31.358 |
| | CTBP1 | 148 | 0.058 | 14901.592 | 0.751 | 2 | 98.5353 | 0.66578 | 1.502 |
| | KMT2A | 394 | 0.109 | 5267.838 | 21.87 | 2 | 9.00695 | 0.02286 | 43.744 |
| | CREBBP | 296 | 0.058 | 48179.347 | 1.061 | 2 | 139.491 | 0.47125 | 2.122 |

| | | | | | | | | | |
|---|---|---|---|---|---|---|---|---|---|
| DEK-NUP214 | ESR1 | 23 | 0.134 | 343.967 | 0.409 | 2 | 28.1174 | 1.22249 | 0.818 |
| | KAT2B | 175 | 0.066 | 18317.722 | 13.38 | 2 | 6.53815 | 0.03736 | 26.766 |
| | SMAD2 | 103 | 0.141 | 4685.809 | 2.25 | 2 | 22.8889 | 0.22222 | 4.5 |
| | SMAD3 | 103 | 0.141 | 4685.809 | 26.25 | 2 | 1.9619 | 0.01905 | 52.5 |
| | CDK2 | 233 | 0.061 | 32363.102 | 8.043 | 2 | 14.4846 | 0.06217 | 16.086 |
| | DEK | 82 | 0.238 | 2197.27 | 11.06 | 2 | 3.70705 | 0.04521 | 22.12 |
| | EP300 | 453 | 0 | 203852 | 1.996 | 2 | 113.477 | 0.2505 | 3.992 |
| | CREBBP | 296 | 0.058 | 48179.347 | 1.061 | 2 | 139.491 | 0.47125 | 2.122 |
| | NXF1 | 61 | 0.312 | 489.736 | 24.52 | 2 | 1.24414 | 0.0204 | 49.03 |
| | DHX15 | 82 | 0.238 | 2197.27 | 21.06 | 2 | 1.94682 | 0.02374 | 42.12 |
| | CUL2 | 833 | 0 | 691392 | 2.998 | 2 | 138.926 | 0.16678 | 5.996 |
| | CUL3 | 70 | 0.081 | 3676.928 | 10.37 | 2 | 3.3748 | 0.04821 | 20.742 |
| RUNX1-CBFA2T3 | KMT2A | 394 | 0.109 | 5267.838 | 21.87 | 2 | 9.00695 | 0.02286 | 43.744 |
| | SMARCC1 | 116 | 0.148 | 4333.831 | 0.559 | 2 | 103.757 | 0.89445 | 1.118 |
| | SMARCA4 | 200 | 0.088 | 20379.264 | 3.512 | 2 | 28.4738 | 0.14237 | 7.024 |
| | CREBBP | 296 | 0.058 | 48179.347 | 1.061 | 2 | 139.491 | 0.47125 | 2.122 |
| | EP300 | 453 | 0 | 203852 | 1.996 | 2 | 113.477 | 0.2505 | 3.992 |
| | CREBBP | 296 | 0.058 | 48179.347 | 1.061 | 2 | 139.491 | 0.47125 | 2.122 |
| NUP98-PSIP1 | HDAC1 | 478 | 0 | 212982 | 1.996 | 2 | 119.739 | 0.2505 | 3.992 |
| | KMT2A | 394 | 0.109 | 5267.838 | 21.87 | 2 | 9.00695 | 0.02286 | 43.744 |
| | ESR1 | 23 | 0.134 | 343.967 | 0.409 | 2 | 28.1174 | 1.22249 | 0.818 |
| | CTNNB1 | 286 | 0.057 | 51076.364 | 18.22 | 2 | 7.85024 | 0.02745 | 36.432 |
| | EP300 | 453 | 0 | 203852 | 1.996 | 2 | 113.477 | 0.2505 | 3.992 |
| | CREBBP | 296 | 0.058 | 48179.347 | 1.061 | 2 | 139.491 | 0.47125 | 2.122 |
| | NXF1 | 61 | 0.312 | 489.736 | 24.52 | 2 | 1.24414 | 0.0204 | 49.03 |
| | EIF4A3 | 833 | 0 | 691392 | 62 | 2 | 6.71796 | 0.00806 | 123.996 |
| | SON | 44 | 0.174 | 955.71 | 7.289 | 2 | 3.01825 | 0.0686 | 14.578 |
| NUP98-HOXC13 | HDAC1 | 478 | 0 | 212982 | 1.996 | 2 | 119.739 | 0.2505 | 3.992 |
| | CTNNB1 | 286 | 0.057 | 51076.364 | 18.22 | 2 | 7.85024 | 0.02745 | 36.432 |
| | MAPK8 | 172 | 0.045 | 22876.817 | 9.512 | 2 | 9.04121 | 0.05257 | 19.024 |
| | EP300 | 453 | 0 | 203852 | 1.996 | 2 | 113.477 | 0.2505 | 3.992 |
| | CREBBP | 296 | 0.058 | 48179.347 | 1.061 | 2 | 139.491 | 0.47125 | 2.122 |
| | NXF1 | 61 | 0.312 | 489.736 | 24.52 | 2 | 1.24414 | 0.0204 | 49.03 |
| | HNRNPAB | 145 | 0.089 | 12651.423 | 28.74 | 2 | 2.52262 | 0.0174 | 57.48 |
| | EED | 66 | 0.322 | 479.736 | 0.167 | 2 | 197.605 | 2.99401 | 0.334 |
| | KPNB1 | 211 | 0.072 | 26556.162 | 16.9 | 2 | 6.2426 | 0.02959 | 33.8 |
| | HNRNPUL1 | 112 | 0.14 | 6163.826 | 17.23 | 2 | 3.24977 | 0.02902 | 34.464 |
| | HDAC1 | 478 | 0 | 212982 | 1.996 | 2 | 119.739 | 0.2505 | 3.992 |
| | CREBBP | 296 | 0.058 | 48179.347 | 1.061 | 2 | 139.491 | 0.47125 | 2.122 |
| | APC | 168 | 0.03 | 24110.735 | 6.869 | 2 | 12.2289 | 0.07279 | 13.738 |
| | CSNK2A1 | 382 | 0 | 144780 | 22 | 2 | 8.68379 | 0.02273 | 43.99 |
| | CTNNB1 | 286 | 0.057 | 51076.364 | 18.22 | 2 | 7.85024 | 0.02745 | 36.432 |
| | APC | 168 | 0.03 | 24110.735 | 6.869 | 2 | 12.2289 | 0.07279 | 13.738 |
| | CREBBP | 296 | 0.058 | 48179.347 | 1.061 | 2 | 139.491 | 0.47125 | 2.122 |
| NUP98-HOXC11 | HDAC1 | 478 | 0 | 212982 | 1.996 | 2 | 119.739 | 0.2505 | 3.992 |

| | | | | | | | | | |
|---|---|---|---|---|---|---|---|---|---|
| | STAT3 | 223 | 0.044 | 37462.83 | 5.641 | 2 | 19.766 | 0.08864 | 11.282 |
| | SP1 | 44 | 0.174 | 955.71 | 7.289 | 2 | 3.01825 | 0.0686 | 14.578 |
| | SMAD3 | 103 | 0.141 | 4685.809 | 26.25 | 2 | 1.9619 | 0.01905 | 52.5 |
| | CTNNB1 | 286 | 0.057 | 51076.364 | 18.22 | 2 | 7.85024 | 0.02745 | 36.432 |
| | MAPK8 | 172 | 0.045 | 22876.817 | 9.512 | 2 | 9.04121 | 0.05257 | 19.024 |
| | EP300 | 453 | 0 | 203852 | 1.996 | 2 | 113.477 | 0.2505 | 3.992 |
| | CREBBP | 296 | 0.058 | 48179.347 | 1.061 | 2 | 139.491 | 0.47125 | 2.122 |
| | CTNNB1 | 286 | 0.057 | 51076.364 | 18.22 | 2 | 7.85024 | 0.02745 | 36.432 |
| | APC | 168 | 0.03 | 24110.735 | 6.869 | 2 | 12.2289 | 0.07279 | 13.738 |
| | CREBBP | 296 | 0.058 | 48179.347 | 1.061 | 2 | 139.491 | 0.47125 | 2.122 |
| PAX5-ETV6 | UBE2I | 429 | 0 | 182756 | 1.995 | 2 | 107.519 | 0.25063 | 3.99 |
| | HDAC3 | 23 | 0.134 | 343.967 | 0.509 | 2 | 22.5933 | 0.98232 | 1.018 |
| | TBP | 156 | 0.071 | 15458.811 | 12.86 | 2 | 6.06579 | 0.03888 | 25.718 |
| | KAT5 | 210 | 0.054 | 31401.393 | 3.238 | 2 | 32.4274 | 0.15442 | 6.476 |
| | RB1 | 217 | 0.059 | 30195.4 | 14.66 | 2 | 7.40059 | 0.0341 | 29.322 |
| | PAX5 | 9 | 0.167 | 51 | 0.2 | 2 | 22.5 | 2.5 | 0.4 |
| | RUNX1 | 68 | 0.149 | 2304.921 | 11.65 | 2 | 2.91921 | 0.04293 | 23.294 |
| | PIN1 | 223 | 0.037 | 40989.149 | 1.089 | 2 | 102.388 | 0.45914 | 2.178 |
| | EP300 | 453 | 0 | 203852 | 1.996 | 2 | 113.477 | 0.2505 | 3.992 |
| | NCOR1 | 151 | 0.095 | 11956.904 | 16.08 | 2 | 4.69557 | 0.0311 | 32.158 |
| | MAPK1 | 214 | 0.042 | 32283.959 | 20.1 | 2 | 5.32418 | 0.02488 | 40.194 |
| | EP300 | 453 | 0 | 203852 | 1.996 | 2 | 113.477 | 0.2505 | 3.992 |
| | HDAC6 | 145 | 0.089 | 12651.423 | 18.74 | 2 | 3.86873 | 0.02668 | 37.48 |
| NUP98-HOXA11 | HDAC1 | 478 | 0 | 212982 | 1.996 | 2 | 119.739 | 0.2505 | 3.992 |
| | HDAC2 | 301 | 0.067 | 40883.166 | 1.874 | 2 | 80.3095 | 0.26681 | 3.748 |
| | YY1 | 121 | 0.098 | 8527.308 | 13.55 | 2 | 4.46363 | 0.03689 | 27.108 |
| | CTNNB1 | 286 | 0.057 | 51076.364 | 18.22 | 2 | 7.85024 | 0.02745 | 36.432 |
| | CSNK2A1 | 382 | 0 | 144780 | 22 | 2 | 8.68379 | 0.02273 | 43.99 |
| | MAPK8 | 172 | 0.045 | 22876.817 | 9.512 | 2 | 9.04121 | 0.05257 | 19.024 |
| | EP300 | 453 | 0 | 203852 | 1.996 | 2 | 113.477 | 0.2505 | 3.992 |
| | CREBBP | 296 | 0.058 | 48179.347 | 1.061 | 2 | 139.491 | 0.47125 | 2.122 |
| | CTNNB1 | 286 | 0.057 | 51076.364 | 18.22 | 2 | 7.85024 | 0.02745 | 36.432 |
| | APC | 168 | 0.03 | 24110.735 | 6.869 | 2 | 12.2289 | 0.07279 | 13.738 |
| | CREBBP | 296 | 0.058 | 48179.347 | 1.061 | 2 | 139.491 | 0.47125 | 2.122 |
| BCR-PDGFRA | TGFBR2 | 82 | 0.046 | 5541.307 | 5.687 | 2 | 7.20943 | 0.08792 | 11.374 |
| | PDGFRA | 48 | 0.095 | 1544.563 | 6.25 | 2 | 3.84 | 0.08 | 12.5 |
| | SHC1 | 214 | 0.062 | 25801.088 | 14.99 | 2 | 7.13762 | 0.03335 | 29.982 |
| | CRKL | 108 | 0.065 | 8323.422 | 8.844 | 2 | 6.10583 | 0.05654 | 17.688 |
| | EGFR | 833 | 0 | 691392 | 52 | 2 | 8.00992 | 0.00962 | 103.996 |
| | PLCG1 | 112 | 0.071 | 8175.982 | 9.805 | 2 | 5.71137 | 0.05099 | 19.61 |
| | FES | 23 | 0.134 | 343.967 | 4.609 | 2 | 2.49512 | 0.10848 | 9.218 |
| | CRK | 148 | 0.058 | 14901.592 | 11.35 | 2 | 6.51925 | 0.04405 | 22.702 |
| | ABL1 | 173 | 0.076 | 17246.646 | 14.9 | 2 | 5.80459 | 0.03355 | 29.804 |
| | HCK | 59 | 0.091 | 2440.137 | 7.133 | 2 | 4.13571 | 0.0701 | 14.266 |
| | PTPN6 | 65 | 0.086 | 3019.11 | 7.364 | 2 | 4.41336 | 0.0679 | 14.728 |
| | BCR | 56 | 0.145 | 1733.391 | 1.789 | 2 | 15.6512 | 0.27949 | 3.578 |
| | UBASH3B | 109 | 0.097 | 6392.818 | 12.26 | 2 | 4.44644 | 0.04079 | 24.514 |

| | | | | | | | | | |
|---|---|---|---|---|---|---|---|---|---|
| | INPP5D | 39 | 0.147 | 977.622 | 7.4 | 2 | 2.63514 | 0.06757 | 14.8 |
| | SOS1 | 44 | 0.174 | 955.71 | 9.289 | 2 | 2.36839 | 0.05383 | 18.578 |
| | GRB2 | 515 | 0 | 263682 | 26 | 2 | 9.90537 | 0.01923 | 51.992 |
| | NTRK1 | 166 | 0.045 | 22876.817 | 9.512 | 2 | 8.72582 | 0.05257 | 19.024 |
| | CBL | 233 | 0.061 | 32363.102 | 6.043 | 2 | 19.2785 | 0.08274 | 12.086 |
| | KIT | 42 | 0.156 | 1050.358 | 8.186 | 2 | 2.56536 | 0.06108 | 16.372 |
| | DOK1 | 33 | 0.29 | 283.697 | 0.167 | 2 | 98.8024 | 2.99401 | 0.334 |
| | PIK3R2 | 109 | 0.111 | 6174.794 | 13.89 | 2 | 3.9234 | 0.03599 | 27.782 |
| | PIK3R1 | 145 | 0.079 | 12813.777 | 13.15 | 2 | 5.51541 | 0.03804 | 26.29 |
| | ABL1 | 173 | 0.076 | 17246.646 | 14.9 | 2 | 5.80459 | 0.03355 | 29.804 |
| | TP53 | 961 | 0 | 920640 | 1.998 | 2 | 240.49 | 0.25025 | 3.996 |
| | RB1 | 217 | 0.059 | 30195.4 | 14.66 | 2 | 7.40059 | 0.0341 | 29.322 |
| | BCR | 56 | 0.145 | 1733.391 | 1.789 | 2 | 15.6512 | 0.27949 | 3.578 |
| BCR-FGFR1 | SRC | 44 | 0.174 | 955.71 | 5.289 | 2 | 4.15958 | 0.09454 | 10.578 |
| | ITK | 41 | 0.133 | 1028.658 | 11.14 | 2 | 1.83972 | 0.04487 | 22.286 |
| | SOS1 | 44 | 0.174 | 955.71 | 9.289 | 2 | 2.36839 | 0.05383 | 18.578 |
| | ERBB3 | 23 | 0.134 | 343.967 | 8.609 | 2 | 1.33581 | 0.05808 | 17.218 |
| | VAV1 | 41 | 0.248 | 760.708 | 9.619 | 2 | 2.1312 | 0.05198 | 19.238 |
| | CBL | 233 | 0.061 | 32363.102 | 6.043 | 2 | 19.2785 | 0.08274 | 12.086 |
| | PLCG1 | 112 | 0.071 | 8175.982 | 9.805 | 2 | 5.71137 | 0.05099 | 19.61 |
| | ABL1 | 173 | 0.076 | 17246.646 | 14.9 | 2 | 5.80459 | 0.03355 | 29.804 |
| | ABL1 | 173 | 0.076 | 17246.646 | 14.9 | 2 | 5.80459 | 0.03355 | 29.804 |
| | HCK | 59 | 0.091 | 2440.137 | 7.133 | 2 | 4.13571 | 0.0701 | 14.266 |
| | BCR | 56 | 0.145 | 1733.391 | 1.789 | 2 | 15.6512 | 0.27949 | 3.578 |
| | CBL | 233 | 0.061 | 32363.102 | 6.043 | 2 | 19.2785 | 0.08274 | 12.086 |
| IGH-BCL6 | HDAC1 | 478 | 0 | 212982 | 1.996 | 2 | 119.739 | 0.2505 | 3.992 |
| | TP53 | 961 | 0 | 920640 | 1.998 | 2 | 240.49 | 0.25025 | 3.996 |
| | CTBP1 | 148 | 0.058 | 14901.592 | 0.751 | 2 | 98.5353 | 0.66578 | 1.502 |
| | EP300 | 453 | 0 | 203852 | 1.996 | 2 | 113.477 | 0.2505 | 3.992 |
| | NCOR2 | 104 | 0.136 | 4824.426 | 15.81 | 2 | 3.28906 | 0.03163 | 31.62 |
| | CREBBP | 296 | 0.058 | 48179.347 | 1.061 | 2 | 139.491 | 0.47125 | 2.122 |
| | HDAC2 | 301 | 0.067 | 40883.166 | 1.874 | 2 | 80.3095 | 0.26681 | 3.748 |
| | SMARCA4 | 200 | 0.088 | 20379.264 | 3.512 | 2 | 28.4738 | 0.14237 | 7.024 |
| | CREBBP | 296 | 0.058 | 48179.347 | 1.061 | 2 | 139.491 | 0.47125 | 2.122 |
| LCP1-BCL6 | HDAC1 | 478 | 0 | 212982 | 1.996 | 2 | 119.739 | 0.2505 | 3.992 |
| | TP53 | 961 | 0 | 920640 | 1.998 | 2 | 240.49 | 0.25025 | 3.996 |
| | CTBP1 | 148 | 0.058 | 14901.592 | 0.751 | 2 | 98.5353 | 0.66578 | 1.502 |
| | EP300 | 453 | 0 | 203852 | 1.996 | 2 | 113.477 | 0.2505 | 3.992 |
| | NCOR2 | 104 | 0.136 | 4824.426 | 15.81 | 2 | 3.28906 | 0.03163 | 31.62 |
| | CREBBP | 296 | 0.058 | 48179.347 | 1.061 | 2 | 139.491 | 0.47125 | 2.122 |
| | HDAC2 | 301 | 0.067 | 40883.166 | 1.874 | 2 | 80.3095 | 0.26681 | 3.748 |
| | SMARCA4 | 200 | 0.088 | 20379.264 | 3.512 | 2 | 28.4738 | 0.14237 | 7.024 |
| | CREBBP | 296 | 0.058 | 48179.347 | 1.061 | 2 | 139.491 | 0.47125 | 2.122 |
| CREBBP-KAT6A | SMARCC2 | 129 | 0.129 | 6769.42 | 0.623 | 2 | 103.531 | 0.80257 | 1.246 |
| | KMT2A | 394 | 0.109 | 5267.838 | 21.87 | 2 | 9.00695 | 0.02286 | 43.744 |
| | HDAC2 | 301 | 0.067 | 40883.166 | 1.874 | 2 | 80.3095 | 0.26681 | 3.748 |
| | CHD3 | 146 | 0.075 | 13797.187 | 12.66 | 2 | 5.7671 | 0.0395 | 25.316 |
| | SMARCC1 | 116 | 0.148 | 4333.831 | 0.559 | 2 | 103.757 | 0.89445 | 1.118 |

| | | | | | | | | | |
|---|---|---|---|---|---|---|---|---|---|
| | POLR2A | 251 | 0.074 | 34918.732 | 1.239 | 2 | 101.291 | 0.40355 | 2.478 |
| | SMARCA2 | 103 | 0.141 | 4685.809 | 16.25 | 2 | 3.16923 | 0.03077 | 32.5 |
| | CREBBP | 296 | 0.058 | 48179.347 | 1.061 | 2 | 139.491 | 0.47125 | 2.122 |
| | SIN3A | 192 | 0.072 | 20856.124 | 15.68 | 2 | 6.12284 | 0.03189 | 31.358 |
| | CTBP1 | 148 | 0.058 | 14901.592 | 0.751 | 2 | 98.5353 | 0.66578 | 1.502 |
| | KMT2A | 394 | 0.109 | 5267.838 | 21.87 | 2 | 9.00695 | 0.02286 | 43.744 |
| | CREBBP | 296 | 0.058 | 48179.347 | 1.061 | 2 | 139.491 | 0.47125 | 2.122 |
| KMT2A-ARHGAP26 | SMARCC2 | 129 | 0.129 | 6769.42 | 0.623 | 2 | 103.531 | 0.80257 | 1.246 |
| | KMT2A | 394 | 0.109 | 5267.838 | 21.87 | 2 | 9.00695 | 0.02286 | 43.744 |
| | HDAC2 | 301 | 0.067 | 40883.166 | 1.874 | 2 | 80.3095 | 0.26681 | 3.748 |
| | CHD3 | 146 | 0.075 | 13797.187 | 12.66 | 2 | 5.7671 | 0.0395 | 25.316 |
| | SMARCC1 | 116 | 0.148 | 4333.831 | 0.559 | 2 | 103.757 | 0.89445 | 1.118 |
| | POLR2A | 251 | 0.074 | 34918.732 | 1.239 | 2 | 101.291 | 0.40355 | 2.478 |
| | SMARCA2 | 103 | 0.141 | 4685.809 | 16.25 | 2 | 3.16923 | 0.03077 | 32.5 |
| | CREBBP | 296 | 0.058 | 48179.347 | 1.061 | 2 | 139.491 | 0.47125 | 2.122 |
| | SIN3A | 192 | 0.072 | 20856.124 | 15.68 | 2 | 6.12284 | 0.03189 | 31.358 |
| | CTBP1 | 148 | 0.058 | 14901.592 | 0.751 | 2 | 98.5353 | 0.66578 | 1.502 |
| | KMT2A | 394 | 0.109 | 5267.838 | 21.87 | 2 | 9.00695 | 0.02286 | 43.744 |
| | CREBBP | 296 | 0.058 | 48179.347 | 1.061 | 2 | 139.491 | 0.47125 | 2.122 |
| FOXO3-KMT2A | SMARCC2 | 129 | 0.129 | 6769.42 | 0.623 | 2 | 103.531 | 0.80257 | 1.246 |
| | KMT2A | 394 | 0.109 | 5267.838 | 21.87 | 2 | 9.00695 | 0.02286 | 43.744 |
| | SMARCC1 | 116 | 0.148 | 4333.831 | 0.559 | 2 | 103.757 | 0.89445 | 1.118 |
| | CHD3 | 146 | 0.075 | 13797.187 | 12.66 | 2 | 5.7671 | 0.0395 | 25.316 |
| | SMARCA2 | 103 | 0.141 | 4685.809 | 16.25 | 2 | 3.16923 | 0.03077 | 32.5 |
| | CREBBP | 296 | 0.058 | 48179.347 | 1.061 | 2 | 139.491 | 0.47125 | 2.122 |
| | SMARCA2 | 103 | 0.141 | 4685.809 | 16.25 | 2 | 3.16923 | 0.03077 | 32.5 |
| | EP300 | 453 | 0 | 203852 | 1.996 | 2 | 113.477 | 0.2505 | 3.992 |
| | CREBBP | 296 | 0.058 | 48179.347 | 1.061 | 2 | 139.491 | 0.47125 | 2.122 |
| KMT2A-DCPS | SMARCC2 | 129 | 0.129 | 6769.42 | 0.623 | 2 | 103.531 | 0.80257 | 1.246 |
| | KMT2A | 394 | 0.109 | 5267.838 | 21.87 | 2 | 9.00695 | 0.02286 | 43.744 |
| | HDAC2 | 301 | 0.067 | 40883.166 | 1.874 | 2 | 80.3095 | 0.26681 | 3.748 |
| | CHD3 | 146 | 0.075 | 13797.187 | 12.66 | 2 | 5.7671 | 0.0395 | 25.316 |
| | SMARCC1 | 116 | 0.148 | 4333.831 | 0.559 | 2 | 103.757 | 0.89445 | 1.118 |
| | POLR2A | 251 | 0.074 | 34918.732 | 1.239 | 2 | 101.291 | 0.40355 | 2.478 |
| | SMARCA2 | 103 | 0.141 | 4685.809 | 16.25 | 2 | 3.16923 | 0.03077 | 32.5 |
| | CREBBP | 296 | 0.058 | 48179.347 | 1.061 | 2 | 139.491 | 0.47125 | 2.122 |
| | SIN3A | 192 | 0.072 | 20856.124 | 15.68 | 2 | 6.12284 | 0.03189 | 31.358 |
| | CTBP1 | 148 | 0.058 | 14901.592 | 0.751 | 2 | 98.5353 | 0.66578 | 1.502 |
| | KMT2A | 394 | 0.109 | 5267.838 | 21.87 | 2 | 9.00695 | 0.02286 | 43.744 |
| | CREBBP | 296 | 0.058 | 48179.347 | 1.061 | 2 | 139.491 | 0.47125 | 2.122 |
| IGH-CEBPE | UBE2I | 429 | 0 | 182756 | 1.995 | 2 | 107.519 | 0.25063 | 3.99 |
| | BATF | 19 | 0.34 | 136.825 | 7.368 | 2 | 1.28936 | 0.06786 | 14.736 |
| | DDIT3 | 70 | 0.081 | 3676.928 | 7.371 | 2 | 4.74834 | 0.06783 | 14.742 |
| | CEBPG | 22 | 0.371 | 143.851 | 9 | 2 | 1.22222 | 0.05556 | 18 |
| | CEBPE | 44 | 0.083 | 1464.862 | 5.364 | 2 | 4.10142 | 0.09321 | 10.728 |
| | FOS | 130 | 0.053 | 12361.493 | 8.769 | 2 | 7.41248 | 0.05702 | 17.538 |

| | | | | | | | | | |
|---|---|---|---|---|---|---|---|---|---|
| | JUN | 211 | 0.072 | 25977.077 | 17.06 | 2 | 6.18406 | 0.02931 | 34.12 |
| | STAT6 | 61 | 0.05 | 3258.819 | 4.935 | 2 | 6.18034 | 0.10132 | 9.87 |
| | RB1 | 217 | 0.059 | 30195.4 | 14.66 | 2 | 7.40059 | 0.0341 | 29.322 |
| | PIAS1 | 98 | 0.08 | 5869.835 | 1.592 | 2 | 30.7789 | 0.31407 | 3.184 |
| | FOSL1 | 28 | 0.204 | 437.762 | 7.241 | 2 | 1.93343 | 0.06905 | 14.482 |
| | BATF3 | 21 | 0.311 | 166.658 | 7.524 | 2 | 1.39553 | 0.06645 | 15.048 |
| | BATF2 | 17 | 0.191 | 187.233 | 0.078 | 2 | 109.254 | 6.42674 | 0.1556 |
| | ATF4 | 73 | 0.073 | 4098.286 | 7.068 | 2 | 5.16412 | 0.07074 | 14.136 |
| | MYB | 48 | 0.135 | 1352.975 | 8.163 | 2 | 2.9401 | 0.06125 | 16.326 |
| | ATF3 | 59 | 0.171 | 1701.127 | 2.525 | 2 | 11.6832 | 0.19802 | 5.05 |

**Table S18: SC-PAS (TRAINING)**

| | Essential Community Vertices | D | CC | BC | Avg.D (avgD) | Net.Diam (ND) | PAS | PAS/D | D/PAS |
|---|---|---|---|---|---|---|---|---|---|
| ASTN2-CNOT2 | CNOT6L | 18 | 0.359 | 136.43 | 7.684 | 2 | 1.171 | 0.065 | 15.37 |
| | AURKA | 81 | 0.074 | 4620.2 | 9.728 | 2 | 4.163 | 0.051 | 19.46 |
| | CNOT8 | 23 | 0.174 | 326.82 | 5.583 | 2 | 2.06 | 0.09 | 11.17 |
| | TNRC6C | 29 | 0.204 | 469.91 | 7.467 | 2 | 1.942 | 0.067 | 14.93 |
| | TNRC6B | 70 | 0.118 | 3119.7 | 10 | 2 | 3.5 | 0.05 | 20 |
| | CNOT3 | 29 | 0.204 | 439.17 | 0.167 | 2 | 86.83 | 2.994 | 0.334 |
| | CNOT2 | 46 | 0.124 | 1505.4 | 7.404 | 2 | 3.106 | 0.068 | 14.81 |
| | CNOT1 | 79 | 0.084 | 4389.2 | 8.475 | 2 | 4.661 | 0.059 | 16.95 |
| | CNOT7 | 44 | 0.162 | 1104.6 | 8.756 | 2 | 2.513 | 0.057 | 17.51 |
| | AGO2 | 88 | 0.085 | 5258.7 | 9.303 | 2 | 4.73 | 0.054 | 18.61 |
| | HDAC3 | 23 | 0.134 | 343.97 | 0.509 | 2 | 22.59 | 0.982 | 1.018 |
| | GPS2 | 13 | 0.246 | 389.67 | 9.333 | 2 | 0.696 | 0.054 | 18.67 |
| | CNOT2 | 46 | 0.124 | 1505.4 | 7.404 | 2 | 3.106 | 0.068 | 14.81 |
| | NCOR2 | 104 | 0.136 | 4824.4 | 15.81 | 2 | 3.289 | 0.032 | 31.62 |
| | NCOR1 | 151 | 0.095 | 11957 | 16.08 | 2 | 4.696 | 0.031 | 32.16 |
| BCOR-ZC3H7B | HDAC3 | 23 | 0.134 | 343.97 | 0.509 | 2 | 22.59 | 0.982 | 1.018 |
| | HDAC4 | 145 | 0.089 | 12651 | 16.74 | 2 | 4.331 | 0.03 | 33.48 |
| | SP1 | 44 | 0.174 | 955.71 | 7.289 | 2 | 3.018 | 0.069 | 14.58 |
| | CTBP1 | 148 | 0.058 | 14902 | 0.751 | 2 | 98.54 | 0.666 | 1.502 |
| | NACC1 | 30 | 0.069 | 693.07 | 3.8 | 2 | 3.947 | 0.132 | 7.6 |
| | NCOR2 | 104 | 0.136 | 4824.4 | 15.81 | 2 | 3.289 | 0.032 | 31.62 |
| | HDAC1 | 478 | 0 | 212982 | 1.996 | 2 | 119.7 | 0.251 | 3.992 |
| | CTBP1 | 148 | 0.058 | 14902 | 0.751 | 2 | 98.54 | 0.666 | 1.502 |
| | NCOR2 | 104 | 0.136 | 4824.4 | 15.81 | 2 | 3.289 | 0.032 | 31.62 |
| BCOR-CCNB3 | HDAC3 | 23 | 0.134 | 343.97 | 0.509 | 2 | 22.59 | 0.982 | 1.018 |
| | HDAC4 | 145 | 0.089 | 12651 | 16.74 | 2 | 4.331 | 0.03 | 33.48 |
| | SP1 | 44 | 0.174 | 955.71 | 7.289 | 2 | 3.018 | 0.069 | 14.58 |
| | CTBP1 | 148 | 0.058 | 14902 | 0.751 | 2 | 98.54 | 0.666 | 1.502 |
| | NACC1 | 30 | 0.069 | 693.07 | 3.8 | 2 | 3.947 | 0.132 | 7.6 |
| | NCOR2 | 104 | 0.136 | 4824.4 | 15.81 | 2 | 3.289 | 0.032 | 31.62 |
| | HDAC1 | 478 | 0 | 212982 | 1.996 | 2 | 119.7 | 0.251 | 3.992 |
| | CTBP1 | 148 | 0.058 | 14902 | 0.751 | 2 | 98.54 | 0.666 | 1.502 |
| | NCOR2 | 104 | 0.136 | 4824.4 | 15.81 | 2 | 3.289 | 0.032 | 31.62 |

| | | | | | | | | | |
|---|---|---|---|---|---|---|---|---|---|
| CDX1-IRF2BP2 | ELAVL1 | 213 | 0.076 | 28937 | 18.66 | 2 | 5.706 | 0.027 | 37.33 |
| | NTRK1 | 166 | 0.045 | 22877 | 9.512 | 2 | 8.726 | 0.053 | 19.02 |
| | IRF2BPL | 58 | 0.162 | 1617 | 21.05 | 2 | 1.378 | 0.024 | 42.1 |
| | IRF2BP2 | 58 | 0.162 | 1617 | 11.05 | 2 | 2.624 | 0.045 | 22.1 |
| | RBM39 | 144 | 0.15 | 8587.6 | 23.28 | 2 | 3.092 | 0.021 | 46.57 |
| CREB1-EWSR1 | BRCA1 | 17 | 0.191 | 187.23 | 0.09 | 2 | 94.44 | 5.556 | 0.18 |
| | EPAS1 | 102 | 0.1 | 6894.1 | 0.202 | 2 | 252.5 | 2.475 | 0.404 |
| | MYOD1 | 68 | 0.111 | 2755.6 | 9.206 | 2 | 3.693 | 0.054 | 18.41 |
| | ESR1 | 23 | 0.134 | 343.97 | 0.409 | 2 | 28.12 | 1.222 | 0.818 |
| | JUN | 211 | 0.072 | 25977 | 17.06 | 2 | 6.184 | 0.029 | 34.12 |
| | POLR2A | 251 | 0.074 | 34919 | 1.239 | 2 | 101.3 | 0.404 | 2.478 |
| | EP300 | 453 | 0 | 203852 | 1.996 | 2 | 113.5 | 0.251 | 3.992 |
| | SMARCA4 | 200 | 0.088 | 20379 | 3.512 | 2 | 28.47 | 0.142 | 7.024 |
| | EWSR1 | 643 | 0 | 411522 | 32 | 2 | 10.05 | 0.016 | 63.99 |
| | CREBBP | 296 | 0.058 | 48179 | 1.061 | 2 | 139.5 | 0.471 | 2.122 |
| | NR3C1 | 157 | 0.069 | 15985 | 12.61 | 2 | 6.225 | 0.04 | 25.22 |
| | EP300 | 453 | 0 | 203852 | 1.996 | 2 | 113.5 | 0.251 | 3.992 |
| | SMARCA4 | 200 | 0.088 | 20379 | 3.512 | 2 | 28.47 | 0.142 | 7.024 |
| | CREBBP | 296 | 0.058 | 48179 | 1.061 | 2 | 139.5 | 0.471 | 2.122 |
| | NONO | 144 | 0.105 | 10404 | 16.86 | 2 | 4.27 | 0.03 | 33.72 |
| | SMARCA4 | 200 | 0.088 | 20379 | 3.512 | 2 | 28.47 | 0.142 | 7.024 |
| | CUL3 | 70 | 0.081 | 3676.9 | 10.37 | 2 | 3.375 | 0.048 | 20.74 |
| CTDSP2-FAM19A2 | SETD1A | 27 | 0.217 | 375.33 | 7.357 | 2 | 1.835 | 0.068 | 14.71 |
| | POLR2A | 251 | 0.074 | 34919 | 1.239 | 2 | 101.3 | 0.404 | 2.478 |
| | INTS6 | 58 | 0.162 | 1617 | 1.051 | 2 | 27.59 | 0.476 | 2.102 |
| | CTDSP1 | 14 | 0.066 | 165.33 | 2.667 | 2 | 2.625 | 0.187 | 5.334 |
| | CTDSP2 | 14 | 0.066 | 165.33 | 1.667 | 2 | 4.199 | 0.3 | 3.334 |
| EPC1-PHF1 | HDAC1 | 478 | 0 | 212982 | 1.996 | 2 | 119.7 | 0.251 | 3.992 |
| | DHX9 | 75 | 0.238 | 2197.3 | 11.06 | 2 | 3.391 | 0.045 | 22.12 |
| | TP53 | 961 | 0 | 920640 | 1.998 | 2 | 240.5 | 0.25 | 3.996 |
| | ELAVL1 | 213 | 0.076 | 28937 | 18.66 | 2 | 5.706 | 0.027 | 37.33 |
| | E2F6 | 37 | 0.18 | 744.18 | 8.263 | 2 | 2.239 | 0.061 | 16.53 |
| | RBBP7 | 151 | 0.094 | 11807 | 15.99 | 2 | 4.723 | 0.031 | 31.97 |
| | RBBP4 | 170 | 0.106 | 12864 | 19.75 | 2 | 4.303 | 0.025 | 39.51 |
| | EZH1 | 9 | 0.571 | 9.333 | 0.033 | 2 | 135.1 | 15.02 | 0.067 |
| | EZH2 | 275 | 0.045 | 49579 | 1.233 | 2 | 111.5 | 0.406 | 2.466 |
| | EED | 66 | 0.322 | 479.74 | 0.167 | 2 | 197.6 | 2.994 | 0.334 |
| | XRCC6 | 264 | 0.064 | 41296 | 0.568 | 2 | 232.4 | 0.88 | 1.136 |
| | XRCC5 | 264 | 0.064 | 41296 | 0.868 | 2 | 152.1 | 0.576 | 1.736 |
| | PHF1 | 23 | 0.194 | 380.67 | 6 | 2 | 1.917 | 0.083 | 12 |
| | HDAC1 | 478 | 0 | 212982 | 1.996 | 2 | 119.7 | 0.251 | 3.992 |
| | TP53 | 961 | 0 | 920640 | 1.998 | 2 | 240.5 | 0.25 | 3.996 |
| | YEATS4 | 99 | 0.061 | 8123.4 | 7.92 | 2 | 6.25 | 0.063 | 15.84 |
| | TRIM27 | 11 | 0.255 | 62.833 | 4.167 | 2 | 1.32 | 0.12 | 8.334 |
| | KAT5 | 210 | 0.054 | 31401 | 3.238 | 2 | 32.43 | 0.154 | 6.476 |
| | TRIM23 | 11 | 0.255 | 62.833 | 4.167 | 2 | 1.32 | 0.12 | 8.334 |
| | HIST1H2BA | 90 | 0.093 | 5635.8 | 10.13 | 2 | 4.442 | 0.049 | 20.26 |
| | DMAP1 | 58 | 0.182 | 1738.4 | 12.17 | 2 | 2.383 | 0.041 | 24.34 |

| | | | | | | | | | |
|---|---|---|---|---|---|---|---|---|---|
| | MYC | 603 | 0 | 361802 | 1.997 | 2 | 151 | 0.25 | 3.994 |
| | XRCC6 | 264 | 0.064 | 41296 | 0.568 | 2 | 232.4 | 0.88 | 1.136 |
| | MORF4L1 | 106 | 0.087 | 6884 | 10.98 | 2 | 4.827 | 0.046 | 21.96 |
| | ING3 | 58 | 0.162 | 1617 | 1.051 | 2 | 27.59 | 0.476 | 2.102 |
| ERG-EWSR1 | TP53 | 961 | 0 | 920640 | 1.998 | 2 | 240.5 | 0.25 | 3.996 |
| | ESR1 | 23 | 0.134 | 343.97 | 0.409 | 2 | 28.12 | 1.222 | 0.818 |
| | PARP1 | 219 | 0.08 | 25770 | 1.233 | 2 | 88.81 | 0.406 | 2.466 |
| | EP300 | 453 | 0 | 203852 | 1.996 | 2 | 113.5 | 0.251 | 3.992 |
| | CREBBP | 296 | 0.058 | 48179 | 1.061 | 2 | 139.5 | 0.471 | 2.122 |
| | XRCC6 | 264 | 0.064 | 41296 | 0.568 | 2 | 232.4 | 0.88 | 1.136 |
| | EPAS1 | 102 | 0.1 | 6894.1 | 0.202 | 2 | 252.5 | 2.475 | 0.404 |
| | EP300 | 453 | 0 | 203852 | 1.996 | 2 | 113.5 | 0.251 | 3.992 |
| | CREBBP | 296 | 0.058 | 48179 | 1.061 | 2 | 139.5 | 0.471 | 2.122 |
| ETV6-NTRK3 | PDGFRB | 55 | 0.122 | 1609.8 | 8.291 | 2 | 3.317 | 0.06 | 16.58 |
| | SHC1 | 214 | 0.062 | 25801 | 14.99 | 2 | 7.138 | 0.033 | 29.98 |
| | CRKL | 108 | 0.065 | 8323.4 | 8.844 | 2 | 6.106 | 0.057 | 17.69 |
| | GAB2 | 33 | 0.246 | 389.67 | 9.333 | 2 | 1.768 | 0.054 | 18.67 |
| | NTRK1 | 166 | 0.045 | 22877 | 9.512 | 2 | 8.726 | 0.053 | 19.02 |
| | PLCG1 | 112 | 0.071 | 8176 | 9.805 | 2 | 5.711 | 0.051 | 19.61 |
| | GRB2 | 515 | 0 | 263682 | 26 | 2 | 9.905 | 0.019 | 51.99 |
| | HDAC3 | 23 | 0.134 | 343.97 | 0.509 | 2 | 22.59 | 0.982 | 1.018 |
| | PIN1 | 223 | 0.037 | 40989 | 1.089 | 2 | 102.4 | 0.459 | 2.178 |
| | ETV6 | 33 | 0.079 | 840.83 | 0.603 | 2 | 27.36 | 0.829 | 1.206 |
| EWSR1-ATF1 | PDGFRB | 55 | 0.122 | 1609.8 | 8.291 | 2 | 3.317 | 0.06 | 16.58 |
| | SHC1 | 214 | 0.062 | 25801 | 14.99 | 2 | 7.138 | 0.033 | 29.98 |
| | CRKL | 108 | 0.065 | 8323.4 | 8.844 | 2 | 6.106 | 0.057 | 17.69 |
| | GAB2 | 33 | 0.246 | 389.67 | 9.333 | 2 | 1.768 | 0.054 | 18.67 |
| | NTRK1 | 166 | 0.045 | 22877 | 9.512 | 2 | 8.726 | 0.053 | 19.02 |
| | PLCG1 | 112 | 0.071 | 8176 | 9.805 | 2 | 5.711 | 0.051 | 19.61 |
| | GRB2 | 515 | 0 | 263682 | 26 | 2 | 9.905 | 0.019 | 51.99 |
| | HDAC3 | 23 | 0.134 | 343.97 | 0.509 | 2 | 22.59 | 0.982 | 1.018 |
| | PIN1 | 223 | 0.037 | 40989 | 1.089 | 2 | 102.4 | 0.459 | 2.178 |
| | ETV6 | 33 | 0.079 | 840.83 | 0.603 | 2 | 27.36 | 0.829 | 1.206 |
| EWSR1-FLI1 | BRCA1 | 17 | 0.191 | 187.23 | 0.09 | 2 | 94.44 | 5.556 | 0.18 |
| | POLR2A | 251 | 0.074 | 34919 | 1.239 | 2 | 101.3 | 0.404 | 2.478 |
| | ESR1 | 23 | 0.134 | 343.97 | 0.409 | 2 | 28.12 | 1.222 | 0.818 |
| | EP300 | 453 | 0 | 203852 | 1.996 | 2 | 113.5 | 0.251 | 3.992 |
| | KAT2B | 175 | 0.066 | 18318 | 13.38 | 2 | 6.538 | 0.037 | 26.77 |
| | CREBBP | 296 | 0.058 | 48179 | 1.061 | 2 | 139.5 | 0.471 | 2.122 |
| | EPAS1 | 102 | 0.1 | 6894.1 | 0.202 | 2 | 252.5 | 2.475 | 0.404 |
| | EP300 | 453 | 0 | 203852 | 1.996 | 2 | 113.5 | 0.251 | 3.992 |
| | CREBBP | 296 | 0.058 | 48179 | 1.061 | 2 | 139.5 | 0.471 | 2.122 |
| EWSR1-NR4A3 | TSG101 | 145 | 0.05 | 15318 | 4.062 | 2 | 17.85 | 0.123 | 8.124 |
| | DHX9 | 75 | 0.238 | 2197.3 | 11.06 | 2 | 3.391 | 0.045 | 22.12 |
| | HDAC3 | 23 | 0.134 | 343.97 | 0.509 | 2 | 22.59 | 0.982 | 1.018 |
| | TRIM28 | 223 | 0.064 | 31811 | 16.02 | 2 | 6.961 | 0.031 | 32.04 |
| | FUS | 319 | 0.082 | 50310 | 8 | 2 | 19.94 | 0.063 | 16 |
| | RAD23A | 147 | 0.113 | 13124 | 18.23 | 2 | 4.032 | 0.027 | 36.46 |

| | | | | | | | | | |
|---|---|---|---|---|---|---|---|---|---|
| | JUN | 211 | 0.072 | 25977 | 17.06 | 2 | 6.184 | 0.029 | 34.12 |
| | PRMT1 | 145 | 0.085 | 13214 | 14.08 | 2 | 5.148 | 0.036 | 28.17 |
| | ILK | 211 | 0.046 | 30385 | 0.526 | 2 | 200.6 | 0.951 | 1.052 |
| | BMI1 | 63 | 0.12 | 2352.9 | 12.31 | 2 | 2.558 | 0.041 | 24.62 |
| | ELK1 | 27 | 0.129 | 470.23 | 5.037 | 2 | 2.68 | 0.099 | 10.07 |
| | POLR2A | 251 | 0.074 | 34919 | 1.239 | 2 | 101.3 | 0.404 | 2.478 |
| | CHERP | 57 | 0.16 | 1692.9 | 10.76 | 2 | 2.649 | 0.046 | 21.52 |
| | ATXN3 | 81 | 0.074 | 4620.2 | 7.728 | 2 | 5.241 | 0.065 | 15.46 |
| | EP300 | 453 | 0 | 203852 | 1.996 | 2 | 113.5 | 0.251 | 3.992 |
| | IRF3 | 67 | 0.077 | 3294.7 | 0.396 | 2 | 84.6 | 1.263 | 0.792 |
| | EPAS1 | 102 | 0.1 | 6894.1 | 0.202 | 2 | 252.5 | 2.475 | 0.404 |
| | TP53 | 961 | 0 | 920640 | 1.998 | 2 | 240.5 | 0.25 | 3.996 |
| | ESR1 | 23 | 0.134 | 343.97 | 0.409 | 2 | 28.12 | 1.222 | 0.818 |
| | NONO | 144 | 0.105 | 10404 | 16.86 | 2 | 4.27 | 0.03 | 33.72 |
| | RPA1 | 463 | 0 | 212982 | 1.996 | 2 | 116 | 0.251 | 3.992 |
| | NTRK1 | 166 | 0.045 | 22877 | 9.512 | 2 | 8.726 | 0.053 | 19.02 |
| | RPA2 | 226 | 0.06 | 22557 | 9.512 | 2 | 11.88 | 0.053 | 19.02 |
| | CUL4A | 279 | 0.072 | 34971 | 21.8 | 2 | 6.399 | 0.023 | 43.6 |
| | CUL4B | 279 | 0.072 | 34971 | 16.8 | 2 | 8.304 | 0.03 | 33.6 |
| | FASN | 86 | 0.12 | 3995 | 11.98 | 2 | 3.59 | 0.042 | 23.95 |
| | CREBBP | 296 | 0.058 | 48179 | 1.061 | 2 | 139.5 | 0.471 | 2.122 |
| | HBP1 | 21 | 0.114 | 335.33 | 0.61 | 2 | 17.21 | 0.82 | 1.22 |
| | CUL5 | 279 | 0.072 | 34971 | 1.799 | 2 | 77.54 | 0.278 | 3.598 |
| | HLTF | 46 | 0.123 | 1363.6 | 1.261 | 2 | 18.24 | 0.397 | 2.522 |
| | EWSR1 | 643 | 0 | 411522 | 32 | 2 | 10.05 | 0.016 | 63.99 |
| | YBX1 | 178 | 0.124 | 12926 | 23.77 | 2 | 3.744 | 0.021 | 47.54 |
| | CUL1 | 670 | 0 | 446892 | 12 | 2 | 27.92 | 0.042 | 23.99 |
| | CUL2 | 833 | 0 | 691392 | 2.998 | 2 | 138.9 | 0.167 | 5.996 |
| | CUL3 | 70 | 0.081 | 3676.9 | 10.37 | 2 | 3.375 | 0.048 | 20.74 |
| | HDAC2 | 301 | 0.067 | 40883 | 1.874 | 2 | 80.31 | 0.267 | 3.748 |
| | CREBBP | 296 | 0.058 | 48179 | 1.061 | 2 | 139.5 | 0.471 | 2.122 |
| | ESR1 | 23 | 0.134 | 343.97 | 0.409 | 2 | 28.12 | 1.222 | 0.818 |
| | NONO | 144 | 0.105 | 10404 | 16.86 | 2 | 4.27 | 0.03 | 33.72 |
| | FXR2 | 116 | 0.045 | 10902 | 7.103 | 2 | 8.166 | 0.07 | 14.21 |
| | CUL3 | 70 | 0.081 | 3676.9 | 10.37 | 2 | 3.375 | 0.048 | 20.74 |
| EWSR1-ETV4 | TSG101 | 145 | 0.05 | 15318 | 4.062 | 2 | 17.85 | 0.123 | 8.124 |
| | DHX9 | 75 | 0.238 | 2197.3 | 11.06 | 2 | 3.391 | 0.045 | 22.12 |
| | HDAC3 | 23 | 0.134 | 343.97 | 0.509 | 2 | 22.59 | 0.982 | 1.018 |
| | RFWD2 | 76 | 0.15 | 3044.3 | 12.95 | 2 | 2.935 | 0.039 | 25.89 |
| | FUS | 319 | 0.082 | 50310 | 8 | 2 | 19.94 | 0.063 | 16 |
| | RAD23A | 147 | 0.113 | 13124 | 18.23 | 2 | 4.032 | 0.027 | 36.46 |
| | JUN | 211 | 0.072 | 25977 | 17.06 | 2 | 6.184 | 0.029 | 34.12 |
| | PRMT1 | 145 | 0.085 | 13214 | 14.08 | 2 | 5.148 | 0.036 | 28.17 |
| | ILK | 211 | 0.046 | 30385 | 0.526 | 2 | 200.6 | 0.951 | 1.052 |
| | SMAD2 | 103 | 0.141 | 4685.8 | 2.25 | 2 | 22.89 | 0.222 | 4.5 |
| | BMI1 | 63 | 0.12 | 2352.9 | 12.31 | 2 | 2.558 | 0.041 | 24.62 |
| | ELK1 | 27 | 0.129 | 470.23 | 5.037 | 2 | 2.68 | 0.099 | 10.07 |
| | POLR2A | 251 | 0.074 | 34919 | 1.239 | 2 | 101.3 | 0.404 | 2.478 |

| | | | | | | | | | |
|---|---|---|---|---|---|---|---|---|---|
| | CHERP | 57 | 0.16 | 1692.9 | 10.76 | 2 | 2.649 | 0.046 | 21.52 |
| | ATXN3 | 81 | 0.074 | 4620.2 | 7.728 | 2 | 5.241 | 0.065 | 15.46 |
| | EP300 | 453 | 0 | 203852 | 1.996 | 2 | 113.5 | 0.251 | 3.992 |
| | IRF3 | 67 | 0.077 | 3294.7 | 0.396 | 2 | 84.6 | 1.263 | 0.792 |
| | EPAS1 | 102 | 0.1 | 6894.1 | 0.202 | 2 | 252.5 | 2.475 | 0.404 |
| | TP53 | 961 | 0 | 920640 | 1.998 | 2 | 240.5 | 0.25 | 3.996 |
| | ESR1 | 23 | 0.134 | 343.97 | 0.409 | 2 | 28.12 | 1.222 | 0.818 |
| | NONO | 144 | 0.105 | 10404 | 16.86 | 2 | 4.27 | 0.03 | 33.72 |
| | RPA1 | 463 | 0 | 212982 | 1.996 | 2 | 116 | 0.251 | 3.992 |
| | NTRK1 | 166 | 0.045 | 22877 | 9.512 | 2 | 8.726 | 0.053 | 19.02 |
| | RPA2 | 226 | 0.06 | 22557 | 9.512 | 2 | 11.88 | 0.053 | 19.02 |
| | CUL4A | 279 | 0.072 | 34971 | 21.8 | 2 | 6.399 | 0.023 | 43.6 |
| | CUL4B | 279 | 0.072 | 34971 | 16.8 | 2 | 8.304 | 0.03 | 33.6 |
| | FASN | 86 | 0.12 | 3995 | 11.98 | 2 | 3.59 | 0.042 | 23.95 |
| | CREBBP | 296 | 0.058 | 48179 | 1.061 | 2 | 139.5 | 0.471 | 2.122 |
| | HBP1 | 21 | 0.114 | 335.33 | 0.61 | 2 | 17.21 | 0.82 | 1.22 |
| | CUL5 | 279 | 0.072 | 34971 | 1.799 | 2 | 77.54 | 0.278 | 3.598 |
| | HLTF | 46 | 0.123 | 1363.6 | 1.261 | 2 | 18.24 | 0.397 | 2.522 |
| | EWSR1 | 643 | 0 | 411522 | 32 | 2 | 10.05 | 0.016 | 63.99 |
| | YBX1 | 178 | 0.124 | 12926 | 23.77 | 2 | 3.744 | 0.021 | 47.54 |
| | CUL1 | 670 | 0 | 446892 | 12 | 2 | 27.92 | 0.042 | 23.99 |
| | CUL2 | 833 | 0 | 691392 | 2.998 | 2 | 138.9 | 0.167 | 5.996 |
| | CUL3 | 70 | 0.081 | 3676.9 | 10.37 | 2 | 3.375 | 0.048 | 20.74 |
| | HDAC2 | 301 | 0.067 | 40883 | 1.874 | 2 | 80.31 | 0.267 | 3.748 |
| | CREBBP | 296 | 0.058 | 48179 | 1.061 | 2 | 139.5 | 0.471 | 2.122 |
| | ESR1 | 23 | 0.134 | 343.97 | 0.409 | 2 | 28.12 | 1.222 | 0.818 |
| | NONO | 144 | 0.105 | 10404 | 16.86 | 2 | 4.27 | 0.03 | 33.72 |
| | FXR2 | 116 | 0.045 | 10902 | 7.103 | 2 | 8.166 | 0.07 | 14.21 |
| | CUL3 | 70 | 0.081 | 3676.9 | 10.37 | 2 | 3.375 | 0.048 | 20.74 |
| EWSR1-PATZ1 | TSG101 | 145 | 0.05 | 15318 | 4.062 | 2 | 17.85 | 0.123 | 8.124 |
| | DHX9 | 75 | 0.238 | 2197.3 | 11.06 | 2 | 3.391 | 0.045 | 22.12 |
| | HDAC3 | 23 | 0.134 | 343.97 | 0.509 | 2 | 22.59 | 0.982 | 1.018 |
| | RFWD2 | 76 | 0.15 | 3044.3 | 12.95 | 2 | 2.935 | 0.039 | 25.89 |
| | FUS | 319 | 0.082 | 50310 | 8 | 2 | 19.94 | 0.063 | 16 |
| | RAD23A | 147 | 0.113 | 13124 | 18.23 | 2 | 4.032 | 0.027 | 36.46 |
| | JUN | 211 | 0.072 | 25977 | 17.06 | 2 | 6.184 | 0.029 | 34.12 |
| | PRMT1 | 145 | 0.085 | 13214 | 14.08 | 2 | 5.148 | 0.036 | 28.17 |
| | ILK | 211 | 0.046 | 30385 | 0.526 | 2 | 200.6 | 0.951 | 1.052 |
| | SMAD2 | 103 | 0.141 | 4685.8 | 2.25 | 2 | 22.89 | 0.222 | 4.5 |
| | BMI1 | 63 | 0.12 | 2352.9 | 12.31 | 2 | 2.558 | 0.041 | 24.62 |
| | ELK1 | 27 | 0.129 | 470.23 | 5.037 | 2 | 2.68 | 0.099 | 10.07 |
| | POLR2A | 251 | 0.074 | 34919 | 1.239 | 2 | 101.3 | 0.404 | 2.478 |
| | CHERP | 57 | 0.16 | 1692.9 | 10.76 | 2 | 2.649 | 0.046 | 21.52 |
| | ATXN3 | 81 | 0.074 | 4620.2 | 7.728 | 2 | 5.241 | 0.065 | 15.46 |
| | EP300 | 453 | 0 | 203852 | 1.996 | 2 | 113.5 | 0.251 | 3.992 |
| | IRF3 | 67 | 0.077 | 3294.7 | 0.396 | 2 | 84.6 | 1.263 | 0.792 |
| | EPAS1 | 102 | 0.1 | 6894.1 | 0.202 | 2 | 252.5 | 2.475 | 0.404 |
| | TP53 | 961 | 0 | 920640 | 1.998 | 2 | 240.5 | 0.25 | 3.996 |

| | | | | | | | | | |
|---|---|---|---|---|---|---|---|---|---|
| | ESR1 | 23 | 0.134 | 343.97 | 0.409 | 2 | 28.12 | 1.222 | 0.818 |
| | NONO | 144 | 0.105 | 10404 | 16.86 | 2 | 4.27 | 0.03 | 33.72 |
| | RPA1 | 463 | 0 | 212982 | 1.996 | 2 | 116 | 0.251 | 3.992 |
| | NTRK1 | 166 | 0.045 | 22877 | 9.512 | 2 | 8.726 | 0.053 | 19.02 |
| | RPA2 | 226 | 0.06 | 22557 | 9.512 | 2 | 11.88 | 0.053 | 19.02 |
| | CUL4A | 279 | 0.072 | 34971 | 21.8 | 2 | 6.399 | 0.023 | 43.6 |
| | CUL4B | 279 | 0.072 | 34971 | 16.8 | 2 | 8.304 | 0.03 | 33.6 |
| | FASN | 86 | 0.12 | 3995 | 11.98 | 2 | 3.59 | 0.042 | 23.95 |
| | CREBBP | 296 | 0.058 | 48179 | 1.061 | 2 | 139.5 | 0.471 | 2.122 |
| | HBP1 | 21 | 0.114 | 335.33 | 0.61 | 2 | 17.21 | 0.82 | 1.22 |
| | CUL5 | 279 | 0.072 | 34971 | 1.799 | 2 | 77.54 | 0.278 | 3.598 |
| | HLTF | 46 | 0.123 | 1363.6 | 1.261 | 2 | 18.24 | 0.397 | 2.522 |
| | EWSR1 | 643 | 0 | 411522 | 32 | 2 | 10.05 | 0.016 | 63.99 |
| | YBX1 | 178 | 0.124 | 12926 | 23.77 | 2 | 3.744 | 0.021 | 47.54 |
| | CUL1 | 670 | 0 | 446892 | 12 | 2 | 27.92 | 0.042 | 23.99 |
| | CUL2 | 833 | 0 | 691392 | 2.998 | 2 | 138.9 | 0.167 | 5.996 |
| | CUL3 | 70 | 0.081 | 3676.9 | 10.37 | 2 | 3.375 | 0.048 | 20.74 |
| | HDAC2 | 301 | 0.067 | 40883 | 1.874 | 2 | 80.31 | 0.267 | 3.748 |
| | CREBBP | 296 | 0.058 | 48179 | 1.061 | 2 | 139.5 | 0.471 | 2.122 |
| | ESR1 | 23 | 0.134 | 343.97 | 0.409 | 2 | 28.12 | 1.222 | 0.818 |
| | NONO | 144 | 0.105 | 10404 | 16.86 | 2 | 4.27 | 0.03 | 33.72 |
| | FXR2 | 116 | 0.045 | 10902 | 7.103 | 2 | 8.166 | 0.07 | 14.21 |
| | CUL3 | 70 | 0.081 | 3676.9 | 10.37 | 2 | 3.375 | 0.048 | 20.74 |
| EWSR1-DDIT3 | HDAC1 | 478 | 0 | 212982 | 1.996 | 2 | 119.7 | 0.251 | 3.992 |
| | EPAS1 | 102 | 0.1 | 6894.1 | 0.202 | 2 | 252.5 | 2.475 | 0.404 |
| | HDAC3 | 23 | 0.134 | 343.97 | 0.509 | 2 | 22.59 | 0.982 | 1.018 |
| | DDIT3 | 70 | 0.081 | 3676.9 | 7.371 | 2 | 4.748 | 0.068 | 14.74 |
| | CEBPB | 66 | 0.215 | 1391.3 | 15.55 | 2 | 2.123 | 0.032 | 31.09 |
| | ESR1 | 23 | 0.134 | 343.97 | 0.409 | 2 | 28.12 | 1.222 | 0.818 |
| | FOS | 130 | 0.053 | 12361 | 8.769 | 2 | 7.412 | 0.057 | 17.54 |
| | JUN | 211 | 0.072 | 25977 | 17.06 | 2 | 6.184 | 0.029 | 34.12 |
| | HBP1 | 21 | 0.114 | 335.33 | 0.61 | 2 | 17.21 | 0.82 | 1.22 |
| | POLR2A | 251 | 0.074 | 34919 | 1.239 | 2 | 101.3 | 0.404 | 2.478 |
| | IRF3 | 67 | 0.077 | 3294.7 | 0.396 | 2 | 84.6 | 1.263 | 0.792 |
| | TP53 | 961 | 0 | 920640 | 1.998 | 2 | 240.5 | 0.25 | 3.996 |
| | EP300 | 453 | 0 | 203852 | 1.996 | 2 | 113.5 | 0.251 | 3.992 |
| | EWSR1 | 643 | 0 | 411522 | 32 | 2 | 10.05 | 0.016 | 63.99 |
| | CREBBP | 296 | 0.058 | 48179 | 1.061 | 2 | 139.5 | 0.471 | 2.122 |
| | DHX9 | 75 | 0.238 | 2197.3 | 11.06 | 2 | 3.391 | 0.045 | 22.12 |
| | CUL4A | 279 | 0.072 | 34971 | 21.8 | 2 | 6.399 | 0.023 | 43.6 |
| | CUL4B | 279 | 0.072 | 34971 | 16.8 | 2 | 8.304 | 0.03 | 33.6 |
| | CUL5 | 279 | 0.072 | 34971 | 1.799 | 2 | 77.54 | 0.278 | 3.598 |
| | CUL1 | 670 | 0 | 446892 | 12 | 2 | 27.92 | 0.042 | 23.99 |
| | CUL2 | 833 | 0 | 691392 | 2.998 | 2 | 138.9 | 0.167 | 5.996 |
| | CUL3 | 70 | 0.081 | 3676.9 | 10.37 | 2 | 3.375 | 0.048 | 20.74 |
| EWSR1-POU5F1 | IRF3 | 67 | 0.077 | 3294.7 | 0.396 | 2 | 84.6 | 1.263 | 0.792 |
| | EPAS1 | 102 | 0.1 | 6894.1 | 0.202 | 2 | 252.5 | 2.475 | 0.404 |
| | HDAC3 | 23 | 0.134 | 343.97 | 0.509 | 2 | 22.59 | 0.982 | 1.018 |

| | | | | | | | | | |
|---|---|---|---|---|---|---|---|---|---|
| | TP53 | 961 | 0 | 920640 | 1.998 | 2 | 240.5 | 0.25 | 3.996 |
| | ESR1 | 23 | 0.134 | 343.97 | 0.409 | 2 | 28.12 | 1.222 | 0.818 |
| | JUN | 211 | 0.072 | 25977 | 17.06 | 2 | 6.184 | 0.029 | 34.12 |
| | HBP1 | 21 | 0.114 | 335.33 | 0.61 | 2 | 17.21 | 0.82 | 1.22 |
| | ETS2 | 29 | 0.165 | 553.37 | 6.4 | 2 | 2.266 | 0.078 | 12.8 |
| | CTNNB1 | 286 | 0.057 | 51076 | 18.22 | 2 | 7.85 | 0.027 | 36.43 |
| | POLR2A | 251 | 0.074 | 34919 | 1.239 | 2 | 101.3 | 0.404 | 2.478 |
| | EP300 | 453 | 0 | 203852 | 1.996 | 2 | 113.5 | 0.251 | 3.992 |
| | EWSR1 | 643 | 0 | 411522 | 32 | 2 | 10.05 | 0.016 | 63.99 |
| | CREBBP | 296 | 0.058 | 48179 | 1.061 | 2 | 139.5 | 0.471 | 2.122 |
| | NONO | 144 | 0.105 | 10404 | 16.86 | 2 | 4.27 | 0.03 | 33.72 |
| | CUL2 | 833 | 0 | 691392 | 2.998 | 2 | 138.9 | 0.167 | 5.996 |
| | CUL3 | 70 | 0.081 | 3676.9 | 10.37 | 2 | 3.375 | 0.048 | 20.74 |
| EWSR1-SP3 | HDAC1 | 478 | 0 | 212982 | 1.996 | 2 | 119.7 | 0.251 | 3.992 |
| | EPAS1 | 102 | 0.1 | 6894.1 | 0.202 | 2 | 252.5 | 2.475 | 0.404 |
| | HDAC3 | 23 | 0.134 | 343.97 | 0.509 | 2 | 22.59 | 0.982 | 1.018 |
| | CEBPB | 66 | 0.215 | 1391.3 | 15.55 | 2 | 2.123 | 0.032 | 31.09 |
| | ESR1 | 23 | 0.134 | 343.97 | 0.409 | 2 | 28.12 | 1.222 | 0.818 |
| | JUN | 211 | 0.072 | 25977 | 17.06 | 2 | 6.184 | 0.029 | 34.12 |
| | HBP1 | 21 | 0.114 | 335.33 | 0.61 | 2 | 17.21 | 0.82 | 1.22 |
| | POLR2A | 251 | 0.074 | 34919 | 1.239 | 2 | 101.3 | 0.404 | 2.478 |
| | IRF3 | 67 | 0.077 | 3294.7 | 0.396 | 2 | 84.6 | 1.263 | 0.792 |
| | TP53 | 961 | 0 | 920640 | 1.998 | 2 | 240.5 | 0.25 | 3.996 |
| | RELA | 178 | 0.06 | 22557 | 9.512 | 2 | 9.357 | 0.053 | 19.02 |
| | EP300 | 453 | 0 | 203852 | 1.996 | 2 | 113.5 | 0.251 | 3.992 |
| | EWSR1 | 643 | 0 | 411522 | 32 | 2 | 10.05 | 0.016 | 63.99 |
| | CREBBP | 296 | 0.058 | 48179 | 1.061 | 2 | 139.5 | 0.471 | 2.122 |
| | DHX9 | 75 | 0.238 | 2197.3 | 11.06 | 2 | 3.391 | 0.045 | 22.12 |
| | CUL4A | 279 | 0.072 | 34971 | 21.8 | 2 | 6.399 | 0.023 | 43.6 |
| | CUL4B | 279 | 0.072 | 34971 | 16.8 | 2 | 8.304 | 0.03 | 33.6 |
| | CUL5 | 279 | 0.072 | 34971 | 1.799 | 2 | 77.54 | 0.278 | 3.598 |
| | CUL1 | 670 | 0 | 446892 | 12 | 2 | 27.92 | 0.042 | 23.99 |
| | CUL2 | 833 | 0 | 691392 | 2.998 | 2 | 138.9 | 0.167 | 5.996 |
| | CUL3 | 70 | 0.081 | 3676.9 | 10.37 | 2 | 3.375 | 0.048 | 20.74 |
| FOXO4-CIC | XPO1 | 89 | 0.109 | 4933.6 | 10.37 | 2 | 4.291 | 0.048 | 20.74 |
| | CTNNB1 | 286 | 0.057 | 51076 | 18.22 | 2 | 7.85 | 0.027 | 36.43 |
| | VDR | 89 | 0.109 | 4933.6 | 0.371 | 2 | 119.9 | 1.348 | 0.742 |
| | ESR1 | 23 | 0.134 | 343.97 | 0.409 | 2 | 28.12 | 1.222 | 0.818 |
| | SMAD4 | 103 | 0.141 | 4685.8 | 2.01 | 2 | 25.62 | 0.249 | 4.02 |
| | SMAD3 | 103 | 0.141 | 4685.8 | 26.25 | 2 | 1.962 | 0.019 | 52.5 |
| | SFN | 250 | 0.053 | 43097 | 14.9 | 2 | 8.389 | 0.034 | 29.8 |
| | AKT1 | 60 | 0.16 | 1929 | 14.25 | 2 | 2.106 | 0.035 | 28.49 |
| | FOXO4 | 20 | 0.284 | 165.8 | 0.48 | 2 | 20.83 | 1.042 | 0.96 |
| | MDM2 | 189 | 0.06 | 22877 | 9.512 | 2 | 9.935 | 0.053 | 19.02 |
| | NLK | 56 | 0.067 | 2469.2 | 5.536 | 2 | 5.058 | 0.09 | 11.07 |
| | CREBBP | 296 | 0.058 | 48179 | 1.061 | 2 | 139.5 | 0.471 | 2.122 |
| FUS-ERG | RPA1 | 463 | 0 | 212982 | 1.996 | 2 | 116 | 0.251 | 3.992 |
| | SF3B2 | 26 | 0.142 | 450.27 | 7.333 | 2 | 1.773 | 0.068 | 14.67 |

| | | | | | | | | | |
|---|---|---|---|---|---|---|---|---|---|
| | PRKDC | 234 | 0.084 | 28338 | 21.36 | 2 | 5.478 | 0.023 | 42.72 |
| | PRPF8 | 153 | 0.177 | 7356.1 | 28.5 | 2 | 2.684 | 0.018 | 56.99 |
| | SF3A2 | 26 | 0.142 | 450.27 | 6.333 | 2 | 2.053 | 0.079 | 12.67 |
| | DHX15 | 82 | 0.238 | 2197.3 | 21.06 | 2 | 1.947 | 0.024 | 42.12 |
| | RPA2 | 226 | 0.06 | 22557 | 9.512 | 2 | 11.88 | 0.053 | 19.02 |
| | CUL3 | 70 | 0.081 | 3676.9 | 10.37 | 2 | 3.375 | 0.048 | 20.74 |
| | ABL1 | 173 | 0.076 | 17247 | 14.9 | 2 | 5.805 | 0.034 | 29.8 |
| | PARP1 | 219 | 0.08 | 25770 | 1.233 | 2 | 88.81 | 0.406 | 2.466 |
| | PRKDC | 234 | 0.084 | 28338 | 21.36 | 2 | 5.478 | 0.023 | 42.72 |
| FUS-CREB3L1 | NONO | 144 | 0.105 | 10404 | 16.86 | 2 | 4.27 | 0.03 | 33.72 |
| | CUL4A | 279 | 0.072 | 34971 | 21.8 | 2 | 6.399 | 0.023 | 43.6 |
| | CUL4B | 279 | 0.072 | 34971 | 16.8 | 2 | 8.304 | 0.03 | 33.6 |
| | DHX15 | 82 | 0.238 | 2197.3 | 21.06 | 2 | 1.947 | 0.024 | 42.12 |
| | CUL5 | 279 | 0.072 | 34971 | 1.799 | 2 | 77.54 | 0.278 | 3.598 |
| | CUL1 | 670 | 0 | 446892 | 12 | 2 | 27.92 | 0.042 | 23.99 |
| | CUL2 | 833 | 0 | 691392 | 2.998 | 2 | 138.9 | 0.167 | 5.996 |
| | CUL3 | 70 | 0.081 | 3676.9 | 10.37 | 2 | 3.375 | 0.048 | 20.74 |
| | VCP | 586 | 0 | 341640 | 1.997 | 2 | 146.7 | 0.25 | 3.994 |
| | FBXW11 | 76 | 0.12 | 3995 | 9.977 | 2 | 3.809 | 0.05 | 19.95 |
| | CUL1 | 670 | 0 | 446892 | 12 | 2 | 27.92 | 0.042 | 23.99 |
| FUS-DDIT3 | HDAC1 | 478 | 0 | 212982 | 1.996 | 2 | 119.7 | 0.251 | 3.992 |
| | DDX17 | 142 | 0.185 | 5363.4 | 27.75 | 2 | 2.559 | 0.018 | 55.49 |
| | EPAS1 | 102 | 0.1 | 6894.1 | 0.202 | 2 | 252.5 | 2.475 | 0.404 |
| | RELA | 178 | 0.06 | 22557 | 9.512 | 2 | 9.357 | 0.053 | 19.02 |
| | ESR1 | 23 | 0.134 | 343.97 | 0.409 | 2 | 28.12 | 1.222 | 0.818 |
| | JUN | 211 | 0.072 | 25977 | 17.06 | 2 | 6.184 | 0.029 | 34.12 |
| | TP73 | 112 | 0.072 | 7458.6 | 1.839 | 2 | 30.45 | 0.272 | 3.678 |
| | EWSR1 | 643 | 0 | 411522 | 32 | 2 | 10.05 | 0.016 | 63.99 |
| | CTNNB1 | 286 | 0.057 | 51076 | 18.22 | 2 | 7.85 | 0.027 | 36.43 |
| | DDX5 | 203 | 0.145 | 13444 | 30.97 | 2 | 3.278 | 0.016 | 61.93 |
| | CDK2 | 233 | 0.061 | 32363 | 4.043 | 2 | 28.82 | 0.124 | 8.086 |
| | DDIT3 | 70 | 0.081 | 3676.9 | 7.371 | 2 | 4.748 | 0.068 | 14.74 |
| | MDM2 | 189 | 0.06 | 22877 | 9.512 | 2 | 9.935 | 0.053 | 19.02 |
| | EP300 | 453 | 0 | 203852 | 1.996 | 2 | 113.5 | 0.251 | 3.992 |
| | TRIP4 | 41 | 0.248 | 760.71 | 11.62 | 2 | 1.764 | 0.043 | 23.24 |
| | CREBBP | 296 | 0.058 | 48179 | 1.061 | 2 | 139.5 | 0.471 | 2.122 |
| | VCP | 586 | 0 | 341640 | 1.997 | 2 | 146.7 | 0.25 | 3.994 |
| | FBXW11 | 76 | 0.12 | 3995 | 9.977 | 2 | 3.809 | 0.05 | 19.95 |
| | CUL1 | 670 | 0 | 446892 | 12 | 2 | 27.92 | 0.042 | 23.99 |
| FUS-ATF1 | HDAC1 | 478 | 0 | 212982 | 1.996 | 2 | 119.7 | 0.251 | 3.992 |
| | DDX17 | 142 | 0.185 | 5363.4 | 27.75 | 2 | 2.559 | 0.018 | 55.49 |
| | EPAS1 | 102 | 0.1 | 6894.1 | 0.202 | 2 | 252.5 | 2.475 | 0.404 |
| | RELA | 178 | 0.06 | 22557 | 9.512 | 2 | 9.357 | 0.053 | 19.02 |
| | ESR1 | 23 | 0.134 | 343.97 | 0.409 | 2 | 28.12 | 1.222 | 0.818 |
| | JUN | 211 | 0.072 | 25977 | 17.06 | 2 | 6.184 | 0.029 | 34.12 |
| | TP73 | 112 | 0.072 | 7458.6 | 1.839 | 2 | 30.45 | 0.272 | 3.678 |
| | EWSR1 | 643 | 0 | 411522 | 32 | 2 | 10.05 | 0.016 | 63.99 |
| | CTNNB1 | 286 | 0.057 | 51076 | 18.22 | 2 | 7.85 | 0.027 | 36.43 |

| | | | | | | | | | |
|---|---|---|---|---|---|---|---|---|---|
| | DDX5 | 203 | 0.145 | 13444 | 30.97 | 2 | 3.278 | 0.016 | 61.93 |
| | CDK2 | 233 | 0.061 | 32363 | 4.043 | 2 | 28.82 | 0.124 | 8.086 |
| | DDIT3 | 70 | 0.081 | 3676.9 | 7.371 | 2 | 4.748 | 0.068 | 14.74 |
| | MDM2 | 189 | 0.06 | 22877 | 9.512 | 2 | 9.935 | 0.053 | 19.02 |
| | EP300 | 453 | 0 | 203852 | 1.996 | 2 | 113.5 | 0.251 | 3.992 |
| | TRIP4 | 41 | 0.248 | 760.71 | 11.62 | 2 | 1.764 | 0.043 | 23.24 |
| | CREBBP | 296 | 0.058 | 48179 | 1.061 | 2 | 139.5 | 0.471 | 2.122 |
| | VCP | 586 | 0 | 341640 | 1.997 | 2 | 146.7 | 0.25 | 3.994 |
| | FBXW11 | 76 | 0.12 | 3995 | 9.977 | 2 | 3.809 | 0.05 | 19.95 |
| | CUL1 | 670 | 0 | 446892 | 12 | 2 | 27.92 | 0.042 | 23.99 |
| FUS-CREB3L2 | NONO | 144 | 0.105 | 10404 | 16.86 | 2 | 4.27 | 0.03 | 33.72 |
| | CUL4A | 279 | 0.072 | 34971 | 21.8 | 2 | 6.399 | 0.023 | 43.6 |
| | CUL4B | 279 | 0.072 | 34971 | 16.8 | 2 | 8.304 | 0.03 | 33.6 |
| | DHX15 | 82 | 0.238 | 2197.3 | 21.06 | 2 | 1.947 | 0.024 | 42.12 |
| | CUL5 | 279 | 0.072 | 34971 | 1.799 | 2 | 77.54 | 0.278 | 3.598 |
| | CUL1 | 670 | 0 | 446892 | 12 | 2 | 27.92 | 0.042 | 23.99 |
| | CUL2 | 833 | 0 | 691392 | 2.998 | 2 | 138.9 | 0.167 | 5.996 |
| | CUL3 | 70 | 0.081 | 3676.9 | 10.37 | 2 | 3.375 | 0.048 | 20.74 |
| | VCP | 586 | 0 | 341640 | 1.997 | 2 | 146.7 | 0.25 | 3.994 |
| | FBXW11 | 76 | 0.12 | 3995 | 9.977 | 2 | 3.809 | 0.05 | 19.95 |
| | CUL1 | 670 | 0 | 446892 | 12 | 2 | 27.92 | 0.042 | 23.99 |
| HEY1-NCOA2 | BRCA1 | 17 | 0.191 | 187.23 | 0.09 | 2 | 94.44 | 5.556 | 0.18 |
| | RARA | 108 | 0.084 | 7253.6 | 10.8 | 2 | 5.002 | 0.046 | 21.59 |
| | NR3C1 | 157 | 0.069 | 15985 | 12.61 | 2 | 6.225 | 0.04 | 25.22 |
| | VDR | 89 | 0.109 | 4933.6 | 0.371 | 2 | 119.9 | 1.348 | 0.742 |
| | STAT6 | 61 | 0.05 | 3258.8 | 4.935 | 2 | 6.18 | 0.101 | 9.87 |
| | HNF4A | 64 | 0.144 | 2232.6 | 0.25 | 2 | 128 | 2 | 0.5 |
| | PRMT1 | 145 | 0.085 | 13214 | 14.08 | 2 | 5.148 | 0.036 | 28.17 |
| | CARM1 | 88 | 0.112 | 4522.9 | 11.57 | 2 | 3.802 | 0.043 | 23.15 |
| | RXRA | 109 | 0.094 | 6829.1 | 11.91 | 2 | 4.577 | 0.042 | 23.82 |
| | PPARG | 130 | 0.086 | 10094 | 12.88 | 2 | 5.048 | 0.039 | 25.75 |
| | PPARD | 44 | 0.104 | 1337.1 | 6.227 | 2 | 3.533 | 0.08 | 12.45 |
| | AR | 60 | 0.16 | 1929 | 14.25 | 2 | 2.106 | 0.035 | 28.49 |
| | PPARA | 42 | 0.095 | 1312.2 | 5.767 | 2 | 3.641 | 0.087 | 11.53 |
| | ESR2 | 100 | 0.074 | 6532.6 | 1.2 | 2 | 41.67 | 0.417 | 2.4 |
| | EP300 | 453 | 0 | 203852 | 1.996 | 2 | 113.5 | 0.251 | 3.992 |
| | NCOA2 | 64 | 0.121 | 2191.9 | 9.446 | 2 | 3.388 | 0.053 | 18.89 |
| | NCOA3 | 106 | 0.109 | 5991.8 | 13.36 | 2 | 3.966 | 0.037 | 26.73 |
| | NCOA1 | 101 | 0.12 | 5004.5 | 13.74 | 2 | 3.675 | 0.036 | 27.49 |
| | TP53 | 961 | 0 | 920640 | 1.998 | 2 | 240.5 | 0.25 | 3.996 |
| | ESR1 | 23 | 0.134 | 343.97 | 0.409 | 2 | 28.12 | 1.222 | 0.818 |
| | AHR | 38 | 0.192 | 639.37 | 0.084 | 3 | 150.8 | 3.968 | 0.252 |
| | ARNT | 42 | 0.07 | 1457.6 | 0.094 | 2 | 222.5 | 5.297 | 0.189 |
| | THRB | 48 | 0.117 | 1383.9 | 7.25 | 2 | 3.31 | 0.069 | 14.5 |
| | THRA | 69 | 0.09 | 3265.2 | 7.942 | 2 | 4.344 | 0.063 | 15.88 |
| | NR1I3 | 29 | 0.167 | 577.51 | 6.467 | 2 | 2.242 | 0.077 | 12.93 |
| | NR1I2 | 32 | 0.093 | 794.13 | 4.727 | 2 | 3.385 | 0.106 | 9.454 |
| | PGR | 62 | 0.106 | 2063.4 | 8.226 | 2 | 3.769 | 0.061 | 16.45 |

| | | | | | | | | | |
|---|---|---|---|---|---|---|---|---|---|
| | CREBBP | 296 | 0.058 | 48179 | 1.061 | 2 | 139.5 | 0.471 | 2.122 |
| | PIAS3 | 50 | 0.089 | 1708.7 | 6.235 | 2 | 4.01 | 0.08 | 12.47 |
| | NCOA2 | 64 | 0.121 | 2191.9 | 9.446 | 2 | 3.388 | 0.053 | 18.89 |
| | NCOA1 | 101 | 0.12 | 5004.5 | 13.74 | 2 | 3.675 | 0.036 | 27.49 |
| | UBR5 | 217 | 0.107 | 18784 | 25.04 | 2 | 4.334 | 0.02 | 50.07 |
| IRX2-TERT | YWHAZ | 335 | 0.049 | 63935 | 18.42 | 2 | 9.094 | 0.027 | 36.84 |
| | AKT1 | 60 | 0.16 | 1929 | 14.25 | 2 | 2.106 | 0.035 | 28.49 |
| | RPS6KB1 | 57 | 0.116 | 1844.1 | 8.246 | 2 | 3.456 | 0.061 | 16.49 |
| | ENO1 | 27 | 0.129 | 470.23 | 2.037 | 2 | 6.627 | 0.245 | 4.074 |
| | MTOR | 140 | 0.061 | 13361 | 10.34 | 2 | 6.768 | 0.048 | 20.69 |
| | MDM2 | 189 | 0.06 | 22877 | 9.512 | 2 | 9.935 | 0.053 | 19.02 |
| | TERT | 66 | 0.075 | 3200.9 | 6.746 | 2 | 4.892 | 0.074 | 13.49 |
| | XRCC6 | 264 | 0.064 | 41296 | 0.568 | 2 | 232.4 | 0.88 | 1.136 |
| | TERF1 | 307 | 0.026 | 71236 | 9.746 | 2 | 15.75 | 0.051 | 19.49 |
| | STUB1 | 239 | 0.055 | 35206 | 15 | 2 | 7.969 | 0.033 | 29.99 |
| | TERT | 66 | 0.075 | 3200.9 | 6.746 | 2 | 4.892 | 0.074 | 13.49 |
| | POT1 | 211 | 0.024 | 35382 | 6.882 | 2 | 15.33 | 0.073 | 13.76 |
| | TERT | 66 | 0.075 | 3200.9 | 6.746 | 2 | 4.892 | 0.074 | 13.49 |
| | MTOR | 140 | 0.061 | 13361 | 10.34 | 2 | 6.768 | 0.048 | 20.69 |
| | YWHAQ | 410 | 0 | 167690 | 22 | 2 | 9.32 | 0.023 | 43.99 |
| | RUVBL2 | 223 | 0.085 | 25036 | 20.62 | 2 | 5.408 | 0.024 | 41.24 |
| | TPP1 | 112 | 0.072 | 7458.6 | 11.84 | 2 | 4.73 | 0.042 | 23.68 |
| | TERT | 66 | 0.075 | 3200.9 | 6.746 | 2 | 4.892 | 0.074 | 13.49 |
| | POT1 | 211 | 0.024 | 35382 | 6.882 | 2 | 15.33 | 0.073 | 13.76 |
| JAZF1-SUZ12 | DHX9 | 75 | 0.238 | 2197.3 | 11.06 | 2 | 3.391 | 0.045 | 22.12 |
| | RBM5 | 31 | 0.161 | 600.02 | 6.625 | 2 | 2.34 | 0.075 | 13.25 |
| | FBXW11 | 76 | 0.12 | 3995 | 9.977 | 2 | 3.809 | 0.05 | 19.95 |
| | DDX3X | 146 | 0.144 | 8758.1 | 0.746 | 2 | 97.86 | 0.67 | 1.492 |
| | NXF1 | 61 | 0.312 | 489.74 | 24.52 | 2 | 1.244 | 0.02 | 49.03 |
| | SF3B4 | 100 | 0.128 | 5654 | 14.54 | 2 | 3.44 | 0.034 | 29.07 |
| | PRMT1 | 145 | 0.085 | 13214 | 14.08 | 2 | 5.148 | 0.036 | 28.17 |
| | SF3B1 | 177 | 0.154 | 11161 | 28.85 | 2 | 3.067 | 0.017 | 57.71 |
| | SF3B2 | 26 | 0.142 | 450.27 | 7.333 | 2 | 1.773 | 0.068 | 14.67 |
| | PRPF8 | 153 | 0.177 | 7356.1 | 28.5 | 2 | 2.684 | 0.018 | 56.99 |
| | EED | 66 | 0.322 | 479.74 | 0.167 | 2 | 197.6 | 2.994 | 0.334 |
| | CRNKL1 | 42 | 0.273 | 655.45 | 9.884 | 2 | 2.125 | 0.051 | 19.77 |
| | RNPS1 | 168 | 0.121 | 13587 | 22.04 | 2 | 3.812 | 0.023 | 44.07 |
| | UBE2I | 217 | 0.107 | 18784 | 25.04 | 2 | 4.334 | 0.02 | 50.07 |
| | SNRNP200 | 152 | 0.168 | 8533.7 | 26.96 | 2 | 2.819 | 0.019 | 53.92 |
| | SRSF7 | 108 | 0.234 | 2992.8 | 26.75 | 2 | 2.019 | 0.019 | 53.5 |
| | SNRPD3 | 104 | 0.2 | 3530.6 | 22.38 | 2 | 2.323 | 0.022 | 44.76 |
| | RALY | 76 | 0.228 | 2341.4 | 18.83 | 2 | 2.018 | 0.027 | 37.66 |
| | DDX5 | 203 | 0.145 | 13444 | 30.97 | 2 | 3.278 | 0.016 | 61.93 |
| | PRPF19 | 139 | 0.182 | 6341.7 | 26.81 | 2 | 2.593 | 0.019 | 53.61 |
| | RNF2 | 710 | 0 | 501972 | 1.997 | 2 | 177.8 | 0.25 | 3.994 |
| | SNRPA1 | 134 | 0.15 | 8049.1 | 21.78 | 2 | 3.076 | 0.023 | 43.56 |
| | SON | 44 | 0.174 | 955.71 | 7.289 | 2 | 3.018 | 0.069 | 14.58 |
| | EFTUD2 | 139 | 0.172 | 6970.7 | 25.34 | 2 | 2.743 | 0.02 | 50.68 |

| | | | | | | | | | |
|---|---|---|---|---|---|---|---|---|---|
| | U2AF1 | 217 | 0.107 | 18784 | 25.04 | 2 | 4.334 | 0.02 | 50.07 |
| | SF3A1 | 153 | 0.162 | 8222 | 26.43 | 2 | 2.895 | 0.019 | 52.86 |
| | EPRS | 104 | 0.144 | 4810.1 | 16.67 | 2 | 3.12 | 0.03 | 33.33 |
| | ILF2 | 65 | 0.109 | 2628.6 | 12.82 | 2 | 2.535 | 0.039 | 25.64 |
| | EIF4A3 | 833 | 0 | 691392 | 62 | 2 | 6.718 | 0.008 | 124 |
| | CDC40 | 26 | 0.4 | 140.33 | 11.56 | 2 | 1.125 | 0.043 | 23.11 |
| | ILF3 | 65 | 0.109 | 2628.6 | 18.82 | 2 | 1.727 | 0.027 | 37.64 |
| | RANBP2 | 124 | 0.124 | 7555.6 | 16.97 | 2 | 3.654 | 0.029 | 33.94 |
| | DNMT3B | 68 | 0.123 | 2784.1 | 0.116 | 2 | 293.1 | 4.31 | 0.232 |
| | HDAC1 | 478 | 0 | 212982 | 1.996 | 2 | 119.7 | 0.251 | 3.992 |
| | HDAC2 | 301 | 0.067 | 40883 | 1.874 | 2 | 80.31 | 0.267 | 3.748 |
| | TRIM28 | 223 | 0.064 | 31811 | 16.02 | 2 | 6.961 | 0.031 | 32.04 |
| | CHD4 | 119 | 0.126 | 6244.5 | 16.61 | 2 | 3.583 | 0.03 | 33.21 |
| | UHRF1 | 134 | 0.076 | 11651 | 11.88 | 2 | 5.639 | 0.042 | 23.76 |
| | DNMT1 | 84 | 0.147 | 3254.8 | 13.91 | 2 | 3.02 | 0.036 | 27.81 |
| | MTA1 | 93 | 0.13 | 4604.9 | 13.81 | 2 | 3.367 | 0.036 | 27.62 |
| | NR2C2 | 13 | 0.682 | 7.021 | 6.78 | 2 | 0.959 | 0.074 | 13.56 |
| | EZH2 | 275 | 0.045 | 49579 | 1.233 | 2 | 111.5 | 0.406 | 2.466 |
| | GATAD2B | 20 | 0.246 | 389.67 | 9.333 | 2 | 1.071 | 0.054 | 18.67 |
| | EED | 66 | 0.322 | 479.74 | 0.167 | 2 | 197.6 | 2.994 | 0.334 |
| | CBX5 | 161 | 0.04 | 20734 | 1.323 | 2 | 60.85 | 0.378 | 2.646 |
| | RBBP4 | 170 | 0.106 | 12864 | 19.75 | 2 | 4.303 | 0.025 | 39.51 |
| | CBX3 | 101 | 0.093 | 6093.6 | 11.07 | 2 | 4.562 | 0.045 | 22.14 |
| | SETDB1 | 126 | 0.036 | 13488 | 6.441 | 2 | 9.781 | 0.078 | 12.88 |
| | VCP | 586 | 0 | 341640 | 1.997 | 2 | 146.7 | 0.25 | 3.994 |
| | BRCA1 | 17 | 0.191 | 187.23 | 0.09 | 2 | 94.44 | 5.556 | 0.18 |
| | MTOR | 140 | 0.061 | 13361 | 10.34 | 2 | 6.768 | 0.048 | 20.69 |
| | NXF1 | 61 | 0.312 | 489.74 | 24.52 | 2 | 1.244 | 0.02 | 49.03 |
| | RUVBL2 | 223 | 0.085 | 25036 | 20.62 | 2 | 5.408 | 0.024 | 41.24 |
| | VCP | 586 | 0 | 341640 | 1.997 | 2 | 146.7 | 0.25 | 3.994 |
| | FBXW11 | 76 | 0.12 | 3995 | 9.977 | 2 | 3.809 | 0.05 | 19.95 |
| | SKP1 | 185 | 0.059 | 20423 | 12.74 | 2 | 7.259 | 0.039 | 25.48 |
| | BTRC | 42 | 0.211 | 765.63 | 12.19 | 2 | 1.723 | 0.041 | 24.38 |
| | EZH2 | 275 | 0.045 | 49579 | 1.233 | 2 | 111.5 | 0.406 | 2.466 |
| | DHX9 | 75 | 0.238 | 2197.3 | 11.06 | 2 | 3.391 | 0.045 | 22.12 |
| | ADAR | 60 | 0.16 | 1929 | 11.25 | 2 | 2.668 | 0.044 | 22.49 |
| | EZH2 | 275 | 0.045 | 49579 | 1.233 | 2 | 111.5 | 0.406 | 2.466 |
| | FBXW11 | 76 | 0.12 | 3995 | 9.977 | 2 | 3.809 | 0.05 | 19.95 |
| | HDAC2 | 301 | 0.067 | 40883 | 1.874 | 2 | 80.31 | 0.267 | 3.748 |
| | NXF1 | 61 | 0.312 | 489.74 | 24.52 | 2 | 1.244 | 0.02 | 49.03 |
| | JARID2 | 58 | 0.162 | 1617 | 21.05 | 2 | 1.378 | 0.024 | 42.1 |
| | SETDB1 | 126 | 0.036 | 13488 | 6.441 | 2 | 9.781 | 0.078 | 12.88 |
| | RELA | 178 | 0.06 | 22557 | 9.512 | 2 | 9.357 | 0.053 | 19.02 |
| | FBXW11 | 76 | 0.12 | 3995 | 9.977 | 2 | 3.809 | 0.05 | 19.95 |
| | BTRC | 42 | 0.211 | 765.63 | 12.19 | 2 | 1.723 | 0.041 | 24.38 |
| | BTRC | 42 | 0.211 | 765.63 | 12.19 | 2 | 1.723 | 0.041 | 24.38 |
| | CSNK2B | 42 | 0.273 | 655.45 | 6.884 | 2 | 3.051 | 0.073 | 13.77 |
| | NXF1 | 61 | 0.312 | 489.74 | 24.52 | 2 | 1.244 | 0.02 | 49.03 |

| | | | | | | | | | |
|---|---|---|---|---|---|---|---|---|---|
| JAZF1-PHF1 | DHX9 | 75 | 0.238 | 2197.3 | 11.06 | 2 | 3.391 | 0.045 | 22.12 |
| | EZH1 | 9 | 0.571 | 9.333 | 0.033 | 2 | 135.1 | 15.02 | 0.067 |
| | PPARG | 130 | 0.086 | 10094 | 12.88 | 2 | 5.048 | 0.039 | 25.75 |
| | EZH2 | 275 | 0.045 | 49579 | 1.233 | 2 | 111.5 | 0.406 | 2.466 |
| | EED | 66 | 0.322 | 479.74 | 0.167 | 2 | 197.6 | 2.994 | 0.334 |
| | XRCC6 | 264 | 0.064 | 41296 | 0.568 | 2 | 232.4 | 0.88 | 1.136 |
| | XRCC5 | 264 | 0.064 | 41296 | 0.868 | 2 | 152.1 | 0.576 | 1.736 |
| | PHF1 | 23 | 0.194 | 380.67 | 6 | 2 | 1.917 | 0.083 | 12 |
| | HDAC1 | 478 | 0 | 212982 | 1.996 | 2 | 119.7 | 0.251 | 3.992 |
| | PPARG | 130 | 0.086 | 10094 | 12.88 | 2 | 5.048 | 0.039 | 25.75 |
| | EZH2 | 275 | 0.045 | 49579 | 1.233 | 2 | 111.5 | 0.406 | 2.466 |
| | PHF1 | 23 | 0.194 | 380.67 | 6 | 2 | 1.917 | 0.083 | 12 |
| | PHF1 | 23 | 0.194 | 380.67 | 6 | 2 | 1.917 | 0.083 | 12 |
| | TP53 | 961 | 0 | 920640 | 1.998 | 2 | 240.5 | 0.25 | 3.996 |
| | XRCC6 | 264 | 0.064 | 41296 | 0.568 | 2 | 232.4 | 0.88 | 1.136 |
| MEAF6-TRERF1 | HDAC1 | 478 | 0 | 212982 | 1.996 | 2 | 119.7 | 0.251 | 3.992 |
| | TRERF1 | 11 | 0.255 | 62.833 | 4.167 | 2 | 1.32 | 0.12 | 8.334 |
| | KAT5 | 210 | 0.054 | 31401 | 3.238 | 2 | 32.43 | 0.154 | 6.476 |
| | ING3 | 58 | 0.162 | 1617 | 1.051 | 2 | 27.59 | 0.476 | 2.102 |
| | CREBBP | 296 | 0.058 | 48179 | 1.061 | 2 | 139.5 | 0.471 | 2.122 |
| | YEATS4 | 99 | 0.061 | 8123.4 | 7.92 | 2 | 6.25 | 0.063 | 15.84 |
| | EP300 | 453 | 0 | 203852 | 1.996 | 2 | 113.5 | 0.251 | 3.992 |
| | MORF4L1 | 106 | 0.087 | 6884 | 10.98 | 2 | 4.827 | 0.046 | 21.96 |
| | HIST1H2BA | 90 | 0.093 | 5635.8 | 10.13 | 2 | 4.442 | 0.049 | 20.26 |
| | ELAVL1 | 213 | 0.076 | 28937 | 18.66 | 2 | 5.706 | 0.027 | 37.33 |
| | TRERF1 | 11 | 0.255 | 62.833 | 4.167 | 2 | 1.32 | 0.12 | 8.334 |
| | KAT6A | 37 | 0.116 | 925.49 | 0.06 | 2 | 308.3 | 8.333 | 0.12 |
| | CREBBP | 296 | 0.058 | 48179 | 1.061 | 2 | 139.5 | 0.471 | 2.122 |
| | NR5A1 | 26 | 0.12 | 476 | 4.815 | 2 | 2.7 | 0.104 | 9.63 |
| | SOX2 | 346 | 0.02 | 104377 | 8.832 | 2 | 19.59 | 0.057 | 17.66 |
| | HDAC1 | 478 | 0 | 212982 | 1.996 | 2 | 119.7 | 0.251 | 3.992 |
| | TRERF1 | 11 | 0.255 | 62.833 | 4.167 | 2 | 1.32 | 0.12 | 8.334 |
| MEAF6-PHF1 | DHX9 | 75 | 0.238 | 2197.3 | 11.06 | 2 | 3.391 | 0.045 | 22.12 |
| | EZH1 | 9 | 0.571 | 9.333 | 0.033 | 2 | 135.1 | 15.02 | 0.067 |
| | EZH2 | 275 | 0.045 | 49579 | 1.233 | 2 | 111.5 | 0.406 | 2.466 |
| | EED | 66 | 0.322 | 479.74 | 0.167 | 2 | 197.6 | 2.994 | 0.334 |
| | XRCC6 | 264 | 0.064 | 41296 | 0.568 | 2 | 232.4 | 0.88 | 1.136 |
| | XRCC5 | 264 | 0.064 | 41296 | 0.868 | 2 | 152.1 | 0.576 | 1.736 |
| | PHF1 | 23 | 0.194 | 380.67 | 6 | 2 | 1.917 | 0.083 | 12 |
| | PHF1 | 23 | 0.194 | 380.67 | 6 | 2 | 1.917 | 0.083 | 12 |
| | TP53 | 961 | 0 | 920640 | 1.998 | 2 | 240.5 | 0.25 | 3.996 |
| | XRCC6 | 264 | 0.064 | 41296 | 0.568 | 2 | 232.4 | 0.88 | 1.136 |
| | KAT6A | 37 | 0.116 | 925.49 | 0.06 | 2 | 308.3 | 8.333 | 0.12 |
| | TP53 | 961 | 0 | 920640 | 1.998 | 2 | 240.5 | 0.25 | 3.996 |
| | ELAVL1 | 213 | 0.076 | 28937 | 18.66 | 2 | 5.706 | 0.027 | 37.33 |
| | HDAC1 | 478 | 0 | 212982 | 1.996 | 2 | 119.7 | 0.251 | 3.992 |
| | EZH2 | 275 | 0.045 | 49579 | 1.233 | 2 | 111.5 | 0.406 | 2.466 |
| | PHF1 | 23 | 0.194 | 380.67 | 6 | 2 | 1.917 | 0.083 | 12 |

| | | | | | | | | | |
|---|---|---|---|---|---|---|---|---|---|
| NR4A3-TAF15 | TRIM28 | 223 | 0.064 | 31811 | 16.02 | 2 | 6.961 | 0.031 | 32.04 |
| | FUS | 319 | 0.082 | 50310 | 8 | 2 | 19.94 | 0.063 | 16 |
| | PRMT1 | 145 | 0.085 | 13214 | 14.08 | 2 | 5.148 | 0.036 | 28.17 |
| | COPS6 | 795 | 0 | 629642 | 1.997 | 2 | 199 | 0.25 | 3.994 |
| | COPS5 | 795 | 0 | 629642 | 1.997 | 2 | 199 | 0.25 | 3.994 |
| | POLR2C | 101 | 0.131 | 4996.6 | 14.85 | 2 | 3.4 | 0.034 | 29.7 |
| | POLR2A | 251 | 0.074 | 34919 | 1.239 | 2 | 101.3 | 0.404 | 2.478 |
| | TAF15 | 58 | 0.178 | 1581.8 | 11.97 | 2 | 2.424 | 0.042 | 23.93 |
| | SF1 | 123 | 0.084 | 9821.5 | 12.08 | 2 | 5.091 | 0.041 | 24.16 |
| | NEDD8 | 257 | 0.089 | 25031 | 24.58 | 2 | 5.227 | 0.02 | 49.17 |
| | POLR2E | 103 | 0.155 | 4501.4 | 17.46 | 2 | 2.95 | 0.029 | 34.91 |
| | RPA1 | 463 | 0 | 212982 | 1.996 | 2 | 116 | 0.251 | 3.992 |
| | RPA2 | 226 | 0.06 | 22557 | 9.512 | 2 | 11.88 | 0.053 | 19.02 |
| | CUL4A | 279 | 0.072 | 34971 | 21.8 | 2 | 6.399 | 0.023 | 43.6 |
| | CUL4B | 279 | 0.072 | 34971 | 16.8 | 2 | 8.304 | 0.03 | 33.6 |
| | CUL5 | 279 | 0.072 | 34971 | 1.799 | 2 | 77.54 | 0.278 | 3.598 |
| | CUL1 | 670 | 0 | 446892 | 12 | 2 | 27.92 | 0.042 | 23.99 |
| | CUL2 | 833 | 0 | 691392 | 2.998 | 2 | 138.9 | 0.167 | 5.996 |
| | CUL3 | 70 | 0.081 | 3676.9 | 10.37 | 2 | 3.375 | 0.048 | 20.74 |
| | EZH2 | 275 | 0.045 | 49579 | 1.233 | 2 | 111.5 | 0.406 | 2.466 |
| | TRIM28 | 223 | 0.064 | 31811 | 16.02 | 2 | 6.961 | 0.031 | 32.04 |
| | CUL1 | 670 | 0 | 446892 | 12 | 2 | 27.92 | 0.042 | 23.99 |
| NR4A3-TFG | CUL4A | 279 | 0.072 | 34971 | 21.8 | 2 | 6.399 | 0.023 | 43.6 |
| | CUL4B | 279 | 0.072 | 34971 | 16.8 | 2 | 8.304 | 0.03 | 33.6 |
| | CUL5 | 279 | 0.072 | 34971 | 1.799 | 2 | 77.54 | 0.278 | 3.598 |
| | CUL1 | 670 | 0 | 446892 | 12 | 2 | 27.92 | 0.042 | 23.99 |
| | CUL2 | 833 | 0 | 691392 | 2.998 | 2 | 138.9 | 0.167 | 5.996 |
| | CUL3 | 70 | 0.081 | 3676.9 | 10.37 | 2 | 3.375 | 0.048 | 20.74 |
| | TRIM28 | 223 | 0.064 | 31811 | 16.02 | 2 | 6.961 | 0.031 | 32.04 |
| | CUL1 | 670 | 0 | 446892 | 12 | 2 | 27.92 | 0.042 | 23.99 |
| | CUL3 | 70 | 0.081 | 3676.9 | 10.37 | 2 | 3.375 | 0.048 | 20.74 |
| NUP107-LGR5 | NUP153 | 105 | 0.12 | 5271.1 | 14.19 | 2 | 3.7 | 0.035 | 28.38 |
| | KPNB1 | 56 | 0.072 | 26556 | 6.9 | 2 | 4.058 | 0.072 | 13.8 |
| | NTRK1 | 166 | 0.045 | 22877 | 9.512 | 2 | 8.726 | 0.053 | 19.02 |
| | CUL3 | 70 | 0.081 | 3676.9 | 10.37 | 2 | 3.375 | 0.048 | 20.74 |
| | NUP153 | 105 | 0.12 | 5271.1 | 14.19 | 2 | 3.7 | 0.035 | 28.38 |
| | EIF4B | 77 | 0.115 | 3123.9 | 10.47 | 2 | 3.678 | 0.048 | 20.94 |
| | CUL3 | 70 | 0.081 | 3676.9 | 10.37 | 2 | 3.375 | 0.048 | 20.74 |
| | EED | 66 | 0.322 | 479.74 | 0.167 | 2 | 197.6 | 2.994 | 0.334 |
| | KPNB1 | 56 | 0.072 | 26556 | 6.9 | 2 | 4.058 | 0.072 | 13.8 |
| | TP53BP1 | 961 | 0 | 920640 | 1.998 | 2 | 240.5 | 0.25 | 3.996 |
| PAPPA-NUP107 | NUP153 | 105 | 0.12 | 5271.1 | 14.19 | 2 | 3.7 | 0.035 | 28.38 |
| | ELAVL1 | 213 | 0.076 | 28937 | 18.66 | 2 | 5.706 | 0.027 | 37.33 |
| | KPNB1 | 56 | 0.072 | 26556 | 6.9 | 2 | 4.058 | 0.072 | 13.8 |
| | NTRK1 | 166 | 0.045 | 22877 | 9.512 | 2 | 8.726 | 0.053 | 19.02 |
| | SMAD3 | 103 | 0.141 | 4685.8 | 26.25 | 2 | 1.962 | 0.019 | 52.5 |
| | VCP | 586 | 0 | 341640 | 1.997 | 2 | 146.7 | 0.25 | 3.994 |
| | CUL3 | 70 | 0.081 | 3676.9 | 10.37 | 2 | 3.375 | 0.048 | 20.74 |

| | | | | | | | | | |
|---|---|---|---|---|---|---|---|---|---|
| | EIF4B | 77 | 0.115 | 3123.9 | 10.47 | 2 | 3.678 | 0.048 | 20.94 |
| | SMAD9 | 111 | 0.019 | 11160 | 4 | 2 | 13.88 | 0.125 | 8 |
| | SKIL | 95 | 0.044 | 7250.5 | 6.063 | 2 | 7.834 | 0.082 | 12.13 |
| | PAPPA | 10 | 0.244 | 53 | 3.818 | 2 | 1.31 | 0.131 | 7.636 |
| | SMAD2 | 103 | 0.141 | 4685.8 | 2.25 | 2 | 22.89 | 0.222 | 4.5 |
| | SMAD3 | 103 | 0.141 | 4685.8 | 26.25 | 2 | 1.962 | 0.019 | 52.5 |
| | TP53BP1 | 961 | 0 | 920640 | 1.998 | 2 | 240.5 | 0.25 | 3.996 |
| | ELAVL1 | 213 | 0.076 | 28937 | 18.66 | 2 | 5.706 | 0.027 | 37.33 |
| | EED | 66 | 0.322 | 479.74 | 0.167 | 2 | 197.6 | 2.994 | 0.334 |
| | KPNB1 | 56 | 0.072 | 26556 | 6.9 | 2 | 4.058 | 0.072 | 13.8 |
| | NUP214 | 59 | 0.157 | 1780.2 | 10.75 | 2 | 2.745 | 0.047 | 21.49 |
| | SMAD2 | 103 | 0.141 | 4685.8 | 2.25 | 2 | 22.89 | 0.222 | 4.5 |
| | SMAD3 | 103 | 0.141 | 4685.8 | 26.25 | 2 | 1.962 | 0.019 | 52.5 |
| PAX3-FOXO1 | NCOA1 | 101 | 0.12 | 5004.5 | 13.74 | 2 | 3.675 | 0.036 | 27.49 |
| | ESR1 | 23 | 0.134 | 343.97 | 0.409 | 2 | 28.12 | 1.222 | 0.818 |
| | PARP1 | 219 | 0.08 | 25770 | 1.233 | 2 | 88.81 | 0.406 | 2.466 |
| | AR | 60 | 0.16 | 1929 | 14.25 | 2 | 2.106 | 0.035 | 28.49 |
| | EP300 | 453 | 0 | 203852 | 1.996 | 2 | 113.5 | 0.251 | 3.992 |
| | CREBBP | 296 | 0.058 | 48179 | 1.061 | 2 | 139.5 | 0.471 | 2.122 |
| | TRIM28 | 223 | 0.064 | 31811 | 16.02 | 2 | 6.961 | 0.031 | 32.04 |
| | PARP1 | 219 | 0.08 | 25770 | 1.233 | 2 | 88.81 | 0.406 | 2.466 |
| | CREBBP | 296 | 0.058 | 48179 | 1.061 | 2 | 139.5 | 0.471 | 2.122 |
| PAX7-FOXO1 | RARA | 108 | 0.084 | 7253.6 | 10.8 | 2 | 5.002 | 0.046 | 21.59 |
| | NCOA1 | 101 | 0.12 | 5004.5 | 13.74 | 2 | 3.675 | 0.036 | 27.49 |
| | MYOD1 | 68 | 0.111 | 2755.6 | 9.206 | 2 | 3.693 | 0.054 | 18.41 |
| | ESR1 | 23 | 0.134 | 343.97 | 0.409 | 2 | 28.12 | 1.222 | 0.818 |
| | HNF4A | 64 | 0.144 | 2232.6 | 0.25 | 2 | 128 | 2 | 0.5 |
| | PARP1 | 219 | 0.08 | 25770 | 1.233 | 2 | 88.81 | 0.406 | 2.466 |
| | SMAD3 | 103 | 0.141 | 4685.8 | 26.25 | 2 | 1.962 | 0.019 | 52.5 |
| | FOXO1 | 54 | 0.137 | 1666.9 | 0.963 | 2 | 28.04 | 0.519 | 1.926 |
| | AR | 60 | 0.16 | 1929 | 14.25 | 2 | 2.106 | 0.035 | 28.49 |
| | MDM2 | 189 | 0.06 | 22877 | 9.512 | 2 | 9.935 | 0.053 | 19.02 |
| | EP300 | 453 | 0 | 203852 | 1.996 | 2 | 113.5 | 0.251 | 3.992 |
| | CREBBP | 296 | 0.058 | 48179 | 1.061 | 2 | 139.5 | 0.471 | 2.122 |
| | AKT1 | 60 | 0.16 | 1929 | 14.25 | 2 | 2.106 | 0.035 | 28.49 |
| | EP300 | 453 | 0 | 203852 | 1.996 | 2 | 113.5 | 0.251 | 3.992 |
| | CREBBP | 296 | 0.058 | 48179 | 1.061 | 2 | 139.5 | 0.471 | 2.122 |
| SS18-SSX2 | DPF2 | 45 | 0.204 | 1113.3 | 10.74 | 2 | 2.095 | 0.047 | 21.48 |
| | SMARCC2 | 129 | 0.129 | 6769.4 | 0.623 | 2 | 103.5 | 0.803 | 1.246 |
| | SMARCC1 | 116 | 0.148 | 4333.8 | 0.559 | 2 | 103.8 | 0.894 | 1.118 |
| | PHF10 | 23 | 0.194 | 380.67 | 6 | 2 | 1.917 | 0.083 | 12 |
| | ELAVL1 | 213 | 0.076 | 28937 | 18.66 | 2 | 5.706 | 0.027 | 37.33 |
| | DPF3 | 25 | 0.487 | 162.74 | 13.15 | 2 | 0.95 | 0.038 | 26.31 |
| | ARID2 | 24 | 0.293 | 307.74 | 0.4 | 2 | 30 | 1.25 | 0.8 |
| | DPF1 | 9 | 0.639 | 19 | 6.4 | 2 | 0.703 | 0.078 | 12.8 |
| | SMARCD3 | 40 | 0.215 | 786.57 | 10.15 | 2 | 1.971 | 0.049 | 20.29 |
| | SMARCE1 | 118 | 0.091 | 8801.9 | 12.54 | 2 | 4.706 | 0.04 | 25.08 |
| | SMARCD1 | 107 | 0.084 | 7929.3 | 10.8 | 2 | 4.956 | 0.046 | 21.59 |

| | | | | | | | | | |
|---|---|---|---|---|---|---|---|---|---|
| | EED | 66 | 0.322 | 479.74 | 0.167 | 2 | 197.6 | 2.994 | 0.334 |
| | SMARCA2 | 103 | 0.141 | 4685.8 | 16.25 | 2 | 3.169 | 0.031 | 32.5 |
| | EP300 | 453 | 0 | 203852 | 1.996 | 2 | 113.5 | 0.251 | 3.992 |
| | SMARCA4 | 200 | 0.088 | 20379 | 3.512 | 2 | 28.47 | 0.142 | 7.024 |
| | HDAC1 | 478 | 0 | 212982 | 1.996 | 2 | 119.7 | 0.251 | 3.992 |
| | ARID1B | 37 | 0.263 | 682.8 | 11.16 | 2 | 1.658 | 0.045 | 22.32 |
| | ARID1A | 55 | 0.249 | 885.76 | 15.18 | 2 | 1.812 | 0.033 | 30.36 |
| | ACTL6A | 103 | 0.125 | 6167.6 | 14.58 | 2 | 3.533 | 0.034 | 29.15 |
| | HDAC2 | 301 | 0.067 | 40883 | 1.874 | 2 | 80.31 | 0.267 | 3.748 |
| | CUL3 | 70 | 0.081 | 3676.9 | 10.37 | 2 | 3.375 | 0.048 | 20.74 |
| | RNF2 | 710 | 0 | 501972 | 1.997 | 2 | 177.8 | 0.25 | 3.994 |
| | SMARCD2 | 46 | 0.225 | 972.23 | 11.87 | 2 | 1.937 | 0.042 | 23.74 |
| | GRB2 | 515 | 0 | 263682 | 26 | 2 | 9.905 | 0.019 | 51.99 |
| | YWHAG | 272 | 0.056 | 37593 | 16.99 | 2 | 8.003 | 0.029 | 33.99 |
| | CUL3 | 70 | 0.081 | 3676.9 | 10.37 | 2 | 3.375 | 0.048 | 20.74 |
| SS18-SSX1 | SMARCC2 | 129 | 0.129 | 6769.4 | 0.623 | 2 | 103.5 | 0.803 | 1.246 |
| | DPF2 | 45 | 0.204 | 1113.3 | 10.74 | 2 | 2.095 | 0.047 | 21.48 |
| | SMARCC1 | 116 | 0.148 | 4333.8 | 0.559 | 2 | 103.8 | 0.894 | 1.118 |
| | PHF10 | 23 | 0.194 | 380.67 | 6 | 2 | 5.155 | 0.224 | 4.462 |
| | ELAVL1 | 213 | 0.076 | 28937 | 18.66 | 2 | 5.706 | 0.027 | 37.33 |
| | DPF3 | 25 | 0.487 | 162.74 | 13.15 | 2 | 0.95 | 0.038 | 26.31 |
| | DPF1 | 9 | 0.639 | 19 | 6.4 | 2 | 0.703 | 0.078 | 12.8 |
| | ARID2 | 24 | 0.293 | 307.74 | 0.4 | 2 | 30 | 1.25 | 0.8 |
| | HDAC2 | 301 | 0.067 | 40883 | 1.874 | 2 | 80.31 | 0.267 | 3.748 |
| | SMARCD3 | 40 | 0.215 | 786.57 | 10.15 | 2 | 1.971 | 0.049 | 20.29 |
| | SMARCE1 | 118 | 0.091 | 8801.9 | 12.54 | 2 | 4.706 | 0.04 | 25.08 |
| | SMARCD1 | 107 | 0.084 | 7929.3 | 10.8 | 2 | 4.956 | 0.046 | 21.59 |
| | EED | 66 | 0.322 | 479.74 | 0.167 | 2 | 197.6 | 2.994 | 0.334 |
| | SMARCA2 | 103 | 0.141 | 4685.8 | 16.25 | 2 | 3.169 | 0.031 | 32.5 |
| | EP300 | 453 | 0 | 203852 | 1.996 | 2 | 113.5 | 0.251 | 3.992 |
| | SMARCA4 | 200 | 0.088 | 20379 | 3.512 | 2 | 28.47 | 0.142 | 7.024 |
| | HDAC1 | 478 | 0 | 212982 | 1.996 | 2 | 119.7 | 0.251 | 3.992 |
| | ARID1B | 37 | 0.263 | 682.8 | 11.16 | 2 | 1.658 | 0.045 | 22.32 |
| | ARID1A | 55 | 0.249 | 885.76 | 15.18 | 2 | 1.812 | 0.033 | 30.36 |
| | ACTL6A | 103 | 0.125 | 6167.6 | 14.58 | 2 | 3.533 | 0.034 | 29.15 |
| | CUL3 | 70 | 0.081 | 3676.9 | 10.37 | 2 | 3.375 | 0.048 | 20.74 |
| | SMARCD2 | 46 | 0.225 | 972.23 | 11.87 | 2 | 1.937 | 0.042 | 23.74 |
| | GRB2 | 515 | 0 | 263682 | 26 | 2 | 9.905 | 0.019 | 51.99 |
| | YWHAG | 272 | 0.056 | 37593 | 16.99 | 2 | 8.003 | 0.029 | 33.99 |
| | CUL3 | 70 | 0.081 | 3676.9 | 10.37 | 2 | 3.375 | 0.048 | 20.74 |
| SS18L1-SSX1 | SMARCC1 | 116 | 0.148 | 4333.8 | 0.559 | 2 | 103.8 | 0.894 | 1.118 |
| | STAT3 | 223 | 0.044 | 37463 | 11.64 | 2 | 9.578 | 0.043 | 23.28 |
| | BMI1 | 63 | 0.12 | 2352.9 | 12.31 | 2 | 2.558 | 0.041 | 24.62 |
| | WHSC1L1 | 89 | 0.109 | 4933.6 | 0.371 | 2 | 119.9 | 1.348 | 0.742 |
| | SMAD3 | 103 | 0.141 | 4685.8 | 26.25 | 2 | 1.962 | 0.019 | 52.5 |
| | HDAC2 | 301 | 0.067 | 40883 | 1.874 | 2 | 80.31 | 0.267 | 3.748 |
| | SMAD1 | 192 | 0.072 | 20856 | 11.68 | 2 | 8.22 | 0.043 | 23.36 |
| | EP300 | 453 | 0 | 203852 | 1.996 | 2 | 113.5 | 0.251 | 3.992 |

| | | | | | | | | | |
|---|---|---|---|---|---|---|---|---|---|
| | SMARCA4 | 200 | 0.088 | 20379 | 3.512 | 2 | 28.47 | 0.142 | 7.024 |
| | CREBBP | 296 | 0.058 | 48179 | 1.061 | 2 | 139.5 | 0.471 | 2.122 |
| | DPF2 | 45 | 0.204 | 1113.3 | 10.74 | 2 | 2.095 | 0.047 | 21.48 |
| | SMARCC1 | 116 | 0.148 | 4333.8 | 0.559 | 2 | 103.8 | 0.894 | 1.118 |
| | SMARCE1 | 118 | 0.091 | 8801.9 | 12.54 | 2 | 4.706 | 0.04 | 25.08 |
| | SMARCA4 | 200 | 0.088 | 20379 | 3.512 | 2 | 28.47 | 0.142 | 7.024 |
| | CUL3 | 70 | 0.081 | 3676.9 | 10.37 | 2 | 3.375 | 0.048 | 20.74 |
| TGFBR3-MGEA5 | MAST1 | 39 | 0.206 | 588.77 | 9.6 | 2 | 2.031 | 0.052 | 19.2 |
| | RNF32 | 42 | 0.206 | 715.69 | 10 | 2 | 2.1 | 0.05 | 20 |
| | PAXIP1 | 257 | 0.03 | 53566 | 9.674 | 2 | 13.28 | 0.052 | 19.35 |
| | CBX8 | 128 | 0.071 | 11127 | 10.89 | 2 | 5.876 | 0.046 | 21.78 |
| | CSNK2B | 42 | 0.273 | 655.45 | 6.884 | 2 | 3.051 | 0.073 | 13.77 |
| | PAXIP1 | 257 | 0.03 | 53566 | 9.674 | 2 | 13.28 | 0.052 | 19.35 |
| TRIO-TERT | YWHAZ | 335 | 0.049 | 63935 | 18.42 | 2 | 9.094 | 0.027 | 36.84 |
| | AKT1 | 60 | 0.16 | 1929 | 14.25 | 2 | 2.106 | 0.035 | 28.49 |
| | RPS6KB1 | 57 | 0.116 | 1844.1 | 8.246 | 2 | 3.456 | 0.061 | 16.49 |
| | ENO1 | 27 | 0.129 | 470.23 | 2.037 | 2 | 6.627 | 0.245 | 4.074 |
| | MTOR | 140 | 0.061 | 13361 | 10.34 | 2 | 6.768 | 0.048 | 20.69 |
| | MDM2 | 189 | 0.06 | 22877 | 9.512 | 2 | 9.935 | 0.053 | 19.02 |
| | TERT | 66 | 0.075 | 3200.9 | 6.746 | 2 | 4.892 | 0.074 | 13.49 |
| | XRCC6 | 264 | 0.064 | 41296 | 0.568 | 2 | 232.4 | 0.88 | 1.136 |
| | TERF1 | 307 | 0.026 | 71236 | 9.746 | 2 | 15.75 | 0.051 | 19.49 |
| | STUB1 | 239 | 0.055 | 35206 | 15 | 2 | 7.969 | 0.033 | 29.99 |
| | TERT | 66 | 0.075 | 3200.9 | 6.746 | 2 | 4.892 | 0.074 | 13.49 |
| | POT1 | 211 | 0.024 | 35382 | 6.882 | 2 | 15.33 | 0.073 | 13.76 |
| | TERT | 66 | 0.075 | 3200.9 | 6.746 | 2 | 4.892 | 0.074 | 13.49 |
| | MTOR | 140 | 0.061 | 13361 | 10.34 | 2 | 6.768 | 0.048 | 20.69 |
| | YWHAQ | 410 | 0 | 167690 | 22 | 2 | 9.32 | 0.023 | 43.99 |
| | RUVBL2 | 223 | 0.085 | 25036 | 20.62 | 2 | 5.408 | 0.024 | 41.24 |
| WDR70-RCOR1 | SMARCC2 | 129 | 0.129 | 6769.4 | 0.623 | 2 | 103.5 | 0.803 | 1.246 |
| | HDAC1 | 478 | 0 | 212982 | 1.996 | 2 | 119.7 | 0.251 | 3.992 |
| | HDAC3 | 23 | 0.134 | 343.97 | 0.509 | 2 | 22.59 | 0.982 | 1.018 |
| | HDAC2 | 301 | 0.067 | 40883 | 1.874 | 2 | 80.31 | 0.267 | 3.748 |
| | KDM1A | 210 | 0.054 | 31401 | 3.238 | 2 | 32.43 | 0.154 | 6.476 |
| | RCOR1 | 52 | 0.125 | 1397.5 | 8.115 | 2 | 3.204 | 0.062 | 16.23 |
| | NR2C1 | 27 | 0.399 | 167.85 | 11.93 | 2 | 1.132 | 0.042 | 23.86 |
| | SMARCE1 | 118 | 0.091 | 8801.9 | 12.54 | 2 | 4.706 | 0.04 | 25.08 |
| | CTBP1 | 148 | 0.058 | 14902 | 0.751 | 2 | 98.54 | 0.666 | 1.502 |
| | NR2E1 | 31 | 0.067 | 804.9 | 3.875 | 2 | 4 | 0.129 | 7.75 |
| | SMARCA4 | 200 | 0.088 | 20379 | 3.512 | 2 | 28.47 | 0.142 | 7.024 |
| | CTBP2 | 99 | 0.035 | 7872.9 | 5.293 | 2 | 9.352 | 0.094 | 10.59 |
| | KDM5B | 210 | 0.054 | 31401 | 11.24 | 2 | 9.343 | 0.044 | 22.48 |
| | MTA3 | 55 | 0.195 | 1541.7 | 12.32 | 2 | 2.232 | 0.041 | 24.64 |
| | KDM1A | 210 | 0.054 | 31401 | 3.238 | 2 | 32.43 | 0.154 | 6.476 |
| | HDAC3 | 23 | 0.134 | 343.97 | 0.509 | 2 | 22.59 | 0.982 | 1.018 |
| | CTBP1 | 148 | 0.058 | 14902 | 0.751 | 2 | 98.54 | 0.666 | 1.502 |
| YWHAE-NUTM2B | LARP1 | 82 | 0.211 | 2946.7 | 18.89 | 2 | 2.17 | 0.026 | 37.78 |
| | NOS2 | 146 | 0.095 | 12856 | 15.63 | 2 | 4.671 | 0.032 | 31.26 |

| | | | | | | | | | |
|---|---|---|---|---|---|---|---|---|---|
| | YWHAQ | 410 | 0 | 167690 | 22 | 2 | 9.32 | 0.023 | 43.99 |
| | YWHAG | 272 | 0.056 | 37593 | 16.99 | 2 | 8.003 | 0.029 | 33.99 |
| | HUWE1 | 453 | 0 | 203852 | 32 | 2 | 7.079 | 0.016 | 63.99 |
| | MAST2 | 20 | 0.121 | 287.33 | 4.095 | 2 | 2.442 | 0.122 | 8.19 |
| | NTRK1 | 166 | 0.045 | 22877 | 9.512 | 2 | 8.726 | 0.053 | 19.02 |
| | VCP | 586 | 0 | 341640 | 1.997 | 2 | 146.7 | 0.25 | 3.994 |
| | AKT1 | 60 | 0.16 | 1929 | 14.25 | 2 | 2.106 | 0.035 | 28.49 |
| | MAP2K1 | 59 | 0.099 | 2307.9 | 7.492 | 2 | 3.938 | 0.067 | 14.98 |
| | FBXW11 | 76 | 0.12 | 3995 | 9.977 | 2 | 3.809 | 0.05 | 19.95 |
| | ARAF | 87 | 0.058 | 5915.6 | 6.886 | 2 | 6.317 | 0.073 | 13.77 |
| | CUL3 | 70 | 0.081 | 3676.9 | 10.37 | 2 | 3.375 | 0.048 | 20.74 |
| | PARK2 | 407 | 0 | 164430 | 1.995 | 2 | 102 | 0.251 | 3.99 |
| | RAF1 | 150 | 0.066 | 15466 | 11.75 | 2 | 6.385 | 0.043 | 23.49 |
| | YWHAZ | 335 | 0.049 | 63935 | 18.42 | 2 | 9.094 | 0.027 | 36.84 |
| | UBXN1 | 134 | 0.076 | 11651 | 11.88 | 2 | 5.639 | 0.042 | 23.76 |
| | TP53 | 961 | 0 | 920640 | 1.998 | 2 | 240.5 | 0.25 | 3.996 |
| | RUVBL2 | 223 | 0.085 | 25036 | 20.62 | 2 | 5.408 | 0.024 | 41.24 |
| | BTRC | 42 | 0.211 | 765.63 | 12.19 | 2 | 1.723 | 0.041 | 24.38 |
| | TUBB | 217 | 0.107 | 18784 | 25.04 | 2 | 4.334 | 0.02 | 50.07 |
| | CDC37 | 44 | 0.083 | 1464.9 | 5.364 | 2 | 4.101 | 0.093 | 10.73 |
| | YWHAH | 180 | 0.063 | 18761 | 13.22 | 2 | 6.807 | 0.038 | 26.44 |
| | MAST3 | 43 | 0.064 | 1549.5 | 4.591 | 2 | 4.683 | 0.109 | 9.182 |
| | BRAF | 42 | 0.211 | 765.63 | 10.19 | 2 | 2.061 | 0.049 | 20.38 |
| | YWHAB | 311 | 0.036 | 61018 | 29.33 | 2 | 5.302 | 0.017 | 58.65 |
| | KSR1 | 16 | 0.21 | 142.33 | 4.625 | 2 | 1.73 | 0.108 | 9.25 |
| | YWHAE | 349 | 0.05 | 71479 | 19.33 | 2 | 9.029 | 0.026 | 38.65 |
| | CUL1 | 670 | 0 | 446892 | 12 | 2 | 27.92 | 0.042 | 23.99 |
| | MAPK7 | 62 | 0.105 | 2585.6 | 8.194 | 2 | 3.783 | 0.061 | 16.39 |
| | YWHAZ | 335 | 0.049 | 63935 | 18.42 | 2 | 9.094 | 0.027 | 36.84 |
| | IGF1R | 65 | 0.109 | 2628.6 | 8.818 | 2 | 3.686 | 0.057 | 17.64 |
| | IRS1 | 58 | 0.162 | 1617 | 11.05 | 2 | 2.624 | 0.045 | 22.1 |
| | YWHAQ | 410 | 0 | 167690 | 22 | 2 | 9.32 | 0.023 | 43.99 |
| | MST1R | 20 | 0.342 | 139.7 | 8.095 | 2 | 1.235 | 0.062 | 16.19 |
| | NTRK1 | 166 | 0.045 | 22877 | 9.512 | 2 | 8.726 | 0.053 | 19.02 |
| | CBL | 233 | 0.061 | 32363 | 6.043 | 2 | 19.28 | 0.083 | 12.09 |
| | TUBB | 217 | 0.107 | 18784 | 25.04 | 2 | 4.334 | 0.02 | 50.07 |
| | YWHAB | 311 | 0.036 | 61018 | 29.33 | 2 | 5.302 | 0.017 | 58.65 |
| | YWHAH | 180 | 0.063 | 18761 | 13.22 | 2 | 6.807 | 0.038 | 26.44 |
| | GRB2 | 515 | 0 | 263682 | 26 | 2 | 9.905 | 0.019 | 51.99 |
| | ABL1 | 173 | 0.076 | 17247 | 14.9 | 2 | 5.805 | 0.034 | 29.8 |
| | SORBS2 | 46 | 0.16 | 1265 | 8.826 | 2 | 2.606 | 0.057 | 17.65 |
| | BCAR1 | 63 | 0.12 | 2352.9 | 9.312 | 2 | 3.383 | 0.054 | 18.62 |
| | YWHAE | 349 | 0.05 | 71479 | 19.33 | 2 | 9.029 | 0.026 | 38.65 |
| | VCP | 586 | 0 | 341640 | 1.997 | 2 | 146.7 | 0.25 | 3.994 |
| | CDK2 | 233 | 0.061 | 32363 | 4.043 | 2 | 28.82 | 0.124 | 8.086 |
| | CUL1 | 670 | 0 | 446892 | 12 | 2 | 27.92 | 0.042 | 23.99 |
| | CDC37 | 44 | 0.083 | 1464.9 | 5.364 | 2 | 4.101 | 0.093 | 10.73 |
| | LRRK2 | 138 | 0.057 | 14507 | 9.638 | 2 | 7.159 | 0.052 | 19.28 |

| | | | | | | | | | |
|---|---|---|---|---|---|---|---|---|---|
| | YWHAE | 349 | 0.05 | 71479 | 19.33 | 2 | 9.029 | 0.026 | 38.65 |
| YWHAE-NUTM2A-AS1 | LARP1 | 82 | 0.211 | 2946.7 | 18.89 | 2 | 2.17 | 0.026 | 37.78 |
| | NOS2 | 146 | 0.095 | 12856 | 15.63 | 2 | 4.671 | 0.032 | 31.26 |
| | YWHAQ | 410 | 0 | 167690 | 22 | 2 | 9.32 | 0.023 | 43.99 |
| | YWHAG | 272 | 0.056 | 37593 | 16.99 | 2 | 8.003 | 0.029 | 33.99 |
| | HUWE1 | 453 | 0 | 203852 | 32 | 2 | 7.079 | 0.016 | 63.99 |
| | MAST2 | 20 | 0.121 | 287.33 | 4.095 | 2 | 2.442 | 0.122 | 8.19 |
| | NTRK1 | 166 | 0.045 | 22877 | 9.512 | 2 | 8.726 | 0.053 | 19.02 |
| | VCP | 586 | 0 | 341640 | 1.997 | 2 | 146.7 | 0.25 | 3.994 |
| | AKT1 | 60 | 0.16 | 1929 | 14.25 | 2 | 2.106 | 0.035 | 28.49 |
| | MAP2K1 | 59 | 0.099 | 2307.9 | 7.492 | 2 | 3.938 | 0.067 | 14.98 |
| | FBXW11 | 76 | 0.12 | 3995 | 9.977 | 2 | 3.809 | 0.05 | 19.95 |
| | ARAF | 87 | 0.058 | 5915.6 | 6.886 | 2 | 6.317 | 0.073 | 13.77 |
| | CUL3 | 70 | 0.081 | 3676.9 | 10.37 | 2 | 3.375 | 0.048 | 20.74 |
| | PARK2 | 407 | 0 | 164430 | 1.995 | 2 | 102 | 0.251 | 3.99 |
| | RAF1 | 150 | 0.066 | 15466 | 11.75 | 2 | 6.385 | 0.043 | 23.49 |
| | YWHAZ | 335 | 0.049 | 63935 | 18.42 | 2 | 9.094 | 0.027 | 36.84 |
| | UBXN1 | 134 | 0.076 | 11651 | 11.88 | 2 | 5.639 | 0.042 | 23.76 |
| | TP53 | 961 | 0 | 920640 | 1.998 | 2 | 240.5 | 0.25 | 3.996 |
| | RUVBL2 | 223 | 0.085 | 25036 | 20.62 | 2 | 5.408 | 0.024 | 41.24 |
| | BTRC | 42 | 0.211 | 765.63 | 12.19 | 2 | 1.723 | 0.041 | 24.38 |
| | TUBB | 217 | 0.107 | 18784 | 25.04 | 2 | 4.334 | 0.02 | 50.07 |
| | CDC37 | 44 | 0.083 | 1464.9 | 5.364 | 2 | 4.101 | 0.093 | 10.73 |
| | YWHAH | 180 | 0.063 | 18761 | 13.22 | 2 | 6.807 | 0.038 | 26.44 |
| | MAST3 | 43 | 0.064 | 1549.5 | 4.591 | 2 | 4.683 | 0.109 | 9.182 |
| | BRAF | 42 | 0.211 | 765.63 | 10.19 | 2 | 2.061 | 0.049 | 20.38 |
| | YWHAB | 311 | 0.036 | 61018 | 29.33 | 2 | 5.302 | 0.017 | 58.65 |
| | KSR1 | 16 | 0.21 | 142.33 | 4.625 | 2 | 1.73 | 0.108 | 9.25 |
| | YWHAE | 349 | 0.05 | 71479 | 19.33 | 2 | 9.029 | 0.026 | 38.65 |
| | CUL1 | 670 | 0 | 446892 | 12 | 2 | 27.92 | 0.042 | 23.99 |
| | MAPK7 | 62 | 0.105 | 2585.6 | 8.194 | 2 | 3.783 | 0.061 | 16.39 |
| | YWHAZ | 335 | 0.049 | 63935 | 18.42 | 2 | 9.094 | 0.027 | 36.84 |
| | IGF1R | 65 | 0.109 | 2628.6 | 8.818 | 2 | 3.686 | 0.057 | 17.64 |
| | IRS1 | 58 | 0.162 | 1617 | 11.05 | 2 | 2.624 | 0.045 | 22.1 |
| | YWHAQ | 410 | 0 | 167690 | 22 | 2 | 9.32 | 0.023 | 43.99 |
| | MST1R | 20 | 0.342 | 139.7 | 8.095 | 2 | 1.235 | 0.062 | 16.19 |
| | NTRK1 | 166 | 0.045 | 22877 | 9.512 | 2 | 8.726 | 0.053 | 19.02 |
| | CBL | 233 | 0.061 | 32363 | 6.043 | 2 | 19.28 | 0.083 | 12.09 |
| | TUBB | 217 | 0.107 | 18784 | 25.04 | 2 | 4.334 | 0.02 | 50.07 |
| | YWHAB | 311 | 0.036 | 61018 | 29.33 | 2 | 5.302 | 0.017 | 58.65 |
| | YWHAH | 180 | 0.063 | 18761 | 13.22 | 2 | 6.807 | 0.038 | 26.44 |
| | GRB2 | 515 | 0 | 263682 | 26 | 2 | 9.905 | 0.019 | 51.99 |
| | ABL1 | 173 | 0.076 | 17247 | 14.9 | 2 | 5.805 | 0.034 | 29.8 |
| | SORBS2 | 46 | 0.16 | 1265 | 8.826 | 2 | 2.606 | 0.057 | 17.65 |
| | BCAR1 | 63 | 0.12 | 2352.9 | 9.312 | 2 | 3.383 | 0.054 | 18.62 |
| | YWHAE | 349 | 0.05 | 71479 | 19.33 | 2 | 9.029 | 0.026 | 38.65 |
| | VCP | 586 | 0 | 341640 | 1.997 | 2 | 146.7 | 0.25 | 3.994 |
| | CDK2 | 233 | 0.061 | 32363 | 4.043 | 2 | 28.82 | 0.124 | 8.086 |

| | | | | | | | | | |
|---|---|---|---|---|---|---|---|---|---|
| | CUL1 | 670 | 0 | 446892 | 12 | 2 | 27.92 | 0.042 | 23.99 |
| | CDC37 | 44 | 0.083 | 1464.9 | 5.364 | 2 | 4.101 | 0.093 | 10.73 |
| | LRRK2 | 138 | 0.057 | 14507 | 9.638 | 2 | 7.159 | 0.052 | 19.28 |
| | YWHAE | 349 | 0.05 | 71479 | 19.33 | 2 | 9.029 | 0.026 | 38.65 |
| YWHAE-NUTM2A | LARP1 | 82 | 0.211 | 2946.7 | 18.89 | 2 | 2.17 | 0.026 | 37.78 |
| | NOS2 | 146 | 0.095 | 12856 | 15.63 | 2 | 4.671 | 0.032 | 31.26 |
| | YWHAQ | 410 | 0 | 167690 | 22 | 2 | 9.32 | 0.023 | 43.99 |
| | YWHAG | 272 | 0.056 | 37593 | 16.99 | 2 | 8.003 | 0.029 | 33.99 |
| | HUWE1 | 453 | 0 | 203852 | 32 | 2 | 7.079 | 0.016 | 63.99 |
| | MAST2 | 20 | 0.121 | 287.33 | 4.095 | 2 | 2.442 | 0.122 | 8.19 |
| | NTRK1 | 166 | 0.045 | 22877 | 9.512 | 2 | 8.726 | 0.053 | 19.02 |
| | VCP | 586 | 0 | 341640 | 1.997 | 2 | 146.7 | 0.25 | 3.994 |
| | AKT1 | 60 | 0.16 | 1929 | 14.25 | 2 | 2.106 | 0.035 | 28.49 |
| | MAP2K1 | 59 | 0.099 | 2307.9 | 7.492 | 2 | 3.938 | 0.067 | 14.98 |
| | FBXW11 | 76 | 0.12 | 3995 | 9.977 | 2 | 3.809 | 0.05 | 19.95 |
| | ARAF | 87 | 0.058 | 5915.6 | 6.886 | 2 | 6.317 | 0.073 | 13.77 |
| | CUL3 | 70 | 0.081 | 3676.9 | 10.37 | 2 | 3.375 | 0.048 | 20.74 |
| | PARK2 | 407 | 0 | 164430 | 1.995 | 2 | 102 | 0.251 | 3.99 |
| | RAF1 | 150 | 0.066 | 15466 | 11.75 | 2 | 6.385 | 0.043 | 23.49 |
| | YWHAZ | 335 | 0.049 | 63935 | 18.42 | 2 | 9.094 | 0.027 | 36.84 |
| | UBXN1 | 134 | 0.076 | 11651 | 11.88 | 2 | 5.639 | 0.042 | 23.76 |
| | TP53 | 961 | 0 | 920640 | 1.998 | 2 | 240.5 | 0.25 | 3.996 |
| | RUVBL2 | 223 | 0.085 | 25036 | 20.62 | 2 | 5.408 | 0.024 | 41.24 |
| | BTRC | 42 | 0.211 | 765.63 | 12.19 | 2 | 1.723 | 0.041 | 24.38 |
| | TUBB | 217 | 0.107 | 18784 | 25.04 | 2 | 4.334 | 0.02 | 50.07 |
| | CDC37 | 44 | 0.083 | 1464.9 | 5.364 | 2 | 4.101 | 0.093 | 10.73 |
| | YWHAH | 180 | 0.063 | 18761 | 13.22 | 2 | 6.807 | 0.038 | 26.44 |
| | MAST3 | 43 | 0.064 | 1549.5 | 4.591 | 2 | 4.683 | 0.109 | 9.182 |
| | BRAF | 42 | 0.211 | 765.63 | 10.19 | 2 | 2.061 | 0.049 | 20.38 |
| | YWHAB | 311 | 0.036 | 61018 | 29.33 | 2 | 5.302 | 0.017 | 58.65 |
| | KSR1 | 16 | 0.21 | 142.33 | 4.625 | 2 | 1.73 | 0.108 | 9.25 |
| | YWHAE | 349 | 0.05 | 71479 | 19.33 | 2 | 9.029 | 0.026 | 38.65 |
| | CUL1 | 670 | 0 | 446892 | 12 | 2 | 27.92 | 0.042 | 23.99 |
| | MAPK7 | 62 | 0.105 | 2585.6 | 8.194 | 2 | 3.783 | 0.061 | 16.39 |
| | YWHAZ | 335 | 0.049 | 63935 | 18.42 | 2 | 9.094 | 0.027 | 36.84 |
| | IGF1R | 65 | 0.109 | 2628.6 | 8.818 | 2 | 3.686 | 0.057 | 17.64 |
| | IRS1 | 58 | 0.162 | 1617 | 11.05 | 2 | 2.624 | 0.045 | 22.1 |
| | YWHAQ | 410 | 0 | 167690 | 22 | 2 | 9.32 | 0.023 | 43.99 |
| | MST1R | 20 | 0.342 | 139.7 | 8.095 | 2 | 1.235 | 0.062 | 16.19 |
| | NTRK1 | 166 | 0.045 | 22877 | 9.512 | 2 | 8.726 | 0.053 | 19.02 |
| | CBL | 233 | 0.061 | 32363 | 6.043 | 2 | 19.28 | 0.083 | 12.09 |
| | TUBB | 217 | 0.107 | 18784 | 25.04 | 2 | 4.334 | 0.02 | 50.07 |
| | YWHAB | 311 | 0.036 | 61018 | 29.33 | 2 | 5.302 | 0.017 | 58.65 |
| | YWHAH | 180 | 0.063 | 18761 | 13.22 | 2 | 6.807 | 0.038 | 26.44 |
| | GRB2 | 515 | 0 | 263682 | 26 | 2 | 9.905 | 0.019 | 51.99 |
| | ABL1 | 173 | 0.076 | 17247 | 14.9 | 2 | 5.805 | 0.034 | 29.8 |
| | SORBS2 | 46 | 0.16 | 1265 | 8.826 | 2 | 2.606 | 0.057 | 17.65 |
| | BCAR1 | 63 | 0.12 | 2352.9 | 9.312 | 2 | 3.383 | 0.054 | 18.62 |

| | | | | | | | | | |
|---|---|---|---|---|---|---|---|---|---|
| | YWHAE | 349 | 0.05 | 71479 | 19.33 | 2 | 9.029 | 0.026 | 38.65 |
| | VCP | 586 | 0 | 341640 | 1.997 | 2 | 146.7 | 0.25 | 3.994 |
| | CDK2 | 233 | 0.061 | 32363 | 4.043 | 2 | 28.82 | 0.124 | 8.086 |
| | CUL1 | 670 | 0 | 446892 | 12 | 2 | 27.92 | 0.042 | 23.99 |
| | CDC37 | 44 | 0.083 | 1464.9 | 5.364 | 2 | 4.101 | 0.093 | 10.73 |
| | LRRK2 | 138 | 0.057 | 14507 | 9.638 | 2 | 7.159 | 0.052 | 19.28 |
| | YWHAE | 349 | 0.05 | 71479 | 19.33 | 2 | 9.029 | 0.026 | 38.65 |

**Table S19: ST-PAS (TRAINING)**

| | Essential Community Vertices | D | CC | BC | Avg.D (avgD) | Net.Diam (ND) | PAS | PAS/D | D/PAS |
|---|---|---|---|---|---|---|---|---|---|
| ARGLU1-CXCR4 | APP | 9 | 0.286 | 18.5 | 3.6 | 2 | 1.25 | 0.139 | 7.2 |
| | CHERP | 57 | 0.16 | 1693 | 10.8 | 2 | 2.649 | 0.046 | 21.5 |
| | SNRNP70 | 125 | 0.178 | 5871 | 23.8 | 2 | 2.622 | 0.021 | 47.7 |
| | SRPK1 | 221 | 0.052 | 31598 | 13.3 | 2 | 8.282 | 0.037 | 26.7 |
| | SRPK2 | 449 | 0 | 2E+05 | 2 | 2 | 112.5 | 0.251 | 3.99 |
| | PTK2 | 114 | 0.084 | 8149 | 11.4 | 2 | 5.022 | 0.044 | 22.7 |
| | JAK2 | 41 | 0.133 | 1029 | 14.1 | 2 | 1.449 | 0.035 | 28.3 |
| | SOCS3 | 70 | 0.071 | 3785 | 6.82 | 2 | 5.134 | 0.073 | 13.6 |
| | PTPN11 | 106 | 0.071 | 7970 | 9.34 | 2 | 5.675 | 0.054 | 18.7 |
| | NTRK1 | 166 | 0.045 | 22877 | 9.51 | 2 | 8.726 | 0.053 | 19 |
| | PTK2 | 114 | 0.084 | 8149 | 11.4 | 2 | 5.022 | 0.044 | 22.7 |
| | ELAVL1 | 213 | 0.076 | 28937 | 18.7 | 2 | 5.706 | 0.027 | 37.3 |
| | SRPK1 | 221 | 0.052 | 31598 | 13.3 | 2 | 8.282 | 0.037 | 26.7 |
| | NTRK1 | 166 | 0.045 | 22877 | 9.51 | 2 | 8.726 | 0.053 | 19 |
| | PTK2 | 114 | 0.084 | 8149 | 11.4 | 2 | 5.022 | 0.044 | 22.7 |
| | JAK2 | 41 | 0.133 | 1029 | 14.1 | 2 | 1.449 | 0.035 | 28.3 |
| | JAK3 | 72 | 0.045 | 4518 | 5.15 | 2 | 6.989 | 0.097 | 10.3 |
| | SOCS3 | 70 | 0.071 | 3785 | 6.82 | 2 | 5.134 | 0.073 | 13.6 |
| | PTPN11 | 106 | 0.071 | 7970 | 9.34 | 2 | 5.675 | 0.054 | 18.7 |
| | STAM | 58 | 0.099 | 2336 | 7.49 | 2 | 3.871 | 0.067 | 15 |
| ATXN10-FBLN1 | FN1 | 23 | 0.134 | 344 | 0.11 | 2 | 105.5 | 4.587 | 0.22 |
| | ATXN10 | 32 | 0.089 | 823.4 | 4.61 | 2 | 3.474 | 0.109 | 9.21 |
| | EGFR | 833 | 0 | 7E+05 | 52 | 2 | 8.01 | 0.01 | 104 |
| | GSTK1 | 45 | 0.068 | 1519 | 4.8 | 2 | 4.688 | 0.104 | 9.6 |
| | ATXN10 | 32 | 0.089 | 823.4 | 4.61 | 2 | 3.474 | 0.109 | 9.21 |
| | VCP | 586 | 0 | 3E+05 | 2 | 2 | 146.7 | 0.25 | 3.99 |
| | BSG | 125 | 0.026 | 13028 | 5.19 | 2 | 12.04 | 0.096 | 10.4 |
| | CUL3 | 70 | 0.081 | 3677 | 10.4 | 2 | 3.375 | 0.048 | 20.7 |
| | VCP | 586 | 0 | 3E+05 | 2 | 2 | 146.7 | 0.25 | 3.99 |
| | ABCE1 | 236 | 0.037 | 39562 | 10.7 | 2 | 11.03 | 0.047 | 21.4 |
| | ATXN10 | 32 | 0.089 | 823.4 | 4.61 | 2 | 3.474 | 0.109 | 9.21 |
| | APP | 9 | 0.286 | 18.5 | 3.6 | 2 | 1.25 | 0.139 | 7.2 |
| | YWHAQ | 180 | 0.063 | 18761 | 13.2 | 2 | 6.807 | 0.038 | 26.4 |
| | CUL3 | 70 | 0.081 | 3677 | 10.4 | 2 | 3.375 | 0.048 | 20.7 |

| | | | | | | | | | |
|---|---|---|---|---|---|---|---|---|---|
| BCAS3-NFS1 | CTBP1 | 148 | 0.058 | 14902 | 0.75 | 2 | 98.54 | 0.666 | 1.5 |
| | CTBP2 | 99 | 0.035 | 7873 | 5.29 | 2 | 9.352 | 0.094 | 10.6 |
| | BCAS3 | 32 | 0.087 | 807.4 | 4.55 | 2 | 3.52 | 0.11 | 9.09 |
| | KAT2B | 175 | 0.066 | 18318 | 13.4 | 2 | 6.538 | 0.037 | 26.8 |
| | CTBP1 | 148 | 0.058 | 14902 | 0.75 | 2 | 98.54 | 0.666 | 1.5 |
| | CTBP2 | 99 | 0.035 | 7873 | 5.29 | 2 | 9.352 | 0.094 | 10.6 |
| | CDC23 | 101 | 0.079 | 7577 | 9.74 | 2 | 5.183 | 0.051 | 19.5 |
| | KAT2B | 175 | 0.066 | 18318 | 13.4 | 2 | 6.538 | 0.037 | 26.8 |
| | BCAS3 | 32 | 0.087 | 807.4 | 4.55 | 2 | 3.52 | 0.11 | 9.09 |
| BCAS4-BCAS3 | CTBP1 | 148 | 0.058 | 14902 | 0.75 | 2 | 98.54 | 0.666 | 1.5 |
| | CTBP2 | 99 | 0.035 | 7873 | 5.29 | 2 | 9.352 | 0.094 | 10.6 |
| | BCAS3 | 32 | 0.087 | 807.4 | 4.55 | 2 | 3.52 | 0.11 | 9.09 |
| | KAT2B | 175 | 0.066 | 18318 | 13.4 | 2 | 6.538 | 0.037 | 26.8 |
| | CTBP1 | 148 | 0.058 | 14902 | 0.75 | 2 | 98.54 | 0.666 | 1.5 |
| | CTBP2 | 99 | 0.035 | 7873 | 5.29 | 2 | 9.352 | 0.094 | 10.6 |
| | CDC23 | 101 | 0.079 | 7577 | 9.74 | 2 | 5.183 | 0.051 | 19.5 |
| | KAT2B | 175 | 0.066 | 18318 | 13.4 | 2 | 6.538 | 0.037 | 26.8 |
| | BCAS3 | 32 | 0.087 | 807.4 | 4.55 | 2 | 3.52 | 0.11 | 9.09 |
| BCL2L12-PRMT1 | NCOA2 | 64 | 0.121 | 2192 | 9.45 | 2 | 3.388 | 0.053 | 18.9 |
| | NCOA3 | 106 | 0.109 | 5992 | 13.4 | 2 | 3.966 | 0.037 | 26.7 |
| | NCOA1 | 101 | 0.12 | 5005 | 13.7 | 2 | 3.675 | 0.036 | 27.5 |
| | TP53 | 961 | 0 | 9E+05 | 2 | 2 | 240.5 | 0.25 | 4 |
| | ESR1 | 23 | 0.134 | 344 | 0.41 | 2 | 28.12 | 1.222 | 0.82 |
| | BRCA1 | 17 | 0.191 | 187.2 | 0.09 | 2 | 94.44 | 5.556 | 0.18 |
| | PRMT1 | 145 | 0.085 | 13214 | 14.1 | 2 | 5.148 | 0.036 | 28.2 |
| | THRB | 48 | 0.117 | 1384 | 7.25 | 2 | 3.31 | 0.069 | 14.5 |
| | CARM1 | 88 | 0.112 | 4523 | 11.6 | 2 | 3.802 | 0.043 | 23.1 |
| | NR1I2 | 32 | 0.093 | 794.1 | 4.73 | 2 | 3.385 | 0.106 | 9.45 |
| | AR | 60 | 0.16 | 1929 | 14.2 | 2 | 2.106 | 0.035 | 28.5 |
| | PPARA | 42 | 0.095 | 1312 | 5.77 | 2 | 3.641 | 0.087 | 11.5 |
| | EP300 | 453 | 0 | 2E+05 | 2 | 2 | 113.5 | 0.251 | 3.99 |
| | NCOA2 | 64 | 0.121 | 2192 | 9.45 | 2 | 3.388 | 0.053 | 18.9 |
| | NCOA3 | 106 | 0.109 | 5992 | 13.4 | 2 | 3.966 | 0.037 | 26.7 |
| | NCOA1 | 101 | 0.12 | 5005 | 13.7 | 2 | 3.675 | 0.036 | 27.5 |
| | EP300 | 453 | 0 | 2E+05 | 2 | 2 | 113.5 | 0.251 | 3.99 |
| | PARP1 | 219 | 0.08 | 25770 | 1.23 | 2 | 88.81 | 0.406 | 2.47 |
| CAPNS1-WDR62 | YWHAZ | 335 | 0.049 | 63935 | 18.4 | 2 | 9.094 | 0.027 | 36.8 |
| | FBXW11 | 76 | 0.12 | 3995 | 9.98 | 2 | 3.809 | 0.05 | 20 |
| | FN1 | 23 | 0.134 | 344 | 0.11 | 2 | 105.5 | 4.587 | 0.22 |
| | HUWE1 | 453 | 0 | 2E+05 | 32 | 2 | 7.079 | 0.016 | 64 |
| | YWHAQ | 180 | 0.063 | 18761 | 13.2 | 2 | 6.807 | 0.038 | 26.4 |
| | GAPDH | 196 | 0.098 | 19326 | 20.9 | 2 | 4.678 | 0.024 | 41.9 |
| | FERMT2 | 50 | 0.131 | 1582 | 8.24 | 2 | 3.036 | 0.061 | 16.5 |
| | YWHAH | 180 | 0.063 | 18761 | 13.2 | 2 | 6.807 | 0.038 | 26.4 |
| | VCAM1 | 442 | 0 | 2E+05 | 2 | 2 | 110.8 | 0.251 | 3.99 |
| | YWHAB | 311 | 0.036 | 61018 | 29.3 | 2 | 5.302 | 0.017 | 58.7 |
| | PAFAH1B1 | 59 | 0.105 | 2238 | 7.86 | 2 | 3.751 | 0.064 | 15.7 |

| | | | | | | | | | |
|---|---|---|---|---|---|---|---|---|---|
| | YWHAG | 272 | 0.056 | 37593 | 17 | 2 | 8.003 | 0.029 | 34 |
| | PAK2 | 72 | 0.091 | 3468 | 8.25 | 2 | 4.364 | 0.061 | 16.5 |
| | YWHAE | 349 | 0.05 | 71479 | 19.3 | 2 | 9.029 | 0.026 | 38.7 |
| | OGFOD1 | 32 | 0.226 | 585.4 | 8.73 | 2 | 1.833 | 0.057 | 17.5 |
| | MYO1E | 55 | 0.174 | 1894 | 11.2 | 2 | 2.46 | 0.045 | 22.4 |
| | ASNS | 82 | 0.102 | 4314 | 10.1 | 2 | 4.051 | 0.049 | 20.2 |
| | TBCB | 43 | 0.175 | 1085 | 9.14 | 2 | 2.353 | 0.055 | 18.3 |
| | CAPN2 | 46 | 0.134 | 1394 | 7.87 | 2 | 2.922 | 0.064 | 15.7 |
| | PROSC | 58 | 0.097 | 2535 | 7.42 | 2 | 3.906 | 0.067 | 14.8 |
| | YWHAE | 349 | 0.05 | 71479 | 19.3 | 2 | 9.029 | 0.026 | 38.7 |
| CCDC6-ANK3 | HDAC1 | 478 | 0 | 2E+05 | 2 | 2 | 119.7 | 0.251 | 3.99 |
| | NR3C1 | 157 | 0.069 | 15985 | 12.6 | 2 | 6.225 | 0.04 | 25.2 |
| | TRIM28 | 223 | 0.064 | 31811 | 16 | 2 | 6.961 | 0.031 | 32 |
| | SF3A1 | 153 | 0.162 | 8222 | 26.4 | 2 | 2.895 | 0.019 | 52.9 |
| | HNRNPR | 191 | 0.166 | 11408 | 33.3 | 2 | 2.866 | 0.015 | 66.6 |
| | BRCC3 | 52 | 0.165 | 1371 | 10 | 2 | 2.59 | 0.05 | 20.1 |
| | HDAC1 | 478 | 0 | 2E+05 | 2 | 2 | 119.7 | 0.251 | 3.99 |
| | NR3C1 | 157 | 0.069 | 15985 | 12.6 | 2 | 6.225 | 0.04 | 25.2 |
| | TRIM28 | 223 | 0.064 | 31811 | 16 | 2 | 6.961 | 0.031 | 32 |
| | ELAVL1 | 213 | 0.076 | 28937 | 18.7 | 2 | 5.706 | 0.027 | 37.3 |
| | SKP1 | 185 | 0.059 | 20423 | 12.7 | 2 | 7.259 | 0.039 | 25.5 |
| | HNRNPR | 191 | 0.166 | 11408 | 33.3 | 2 | 2.866 | 0.015 | 66.6 |
| | PPP1CA | 285 | 0.045 | 53592 | 14.8 | 2 | 9.653 | 0.034 | 29.5 |
| | BRCC3 | 52 | 0.165 | 1371 | 10 | 2 | 2.59 | 0.05 | 20.1 |
| | NTRK1 | 166 | 0.045 | 22877 | 9.51 | 2 | 8.726 | 0.053 | 19 |
| | SF3A1 | 153 | 0.162 | 8222 | 26.4 | 2 | 2.895 | 0.019 | 52.9 |
| | CUL1 | 670 | 0 | 4E+05 | 12 | 2 | 27.92 | 0.042 | 24 |
| | FBXW7 | 288 | 0.021 | 74536 | 8.01 | 2 | 17.98 | 0.062 | 16 |
| CCDC9-DHX34 | EIF4A3 | 833 | 0 | 7E+05 | 62 | 2 | 6.718 | 0.008 | 124 |
| | SNIP1 | 66 | 0.116 | 2646 | 9.37 | 2 | 3.521 | 0.053 | 18.7 |
| | CCDC9 | 16 | 0.175 | 165.3 | 4.35 | 2 | 1.838 | 0.115 | 8.71 |
| | PRPF40A | 178 | 0.091 | 18728 | 17.9 | 2 | 4.985 | 0.028 | 35.7 |
| CDC27-ST7L | CDC16 | 61 | 0.199 | 1567 | 13.7 | 2 | 2.219 | 0.036 | 27.5 |
| | CDC27 | 100 | 0.116 | 5243 | 13.3 | 2 | 3.771 | 0.038 | 26.5 |
| | MDC1 | 188 | 0.067 | 22418 | 14.4 | 2 | 6.512 | 0.035 | 28.9 |
| | CDC20 | 148 | 0.096 | 12082 | 16 | 2 | 4.633 | 0.031 | 31.9 |
| | CREBBP | 296 | 0.058 | 48179 | 1.06 | 2 | 139.5 | 0.471 | 2.12 |
| | ANAPC2 | 44 | 0.264 | 654.6 | 13.1 | 2 | 1.684 | 0.038 | 26.1 |
| | ANAPC7 | 55 | 0.239 | 964.6 | 14.4 | 2 | 1.91 | 0.035 | 28.8 |
| | CREBBP | 296 | 0.058 | 48179 | 1.06 | 2 | 139.5 | 0.471 | 2.12 |
| | E2F1 | 110 | 0.101 | 6727 | 12.9 | 2 | 4.257 | 0.039 | 25.8 |
| | RB1 | 217 | 0.059 | 30195 | 14.7 | 2 | 7.401 | 0.034 | 29.3 |
| | TFDP1 | 35 | 0.185 | 674.8 | 8.06 | 2 | 2.172 | 0.062 | 16.1 |
| | CDC16 | 61 | 0.199 | 1567 | 13.7 | 2 | 2.219 | 0.036 | 27.5 |
| | CDC27 | 100 | 0.116 | 5243 | 13.3 | 2 | 3.771 | 0.038 | 26.5 |
| | MDC1 | 188 | 0.067 | 22418 | 14.4 | 2 | 6.512 | 0.035 | 28.9 |
| | SMAD2 | 103 | 0.141 | 4686 | 2.25 | 2 | 22.89 | 0.222 | 4.5 |
| | TP53BP1 | 961 | 0 | 9E+05 | 2 | 2 | 240.5 | 0.25 | 4 |

| | | | | | | | | | |
|---|---|---|---|---|---|---|---|---|---|
| | CREBBP | 296 | 0.058 | 48179 | 1.06 | 2 | 139.5 | 0.471 | 2.12 |
| | UBE2S | 63 | 0.094 | 2807 | 7.59 | 2 | 4.152 | 0.066 | 15.2 |
| | ANAPC2 | 44 | 0.264 | 654.6 | 13.1 | 2 | 1.684 | 0.038 | 26.1 |
| | ANAPC7 | 55 | 0.239 | 964.6 | 14.4 | 2 | 1.91 | 0.035 | 28.8 |
| CDK7-RIN3 | BRCA1 | 17 | 0.191 | 187.2 | 0.09 | 2 | 94.44 | 5.556 | 0.18 |
| | TP53 | 961 | 0 | 9E+05 | 2 | 2 | 240.5 | 0.25 | 4 |
| | RUVBL2 | 223 | 0.085 | 25036 | 20.6 | 2 | 5.408 | 0.024 | 41.2 |
| | SUPT5H | 123 | 0.088 | 9108 | 12.6 | 2 | 4.864 | 0.04 | 25.3 |
| | ESR1 | 23 | 0.134 | 344 | 0.41 | 2 | 28.12 | 1.222 | 0.82 |
| | RPA1 | 463 | 0 | 2E+05 | 2 | 2 | 116 | 0.251 | 3.99 |
| | HNRNPU | 503 | 0 | 3E+05 | 2 | 2 | 126 | 0.251 | 3.99 |
| | RPA2 | 226 | 0.06 | 22557 | 9.51 | 2 | 11.88 | 0.053 | 19 |
| | CDK2 | 233 | 0.061 | 32363 | 4.04 | 2 | 28.82 | 0.124 | 8.09 |
| | POLR2A | 251 | 0.074 | 34919 | 1.24 | 2 | 101.3 | 0.404 | 2.48 |
| | GTF2H1 | 53 | 0.12 | 1854 | 8.07 | 2 | 3.282 | 0.062 | 16.1 |
| | RPA1 | 463 | 0 | 2E+05 | 2 | 2 | 116 | 0.251 | 3.99 |
| | RPA2 | 226 | 0.06 | 22557 | 9.51 | 2 | 11.88 | 0.053 | 19 |
| | POLR2A | 251 | 0.074 | 34919 | 1.24 | 2 | 101.3 | 0.404 | 2.48 |
| | CCNH | 51 | 0.143 | 1534 | 8.96 | 2 | 2.845 | 0.056 | 17.9 |
| | HDAC2 | 301 | 0.067 | 40883 | 1.87 | 2 | 80.31 | 0.267 | 3.75 |
| | TP53 | 961 | 0 | 9E+05 | 2 | 2 | 240.5 | 0.25 | 4 |
| | ESR1 | 23 | 0.134 | 344 | 0.41 | 2 | 28.12 | 1.222 | 0.82 |
| | MTA1 | 93 | 0.13 | 4605 | 13.8 | 2 | 3.367 | 0.036 | 27.6 |
| | GTF2H1 | 53 | 0.12 | 1854 | 8.07 | 2 | 3.282 | 0.062 | 16.1 |
| | ERCC3 | 46 | 0.137 | 1257 | 8 | 2 | 2.875 | 0.063 | 16 |
| | POLR2A | 251 | 0.074 | 34919 | 1.24 | 2 | 101.3 | 0.404 | 2.48 |
| | ERCC5 | 18 | 0.196 | 206.6 | 5.05 | 2 | 1.781 | 0.099 | 10.1 |
| | BRCA1 | 17 | 0.191 | 187.2 | 0.09 | 2 | 94.44 | 5.556 | 0.18 |
| | POLR2A | 251 | 0.074 | 34919 | 1.24 | 2 | 101.3 | 0.404 | 2.48 |
| | RUVBL2 | 223 | 0.085 | 25036 | 20.6 | 2 | 5.408 | 0.024 | 41.2 |
| | GTF2H1 | 53 | 0.12 | 1854 | 8.07 | 2 | 3.282 | 0.062 | 16.1 |
| | RPA1 | 463 | 0 | 2E+05 | 2 | 2 | 116 | 0.251 | 3.99 |
| | RPA2 | 226 | 0.06 | 22557 | 9.51 | 2 | 11.88 | 0.053 | 19 |
| | PRKCI | 69 | 0.09 | 3320 | 7.97 | 2 | 4.328 | 0.063 | 15.9 |
| | APP | 9 | 0.286 | 18.5 | 3.6 | 2 | 1.25 | 0.139 | 7.2 |
| | CDK7 | 76 | 0.117 | 3334 | 10.5 | 2 | 3.61 | 0.048 | 21.1 |
| | CDC37 | 44 | 0.083 | 1465 | 5.36 | 2 | 4.101 | 0.093 | 10.7 |
| CHERP-CPAMD8 | DHX8 | 63 | 0.134 | 2181 | 10 | 2 | 3.14 | 0.05 | 20.1 |
| | U2AF1 | 114 | 0.212 | 3674 | 25.8 | 2 | 2.213 | 0.019 | 51.5 |
| | RPA1 | 463 | 0 | 2E+05 | 2 | 2 | 116 | 0.251 | 3.99 |
| | PRPF40A | 178 | 0.091 | 18728 | 17.9 | 2 | 4.985 | 0.028 | 35.7 |
| | RPA2 | 226 | 0.06 | 22557 | 9.51 | 2 | 11.88 | 0.053 | 19 |
| | CHERP | 57 | 0.16 | 1693 | 10.8 | 2 | 2.649 | 0.046 | 21.5 |
| | SF3A2 | 26 | 0.142 | 450.3 | 6.33 | 2 | 2.053 | 0.079 | 12.7 |
| | RBM39 | 144 | 0.15 | 8588 | 23.3 | 2 | 3.092 | 0.021 | 46.6 |
| | EWSR1 | 643 | 0 | 4E+05 | 32 | 2 | 10.05 | 0.016 | 64 |
| | DHX8 | 63 | 0.134 | 2181 | 10 | 2 | 3.14 | 0.05 | 20.1 |
| | AGGF1 | 21 | 0.042 | 354.3 | 2.67 | 2 | 3.937 | 0.187 | 5.33 |

| | | | | | | | | | |
|---|---|---|---|---|---|---|---|---|---|
| | RNPS1 | 168 | 0.121 | 13587 | 22 | 2 | 3.812 | 0.023 | 44.1 |
| | SNIP1 | 66 | 0.116 | 2646 | 9.37 | 2 | 3.521 | 0.053 | 18.7 |
| | SF3B4 | 100 | 0.128 | 5654 | 14.5 | 2 | 3.44 | 0.034 | 29.1 |
| | NTRK1 | 166 | 0.045 | 22877 | 9.51 | 2 | 8.726 | 0.053 | 19 |
| | CHERP | 57 | 0.16 | 1693 | 10.8 | 2 | 2.649 | 0.046 | 21.5 |
| | SRPK1 | 221 | 0.052 | 31598 | 13.3 | 2 | 8.282 | 0.037 | 26.7 |
| | SRPK2 | 449 | 0 | 2E+05 | 2 | 2 | 112.5 | 0.251 | 3.99 |
| | U2AF1 | 114 | 0.212 | 3674 | 25.8 | 2 | 2.213 | 0.019 | 51.5 |
| | U2AF2 | 349 | 0.094 | 46692 | 34.6 | 2 | 5.046 | 0.014 | 69.2 |
| | RPA1 | 463 | 0 | 2E+05 | 2 | 2 | 116 | 0.251 | 3.99 |
| | APBB1 | 81 | 0.072 | 4892 | 7.58 | 2 | 5.343 | 0.066 | 15.2 |
| | PRPF40A | 178 | 0.091 | 18728 | 17.9 | 2 | 4.985 | 0.028 | 35.7 |
| | RPA2 | 226 | 0.06 | 22557 | 9.51 | 2 | 11.88 | 0.053 | 19 |
| | TTC14 | 11 | 0.345 | 43.9 | 5 | 2 | 1.1 | 0.1 | 10 |
| | RBM23 | 40 | 0.062 | 1364 | 4.29 | 2 | 4.659 | 0.116 | 8.59 |
| | SF3A2 | 26 | 0.142 | 450.3 | 6.33 | 2 | 2.053 | 0.079 | 12.7 |
| | SNRNP70 | 125 | 0.178 | 5871 | 23.8 | 2 | 2.622 | 0.021 | 47.7 |
| | RBM39 | 144 | 0.15 | 8588 | 23.3 | 2 | 3.092 | 0.021 | 46.6 |
| | EWSR1 | 643 | 0 | 4E+05 | 32 | 2 | 10.05 | 0.016 | 64 |
| | WBP4 | 44 | 0.174 | 1068 | 9.29 | 2 | 2.368 | 0.054 | 18.6 |
| CYTH1-PRPSAP1 | DDX17 | 142 | 0.185 | 5363 | 27.7 | 2 | 2.559 | 0.018 | 55.5 |
| | DDX5 | 203 | 0.145 | 13444 | 31 | 2 | 3.278 | 0.016 | 61.9 |
| | ILK | 211 | 0.046 | 30385 | 0.53 | 2 | 200.6 | 0.951 | 1.05 |
| | COPS5 | 795 | 0 | 6E+05 | 2 | 2 | 199 | 0.25 | 3.99 |
| | CYTH1 | 11 | 0.091 | 95 | 2.67 | 2 | 2.062 | 0.187 | 5.33 |
| | ARRB2 | 10 | 0.333 | 24.17 | 21 | 2 | 0.238 | 0.024 | 42 |
| | ARF6 | 44 | 0.069 | 1584 | 4.84 | 2 | 4.542 | 0.103 | 9.69 |
| | ARRB1 | 7 | 0.524 | 6.5 | 12.4 | 2 | 0.282 | 0.04 | 24.9 |
| | DDX5 | 203 | 0.145 | 13444 | 31 | 2 | 3.278 | 0.016 | 61.9 |
| | FBXW11 | 76 | 0.12 | 3995 | 9.98 | 2 | 3.809 | 0.05 | 20 |
| | DDX17 | 142 | 0.185 | 5363 | 27.7 | 2 | 2.559 | 0.018 | 55.5 |
| | ILK | 211 | 0.046 | 30385 | 0.53 | 2 | 200.6 | 0.951 | 1.05 |
| | ITGB2 | 26 | 0.086 | 533.3 | 4 | 2 | 3.25 | 0.125 | 8 |
| | COPS5 | 795 | 0 | 6E+05 | 2 | 2 | 199 | 0.25 | 3.99 |
| DLG1-CRYBG3 | DLG1 | 50 | 0.052 | 2113 | 4.47 | 2 | 5.592 | 0.112 | 8.94 |
| | LIN7A | 29 | 0.081 | 684.5 | 4.13 | 2 | 3.508 | 0.121 | 8.27 |
| | LIN7C | 35 | 0.126 | 836.2 | 6.11 | 2 | 2.864 | 0.082 | 12.2 |
| | APBA1 | 94 | 0.119 | 5309 | 3.85 | 2 | 12.22 | 0.13 | 7.69 |
| | CASK | 62 | 0.043 | 3276 | 4.57 | 2 | 6.782 | 0.109 | 9.14 |
| | DLG1 | 50 | 0.052 | 2113 | 4.47 | 2 | 5.592 | 0.112 | 8.94 |
| | NTRK1 | 166 | 0.045 | 22877 | 9.51 | 2 | 8.726 | 0.053 | 19 |
| | CASK | 62 | 0.043 | 3276 | 4.57 | 2 | 6.782 | 0.109 | 9.14 |
| | EPB41 | 29 | 0.126 | 505.3 | 5.33 | 2 | 2.719 | 0.094 | 10.7 |
| | DLG1 | 50 | 0.052 | 2113 | 4.47 | 2 | 5.592 | 0.112 | 8.94 |
| | KHDRBS1 | 161 | 0.069 | 17920 | 12.8 | 2 | 6.273 | 0.039 | 25.7 |
| | LCK | 83 | 0.061 | 5230 | 6.89 | 2 | 6.021 | 0.073 | 13.8 |
| | NTRK1 | 166 | 0.045 | 22877 | 9.51 | 2 | 8.726 | 0.053 | 19 |

| | | | | | | | | | |
|---|---|---|---|---|---|---|---|---|---|
| | MAPK1 | 214 | 0.042 | 32284 | 20.1 | 2 | 5.324 | 0.025 | 40.2 |
| | ARRB2 | 10 | 0.333 | 24.17 | 21 | 2 | 0.238 | 0.024 | 42 |
| | ARRB1 | 7 | 0.524 | 6.5 | 12.4 | 2 | 0.282 | 0.04 | 24.9 |
| DTX4-CCDC102B | MCM7 | 151 | 0.085 | 13247 | 14.5 | 2 | 5.206 | 0.034 | 29 |
| | CDK18 | 55 | 0.056 | 2535 | 4.93 | 2 | 5.579 | 0.101 | 9.86 |
| | LENG1 | 43 | 0.068 | 1460 | 4.73 | 2 | 4.548 | 0.106 | 9.45 |
| | TRIM54 | 127 | 0.031 | 13722 | 5.84 | 2 | 10.87 | 0.086 | 11.7 |
| | TRIM27 | 11 | 0.255 | 62.83 | 4.17 | 2 | 1.32 | 0.12 | 8.33 |
| | KIFC3 | 114 | 0.06 | 9573 | 8.67 | 2 | 6.577 | 0.058 | 17.3 |
| | SFN | 250 | 0.053 | 43097 | 15 | 2 | 8.356 | 0.033 | 29.9 |
| | MARK1 | 11 | 0.291 | 58.17 | 4.5 | 2 | 1.222 | 0.111 | 9 |
| | CCDC102B | 55 | 0.054 | 2313 | 4.76 | 2 | 5.772 | 0.105 | 9.53 |
| EHD4-FSIP1 | EHD4 | 36 | 0.111 | 882.8 | 5.73 | 2 | 3.141 | 0.087 | 11.5 |
| | CTPS2 | 28 | 0.111 | 559.3 | 4.71 | 2 | 2.97 | 0.106 | 9.43 |
| | EHD1 | 65 | 0.067 | 3116 | 6.15 | 2 | 5.281 | 0.081 | 12.3 |
| | EGFR | 833 | 0 | 7E+05 | 52 | 2 | 8.01 | 0.01 | 104 |
| | NTRK1 | 166 | 0.045 | 22877 | 9.51 | 2 | 8.726 | 0.053 | 19 |
| | WARS | 60 | 0.088 | 2479 | 7.05 | 2 | 4.256 | 0.071 | 14.1 |
| | PLCG1 | 112 | 0.071 | 8176 | 9.81 | 2 | 5.711 | 0.051 | 19.6 |
| | UBA2 | 78 | 0.098 | 3888 | 9.42 | 2 | 4.141 | 0.053 | 18.8 |
| | ADSL | 64 | 0.081 | 3019 | 7.02 | 2 | 4.562 | 0.071 | 14 |
| | UQCRC2 | 131 | 0.126 | 7740 | 18.3 | 2 | 3.585 | 0.027 | 36.5 |
| | PLCG1 | 112 | 0.071 | 8176 | 9.81 | 2 | 5.711 | 0.051 | 19.6 |
| | EHD4 | 36 | 0.111 | 882.8 | 5.73 | 2 | 3.141 | 0.087 | 11.5 |
| | EGFR | 833 | 0 | 7E+05 | 52 | 2 | 8.01 | 0.01 | 104 |
| | NTRK1 | 166 | 0.045 | 22877 | 9.51 | 2 | 8.726 | 0.053 | 19 |
| ELK4-SLC26A9 | BRCA1 | 17 | 0.191 | 187.2 | 0.09 | 2 | 94.44 | 5.556 | 0.18 |
| | MAPK3 | 176 | 0.027 | 24611 | 6.67 | 2 | 13.19 | 0.075 | 13.3 |
| | MAPK1 | 214 | 0.042 | 32284 | 20.1 | 2 | 5.324 | 0.025 | 40.2 |
| | ELK4 | 8 | 0.143 | 46 | 2.67 | 2 | 1.5 | 0.187 | 5.33 |
| | BRCA1 | 17 | 0.191 | 187.2 | 0.09 | 2 | 94.44 | 5.556 | 0.18 |
| | MAPK3 | 176 | 0.027 | 24611 | 6.67 | 2 | 13.19 | 0.075 | 13.3 |
| | MAPK1 | 214 | 0.042 | 32284 | 20.1 | 2 | 5.324 | 0.025 | 40.2 |
| | ELK4 | 8 | 0.143 | 46 | 2.67 | 2 | 1.5 | 0.187 | 5.33 |
| | BLM | 111 | 0.084 | 8357 | 11 | 2 | 5.025 | 0.045 | 22.1 |
| ERAL1-DIDO1 | HNRNPDL | 109 | 0.212 | 3840 | 24.6 | 2 | 2.214 | 0.02 | 49.2 |
| | RPA1 | 463 | 0 | 2E+05 | 2 | 2 | 116 | 0.251 | 3.99 |
| | RPA2 | 226 | 0.06 | 22557 | 9.51 | 2 | 11.88 | 0.053 | 19 |
| | HNRNPK | 243 | 0.127 | 23574 | 32.5 | 2 | 3.744 | 0.015 | 64.9 |
| | RBM15 | 42 | 0.152 | 1031 | 8.05 | 2 | 2.61 | 0.062 | 16.1 |
| | CUL3 | 70 | 0.081 | 3677 | 10.4 | 2 | 3.375 | 0.048 | 20.7 |
| | DIDO1 | 32 | 0.129 | 700.7 | 5.82 | 2 | 2.75 | 0.086 | 11.6 |
| | FUS | 319 | 0.082 | 50310 | 8 | 2 | 19.94 | 0.063 | 16 |
| | FUS | 319 | 0.082 | 50310 | 8 | 2 | 19.94 | 0.063 | 16 |
| | APP | 9 | 0.286 | 18.5 | 3.6 | 2 | 1.25 | 0.139 | 7.2 |
| | RPA1 | 463 | 0 | 2E+05 | 2 | 2 | 116 | 0.251 | 3.99 |
| | RPA2 | 226 | 0.06 | 22557 | 9.51 | 2 | 11.88 | 0.053 | 19 |

| | | | | | | | | | |
|---|---|---|---|---|---|---|---|---|---|
| | RBM15 | 42 | 0.152 | 1031 | 8.05 | 2 | 2.61 | 0.062 | 16.1 |
| | CUL3 | 70 | 0.081 | 3677 | 10.4 | 2 | 3.375 | 0.048 | 20.7 |
| | DIDO1 | 32 | 0.129 | 700.7 | 5.82 | 2 | 2.75 | 0.086 | 11.6 |
| | SRPK2 | 449 | 0 | 2E+05 | 2 | 2 | 112.5 | 0.251 | 3.99 |
| GMDS-CCND3 | PCNA | 274 | 0.05 | 50200 | 15.6 | 2 | 8.779 | 0.032 | 31.2 |
| | PPP1CC | 274 | 0.036 | 53066 | 11.8 | 2 | 11.62 | 0.042 | 23.6 |
| | RBL2 | 61 | 0.121 | 2052 | 9.13 | 2 | 3.341 | 0.055 | 18.3 |
| | PPP1CA | 285 | 0.045 | 53592 | 14.8 | 2 | 9.653 | 0.034 | 29.5 |
| | CCND3 | 49 | 0.075 | 1883 | 5.48 | 2 | 4.471 | 0.091 | 11 |
| | RB1 | 217 | 0.059 | 30195 | 14.7 | 2 | 7.401 | 0.034 | 29.3 |
| | POLD1 | 61 | 0.101 | 2589 | 7.9 | 2 | 3.859 | 0.063 | 15.8 |
| | CDK2 | 233 | 0.061 | 32363 | 4.04 | 2 | 28.82 | 0.124 | 8.09 |
| | CDK4 | 143 | 0.058 | 14433 | 10.2 | 2 | 7.033 | 0.049 | 20.3 |
| | CDK6 | 115 | 0.049 | 10296 | 7.44 | 2 | 7.725 | 0.067 | 14.9 |
| | CREBBP | 296 | 0.058 | 48179 | 1.06 | 2 | 139.5 | 0.471 | 2.12 |
| | GMDS | 22 | 0.09 | 370 | 3.64 | 2 | 3.025 | 0.138 | 7.27 |
| | NSFL1C | 77 | 0.092 | 4119 | 8.85 | 2 | 4.352 | 0.057 | 17.7 |
| | CTH | 35 | 0.052 | 1012 | 3.6 | 2 | 4.861 | 0.139 | 7.2 |
| | CAPN2 | 46 | 0.134 | 1394 | 7.87 | 2 | 2.922 | 0.064 | 15.7 |
| | ATIC | 70 | 0.087 | 3421 | 7.92 | 2 | 4.422 | 0.063 | 15.8 |
| | RARA | 108 | 0.084 | 7254 | 10.8 | 2 | 5.002 | 0.046 | 21.6 |
| | NCOA2 | 64 | 0.121 | 2192 | 9.45 | 2 | 3.388 | 0.053 | 18.9 |
| | VDR | 89 | 0.109 | 4934 | 0.37 | 2 | 119.9 | 1.348 | 0.74 |
| | CCND3 | 49 | 0.075 | 1883 | 5.48 | 2 | 4.471 | 0.091 | 11 |
| | CREBBP | 296 | 0.058 | 48179 | 1.06 | 2 | 139.5 | 0.471 | 2.12 |
| | MCM10 | 49 | 0.127 | 1613 | 7.92 | 2 | 3.093 | 0.063 | 15.8 |
| | RBX1 | 158 | 0.08 | 13795 | 14.3 | 2 | 5.513 | 0.035 | 28.7 |
| | CCND3 | 49 | 0.075 | 1883 | 5.48 | 2 | 4.471 | 0.091 | 11 |
| | APP | 9 | 0.286 | 18.5 | 3.6 | 2 | 1.25 | 0.139 | 7.2 |
| | NCOA2 | 64 | 0.121 | 2192 | 9.45 | 2 | 3.388 | 0.053 | 18.9 |
| | RARA | 108 | 0.084 | 7254 | 10.8 | 2 | 5.002 | 0.046 | 21.6 |
| | VDR | 89 | 0.109 | 4934 | 0.37 | 2 | 119.9 | 1.348 | 0.74 |
| | CCND3 | 49 | 0.075 | 1883 | 5.48 | 2 | 4.471 | 0.091 | 11 |
| | CREBBP | 296 | 0.058 | 48179 | 1.06 | 2 | 139.5 | 0.471 | 2.12 |
| HJURP-EIF4E2 | FBXW11 | 76 | 0.12 | 3995 | 9.98 | 2 | 3.809 | 0.05 | 20 |
| | TP53 | 961 | 0 | 9E+05 | 2 | 2 | 240.5 | 0.25 | 4 |
| | GIGYF2 | 53 | 0.158 | 1373 | 10 | 2 | 2.64 | 0.05 | 20.1 |
| | APP | 9 | 0.286 | 18.5 | 3.6 | 2 | 1.25 | 0.139 | 7.2 |
| | HUWE1 | 453 | 0 | 2E+05 | 32 | 2 | 7.079 | 0.016 | 64 |
| | EIF4E2 | 60 | 0.066 | 2914 | 5.8 | 2 | 5.17 | 0.086 | 11.6 |
| | YWHAB | 311 | 0.036 | 61018 | 29.3 | 2 | 5.302 | 0.017 | 58.7 |
| | SHMT2 | 427 | 0 | 2E+05 | 2 | 2 | 107 | 0.251 | 3.99 |
| | YWHAE | 349 | 0.05 | 71479 | 19.3 | 2 | 9.029 | 0.026 | 38.7 |
| INTS4-GAB2 | SRC | 221 | 0.056 | 31406 | 14.2 | 2 | 7.782 | 0.035 | 28.4 |
| | PLCG1 | 112 | 0.071 | 8176 | 9.81 | 2 | 5.711 | 0.051 | 19.6 |
| | GRB2 | 515 | 0 | 3E+05 | 26 | 2 | 9.905 | 0.019 | 52 |
| | ZAP70 | 48 | 0.187 | 1071 | 10.4 | 2 | 2.313 | 0.048 | 20.8 |
| | NTRK1 | 166 | 0.045 | 22877 | 9.51 | 2 | 8.726 | 0.053 | 19 |

| | | | | | | | | | |
|---|---|---|---|---|---|---|---|---|---|
| | SHC1 | 214 | 0.062 | 25801 | 15 | 2 | 7.138 | 0.033 | 30 |
| | PIK3CB | 25 | 0.23 | 298 | 7.23 | 2 | 1.729 | 0.069 | 14.5 |
| | PIK3R2 | 109 | 0.111 | 6175 | 13.9 | 2 | 3.923 | 0.036 | 27.8 |
| | PIK3R1 | 145 | 0.079 | 12814 | 13.1 | 2 | 5.515 | 0.038 | 26.3 |
| KDM5A-ANO2 | HDAC1 | 478 | 0 | 2E+05 | 2 | 2 | 119.7 | 0.251 | 3.99 |
| | HDAC2 | 301 | 0.067 | 40883 | 1.87 | 2 | 80.31 | 0.267 | 3.75 |
| | RBL1 | 69 | 0.108 | 2840 | 9.2 | 2 | 3.75 | 0.054 | 18.4 |
| | TBP | 156 | 0.071 | 15459 | 12.9 | 2 | 6.066 | 0.039 | 25.7 |
| | RB1 | 217 | 0.059 | 30195 | 14.7 | 2 | 7.401 | 0.034 | 29.3 |
| | VDR | 89 | 0.109 | 4934 | 0.37 | 2 | 119.9 | 1.348 | 0.74 |
| | KDM5A | 25 | 0.293 | 232.7 | 8.69 | 2 | 1.438 | 0.058 | 17.4 |
| | MORF4L1 | 106 | 0.087 | 6884 | 11 | 2 | 4.827 | 0.046 | 22 |
| | HDAC2 | 301 | 0.067 | 40883 | 1.87 | 2 | 80.31 | 0.267 | 3.75 |
| | EZH2 | 275 | 0.045 | 49579 | 1.23 | 2 | 111.5 | 0.406 | 2.47 |
| | KDM5A | 25 | 0.293 | 232.7 | 8.69 | 2 | 1.438 | 0.058 | 17.4 |
| | ESR1 | 23 | 0.134 | 344 | 0.41 | 2 | 28.12 | 1.222 | 0.82 |
| MAPK10-FAM13A | HDAC1 | 478 | 0 | 2E+05 | 2 | 2 | 119.7 | 0.251 | 3.99 |
| | TP53 | 961 | 0 | 9E+05 | 2 | 2 | 240.5 | 0.25 | 4 |
| | HDAC9 | 145 | 0.089 | 12651 | 28.7 | 2 | 2.523 | 0.017 | 57.5 |
| | JUN | 211 | 0.072 | 25977 | 17.1 | 2 | 6.184 | 0.029 | 34.1 |
| | DDX5 | 203 | 0.145 | 13444 | 31 | 2 | 3.278 | 0.016 | 61.9 |
| | ELK1 | 27 | 0.129 | 470.2 | 5.04 | 2 | 2.68 | 0.099 | 10.1 |
| | MAPK10 | 43 | 0.085 | 1338 | 5.35 | 2 | 4.019 | 0.093 | 10.7 |
| | RELA | 265 | 0.056 | 43653 | 16.6 | 2 | 7.966 | 0.03 | 33.3 |
| | CREBBP | 296 | 0.058 | 48179 | 1.06 | 2 | 139.5 | 0.471 | 2.12 |
| | ATF2 | 202 | 0.046 | 31394 | 11.1 | 2 | 9.133 | 0.045 | 22.1 |
| | APP | 9 | 0.286 | 18.5 | 3.6 | 2 | 1.25 | 0.139 | 7.2 |
| | MAPK10 | 43 | 0.085 | 1338 | 5.35 | 2 | 4.019 | 0.093 | 10.7 |
| | MAP2K4 | 40 | 0.147 | 903.7 | 7.4 | 2 | 2.703 | 0.068 | 14.8 |
| MAPRE1-TM9SF4 | YWHAZ | 335 | 0.049 | 63935 | 18.4 | 2 | 9.094 | 0.027 | 36.8 |
| | FN1 | 23 | 0.134 | 344 | 0.11 | 2 | 105.5 | 4.587 | 0.22 |
| | APP | 9 | 0.286 | 18.5 | 3.6 | 2 | 1.25 | 0.139 | 7.2 |
| | TUBB | 217 | 0.107 | 18784 | 25 | 2 | 4.334 | 0.02 | 50.1 |
| | NTRK1 | 166 | 0.045 | 22877 | 9.51 | 2 | 8.726 | 0.053 | 19 |
| | VCAM1 | 442 | 0 | 2E+05 | 2 | 2 | 110.8 | 0.251 | 3.99 |
| | UNK | 293 | 0.027 | 72709 | 9.82 | 2 | 14.91 | 0.051 | 19.6 |
| | COPS5 | 795 | 0 | 6E+05 | 2 | 2 | 199 | 0.25 | 3.99 |
| | CDK5RAP2 | 53 | 0.06 | 2393 | 5 | 2 | 5.3 | 0.1 | 10 |
| | PRKACA | 128 | 0.043 | 13502 | 7.46 | 2 | 8.583 | 0.067 | 14.9 |
| | AKAP9 | 57 | 0.061 | 2689 | 5.3 | 2 | 5.377 | 0.094 | 10.6 |
| | PRKACB | 58 | 0.099 | 2255 | 7.53 | 2 | 3.854 | 0.066 | 15.1 |
| | CLIP1 | 24 | 0.098 | 459.2 | 4.08 | 2 | 2.941 | 0.123 | 8.16 |
| | TUBB | 217 | 0.107 | 18784 | 25 | 2 | 4.334 | 0.02 | 50.1 |
| | TUBA1A | 208 | 0.057 | 28221 | 13.7 | 2 | 7.605 | 0.037 | 27.4 |
| | HDAC6 | 145 | 0.089 | 12651 | 18.7 | 2 | 3.869 | 0.027 | 37.5 |
| | PDE4DIP | 75 | 0.044 | 4735 | 5.18 | 2 | 7.234 | 0.096 | 10.4 |
| | PRKACA | 128 | 0.043 | 13502 | 7.46 | 2 | 8.583 | 0.067 | 14.9 |

| | | | | | | | | | |
|---|---|---|---|---|---|---|---|---|---|
| | PRKACB | 58 | 0.099 | 2255 | 7.53 | 2 | 3.854 | 0.066 | 15.1 |
| | CDK5RAP2 | 53 | 0.06 | 2393 | 5 | 2 | 5.3 | 0.1 | 10 |
| | AKAP9 | 57 | 0.061 | 2689 | 5.3 | 2 | 5.377 | 0.094 | 10.6 |
| | TERF1 | 307 | 0.026 | 71236 | 9.75 | 2 | 15.75 | 0.051 | 19.5 |
| | SPTAN1 | 149 | 0.089 | 13292 | 15 | 2 | 4.962 | 0.033 | 30 |
| | DST | 54 | 0.092 | 2238 | 6.73 | 2 | 4.014 | 0.074 | 13.5 |
| | MAPRE1 | 156 | 0.07 | 16022 | 12.7 | 2 | 6.158 | 0.039 | 25.3 |
| NUMB-ALDH6A1 | TP53 | 961 | 0 | 9E+05 | 2 | 2 | 240.5 | 0.25 | 4 |
| | NUMB | 38 | 0.132 | 993.1 | 6.72 | 2 | 2.828 | 0.074 | 13.4 |
| | ITCH | 164 | 0.057 | 18214 | 11.2 | 2 | 7.348 | 0.045 | 22.3 |
| | MDM2 | 189 | 0.06 | 22877 | 9.51 | 2 | 9.935 | 0.053 | 19 |
| | EGFR | 833 | 0 | 7E+05 | 52 | 2 | 8.01 | 0.01 | 104 |
| | EPS15 | 131 | 0.097 | 9575 | 14.4 | 2 | 4.535 | 0.035 | 28.9 |
| | EGFR | 833 | 0 | 7E+05 | 52 | 2 | 8.01 | 0.01 | 104 |
| | AP2A1 | 85 | 0.179 | 2960 | 16.9 | 2 | 2.521 | 0.03 | 33.7 |
| | NUMB | 38 | 0.132 | 993.1 | 6.72 | 2 | 2.828 | 0.074 | 13.4 |
| | PRKCZ | 79 | 0.085 | 4314 | 8.41 | 2 | 4.7 | 0.059 | 16.8 |
| | NUMB | 38 | 0.132 | 993.1 | 6.72 | 2 | 2.828 | 0.074 | 13.4 |
| | APP | 9 | 0.286 | 18.5 | 3.6 | 2 | 1.25 | 0.139 | 7.2 |
| | EGFR | 833 | 0 | 7E+05 | 52 | 2 | 8.01 | 0.01 | 104 |
| PARD6B-CD48 | PRKCI | 69 | 0.09 | 3320 | 7.97 | 2 | 4.328 | 0.063 | 15.9 |
| | RASSF8 | 44 | 0.081 | 1526 | 5.38 | 2 | 4.091 | 0.093 | 10.8 |
| | PARD3 | 60 | 0.107 | 2489 | 8.2 | 2 | 3.66 | 0.061 | 16.4 |
| | PARD6G | 17 | 0.213 | 168.2 | 5.11 | 2 | 1.663 | 0.098 | 10.2 |
| | APP | 9 | 0.286 | 18.5 | 3.6 | 2 | 1.25 | 0.139 | 7.2 |
| | PARD6B | 75 | 0.033 | 5068 | 4.4 | 2 | 8.532 | 0.114 | 8.79 |
| | PARD6A | 55 | 0.067 | 2433 | 5.54 | 2 | 4.967 | 0.09 | 11.1 |
| | YWHAH | 180 | 0.063 | 18761 | 13.2 | 2 | 6.807 | 0.038 | 26.4 |
| | PRKCZ | 79 | 0.085 | 4314 | 8.41 | 2 | 4.7 | 0.059 | 16.8 |
| | WWC1 | 37 | 0.161 | 794.4 | 7.58 | 2 | 2.441 | 0.066 | 15.2 |
| | PRKCI | 69 | 0.09 | 3320 | 7.97 | 2 | 4.328 | 0.063 | 15.9 |
| | PARD3 | 60 | 0.107 | 2489 | 8.2 | 2 | 3.66 | 0.061 | 16.4 |
| | PARD6G | 17 | 0.213 | 168.2 | 5.11 | 2 | 1.663 | 0.098 | 10.2 |
| | APP | 9 | 0.286 | 18.5 | 3.6 | 2 | 1.25 | 0.139 | 7.2 |
| | PARD6B | 75 | 0.033 | 5068 | 4.4 | 2 | 8.532 | 0.114 | 8.79 |
| | PARD6A | 55 | 0.067 | 2433 | 5.54 | 2 | 4.967 | 0.09 | 11.1 |
| | YWHAH | 180 | 0.063 | 18761 | 13.2 | 2 | 6.807 | 0.038 | 26.4 |
| | PRKCZ | 79 | 0.085 | 4314 | 8.41 | 2 | 4.7 | 0.059 | 16.8 |
| | RAC1 | 152 | 0.025 | 20797 | 5.71 | 2 | 13.31 | 0.088 | 11.4 |
| | PARD6G | 17 | 0.213 | 168.2 | 5.11 | 2 | 1.663 | 0.098 | 10.2 |
| | PARD6B | 75 | 0.033 | 5068 | 4.4 | 2 | 8.532 | 0.114 | 8.79 |
| | PARD6A | 55 | 0.067 | 2433 | 5.54 | 2 | 4.967 | 0.09 | 11.1 |
| PPP1R12A-MGAT4C | KDM1A | 210 | 0.054 | 31401 | 3.24 | 2 | 32.43 | 0.154 | 6.48 |
| | ELAVL1 | 213 | 0.076 | 28937 | 18.7 | 2 | 5.706 | 0.027 | 37.3 |
| | RPA1 | 463 | 0 | 2E+05 | 2 | 2 | 116 | 0.251 | 3.99 |
| | RPA2 | 226 | 0.06 | 22557 | 9.51 | 2 | 11.88 | 0.053 | 19 |
| | PPP1R12A | 64 | 0.145 | 2469 | 11 | 2 | 2.913 | 0.046 | 22 |

| | | | | | | | | | |
|---|---|---|---|---|---|---|---|---|---|
| | CUL1 | 670 | 0 | 4E+05 | 12 | 2 | 27.92 | 0.042 | 24 |
| | KDM1A | 210 | 0.054 | 31401 | 3.24 | 2 | 32.43 | 0.154 | 6.48 |
| | NUDT5 | 16 | 0.133 | 191.3 | 3.77 | 2 | 2.125 | 0.133 | 7.53 |
| | TP53 | 961 | 0 | 9E+05 | 2 | 2 | 240.5 | 0.25 | 4 |
| | ELAVL1 | 213 | 0.076 | 28937 | 18.7 | 2 | 5.706 | 0.027 | 37.3 |
| | PUS1 | 35 | 0.081 | 1017 | 4.61 | 2 | 3.795 | 0.108 | 9.22 |
| | NUAK1 | 23 | 0.158 | 338.6 | 5.25 | 2 | 2.19 | 0.095 | 10.5 |
| | AARSD1 | 47 | 0.131 | 1436 | 7.88 | 2 | 2.984 | 0.063 | 15.8 |
| | RPA1 | 463 | 0 | 2E+05 | 2 | 2 | 116 | 0.251 | 3.99 |
| | NTRK1 | 166 | 0.045 | 22877 | 9.51 | 2 | 8.726 | 0.053 | 19 |
| | RPA2 | 226 | 0.06 | 22557 | 9.51 | 2 | 11.88 | 0.053 | 19 |
| | RPRD1B | 68 | 0.14 | 2893 | 11.2 | 2 | 3.023 | 0.044 | 22.5 |
| | PPP1R12A | 64 | 0.145 | 2469 | 11 | 2 | 2.913 | 0.046 | 22 |
| | TRIM47 | 15 | 0.124 | 172.3 | 3.5 | 2 | 2.143 | 0.143 | 7 |
| | PAXIP1 | 257 | 0.03 | 53566 | 9.67 | 2 | 13.28 | 0.052 | 19.3 |
| | ACTR3 | 80 | 0.186 | 2461 | 16.5 | 2 | 2.425 | 0.03 | 33 |
| | CUL1 | 670 | 0 | 4E+05 | 12 | 2 | 27.92 | 0.042 | 24 |
| RNF11-C8A | CBLB | 44 | 0.083 | 1465 | 6.36 | 2 | 3.457 | 0.079 | 12.7 |
| | RNF11 | 92 | 0.087 | 5248 | 9.7 | 2 | 4.744 | 0.052 | 19.4 |
| | ITCH | 164 | 0.057 | 18214 | 11.2 | 2 | 7.348 | 0.045 | 22.3 |
| | SMAD4 | 103 | 0.141 | 4686 | 2.01 | 2 | 25.62 | 0.249 | 4.02 |
| | EPN1 | 63 | 0.097 | 2787 | 7.88 | 2 | 4 | 0.063 | 15.8 |
| | RABGEF1 | 36 | 0.061 | 1026 | 3.94 | 2 | 4.564 | 0.127 | 7.89 |
| | UBE2E1 | 114 | 0.026 | 11458 | 4.83 | 2 | 11.81 | 0.104 | 9.65 |
| | UBE2D3 | 203 | 0.032 | 31872 | 8.34 | 2 | 12.18 | 0.06 | 16.7 |
| | UBE2E3 | 88 | 0.037 | 6296 | 5.14 | 2 | 8.567 | 0.097 | 10.3 |
| | HGS | 200 | 0.035 | 31840 | 8.88 | 2 | 11.27 | 0.056 | 17.8 |
| | GGA1 | 50 | 0.078 | 1682 | 5.64 | 2 | 4.433 | 0.089 | 11.3 |
| | AKT1 | 60 | 0.16 | 1929 | 14.2 | 2 | 2.106 | 0.035 | 28.5 |
| | GGA3 | 33 | 0.138 | 605.6 | 6.24 | 2 | 2.646 | 0.08 | 12.5 |
| | GGA2 | 33 | 0.106 | 674.2 | 5.24 | 2 | 3.152 | 0.096 | 10.5 |
| | AP2A1 | 85 | 0.179 | 2960 | 16.9 | 2 | 2.521 | 0.03 | 33.7 |
| | EPN3 | 28 | 0.037 | 701.7 | 2.9 | 2 | 4.833 | 0.173 | 5.79 |
| | UBE2D1 | 244 | 0.036 | 38557 | 10.6 | 2 | 11.54 | 0.047 | 21.1 |
| | AP2B1 | 94 | 0.119 | 5309 | 12.9 | 2 | 3.636 | 0.039 | 25.9 |
| | CSNK2A1 | 382 | 0 | 1E+05 | 2 | 2 | 95.74 | 0.251 | 3.99 |
| | SMURF1 | 252 | 0.058 | 42527 | 16.4 | 2 | 7.662 | 0.03 | 32.9 |
| | EPS15 | 131 | 0.097 | 9575 | 14.4 | 2 | 4.535 | 0.035 | 28.9 |
| | SMURF2 | 86 | 0.071 | 5165 | 7.84 | 2 | 5.487 | 0.064 | 15.7 |
| | UBQLN2 | 79 | 0.098 | 3762 | 9.44 | 2 | 4.183 | 0.053 | 18.9 |
| | STAM2 | 60 | 0.086 | 2578 | 6.98 | 2 | 4.296 | 0.072 | 14 |
| | NEDD4 | 282 | 0.02 | 69819 | 7.51 | 2 | 18.77 | 0.067 | 15 |
| | UBQLN4 | 185 | 0.019 | 29750 | 5.51 | 2 | 16.78 | 0.091 | 11 |
| | NEDD4L | 174 | 0.021 | 26905 | 5.6 | 2 | 15.54 | 0.089 | 11.2 |
| | APP | 9 | 0.286 | 18.5 | 3.6 | 2 | 1.25 | 0.139 | 7.2 |
| | RNF11 | 92 | 0.087 | 5248 | 9.7 | 2 | 4.744 | 0.052 | 19.4 |
| | PSMD4 | 158 | 0.136 | 12680 | 23.2 | 2 | 3.411 | 0.022 | 46.3 |
| | PSMD7 | 104 | 0.233 | 4497 | 25.8 | 2 | 2.016 | 0.019 | 51.6 |

| | | | | | | | | | |
|---|---|---|---|---|---|---|---|---|---|
| | PSMD6 | 86 | 0.327 | 2225 | 29.5 | 2 | 1.459 | 0.017 | 58.9 |
| | PSMD11 | 129 | 0.18 | 7645 | 24.8 | 2 | 2.596 | 0.02 | 49.7 |
| | PSMD10 | 64 | 0.214 | 2368 | 15.2 | 2 | 2.101 | 0.033 | 30.5 |
| | PSMD3 | 121 | 0.214 | 5219 | 27.5 | 2 | 2.201 | 0.018 | 55 |
| | PSMD12 | 92 | 0.324 | 1981 | 30.8 | 2 | 1.493 | 0.016 | 61.6 |
| | PSMD13 | 89 | 0.318 | 2172 | 29.7 | 2 | 1.499 | 0.017 | 59.4 |
| | USP14 | 66 | 0.23 | 2222 | 16.7 | 2 | 1.978 | 0.03 | 33.4 |
| | PSMD14 | 106 | 0.246 | 4362 | 27.5 | 2 | 1.925 | 0.018 | 55.1 |
| | PSMD1 | 117 | 0.249 | 3404 | 30.6 | 2 | 1.91 | 0.016 | 61.3 |
| | PSMD2 | 161 | 0.148 | 11360 | 25.6 | 2 | 3.15 | 0.02 | 51.1 |
| | APP | 9 | 0.286 | 18.5 | 3.6 | 2 | 1.25 | 0.139 | 7.2 |
| | GGA1 | 50 | 0.078 | 1682 | 5.64 | 2 | 4.433 | 0.089 | 11.3 |
| | GGA3 | 33 | 0.138 | 605.6 | 6.24 | 2 | 2.646 | 0.08 | 12.5 |
| | GGA2 | 33 | 0.106 | 674.2 | 5.24 | 2 | 3.152 | 0.096 | 10.5 |
| | APP | 9 | 0.286 | 18.5 | 3.6 | 2 | 1.25 | 0.139 | 7.2 |
| | AKT1 | 60 | 0.16 | 1929 | 14.2 | 2 | 2.106 | 0.035 | 28.5 |
| | RNF11 | 92 | 0.087 | 5248 | 9.7 | 2 | 4.744 | 0.052 | 19.4 |
| | TBK1 | 111 | 0.085 | 7850 | 11.2 | 2 | 4.968 | 0.045 | 22.3 |
| | APP | 9 | 0.286 | 18.5 | 3.6 | 2 | 1.25 | 0.139 | 7.2 |
| | NEDD4 | 282 | 0.02 | 69819 | 7.51 | 2 | 18.77 | 0.067 | 15 |
| SIPA1L3-WDR62 | MAPK10 | 43 | 0.085 | 1338 | 5.35 | 2 | 4.019 | 0.093 | 10.7 |
| | WDR62 | 35 | 0.077 | 944.6 | 4.4 | 2 | 3.977 | 0.114 | 8.8 |
| | MAPK8 | 172 | 0.045 | 22877 | 9.51 | 2 | 9.041 | 0.053 | 19 |
| | MAPK9 | 97 | 0.041 | 7629 | 5.81 | 2 | 8.342 | 0.086 | 11.6 |
| | SFN | 250 | 0.053 | 43097 | 15 | 2 | 8.356 | 0.033 | 29.9 |
| | YWHAB | 311 | 0.036 | 61018 | 29.3 | 2 | 5.302 | 0.017 | 58.7 |
| | SIPA1L3 | 22 | 0.216 | 305 | 6.26 | 2 | 1.757 | 0.08 | 12.5 |
| | YWHAQ | 180 | 0.063 | 18761 | 13.2 | 2 | 6.807 | 0.038 | 26.4 |
| | YWHAB | 311 | 0.036 | 61018 | 29.3 | 2 | 5.302 | 0.017 | 58.7 |
| | FBXW11 | 76 | 0.12 | 3995 | 9.98 | 2 | 3.809 | 0.05 | 20 |
| | MAPK10 | 43 | 0.085 | 1338 | 5.35 | 2 | 4.019 | 0.093 | 10.7 |
| | ELAVL1 | 213 | 0.076 | 28937 | 18.7 | 2 | 5.706 | 0.027 | 37.3 |
| | WDR62 | 35 | 0.077 | 944.6 | 4.4 | 2 | 3.977 | 0.114 | 8.8 |
| | TBP | 156 | 0.071 | 15459 | 12.9 | 2 | 6.066 | 0.039 | 25.7 |
| | MAPK8 | 172 | 0.045 | 22877 | 9.51 | 2 | 9.041 | 0.053 | 19 |
| | MAPK9 | 97 | 0.041 | 7629 | 5.81 | 2 | 8.342 | 0.086 | 11.6 |
| SLC26A6-PRKAR2A | AKAP7 | 9 | 0.25 | 50 | 3.6 | 2 | 1.25 | 0.139 | 7.2 |
| | AKAP9 | 57 | 0.061 | 2689 | 5.3 | 2 | 5.377 | 0.094 | 10.6 |
| | PRKAR2A | 48 | 0.091 | 1644 | 6.16 | 2 | 3.894 | 0.081 | 12.3 |
| | PRKAR2B | 50 | 0.068 | 2013 | 5.22 | 2 | 4.793 | 0.096 | 10.4 |
| | PRKACA | 128 | 0.043 | 13502 | 7.46 | 2 | 8.583 | 0.067 | 14.9 |
| | PRKACB | 58 | 0.099 | 2255 | 7.53 | 2 | 3.854 | 0.066 | 15.1 |
| | PRKAR2A | 48 | 0.091 | 1644 | 6.16 | 2 | 3.894 | 0.081 | 12.3 |
| | AKAP7 | 9 | 0.25 | 50 | 3.6 | 2 | 1.25 | 0.139 | 7.2 |
| | PRKACA | 128 | 0.043 | 13502 | 7.46 | 2 | 8.583 | 0.067 | 14.9 |
| | PRKACB | 58 | 0.099 | 2255 | 7.53 | 2 | 3.854 | 0.066 | 15.1 |
| | PRKAR2B | 50 | 0.068 | 2013 | 5.22 | 2 | 4.793 | 0.096 | 10.4 |

| | | | | | | | | | |
|---|---|---|---|---|---|---|---|---|---|
| | GCH1 | 30 | 0.02 | 794 | 2.47 | 2 | 6.08 | 0.203 | 4.93 |
| ST14-APLP2 | BRCA1 | 17 | 0.191 | 187.2 | 0.09 | 2 | 94.44 | 5.556 | 0.18 |
| | APLP2 | 22 | 0.056 | 422.3 | 3.04 | 2 | 3.615 | 0.164 | 6.09 |
| | ETS1 | 52 | 0.149 | 1498 | 9.4 | 2 | 2.767 | 0.053 | 18.8 |
| | JUN | 211 | 0.072 | 25977 | 17.1 | 2 | 6.184 | 0.029 | 34.1 |
| | SFN | 250 | 0.053 | 43097 | 15 | 2 | 8.356 | 0.033 | 29.9 |
| | HDAC5 | 339 | 0.061 | 59605 | 22.5 | 2 | 7.535 | 0.022 | 45 |
| | APLP2 | 22 | 0.056 | 422.3 | 3.04 | 2 | 3.615 | 0.164 | 6.09 |
| | RPL26 | 92 | 0.425 | 1353 | 40.2 | 2 | 1.144 | 0.012 | 80.4 |
| | BRCA1 | 17 | 0.191 | 187.2 | 0.09 | 2 | 94.44 | 5.556 | 0.18 |
| | JUNB | 41 | 0.113 | 1175 | 6.38 | 2 | 3.213 | 0.078 | 12.8 |
| | JUN | 211 | 0.072 | 25977 | 17.1 | 2 | 6.184 | 0.029 | 34.1 |
| | APBB1 | 81 | 0.072 | 4892 | 7.58 | 2 | 5.343 | 0.066 | 15.2 |
| | APBB2 | 27 | 0.06 | 617.7 | 3.43 | 2 | 3.937 | 0.146 | 6.86 |
| | KAT5 | 210 | 0.054 | 31401 | 3.24 | 2 | 32.43 | 0.154 | 6.48 |
| | ETS1 | 52 | 0.149 | 1498 | 9.4 | 2 | 2.767 | 0.053 | 18.8 |
| | APLP2 | 22 | 0.056 | 422.3 | 3.04 | 2 | 3.615 | 0.164 | 6.09 |
| | MAPK8 | 172 | 0.045 | 22877 | 9.51 | 2 | 9.041 | 0.053 | 19 |
| STRADB-NOP58 | NIFK | 140 | 0.141 | 8754 | 21.5 | 2 | 3.255 | 0.023 | 43 |
| | NOP56 | 249 | 0.139 | 22329 | 36.4 | 2 | 3.419 | 0.014 | 72.8 |
| | HNRNPU | 503 | 0 | 3E+05 | 2 | 2 | 126 | 0.251 | 3.99 |
| | RUVBL2 | 223 | 0.085 | 25036 | 20.6 | 2 | 5.408 | 0.024 | 41.2 |
| | NOLC1 | 97 | 0.142 | 4793 | 15.5 | 2 | 3.131 | 0.032 | 31 |
| | SNU13 | 164 | 0.138 | 13747 | 24.1 | 2 | 3.398 | 0.021 | 48.3 |
| | NTRK1 | 166 | 0.045 | 22877 | 9.51 | 2 | 8.726 | 0.053 | 19 |
| | PUM3 | 65 | 0.358 | 586.7 | 24.5 | 2 | 1.326 | 0.02 | 49 |
| | RPS15A | 168 | 0.371 | 3578 | 63.6 | 2 | 1.32 | 0.008 | 127 |
| | RSL1D1 | 57 | 0.299 | 838.1 | 18.4 | 2 | 1.545 | 0.027 | 36.9 |
| | KRR1 | 63 | 0.283 | 1087 | 19.3 | 2 | 1.636 | 0.026 | 38.5 |
| | DDX18 | 98 | 0.273 | 1704 | 28.2 | 2 | 1.737 | 0.018 | 56.4 |
| | EIF6 | 142 | 0.101 | 11875 | 16.1 | 2 | 4.407 | 0.031 | 32.2 |
| | DHX15 | 82 | 0.238 | 2197 | 21.1 | 2 | 1.947 | 0.024 | 42.1 |
| | RPS4X | 200 | 0.329 | 4491 | 67.2 | 2 | 1.488 | 0.007 | 134 |
| | EED | 66 | 0.322 | 479.7 | 0.17 | 2 | 197.6 | 2.994 | 0.33 |
| | PRPF3 | 96 | 0.168 | 3879 | 17.6 | 2 | 2.727 | 0.028 | 35.2 |
| | TARDBP | 299 | 0.124 | 26560 | 38.7 | 2 | 3.859 | 0.013 | 77.5 |
| | RPL11 | 175 | 0.355 | 5479 | 63.5 | 2 | 1.378 | 0.008 | 127 |
| | EIF2S2 | 65 | 0.101 | 2804 | 8.33 | 2 | 3.9 | 0.06 | 16.7 |
| | DDX27 | 58 | 0.304 | 817.8 | 19 | 2 | 1.525 | 0.026 | 38 |
| | NOP58 | 101 | 0.262 | 2629 | 27.9 | 2 | 1.809 | 0.018 | 55.8 |
| | FN1 | 23 | 0.134 | 344 | 0.11 | 2 | 105.5 | 4.587 | 0.22 |
| | DDX24 | 115 | 0.177 | 5582 | 21.9 | 2 | 2.62 | 0.023 | 43.9 |
| | U2AF1 | 114 | 0.212 | 3674 | 25.8 | 2 | 2.213 | 0.019 | 51.5 |
| | RPL30 | 131 | 0.445 | 3288 | 59.3 | 2 | 1.104 | 0.008 | 119 |
| | ESR1 | 23 | 0.134 | 344 | 0.41 | 2 | 28.12 | 1.222 | 0.82 |
| | DDX56 | 82 | 0.238 | 2197 | 21.1 | 2 | 1.947 | 0.024 | 42.1 |
| | FTSJ3 | 99 | 0.219 | 2785 | 23.2 | 2 | 2.13 | 0.022 | 46.5 |
| | DDX47 | 46 | 0.144 | 1362 | 8.3 | 2 | 2.772 | 0.06 | 16.6 |

| | | | | | | | | | |
|---|---|---|---|---|---|---|---|---|---|
| | KPNA6 | 55 | 0.098 | 2044 | 7.18 | 2 | 3.831 | 0.07 | 14.4 |
| | GTPBP4 | 87 | 0.244 | 2174 | 22.7 | 2 | 1.916 | 0.022 | 45.4 |
| | WDR36 | 57 | 0.204 | 1484 | 13.2 | 2 | 2.158 | 0.038 | 26.4 |
| | KPNA1 | 101 | 0.085 | 6590 | 10.4 | 2 | 4.85 | 0.048 | 20.8 |
| | DKC1 | 98 | 0.177 | 3896 | 18.8 | 2 | 2.607 | 0.027 | 37.6 |
| | RRP12 | 60 | 0.248 | 1097 | 16.4 | 2 | 1.834 | 0.031 | 32.7 |
| | BOP1 | 48 | 0.165 | 1324 | 9.38 | 2 | 2.56 | 0.053 | 18.8 |
| | FBL | 227 | 0.116 | 24046 | 28.1 | 2 | 4.035 | 0.018 | 56.3 |
| | TBL3 | 47 | 0.254 | 1044 | 13.4 | 2 | 1.752 | 0.037 | 26.8 |
| | WDR36 | 57 | 0.204 | 1484 | 13.2 | 2 | 2.158 | 0.038 | 26.4 |
| | NOP58 | 101 | 0.262 | 2629 | 27.9 | 2 | 1.809 | 0.018 | 55.8 |
| | DHX15 | 82 | 0.238 | 2197 | 21.1 | 2 | 1.947 | 0.024 | 42.1 |
| | NOP56 | 249 | 0.139 | 22329 | 36.4 | 2 | 3.419 | 0.014 | 72.8 |
| STX16-RAE1 | FBXW11 | 76 | 0.12 | 3995 | 9.98 | 2 | 3.809 | 0.05 | 20 |
| | NXF1 | 61 | 0.312 | 489.7 | 24.5 | 2 | 1.244 | 0.02 | 49 |
| | FAF1 | 124 | 0.075 | 9914 | 11 | 2 | 5.636 | 0.045 | 22 |
| | RAE1 | 120 | 0.064 | 10495 | 9.52 | 2 | 6.302 | 0.053 | 19 |
| | ILF3 | 65 | 0.109 | 2629 | 18.8 | 2 | 1.727 | 0.027 | 37.6 |
| | CUL1 | 670 | 0 | 4E+05 | 12 | 2 | 27.92 | 0.042 | 24 |
| | HNRNPUL1 | 112 | 0.14 | 6164 | 17.2 | 2 | 3.25 | 0.029 | 34.5 |
| | CUL3 | 70 | 0.081 | 3677 | 10.4 | 2 | 3.375 | 0.048 | 20.7 |
| | NXF1 | 61 | 0.312 | 489.7 | 24.5 | 2 | 1.244 | 0.02 | 49 |
| | CUL1 | 670 | 0 | 4E+05 | 12 | 2 | 27.92 | 0.042 | 24 |
| | ILF3 | 65 | 0.109 | 2629 | 18.8 | 2 | 1.727 | 0.027 | 37.6 |
| | CUL3 | 70 | 0.081 | 3677 | 10.4 | 2 | 3.375 | 0.048 | 20.7 |
| TANC2-CHD6 | ZFYVE9 | 39 | 0.073 | 1245 | 4.65 | 2 | 4.194 | 0.108 | 9.3 |
| | PPP1CC | 274 | 0.036 | 53066 | 11.8 | 2 | 11.62 | 0.042 | 23.6 |
| | PPP1CA | 285 | 0.045 | 53592 | 14.8 | 2 | 9.653 | 0.034 | 29.5 |
| | TANC2 | 8 | 0.107 | 47 | 2.44 | 2 | 1.637 | 0.205 | 4.89 |
| | ZFYVE9 | 39 | 0.073 | 1245 | 4.65 | 2 | 4.194 | 0.108 | 9.3 |
| | PPP1CC | 274 | 0.036 | 53066 | 11.8 | 2 | 11.62 | 0.042 | 23.6 |
| | PPP1CA | 285 | 0.045 | 53592 | 14.8 | 2 | 9.653 | 0.034 | 29.5 |
| | TANC2 | 8 | 0.107 | 47 | 2.44 | 2 | 1.637 | 0.205 | 4.89 |
| | MAEA | 18 | 0.275 | 139.1 | 6.32 | 2 | 1.425 | 0.079 | 12.6 |
| | RANBP9 | 111 | 0.059 | 9201 | 8.31 | 2 | 6.682 | 0.06 | 16.6 |
| | MKLN1 | 25 | 0.196 | 297.8 | 6.24 | 2 | 2.003 | 0.08 | 12.5 |
| | MMP7 | 23 | 0.206 | 358.9 | 6.25 | 2 | 1.84 | 0.08 | 12.5 |
| | RMND5A | 33 | 0.184 | 645.3 | 7.65 | 2 | 2.158 | 0.065 | 15.3 |
| | MAEA | 18 | 0.275 | 139.1 | 6.32 | 2 | 1.425 | 0.079 | 12.6 |
| | RANBP9 | 111 | 0.059 | 9201 | 8.31 | 2 | 6.682 | 0.06 | 16.6 |
| | MKLN1 | 25 | 0.196 | 297.8 | 6.24 | 2 | 2.003 | 0.08 | 12.5 |
| | RANBP10 | 25 | 0.093 | 488.5 | 4.08 | 2 | 3.066 | 0.123 | 8.15 |
| | MMP7 | 23 | 0.206 | 358.9 | 6.25 | 2 | 1.84 | 0.08 | 12.5 |
| | RMND5A | 33 | 0.184 | 645.3 | 7.65 | 2 | 2.158 | 0.065 | 15.3 |
| TMPRSS2-ERG | CDC5L | 580 | 0 | 3E+05 | 2 | 2 | 145.2 | 0.25 | 3.99 |
| | DDX3X | 146 | 0.144 | 8758 | 22.7 | 2 | 3.219 | 0.022 | 45.4 |
| | ELAVL1 | 213 | 0.076 | 28937 | 18.7 | 2 | 5.706 | 0.027 | 37.3 |
| | CAD | 100 | 0.182 | 3220 | 19.9 | 2 | 2.517 | 0.025 | 39.7 |

| | | | | | | | | | |
|---|---|---|---|---|---|---|---|---|---|
| | NEDD4 | 282 | 0.02 | 69819 | 7.51 | 2 | 18.77 | 0.067 | 15 |
| | SF3B1 | 177 | 0.154 | 11161 | 28.9 | 2 | 3.067 | 0.017 | 57.7 |
| | PARP1 | 219 | 0.08 | 25770 | 1.23 | 2 | 88.81 | 0.406 | 2.47 |
| | PRKDC | 234 | 0.084 | 28338 | 21.4 | 2 | 5.478 | 0.023 | 42.7 |
| | PRPF8 | 153 | 0.177 | 7356 | 28.5 | 2 | 2.684 | 0.018 | 57 |
| | SFPQ | 162 | 0.14 | 10234 | 24.2 | 2 | 3.346 | 0.021 | 48.4 |
| | ERG | 79 | 0.147 | 2628 | 13.3 | 2 | 2.976 | 0.038 | 26.6 |
| | POLR2A | 251 | 0.074 | 34919 | 1.24 | 2 | 101.3 | 0.404 | 2.48 |
| | TOP1 | 146 | 0.154 | 8689 | 24.2 | 2 | 3.021 | 0.021 | 48.3 |
| | CLTC | 277 | 0.066 | 45931 | 20.1 | 2 | 6.885 | 0.025 | 40.2 |
| | SF3B2 | 26 | 0.142 | 450.3 | 7.33 | 2 | 1.773 | 0.068 | 14.7 |
| | XRCC5 | 264 | 0.064 | 41296 | 0.87 | 2 | 152.1 | 0.576 | 1.74 |
| | XRCC6 | 264 | 0.064 | 41296 | 0.57 | 2 | 232.4 | 0.88 | 1.14 |
| | DDX23 | 69 | 0.168 | 2296 | 13.2 | 2 | 2.608 | 0.038 | 26.5 |
| | SNRNP200 | 152 | 0.168 | 8534 | 27 | 2 | 2.819 | 0.019 | 53.9 |
| | DDX21 | 111 | 0.181 | 4280 | 21.7 | 2 | 2.554 | 0.023 | 43.5 |
| | TUBB | 217 | 0.107 | 18784 | 25 | 2 | 4.334 | 0.02 | 50.1 |
| | NONO | 144 | 0.105 | 10404 | 16.9 | 2 | 4.27 | 0.03 | 33.7 |
| | JUN | 211 | 0.072 | 25977 | 17.1 | 2 | 6.184 | 0.029 | 34.1 |
| | HNRNPU | 503 | 0 | 3E+05 | 2 | 2 | 126 | 0.251 | 3.99 |
| | PRPF40A | 178 | 0.091 | 18728 | 17.9 | 2 | 4.985 | 0.028 | 35.7 |
| | AR | 60 | 0.16 | 1929 | 14.2 | 2 | 2.106 | 0.035 | 28.5 |
| | NCL | 228 | 0.135 | 19658 | 32.3 | 2 | 3.53 | 0.015 | 64.6 |
| | HNRNPM | 240 | 0.157 | 18323 | 39.4 | 2 | 3.043 | 0.013 | 78.9 |
| | HNRNPC | 164 | 0.143 | 10541 | 25.1 | 2 | 3.27 | 0.02 | 50.1 |
| | TOP2B | 62 | 0.185 | 1571 | 13.1 | 2 | 2.37 | 0.038 | 26.2 |
| | ILF3 | 65 | 0.109 | 2629 | 18.8 | 2 | 1.727 | 0.027 | 37.6 |
| | ILF2 | 65 | 0.109 | 2629 | 12.8 | 2 | 2.535 | 0.039 | 25.6 |
| | PRPF8 | 153 | 0.177 | 7356 | 28.5 | 2 | 2.684 | 0.018 | 57 |
| | ERG | 79 | 0.147 | 2628 | 13.3 | 2 | 2.976 | 0.038 | 26.6 |
| | SF3B2 | 26 | 0.142 | 450.3 | 7.33 | 2 | 1.773 | 0.068 | 14.7 |
| | SF3B1 | 177 | 0.154 | 11161 | 28.9 | 2 | 3.067 | 0.017 | 57.7 |
| | PARP1 | 219 | 0.08 | 25770 | 1.23 | 2 | 88.81 | 0.406 | 2.47 |
| | PRKDC | 234 | 0.084 | 28338 | 21.4 | 2 | 5.478 | 0.023 | 42.7 |
